# Flux organizations and control modes in antagonistically combined negative feedback loops

**DOI:** 10.1101/2025.11.09.687434

**Authors:** Peter Ruoff

## Abstract

The purpose of this paper is to show the different compensatory responses and control models towards environmental perturbations when two antagonistic integral controllers are combined at the level of the controlled variable. Dependent on the controllers’ relative setpoints two types of compensatory flux regulations occur, which have been termed delegated and isolated control. In delegated control one of the feedbacks submits its entire compensatory flux to the system, while the other feedback is the actual controller by neutralizing the excess flux of its antagonistic partner. In isolated control the compensatory flux of one of the controllers is negligible while the other feedback regulates the controlled variable alone. One of the findings is that these interacting feedbacks can exhibit environmentally driven setpoint changes known as rheostasis. A striking example is the photoperiodic control of Siberian hamsters’ body weights, which can be rationalized by the interacting feedbacks between circadian morning (M) and evening (E) oscillators. A third control type is metastable control. Here, additions or removals of the controller variable can cause a switch to the antagonistic partner’s control regime, but resets to its original control mode once additions or removals stop. Integral windup can induce temporary metastable setpoint changes or even lead to robust perfect adaptation without integral feedback! How the setpoint in blood glucose homeostasis arises is still debated. A dual-controller approach with two setpoints can describe many properties of blood glucose homeostasis and the roles of insulin, glucagon, and somatostatin.

**Graphical Abstract:** 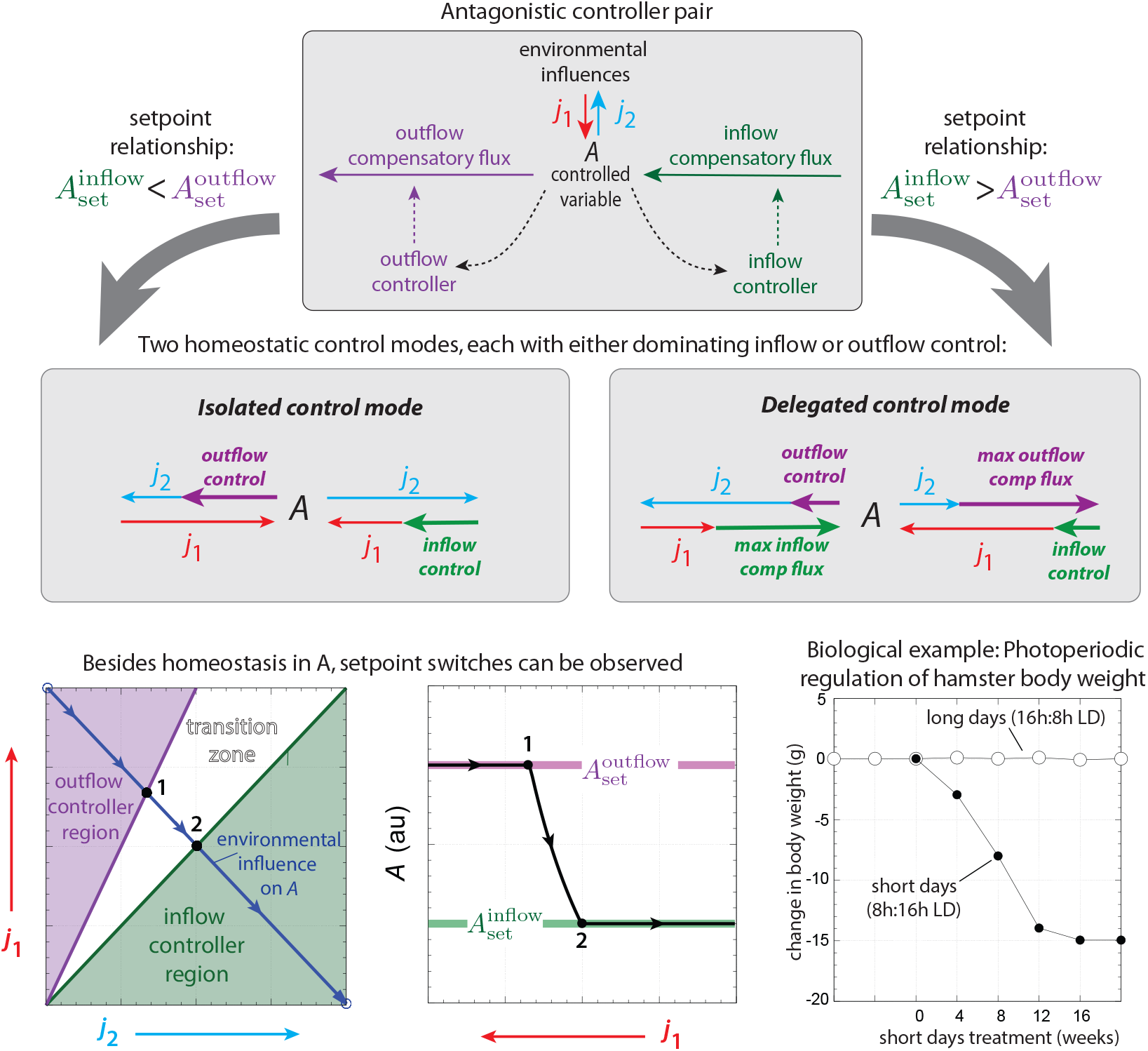

**Highlights:**

- A set of 16 inflow/outflow controllers (negative feedbacks) combined by a common controlled variable have been studied.
- Dependent on the relationship between the inflow/outflow controllers’ setpoints two distinct regulatory modes termed ‘delegated control’ and ‘isolated control’ have been identified.
- Regions of control and their extensions are visualized by a ‘perturbation phase diagram’.
- Metastable control may temporarily occur when one of the controller variables are environmentally increased or decreased.
- Robust perfect or near-perfect adaptation may occur even in the absence of integral feedback.
- The combined controllers can show homeostatic as well as rheostatic behaviors. A striking biological example of rheostasis is found in the photoperiodic regulation of the Siberian hamster’s body weight.
- The origin of the glycemic setpoint in blood glucose homeostasis is still debated. A glucagon- and insulin-based two-setpoint model suggests that in diabetic individuals the upper insulin-dependent setpoint increases with decreasing insulin generation rates. This setpoint is defended irrespective of its value, but the accuracy of regulation depends how tightly the insulin degrading enzyme (IDE, insulysin) binds to insulin. In addition, both glucagon- and insulin-based setpoints depend (rheostatically) on the level of somatostatin.

## 1. Introduction

The term ‘homeostasis’ was introduced by Walter Cannon (Cannon, 1929, 1939; Langley, 1973) to address the ability of organisms to defend themselves against internal and external disturbances. This is done by keeping their physiological steady states within narrow limits (Clancy and McVicar, 2002; Billman, 2020). Wiener’s cybernetics and the findings that negative feedback mechanisms play a major role in homeostatic regulation (Curtis et al., 1958; Hughes, 1964) were the start to describe homeostasis and adaptation mechanisms by combined methods from control engineering and reaction kinetics. The success to describe homeostatic mechanisms by feedback regulation led some researchers to equate homeostasis with single negative feedback regulations and a defined constant setpoint. However, the facets of homeostasis involve more than regulation by a single negative feedback loop. Thus, alternative terminologies were introduced. Selye (1974) coined the term ‘heterostasis’ to point to alternative regulations when a control system faces unusually high demands. The term ‘enantiostasis’ was introduced by Mangum and Towle (1977) to describe a preserved relationship between an environmental influence and a regulated variable (Hochachka and Somero, 1984). Moore-Ede (1986), focussing on the anticipatory adaptation of circadian rhythms, introduced the term ‘predictive homeostasis’ to include regulations by these biochemical oscillators. The term ‘rheostasis’, introduced by Mrosovsky (1990), points to the fact that homeostatic setpoints may change dependent on environmental conditions and may include the regulation by circadian or other biochemical oscillators on a hierarchical basis (Stevenson, 2024).

‘Allostasis’ (Schulkin, 2003, 2004), a term suggested by Sterling and Eyer (1988) and closely connected to the concept of rheostasis, focuses on coordinated changes in the brain by neural signaling pathways, which may include setpoint changes with respect to external conditions, such as environmental stress. Although these aspects of homeostatic regulation, such as changes of setpoints, suggested hierarchical organization (Stevenson, 2024), or the presence of nonlinear dynamics (Lloyd et al., 2001), are clearly important, they still fall within Cannon’s original definition. As argued by Carpenter (2004) and as discussed at the end of the paper with respect to rheostasis, Cannon’s homeostasis definition can still serve as a unifying concept.

Physiological controllers come often in antagonistic pairs and regulate inflow and outflow fluxes to and from a homeostatic controlled compound. Blood glucose regulation by insulin and glucagon is one of many examples. Saunders’ ‘rein control’ model (Saunders et al., 1998, 2000) is an example of how antagonistic compensatory fluxes induced by insulin and glucagon contribute to an overall blood glucose homeostasis.

In the present work I combine integral inflow and outflow feedbacks, such that each feedback has a defined setpoint and contributes to the regulation of the controlled variable. Fig 1 shows how controller combinations are arranged. Unlike hierarchical control arrangements (Thorsen et al., 2018; Nygård and Ruoff, 2023) there is no ‘cross-talk’ between the inflow and out-flow controllers. Compound *A* is the controlled variable which is influenced by the environmental perturbations *k*_1_ and *k*_2_. During homeostasis, but dependent on *k*_1_ and *k*_2_, *A* can be kept at the setpoint of one of the controllers or in-between these setpoints.

**Figure 1.**
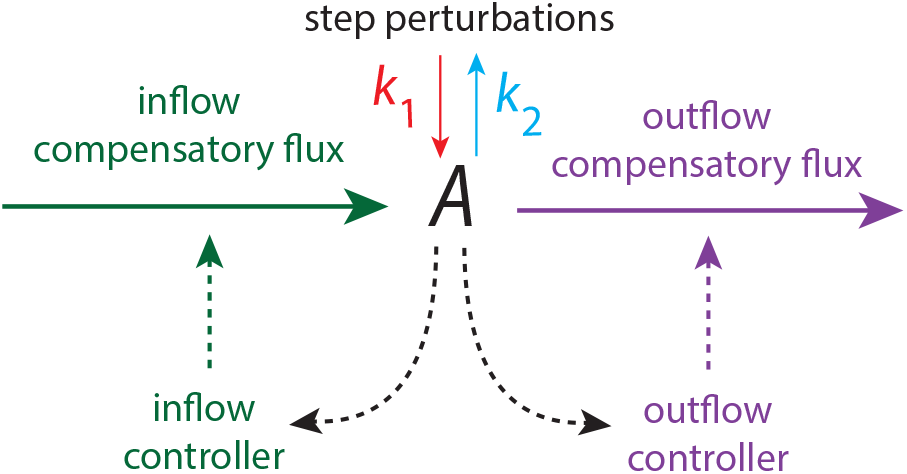
The arrangement of the investigated combinations between inflow and outflow controllers to keep homeostasis in *A*. Dependent on the step perturbations *k*_1_ and *k*_2_ (indicated respectively as red and blue arrows), inflow and/or outflow controllers (outlined in green and violet) will be activated to maintain *A* homeostasis. Dashed lines indicate signaling from *A* to the controllers and the activation of the compensatory fluxes.

The signaling between *A* and the individual controllers is described by saturation kinetics, i.e. by activation or inhibition of a compensatory flux. Thus, control of *A* may break down when perturbations become too large and compensatory fluxes reach their maximum limits (Ang et al., 2010).

## 2. Materials and methods

### 2.1. Computational methods

Computations were performed with the Fortran subroutine LSODE (Radhakrishnan and Hindmarsh, 1993) with plots generated by gnuplot (www.gnuplot.info). Reaction schemes and plot annotations were prepared with Adobe Illustrator (www.adobe.com). Rate constants and other parameters are in arbitrary units (au) and represented by *k*_1_, *k*_2_, *k*_3_,… independent of their kinetic meaning, i.e. whether they are turnover numbers, Michaelis constants, or activation/inhibition constants.

### 2.2. The studied individual controllers (negative feedbacks)

Because many physiological controllers come in antagonistic pairs, I have looked at the combinations between inflow and outflow controllers (Ni et al., 2009; Drengstig et al., 2012a), which act directly on the controlled variable as indicated in Fig 1. Fig 2 shows a set of eight single-loop negative feedback loops (Drengstig et al., 2012a), which have been employed in this study. In these feedbacks integral control is achieved by zero-order kinetic removal of the *E*_*i*_ (‘control of the controller principle’ (Ni et al., 2009); see section 2.5 for further details). The eight feedbacks divide into two equal sets which have been termed inflow and outflow controllers (Fig 2). The controlled variable and its concentration is described as *A*, while the controller variables are the *E*_*i*_’s, where *i* identifies one of the eight controller motifs M*i* with *i* ∈ {1, 2,.., 7, 8}.

**Figure 2.**
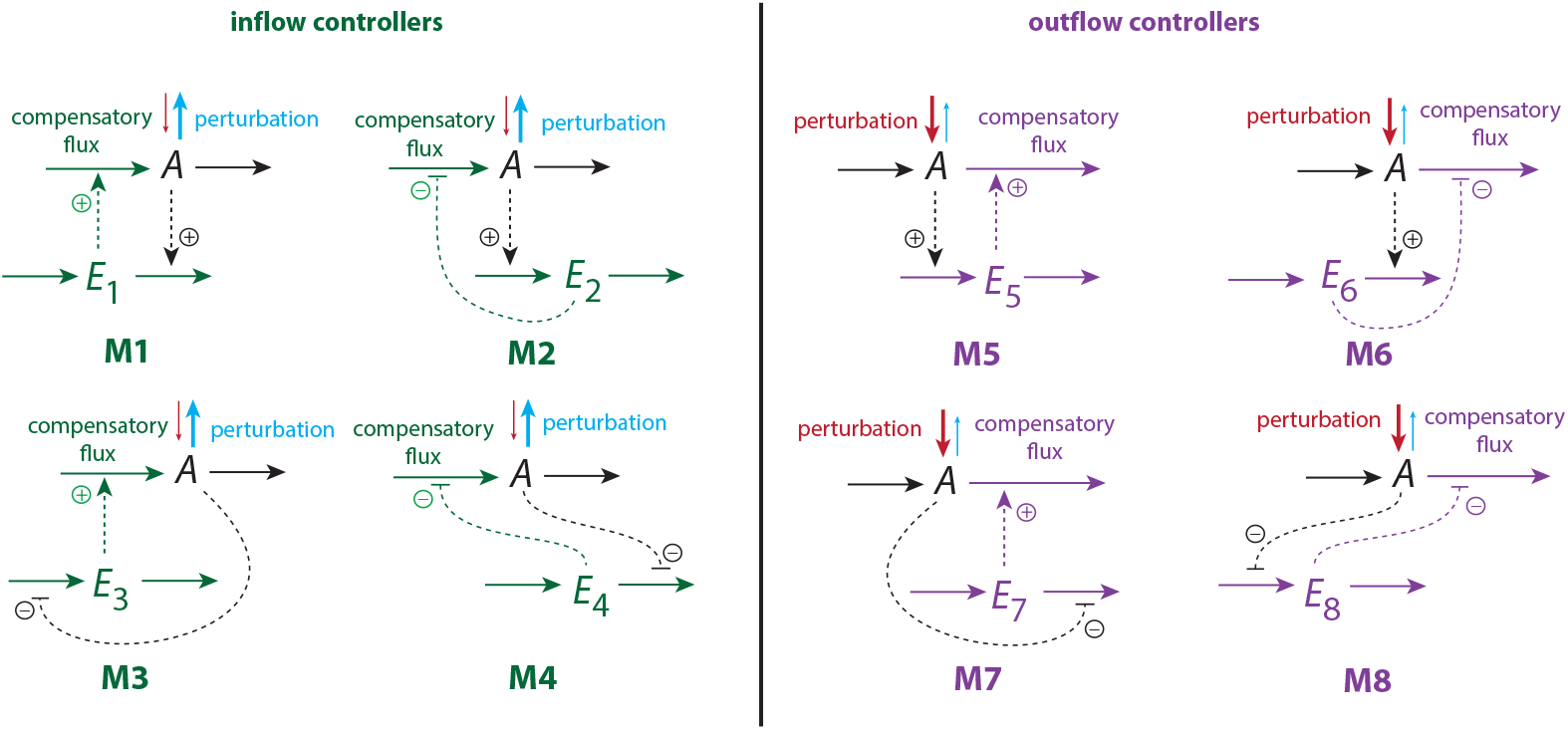
The eight negative feedback loops used in this study. The four inflow controllers M1-M4 are combined with the four outflow controllers M5-M8 which leads to sixteen possible combinations (see below). *A* is the controlled variable while the *E*_*i*_’s (*i* = 1, 2,.., 8) are the controller (manipulated) variables. Dashed lines indicate the signaling between *A* and the *E*_*i*_’s in terms of activations (plus signs) or inhibitions (minus signs). It may be mentioned that a similar negative feedback classification was used by Franck in his 1980 paper on physicochemical oscillators (see Fig 4 in Franck (1980)).

### 2.3. Implementation of inhibition and activation

Inhibitions of a reaction rate *r* by *X* (here X is *A*, or *E*_*k*_ with *k* ∈ {1, 2,.., 7, 8}) is achieved by multiplying *r* with the inhibition factor *f*_*inhib*_

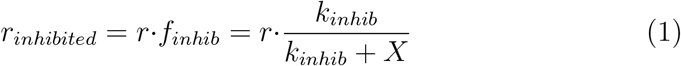

where *k*_*inhib*_ is a inhibition constant. Kinetically this is equivalent to what is known as a special case of ‘mixed inhibition’ (Segel, 1975), where two inhibition constants have the same value and are written as one.

Activations are described by two scenarios. In the first one activation is described in terms of a ‘mixed activation’ (Cornish-Bowden, 2004) by activator *Y* with activation constant *k*_*act*_, i.e.

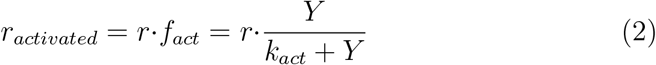

In the other scenario we have first-order kinetics with respect to the activator *Y*, i.e.

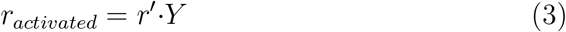

This approach avoids saturation by *Y* in Eq 2 and saturation-related break-downs (Ang and McMillen, 2013).

To compare the effects between these two activation types I have included computations when *activated compensatory fluxes* use either Eq 2 or Eq 3. It may be noted that first-order activation with respect to activator *Y* (Eq 3) represents a special case of Eq 2, i.e. when *k*_*act*_ becomes much larger than *Y* and *r*^*′*^ can be written as *r*^*′*^=*r/k*_*act*_. As *k*_*act*_ is a dissociation constant (the Michaelis constant *K*_*M*_) between the binding of the activator *Y* and an enzyme catalyzing reaction *r* Eq 3 reflects a ‘low-affinity’ activation and regulation. On the other hand, Eq 2 can be considered as a ‘high-affinity’ regulation when *k*_*act*_ is low. The distinction between high-affinity and low-affinity kinetics is well documented for transport proteins, and may also be used to distinguish compensatory fluxes in homeostatic regulations (for an example, see Huang et al. (2011)).

It should be further mentioned that these formulations represent simplifications and that biochemical activations and inhibitions are generally more complex, as they often involve positive or/and negative cooperativities Segel (1975), along possible hysteretic or allosteric behaviors (Frieden, 1970; Farnsworth, 2025). Such complexities are neglected here.

### 2.4. Keeping oscillatory behavior of feedbacks at a minimum

Although oscillations are often observed in biological systems, I decided to focus here on nonoscillatory regulation and keep oscillatory behavior at a minimum. It is well known that negative feedbacks can oscillate (Goodwin, 1963; Franck, 1980; Kholodenko, 2000; Goldbeter et al., 2001; Pigolotti et al., 2007; Jolma et al., 2010), especially when aggressiveness (or loop-gain) of the feedback is increased (Blanchini et al., 2018), or when the removal of the controlled variable becomes zero-order with respect to *A*. In the latter case the average of the controlled variable can show robust perfect adaptation when integral control is present (Thorsen et al., 2014). Since first-order removals with respect to the controlled variable *A* can effectively quench oscillatory behavior of a negative feedback (Thorsen et al., 2014) (but not always ovoid it) all *A* removals are formulated in this study as first-order reactions with respect to *A*.

### 2.5. Implementation of integral control

Since the start of cybernetics during the middle of the 20th century there is now a repeated interest to understand the molecular and topological aspects behind perfect (and near perfect) adaptations and its relevance for homeostasis in biochemical/physiological systems or synthetic biology (Barkai and Leibler, 1997; Saunders et al., 1998, 2000; Yi et al., 2000; Levchenko and Iglesias, 2002; Ni et al., 2009; Drengstig et al., 2008; Ang et al., 2010; Iglesias and Ingalls, 2010; Araujo and Liotta, 2018; Krishnan and Floros, 2019; Aoki et al., 2019; Khammash, 2021; Hirono et al., 2023; Ponsiglione et al., 2023; Bhattacharya et al., 2024; Andrews et al., 2024; Hirono et al., 2025).

To ensure that the combined controllers show robust homeostasis with respect to a defined setpoint of the controlled variable I implemented integral control into the negative feedbacks. Integral control is a concept from control engineering (Warwick, 1996; Wilkie et al., 2002; Yi et al., 2000; Iglesias and Ingalls, 2010), which allows to regulate a controlled variable via a negative feedback precisely to a defined setpoint independent of step perturbations that are applied to the controlled variable. Integral feedback results in so-called ‘robust perfect adaptation’, unless the feedback encounters certain physical constraints (Ang et al., 2010).

Fig 3a illustrates the mechanism of integral control. The function *f* (*A*) of a controlled variable *A* (outlined in blue), is compared with the red outlined fixed reference *A*_*ref*_. The error, i.e. the difference *A*_*ref*_ − *f* (*A*) is calculated and integrated in time. The integrated error, often multiplied with a gain factor, results in *E* (outlined in green), which is then fed into the process to counteract stepwise perturbations applied to *A*. The general proof of this method can be found in textbooks (see for example Warwick (1996) or Wilkie et al. (2002)).

**Figure 3.**
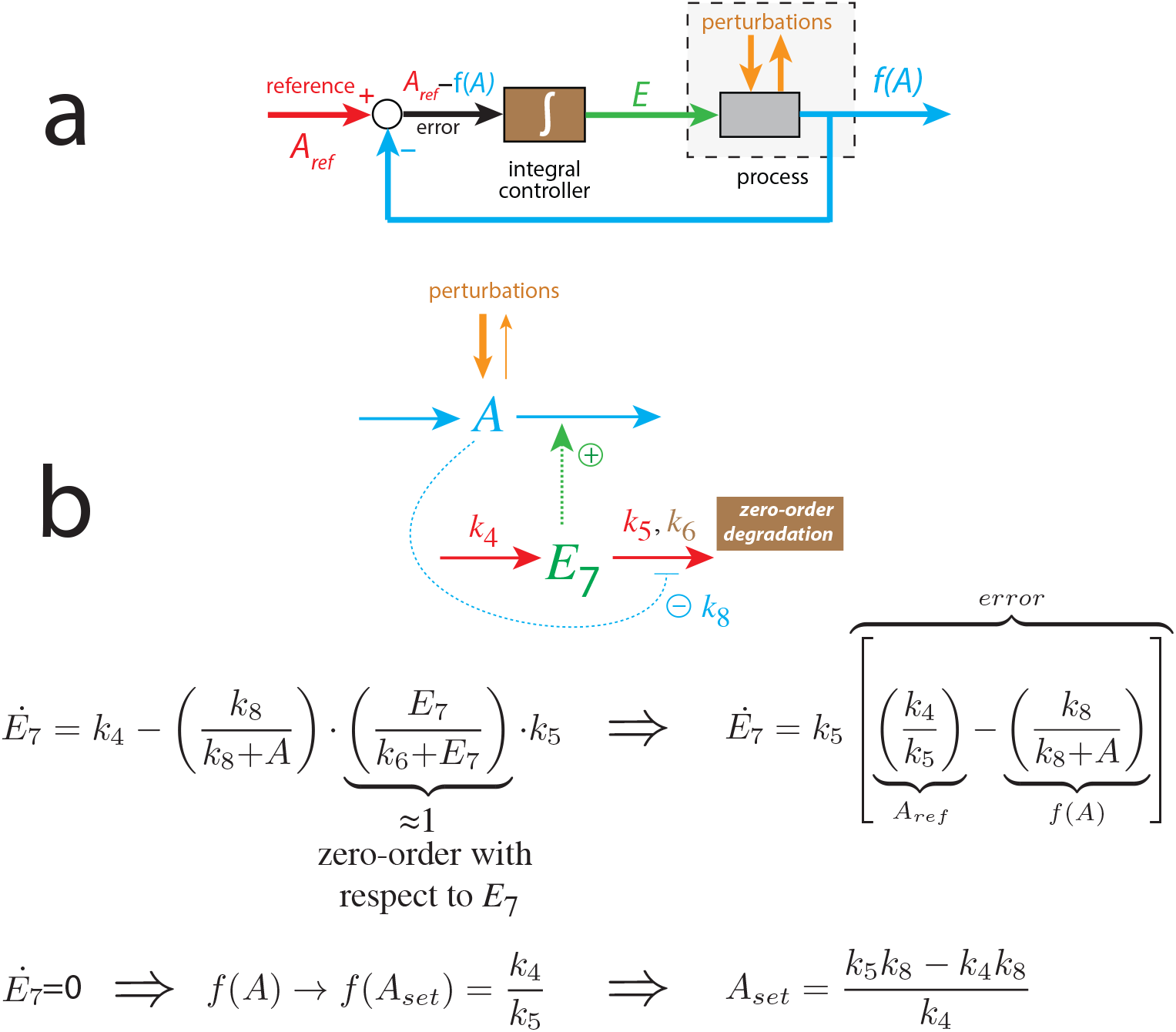
Mechanism of integral control. Panel a: General scheme to obtain integral control. The output *f* (*A*) is subtracted from the reference *A*_*ref*_ and integrated. The integrated error *E* is then fed into the negative feedback with the result to oppose perturbations acting on the controlled variable *A*. The setpoint of *A* is obtained from the relationship *f* (*A*)=*A*_*ref*_ by solving for *A*. Panel b: Motif M7 (Fig 2) is used as an illustration with color coding corresponding to panel a. The reaction scheme shows the motif with zero-order degradation of *E*_7_ by using a low *k*_6_. In all calculations a value of 1.0 × 10^−6^ is used. It reflects a strong binding between the *E*_*i*_’s (here *E*_7_) and a putative *E*_*i*_-removing/processing enzyme. When *f* (*A*)=*A*_*ref*_ steady state of *A* is reached with *A* at its setpoint *A*_*set*_.

Fig 3b shows how zero-order degradation of *E*_7_ incorporates integral control in motif M7 and defines the setpoint of *A*. The color coding in Fig 3b corresponds to that in panel a indicating the contributions of the individual rate parameters to reach integral control. For motif M7 the reference function *f* (*A*) is the strength of *A*’s inhibition on *E*_7_’s degradation. With respect to the other motifs *f* (*A*) may be simply *A* or be described by an activation of the form *K*_*a*_*/*(*K*_*a*_+*A*). When the error goes to zero *A* settles at its setpoint, which can be deduced from the relationship *f* (*A*)=*A*_*ref*_. The condition *Ė* _7_=[*A*_*ref*_−*f*(*A*)] in Fig 3b shows that *E* is proportional to the integrated error and *A* inhibits the degradation of *E*_7_ with constant strength. To show when integral control is working properly and when controller break-down occurs, a phase diagram is employed, which divides the perturbation space into regions of control, their transitions, and regions of breakdown (see section ‘Perturbation phase diagram’ below).

The usage of zero-order (or near zero-order) kinetics in the removal of *E*_*i*_ can be interpreted in form of *E*_*i*_-processing enzymes which work under saturation or near saturation conditions (Ni et al., 2009; Drengstig et al., 2008). In this respect, the Michaelis constants of these enzymes are closely related to the accuracy/inaccuracy of the controlling feedbacks (Drengstig et al., 2012a).

Other mechanisms to achieve integral control employs first-order auto-catalysis in association with a first-order removal of the controller variable *E*_*i*_ (Shoval et al., 2010; Drengstig et al., 2012b; Briat et al., 2016; M. Zand et al., 2024). This type of integral control implementation is generally fast (Nishiyama et al., 2023) and can counteract perturbations which increase rapidly in time (Fjeld et al., 2017).

In addition, topological methods have been put forward to achieve or predict robust perfect adaptation (Levchenko and Iglesias, 2002; Drengstig et al., 2008; Shinar and Feinberg, 2010; Araujo and Liotta, 2018, 2023; Hirono et al., 2023; Bhattacharya et al., 2024; Andrews et al., 2024; Hirono et al., 2025).

### 2.6. Integral windup

Windup is observed as a continuous growth or decrease in the controller variable *E*_*i*_ (see supplementary material in Drengstig et al. (2012a), p.22ff). Here, it will be shown that windup, besides leading to controller breakdown can even result in homeostatic behaviors.

### 2.7. The combined feedbacks

Fig 4 shows the sixteen inflow/outflow controller combinations between the eight feedback motifs of Fig 2. The individual inflow/outflow controllers act directly on *A* by their respective compensatory fluxes.

**Figure 4.**
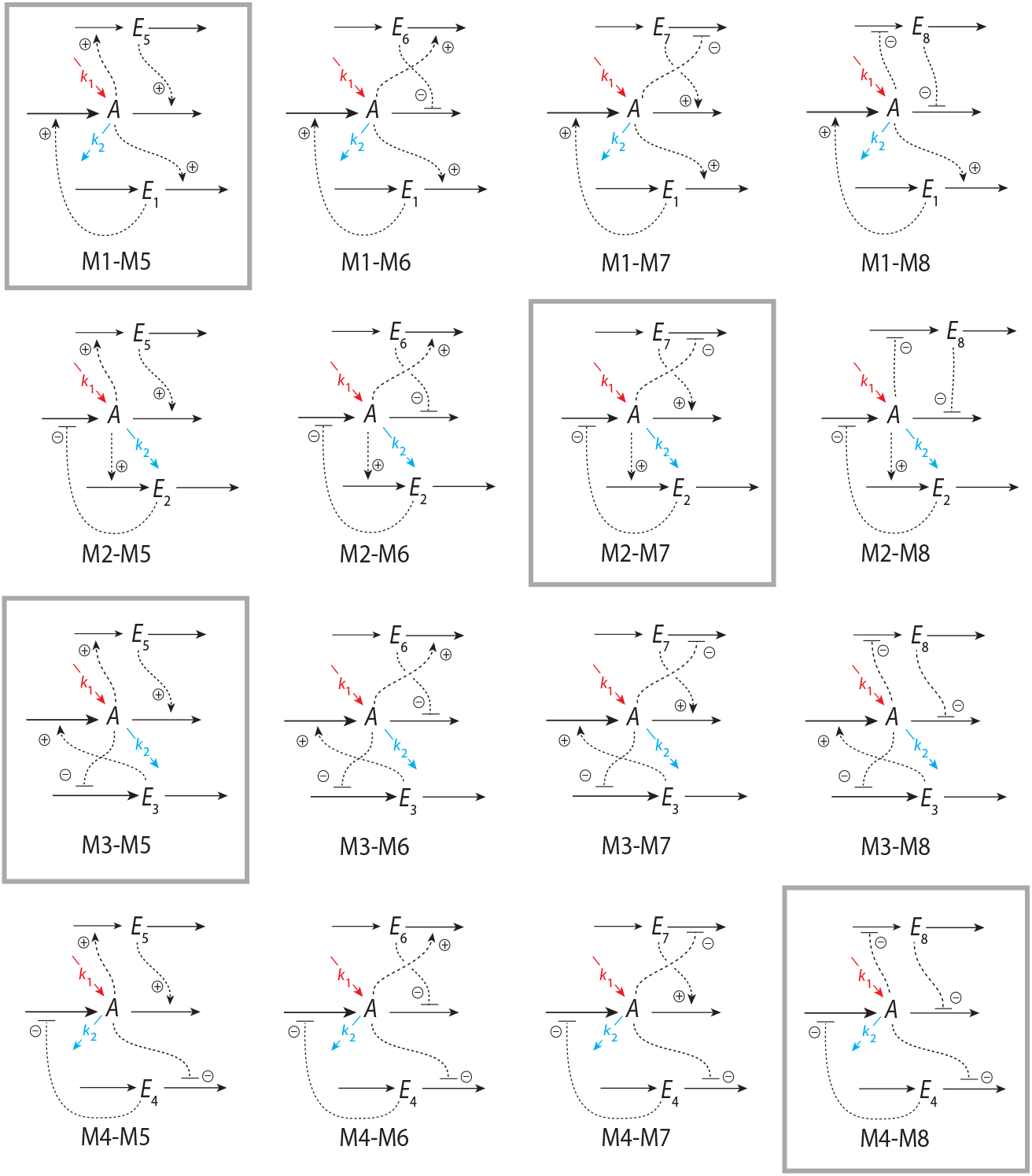
An overview of the sixteen combinations between inflow and outflow controllers. A synopsis of all controller combinations with the conditions for delegated and isolated control is found in the supplementary material Sup1.pdf. The framed schemes are described in detail in the main text (combination M1-M5), and in the supplementary material Sup2.pdf (combinations M2-M7, M3-M5, and M4-M8).

### 2.8. Perturbation phase diagram

To characterize the homeostatic behaviors of the combined controllers I make use of a *perturbation phase diagram*. The diagram, schematically outlined in Fig 5, consists of regions of control and breakdown as a function of the perturbations *k*_1_ and *k*_2_. Depending on the individual controllers’ setpoints and their signaling kinetics a *transition zone* between the regions of control may be observed. The regions of control and the transition zone are separated by border lines, which are described by linear equations of the form

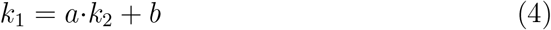

where *a* and *b* depend on the rate constants and setpoints of the combined Mi’s. The derivations of the borderlines are described in detail in the main text for combination M1-M5 and in the supplementary material Sup2.pdf for the combinations M2-M7, M3-M5, and M4-M8.

**Figure 5.**
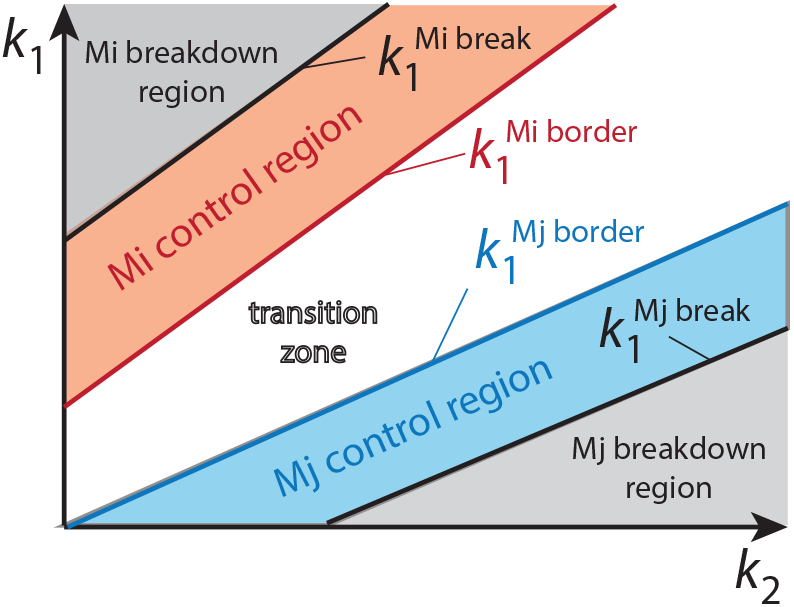
Outline of a perturbation phase diagram between combined controllers Mi and Mj. Ordinate and abscissa indicate the values of *k*_1_ and *k*_2_ perturbations acting on *A* (see Fig 1). The regions of *A*-control by Mi and Mj are outlined in red and blue, respectively. A transition zone (outlined in white) which separates the two control regions may be present. The two controllers’ breakdown regions are indicated in grey. 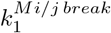 and 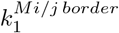 describe the borders (Eq 4) between the controlled regions and the breakdown regions, and the borders between the controlled regions and the transition zone, respectively.

## 3. Results

### 3.1. Overview

First, a brief outline of delegated and isolated control is given with detailed descriptions of the combined M1-M5 controller’s features. The supplementary material `Sup1.pdf` provides a brief overview of all controller combinations with setpoints, windup behaviors, and borderlines. In addition, supplementary material `Sup2.pdf` gives more detailed descriptions of the M2-M7, M3-M5, and M4-M8 controller combinations. Then ‘metastable control’ is described with examples in the main text and in supplementary material `Sup3.pdf`. This is followed by a chapter how robust homeostasis may even occur in open-loop designs without integral feedback.

At the end of the paper two biological examples are discussed in relationship with combined antagonistic feedbacks having different setpoints. One example concerns the photoperiodic regulation of the Siberian hamster’s body weight in terms of combined feedbacks between evening (E) and morning (M) circadian oscillators. The other example shows how blood glucose homeostasis can be understood on basis of an insulin- and glucagon-based two-setpoint mechanism and how the setpoints are affected by somatostatin.

The four supplementary files `Sup4.mp4`-`Sup7.mp4` show animations to illustrate how the rate constants of inflow and outflow compensatory fluxes affect controllers’ borderlines and their control regions. For documentation, a selected set of the Fortran results are provided in Python code.

#### 3.1.1. Flux organization in delegated control

Delegated control occurs when the setpoint of the inflow controller is larger than the setpoint of the outflow controller. Dependent on the values of *k*_1_ and *k*_2_ *A* can be controlled either by the inflow or by the outflow controller, meaning that the steady state concentration of *A* is either at 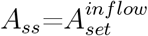 or at 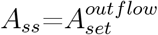. In case an outflow controller controls *A* at 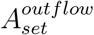 (the four motifs M5 to M8) the inflow controller will submit its fully developed compensatory flux to the system. We can distinguish two cases: (i) For inflow compensatory fluxes based on activation/deactivation (motifs M1 or M3) the compensatory fluxes take the form (Eq 2; see also Tables S1-S4 and S9-S12 in supplementary material Sup1.pdf

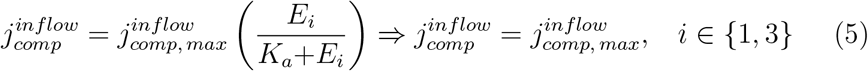

with positive windup in both *E*_1_ or *E*_3_, i.e. *Ė*_1_>0 or *Ė*_3_>0. (ii) For inflow compensatory fluxes based on repression/derepression (motifs M2 or M4) we have (Eq 1; Tables S5-S8 and S13-S16 (Sup1.pdf)):

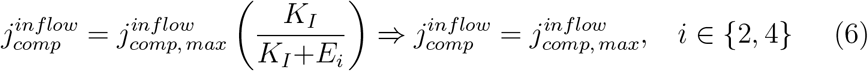

with negative windup in both *E*_2_ or *E*_4_, i.e. *Ė*_2_<0 or *Ė*_4_<0.

On the other hand, when the inflow controller determines *A* (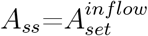, motifs M1-M4), it is the outflow controller which contributes with its fully developed compensatory flux. Also here we distinguish between compensatory fluxes based on activation (M5 or M7) and compensatory fluxes based on repression/derepression (M6 or M8). For the outflow compensatory fluxes based on activation/deactivation they take the form (see also Tables S1, S5, S9, and S13; and S3, S7, S11, and S15 in Sup1.pdf)

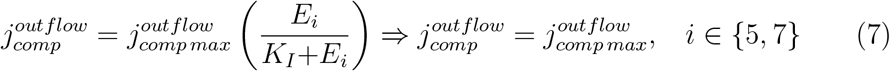

with positive windup in both *E*_5_ or *E*_7_, i.e. *Ė*_5_>0 or *Ė*_7_>0. For the compensatory fluxes based on repression/derepression (motifs M6 and M8) we get (see also Tables S2, S6, S10, and S14; and S4, S8, S12, and S16 (Sup1.pdf))

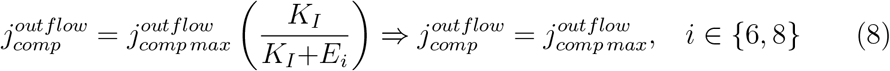

with negative windup in both *E*_6_ or *E*_8_, i.e. *Ė*_6_<0 or *Ė*_8_<0.

Fig 6 illustrates the regulation of the compensatory fluxes under delegated control. In panel a the outflow controller (outlined in violet) regulates the concentration of *A* to 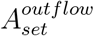 while the combined inflow controller will contribute with its maximum compensatory flux (outlined in green). Perfect adaptation at 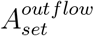 is kept as long as increasing inflow perturbations can be met with increased outflow compensatory fluxes, while increasing outflow perturbations can be balanced by decreased outflow compensatory fluxes. Panel b shows the situation when an inflow controller regulates the concentration of *A* at 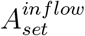. In this case the connected outflow controller will contribute with its maximum compensatory flux and the inflow controller will adjust its compensatory flux according to the inflow/outflow perturbations. Controller breakdown occurs whenever adjusted compensatory fluxes have either reached their maximum capacities or have become zero.

**Figure 6.**
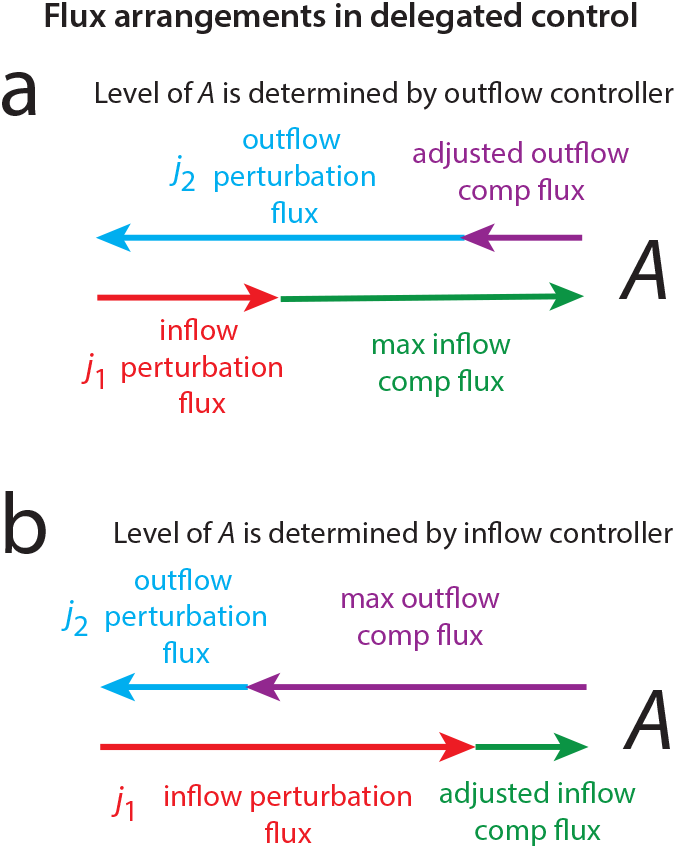
Compensatory flux arrangements in delegated control. Delegated control occurs when the setpoint of the inflow controller is larger than the setpoint of the outflow controller. Panel a: When the outflow controller determines the steady state of *A* at 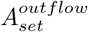 the inflow controller contributes with its maximum compensatory flux (outlined in green). To keep homeostasis in *A* the outflow controller adjusts its compensatory flux (outlined in violet) in dependence how inflow/outflow perturbations to or from *A* change (outlined in red and blue). Panel b: When the inflow controller determines the level of *A* the outflow controller’s compensatory flux is at its maximum (outlined in violet) while the green inflow controller’s compensatory flux is adjusted according to the level of the inflow/outflow perturbations (outlined in red and blue).

#### 3.1.2. Flux organization in isolated control

Isolated control occurs when the outflow controller’s setpoint is larger than the setpoint of the inflow controller. For example, when the outflow controller moves *A* to its setpoint 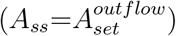 the inflow compensatory flux becomes negligible, practically zero. The reason for this is that for inflow compensatory fluxes based on activation/deactivation (M1 or M3) the corresponding *E*_*i*_’s undergo negative windup (*Ė*_1_<0 or *Ė*_3_<0) such that (see also Tables S17 to S20, and S25 to S28 in Sup1.pdf)

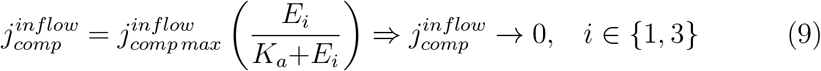

For the inflow compensatory fluxes based on repression/derepression (M2 or M4) the corresponding *E*_*i*_’s show positive windup (*Ė*_2_>0 or *Ė*_4_>0; see also Tables S21 to S24, and S29 to S32 in Sup1.pdf), which leads to

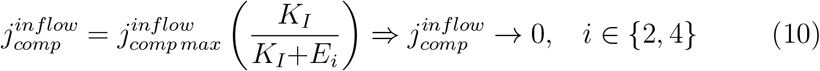

When the inflow controller determines the steady state concentration of *A* at 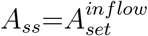 the outflow compensatory flux is negligible, i.e. the inflow controller determines the concentration of *A* alone. Here, outflow compensatory fluxes based on activation/deactivation (M5 or M7) show negative windup (*Ė*_5_<0 or *Ė*_7_<0; Tables S17, S21, S25, and S29; and Tables S19, S23, S27, and S31 in Sup1.pdf) which leads lead to

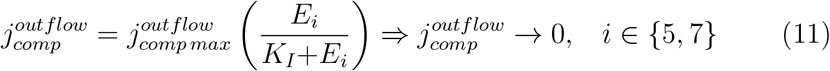

while outflow compensatory fluxes based on repression/derepression (M6 or M8) show positive windup (*Ė*_6_>0 or *Ė*_8_>0; Tables S18, S22, S26, and S30; and Tables S20, S24, S28, and S32 in Sup1.pdf) leading likewise to

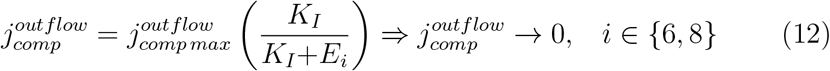

Fig 7 summarizes the compensatory flux behaviors in isolated control. In panel a the outflow controller determines *A* at 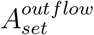 with a negligible compensatory flux of the combined inflow controller; the outflow controller solely controls *A* when inflow perturbations dominate. In panel b an outflow perturbation is dominating and the inflow controller determines *A* at 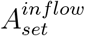. The combined outflow controller does not contribute in the regulation of *A* and has a negligible compensatory flux.

**Figure 7.**
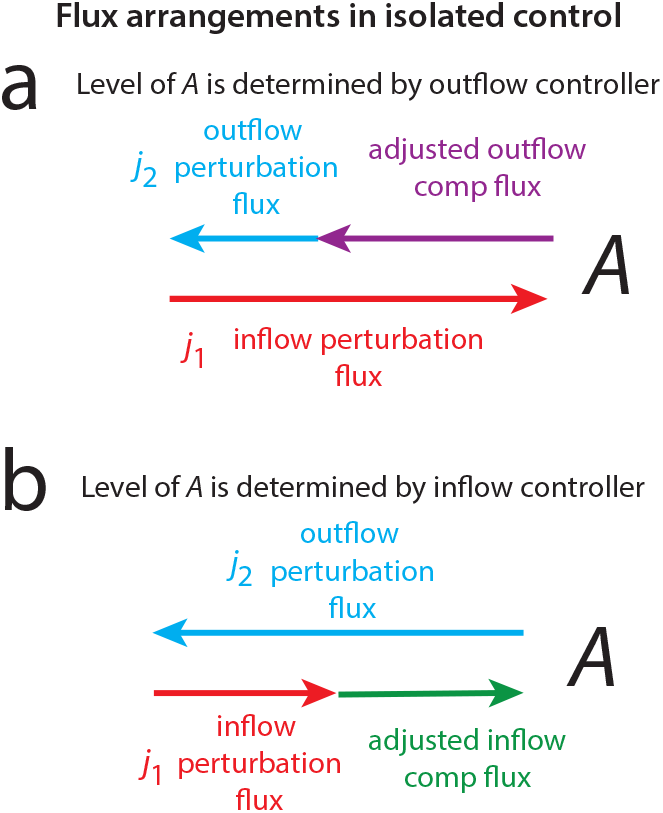
Compensatory flux arrangement in isolated control when 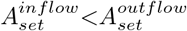. Panel a: When inflow perturbations to *A* dominate, the level of *A* is determined by the outflow controller (its compensatory flux is outlined in violet) and the inflow controller stays silent; its compensatory flux is negligible. The situation is reversed in panel b when an outflow perturbation from *A* dominates and the inflow controller (its compensatory flux is outlined in green) determines the level of *A* without any significant compensatory flux from the outflow controller.

### 3.2. Combined M1 and M5 controllers

In the following the dynamic behaviors and perturbation phase diagrams of combined M1 and M5 controllers are described in dependence to the set-points of the two controllers. Fig 8 shows the reaction scheme.

**Figure 8.**
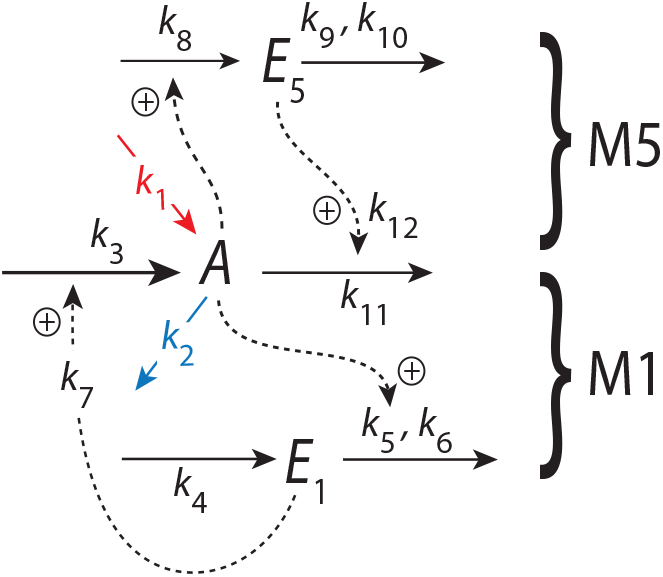
The combined M1-M5 controller. Outlined in red and blue indicate the stepwise inflow (*k*_1_) and outflow (*k*_2_) perturbations applied to *A*. M1 and M5 work by directly activating their respective compensatory fluxes *j*_3_ and *j*_11_ with activation constants *k*_7_ and *k*_12_ (Eq 13). *A* on its side activates the removal of *E*_1_ and the synthesis of *E*_5_ by first-order reactions with respect to *A*.

The rate equations are:

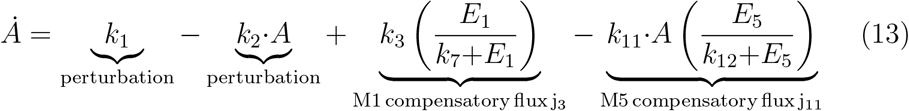

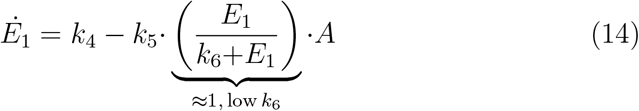

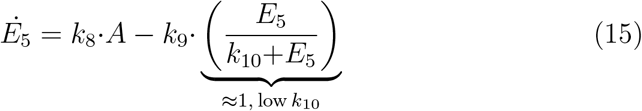

Activation of M1’s and M5’s compensatory fluxes *j*_3_ and *j*_11_ involve activation constants *k*_7_ and *k*_12_, respectively. Activations by *A* are described by first-order processes with respect to *A*. Integral control by the M1 and M5 controllers are introduced by zero-order kinetics with respect to *E*_1_ (Eq 14) and *E*_5_ (Eq 15), which give *E*_1_*/*(*k*_6_+*E*_1_) and *E*_5_*/*(*k*_10_+*E*_5_) ratios close to 1. Practically this is done by setting *k*_6_ and *k*_10_ to 1 × 10^−6^. From Eqs 14 and 15 the setpoints of the M1 and M5 controllers are then found to be, respectively,

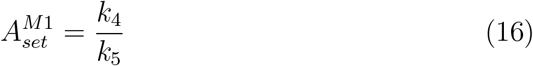

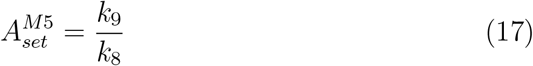

#### 3.2.1. Delegated control 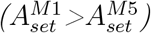

To illustrate delegated control 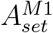 is set (rather arbitrarily) to 2.0 and 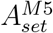 to 1.0. When M1 controls *A* M5 contributes with a fully developed compensatory flux (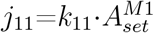, Eq 13) by windup of *E*_5_. *E*_5_ windup occurs because *A* is kept lower than 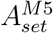, which leads to *Ė*_5_>0 (Eq 15; see also Table S1 in Sup1.pdf). On the other hand, when M5 controls *A, E*_1_ goes into windup since *A* is now kept lower than 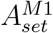 thereby resulting in *Ė*_1_>0. Thus, M1 contributes with its entire compensatory flux *j*_3_=*k*_3_ (Eq 13; Table S1, Sup1.pdf). The controller which actively regulates *A* does so by adjusting/reducing its compensatory flux in order to neutralize the exceeding compensatory inflow or outflow fluxes by the other controller (illustrated in Fig 6).

Fig 9a shows the perturbation phase diagram with four different *k*_1_ and *k*_2_ combinations during four phases which alternate between M5 and M1 control (panels b-d). Typically, the windup of *E*_1_ or *E*_5_ during respective M5 control (phases 1 and 3) or M1 control (phases 2 and 4) causes increased settling times of *A* in response to the changed setpoint (indicated by the logarithmic time scale in Figs 9b-d).

**Figure 9.**
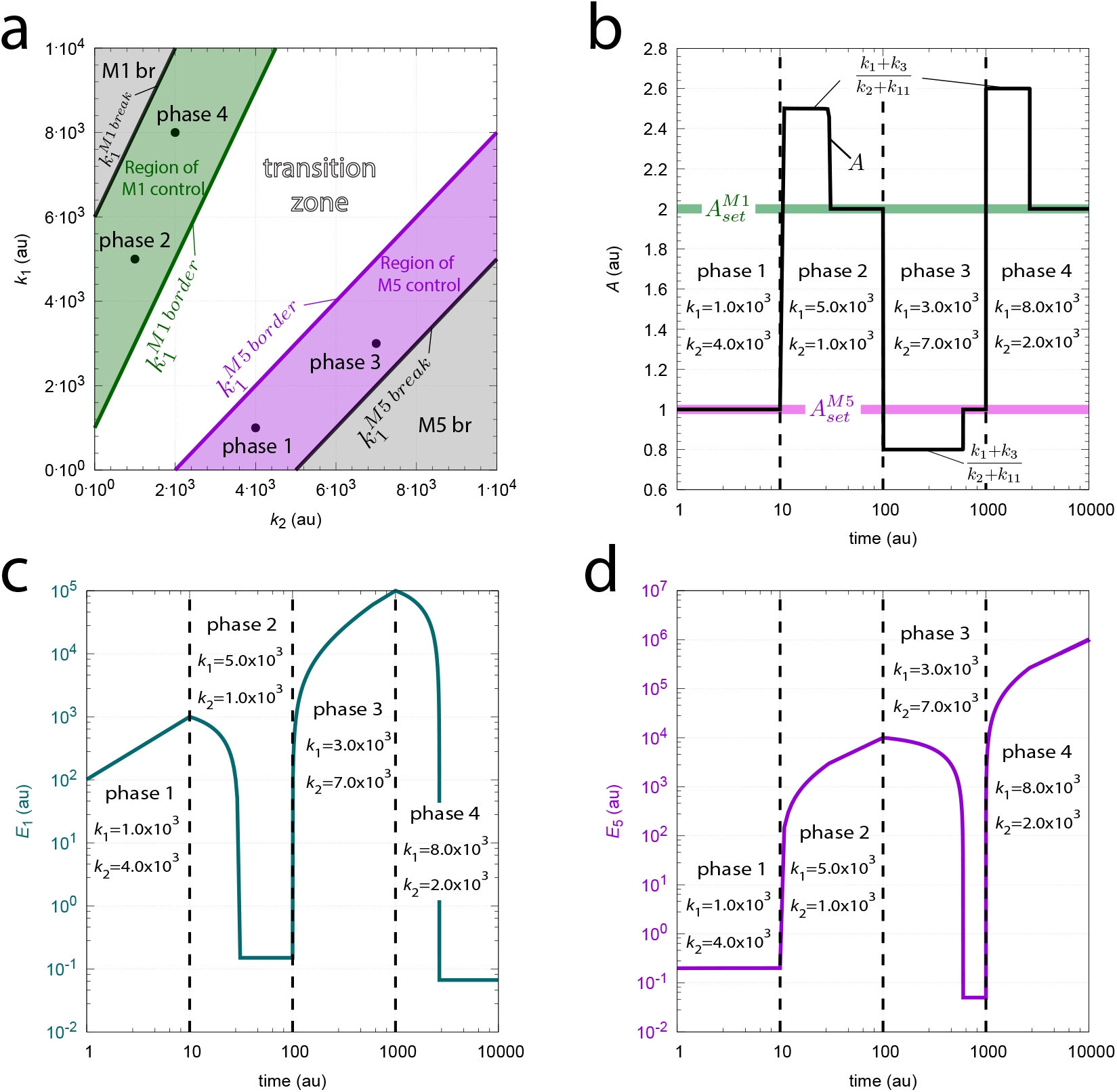
Example of delegated control in the combined M1-M5 controller when 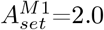 and 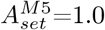. Panel a shows the perturbation phase diagram indicating the (*k*_2_, *k*_1_) values during the four phases when alternating between M5 and M1 control. Panel b: *A* as a function of time during the four phases. The temporary *A*-plateaus prior *A* reaches its setpoint occur because *j*_3_ and *j*_11_ become temporarily saturated. The values of these plateaus can be calculated by Eq 21 below. Panel c: *E*_1_ as a function of time. Panel d: *E*_5_ as a function of time. Note the logarithmic times scales in panels b-d. Rate constants: *k*_3_=5.0 × 10^3^, *k*_4_=200.0, *k*_5_=100.0, *k*_6_=1.0 × 10^−6^, *k*_7_=0.1, *k*_8_=100.0, *k*_9_=100.0, *k*_10_=1.0 × 10^−6^, *k*_11_=3.0 × 10^3^, *k*_12_=0.1. Initial concentrations: *A*_0_=1.0, *E*_1,0_=1.57, and *E*_5,0_=0.1. See Python code for details.

##### Borderlines in M1-M5 delegated control

The four borderlines between regions of control, breakdown, and the transition zone are indicated in Fig 9a. The greyed M1 breakdown region ‘M1 br’ is characterized by a low/insufficient M1 compensatory flux *j*_3_ ^∼^= 0 and a high *A*-dependent M5 compensatory flux, which is approximately described by *j*_11_=*k*_11_*A*. Thus, during M1 breakdown, *A* becomes dependent upon *k*_1_ and *k*_2_, which is approximately described by

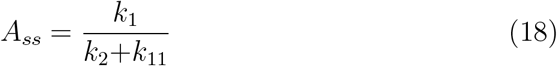

The borderline 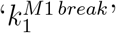, which separates the greyed M1 breakdown region from the green ‘Region of M1 control’, shows *k*_1_ as a function of *k*_2_, which is characterized by M5’s high compensatory flux at 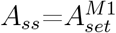 and M1’s incapability to further neutralize 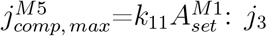 is approximately zero. Inserting these conditions into Eq 13 with steady state in *A* leads to the expression how *k*_1_ depends on *k*_2_ along the border, i.e.

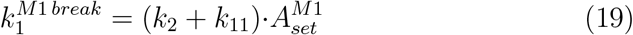

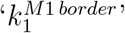 which separates the region of M1 control (outlined in green) and the transition zone (white region), is characterized that both M1 and M5 compensatory fluxes are fully developed, i.e. 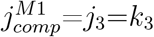 and 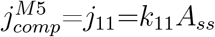. Inserting these conditions into the rate equation of *A* and setting the steady state at 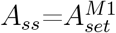 leads to the *k*_1_-*k*_2_ relationship

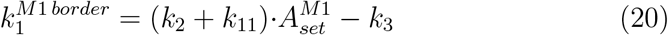

The ‘Region of M1 control’ is characterized by M5’s fully developed compensatory flux 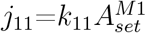. However, dependent on the *k*_1_ and *k*_2_ values M1’s compensatory flux is adjusted in such a way that it neutralizes *j*_11_ and thereby keeps *A*_*ss*_ at 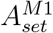. In the transition zone both M1 and M5 have maximum developed compensatory fluxes, but *A*’s steady state depends on both *k*_1_ and *k*_2_, i.e.

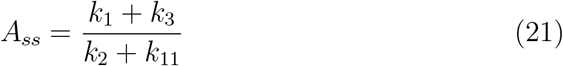

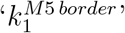 describes the *k*_1_-*k*_2_ relationship when M1 and M5 compensatory fluxes are both at their maxima, but M5 has just gained control over *A* with 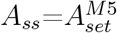:

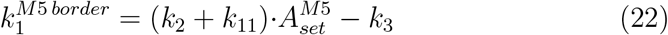

In the ‘Region of M5 control’ M1’s compensatory flux is at its maximum, but now M5 adjusts its compensatory flux such that *A*_*ss*_ can be kept at 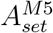. The border 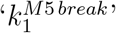 separates the region of M5 control from its breakdown region ‘M5 br’, i.e. when M5’s compensatory flux *j*_11_ becomes negligible. Assuming that *j*_11_ ≈ 0 this border is described by

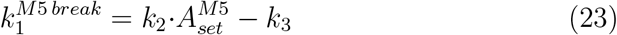

Taking that within the M5 breakdown region M5’s compensatory flux is approximately zero the steady state of *A* becomes dependent upon *k*_1_ and *k*_2_ described by

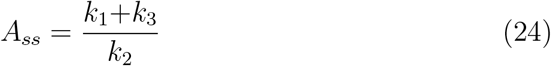

##### M1-M5 delegated control

*Moving across the perturbation phase diagram*. To test the above expressions of the borderlines and to view the behaviors of the compensatory fluxes one can look at the system when perturbations move along a trajectory across the perturbation phase diagram. Fig 10 shows the changes of *A, E*_1_, *E*_5_, and the compensatory fluxes *j*_3_ and *j*_11_ when *k*_1_, *k*_2_ values change along the blue straight line indicated in panel a. In the M1 breakdown region ‘M1 br’ (panel a) M1 cannot compensate the high *A*-dependent M5 compensatory flux *j*_11_=*k*_11_*A*. In this region *A* decreases as *k*_1_ decreases and *k*_2_ increases (panel b) according to Eq 18. At point ‘1’ the trajectory crosses the 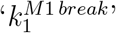 border. Within the region of M1 control *E*_1_ and the compensatory flux *j*_3_ increase. In other words, M1 now adjusts its compensatory flux to keep *A* at 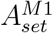 as *k*_1_ and *k*_2_ change. M5’s compensatory flux is now high and constant, described by 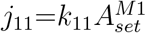 (panel d, right ordinate). When crossing 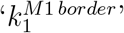 at point ‘2’ and entering the transition zone M1’s compensatory flux *j*_3_ is at its maximum value *k*_3_. M5’s compensatory flux decreases and *A* decreases in dependence of *k*_1_ and *k*_2_ according to Eq 21. Entering the M5 controlled region at point ‘3’ keeps *A* at 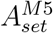 as M5 adjust its compensatory flux while *j*_3_ is at its maximum. Finally, when entering the M5 breakdown region ‘M5 br’, *A* control by M5 collapses and *A* decreases according to Eq 24.

**Figure 10.**
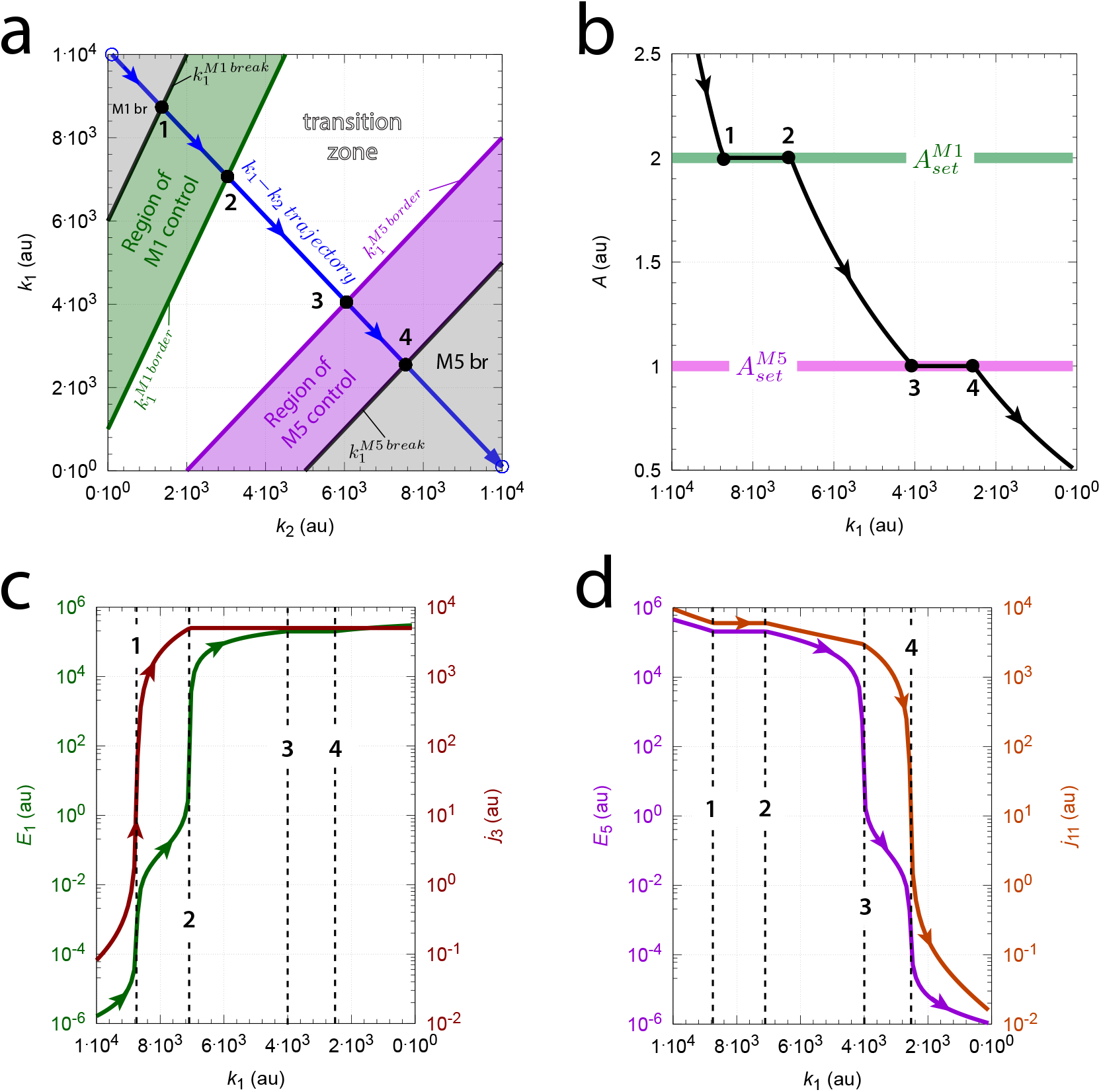
Values of *A, E*_1_, and *E*_5_ calculated along a straight line across the perturbation phase diagram (outlined in blue, panel a). Start values: 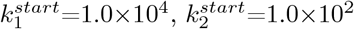. End values: 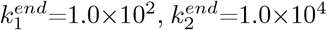. The calculation was performed by dividing the line into 100 data points and determining the values of *A, E*_1_, and *E*_5_ after 2000 time units for each point (to ensure steady state in *A*). Panel b: *A* steady state values as a function of (decreasing) *k*_1_. Panel c: *E*_1_ (left ordinate) and compensatory flux *j*_3_ (right ordinate) as a function of *k*_1_. Panel d: *E*_5_ (left ordinate) and compensatory flux *j*_11_ (right ordinate) as a function of *k*_1_. Solid black points and vertical dashed lines show the calculated *k*_1_, *k*_2_ intersection points between the blue trajectory and the border lines Eqs 19, 20, 22, and 23. Rate constants as in Fig 9. Start concentrations (for all 100 data points): *A*=3.226, *E*_1_=1.63×10^−6^, and *E*_5_=22.26.

#### 3.2.2. Isolated control 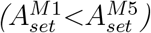

When 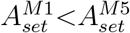 the combined M1-M5 controller shows isolated control. In this case, dependent on the perturbations, either M1 alone, or M5 alone controls *A* while the other controller remains silent. Fig 11 shows a calculation for 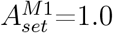 and 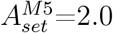 with five different *k*_1_, *k*_2_ perturbations/phases. The regions of M1 and M5 control, the transition zone, and the regions of the controllers’ breakdown is illustrated in the phase diagram shown in panel a.

**Figure 11.**
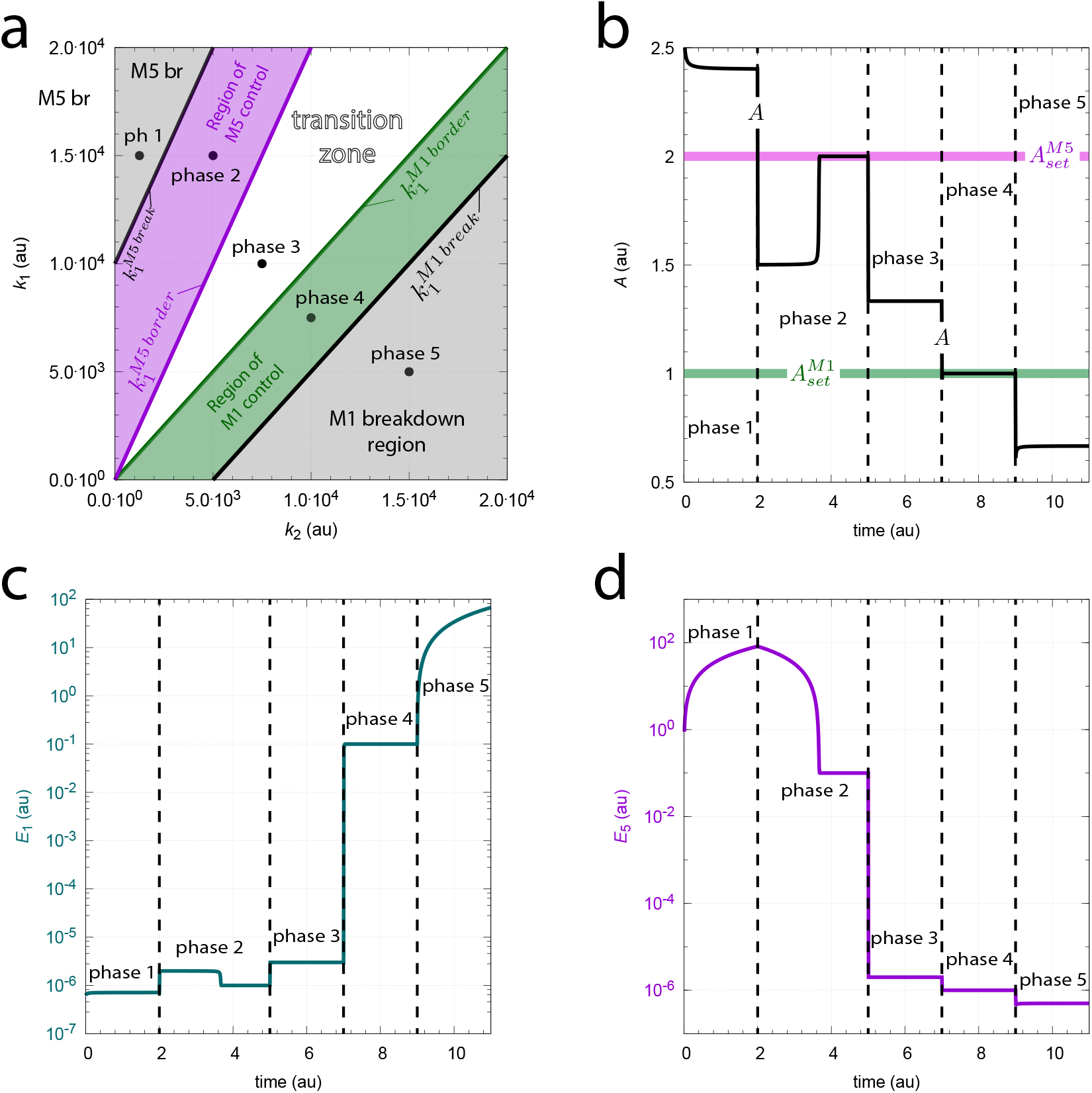
Example calculation of isolated control when 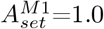 and 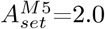. Panel a: Perturbation phase diagram with regions of M1 and M5 control (colored respectively in green and violet) and their greyed breakdown regions. In the calculation five *k*_1_, *k*_2_ perturbations were applied. Phase 1 (ph 1): *k*_1_=1.5 × 10^4^, *k*_2_=1.25 × 10^3^; phase 2: *k*_1_=1.5 × 10^4^, *k*_2_=5.0 × 10^3^; phase 3: *k*_1_=1.0 × 10^4^, *k*_2_=7.5 × 10^3^; phase 4: *k*_1_=7.5 × 10^3^, *k*_2_=1.0 × 10^4^; phase 5: *k*_1_=5.0 × 10^3^, *k*_2_=1.5 × 10^4^. Panel b: Concentration of *A* as a function of time. Panel c: Concentration of *E*_1_ as a function of time. Panel d: Concentration of *E*_5_ as a function of time. Note that the duration of phase 2 has been increased to allow *E*_5_ to reach a concentration where M5 can exert control on *A*. Rate constants: *k*_3_=5.0×10^3^, *k*_4_=100.0, *k*_5_=100.0, *k*_6_=1.0×10^−6^, *k*_7_=0.1, *k*_8_=100.0, *k*_9_=200.0, *k*_10_=1.0×10^−6^, *k*_11_=5.0×10^3^, *k*_12_=0.1. Initial concentrations: *A*_0_=2.4046, *E*_1,0_=7.12×10^−7^, and *E*_5,0_=0.0. See Python code for details.

In the transition zone both controllers have ‘shut down’ with practically zero compensatory fluxes. The *A* concentration inside the transition zone is therefore described by the ratio *k*_1_/*k*_2_. Panel b shows the concentration of *A* during the five phases. Panels c and d show respectively *E*_1_ and *E*_5_ as a function of time. Windup of *E*_1_ and *E*_5_ is only observed in respectively phase 5 and phase 1 when M1 or M5 controllers have broken down.

The borderlines between the different regions inside the phase diagram can be rationalized as follows. The 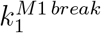 border between M1 control (outline in green, Fig 11a) and M1 breakdown (lower greyed region) has a low/negligible M5 compensatory flux (*j*_11_ ≈ 0), but a maximum (saturated) M1 compensatory flux (*j*_3_=*k*_3_) with windup in *E*_1_. Inserting these conditions into Eq 13, and allowing steady state conditions in *A*, leads to the 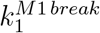 border expressed as a function of *k*_2_:

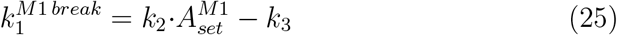

Concerning the borders between the M1 and M5 control regions and the transition zone, termed 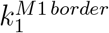 and 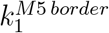, both M1 and M5 controllers are inactive with negligible compensatory fluxes *j*_3_ and *j*_11_. This leads respectively to Eqs 26 and 27:

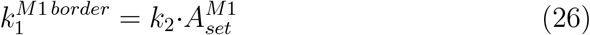

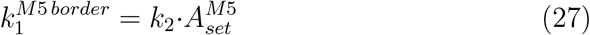

The border 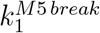 between the region of M5 control (outlined in violet in Fig 11a) and M5 breakdown (upper greyed region ‘M5 br’) is characterized by a completely saturated M5 compensatory flux 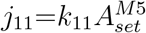 with *E*_5_ windup and a negligible M1 compensatory flux *j*_3_≈0, which leads to the expression:

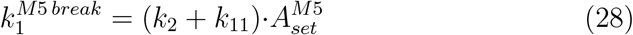

##### M1-M5 isolated control

*Moving across the perturbation phase diagram*. To test the validity of the borderline expressions Eqs 25-28 Fig 12 shows a calculation across the perturbation phase diagram along a straight line, which is indicated in blue (see panel a). The calculation starts out at high *k*_1_ (2.0 × 10^4^) and low *k*_2_ (1.0 × 10^2^) values and ends at low *k*_1_ (1.0 × 10^2^) and high *k*_2_ (2.0 × 10^4^) values. The line was divided into 100 data points and for each point the steady states of *A* were calculated. Panel b shows the numerically calculated *A* values as a function of (decreasing) *k*_1_. In addition, the solid dots 1-4 in panels a and b show the calculated intersection points between the blue line and the borderline expressions Eqs 25-28.

**Figure 12.**
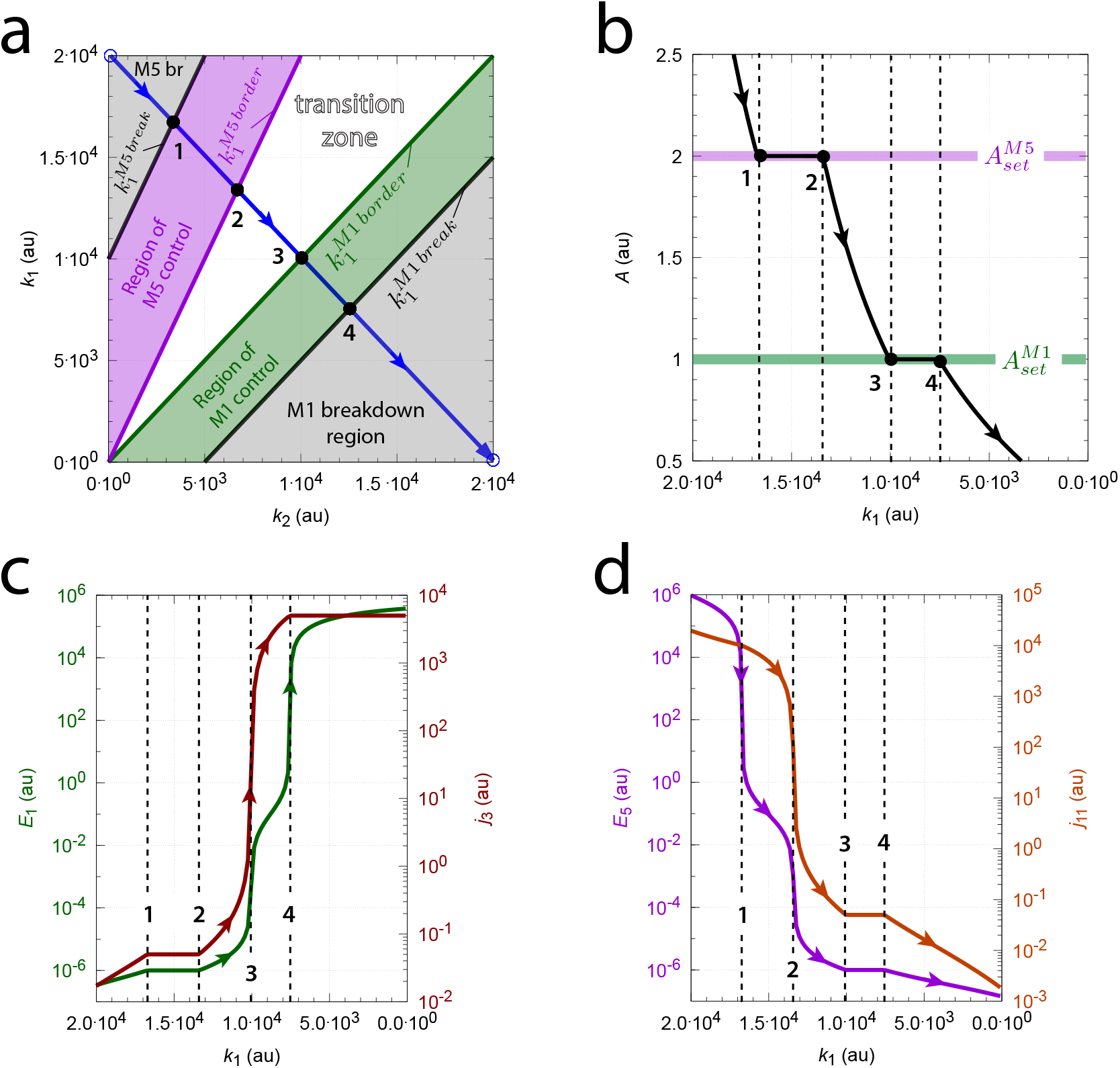
Moving across the perturbation phase diagram at isolated M1-M5 control. Panel a: Perturbation phase diagram. Start values: 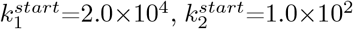. End values: 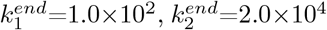. The blue line indicates the path of calculation. The path was divided into 100 sections and for each section the steady states of *A, E*_1_, and *E*_5_ were calculated after 5000 time units. Solid black dots and numbers show the calculated intersection points with the borderlines Eqs 25-28. Panel b: *A* as a function of (decreasing) *k*_1_. Panel c: *E*_1_ (left ordinate) and compensatory flux *j*_3_ (right ordinate) as a function of *k*_1_. Panel d: *E*_5_ (left ordinate) and compensatory flux *j*_11_ (right ordinate) as a function of *k*_1_. Rate constants as in Fig 11. Start concentrations (for all 100 data points): *A*=2.0, *E*_1_=1.63×10^−6^, and *E*_5_=4.901.

Fig 12c shows *E*_1_ (left ordinate) and its compensatory flux *j*_3_ (right ordinate) as a function of decreasing *k*_1_. The *k*_1_ values at the intersection points 1-4 are indicated as vertical dashed lines. Note the relative large changes of *E*_1_ and *j*_3_ at points 3 and 4 indicating active M1 control while the M5 compensatory *j*_11_ (panel d) is negligible. Likewise, during M5 control between points 1 and 2 *j*_11_ varies significantly to obtain *A* homeostasis (panel d), while M1’s compensatory flux does not contribute to the homeostasis (panel c). Fig 12d shows the corresponding *E*_5_ and *j*_11_ values as a function of decreasing *k*_1_. During M5 control (crossing points 1 and 2) there is a large change in *E*_5_ and compensatory flux *j*_11_, while during M1 control (crossing points 3 and 4) *E*_5_ and *j*_11_ are low and constant. Comparing panels c and d shows the ‘isolated control type’ by M1 or M5, i.e. the absence of any significant compensatory flux by one of the controller when the other one is in charge to keep *A* at its setpoint.

#### 3.2.3. 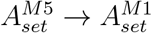: Switching from isolated to delegated control

I wondered how the system would respond when 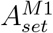 and 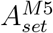 become identical. Would the system behave somewhat between delegated and isolated control? To investigate this question the controllers were compared when 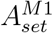 and 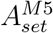 are close (but the system still shows isolated control), and when 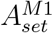 and 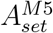 have become equal. Fig 13 (panels a1-a4) show the system’s behavior when 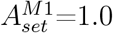 and 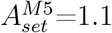. Panel a1 shows the perturbation phase diagram with the *k*_1_ and *k*_2_ values changing along the blue straight line. The other panels (a2-a4) show *A, E*_1_, *j*_3_, *E*_5_, and *j*_11_ as a function of the decreasing *k*_1_ values. As in Fig 12 the system shows isolated control, i.e. when M5 actively keeps *A* at 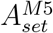 by compensatory flux *j*_11_ (with *k*_1_ values between points 1 and 2) the compensatory flux *j*_3_ of M1 is kept low and constant (panel a3). On the other hand, when M1 controls *A* at 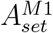 (with *k*_1_ values between points 3 and 4) M5’s compensatory flux *j*_11_ is kept low while M1’s compensatory flux *j*_3_ is high and changing with changing *k*_1_/*k*_2_. The border lines in panel a1 are described by Eqs 25-28.

**Figure 13.**
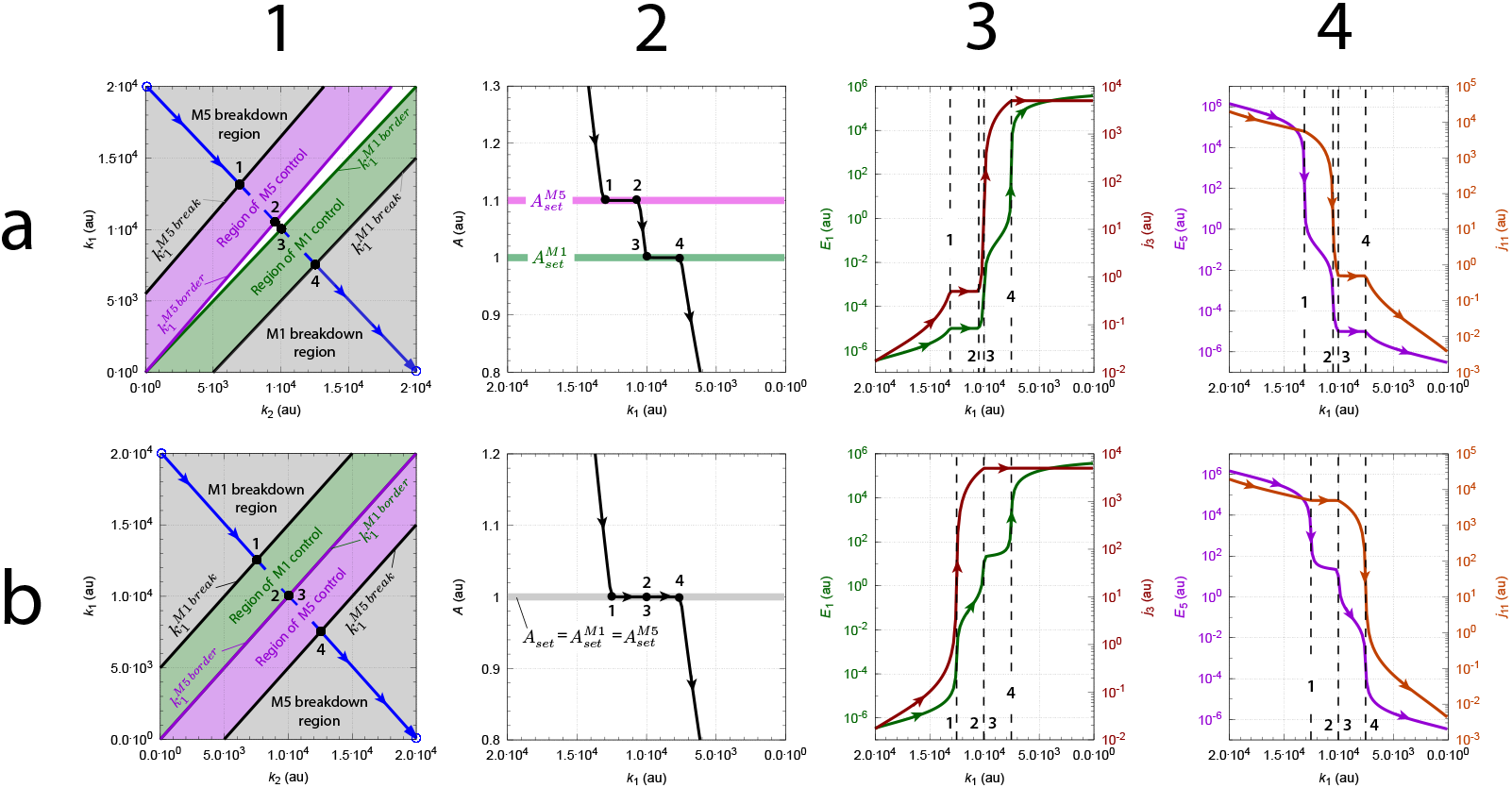
Switch from isolated to delegated control when 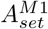 and 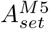 become equal. Panels a1-a4 show the perturbation phase diagram, *A, E*_1_, *j*_3_, *E*_5_, and *j*_11_ as a function of *k*_1_ when 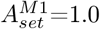 and 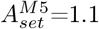. The system shows isolated control. Panels b1-b4 show the same system, but now with 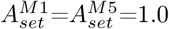. The system has switched to delegated control. Rate constants: *k*_3_=5.0×10^3^, *k*_4_=100.0, *k*_5_=100.0, *k*_6_=1.0×10^−6^, *k*_7_=0.1, *k*_8_=100.0, *k*_9_=110.0 (panels a1-a4), *k*_9_=100.0 (panels b1-b4), *k*_10_=1.0×10^−6^, *k*_11_=5.0×10^3^, *k*_12_=0.1. The blue straight lines in panels a1 and b1 are defined by the following *k*_1_, *k*_2_ values: 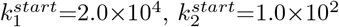 and 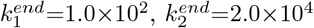. The blue lines were divided into 100 (*k*_1_, *k*_2_) data points. For each point the steady state in *A* and the other values were determined after 5000 time units. Vertical dashed lines in panels a3-a4 and b3-b4 correspond to the solid dots 1-4, which are the calculated intersection points between the blue line and the border lines Eqs 25-28 (panel a1 for isolated control) and Eqs 19-23 (panel b1 for delegated control). Initial concentrations (panels a1-a4, all 100 data points): *A*_0_=2.0, *E*_1,0_=1.6315 × 10^−6^, and *E*_5,0_=4.901. Initial concentrations (panels b1-b4, all 100 data points): *A*_0_=3.226, *E*_1,0_=1.6315 × 10^−6^, and *E*_5,0_=22.26.

By changing *k*_9_ from 110.0 to 100.0 Fig 13 (panels b1-b4) shows the same system, but now with 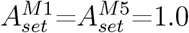: The combined controllers have switched to delegated control. The solid dots in panel b1 represent the calculated intersection points between the straight blue line of the changing *k*_1_’s and *k*_2_’s and the borderlines of delegated control, i.e. Eqs 19, 20, 22, and 23. Points 2 and 3 have now merged into one point. Panel b2 shows the value of *A* changing with decreasing *k*_1_. Panel b3 shows the delegated control by M1: While M5 controls *A* between merged points (2, 3) and 4, M1 submits its entire compensatory flux *j*_3_ to the system (right ordinate). When M1 controls *A* between points 1 and merged points 2, 3 it is now M5’s compensatory flux *j*_11_ that is high and constant (panel b4, right ordinate). Since 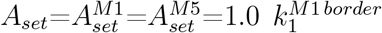 and 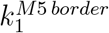 have merged. Furthermore, since *k*_3_=*k*_11_=5.0×10^3^ we have:

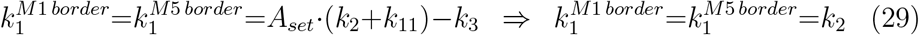

#### 3.2.4. Combined M1-M5 controllers with all four activation constants

I further investigated how Michaelis-Menten signaling kinetics from *A* to *E*_1_ synthesis and from *A* to *E*_5_ removal would affect the controlled regions of the combined M1 and M5 controllers. For this purpose I used ‘mixed activations’ as described by Eq 2. Fig 14 shows the updated reaction scheme of the combined M1 and M5 controller with the additional activation constants *k*_13_ and *k*_14_.

**Figure 14.**
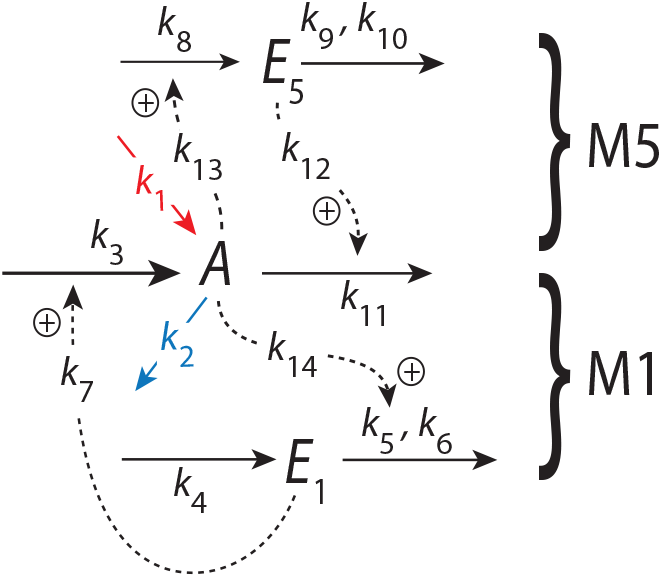
The combined M1-M5 controller with the four activation constants *k*_7_, *k*_12_, *k*_13_, and *k*_14_. M1 and M5 activate their respective compensatory fluxes *j*_3_ and *j*_11_ via activation constants *k*_7_ and *k*_12_ (Eq 30). *A*-induced activations of *E*_5_ synthesis and *E*_1_ removal have the respective activation constants *k*_13_ and *k*_14_ (see Eqs 32 and 31).

The rate equations are:

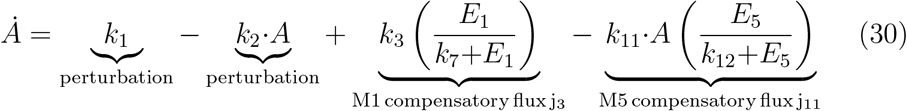

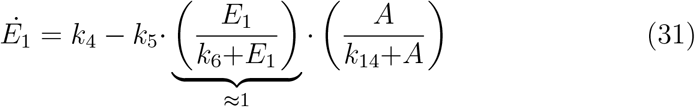

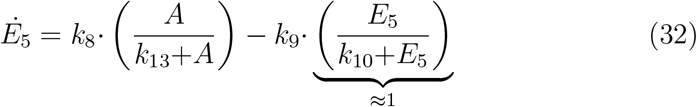

Also here integral control is introduced by zero-order kinetic removals of *E*_1_ and *E*_5_ with *E*_1_*/*(*k*_6_+*E*_1_)≈ 1 and *E*_5_*/*(*k*_10_+*E*_5_)≈ 1 by using low (1 × 10^−6^) *k*_6_ and *k*_10_ values. From the steady state conditions of Eqs 31 and 32 the setpoints by M1 and M5 are calculated to be:

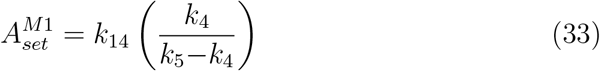

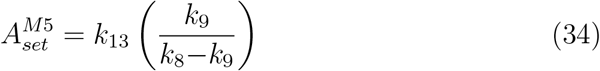

Fig 15 gives an overview of the combined M1-M5 behaviors when *k*_13_ and *k*_14_ are chosen such that 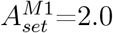 and 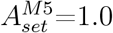 (row a), 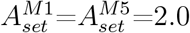 (row b), 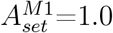 and 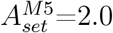 (row c). In all three instances the systems’ behaviors are calculated along a straight *k*_1_-*k*_2_ line, which starts out at 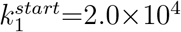 and 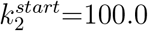 and ends at *k*_1_=100.0 and *k*_2_=2.0 × 10^4^. Columns 2-4 show respectively the concentrations of *A, E*_1_ & *j*_3_, and *E*_5_ & *j*_11_, as a function of decreasing *k*_1_.

**Figure 15.**
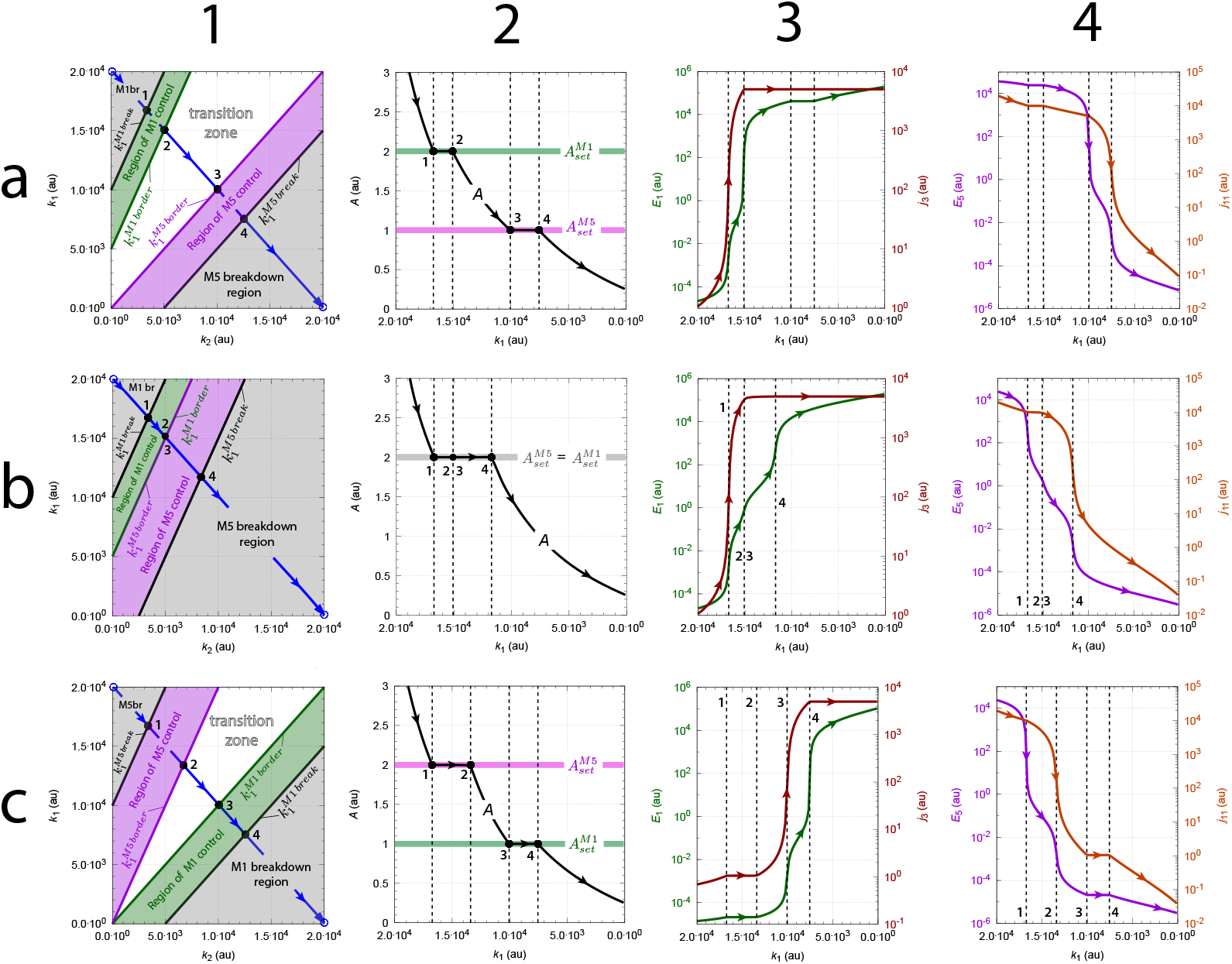
Delegated and isolated control in dependence of three 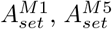 combinations (a-c; Eqs 33-34). In all three instances calculations were performed along a straight line in the perturbation phase diagram (outlined in blue in panels a1, b1, c1) and defined by the blue open circles 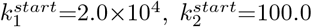, and 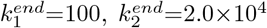. The line was divided into 100 data points and for each *k*_1_, *k*_2_ set *A, E*_1_, *E*_5_ and the respective compensatory fluxes *j*_3_ and *j*_11_ were determined after 5000 time units. Columns 2-4 show the values of *A, E*_1_, *E*_5_ and the corresponding compensatory fluxes *j*_3_ and *j*_11_. Row a: 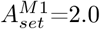 and 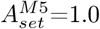 by *k*_13_=0.05, and *k*_14_=0.2. Row b: 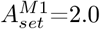 and 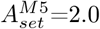 by *k*_13_=0.1, and *k*_14_=0.2. Row c: 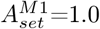 and 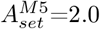 by *k*_13_=0.1, and *k*_14_=0.1. Rate constants: *k*_3_=5.0×10^3^, *k*_4_=100.0, *k*_5_=110.0, *k*_6_=1.0×10^−6^, *k*_7_=0.1, *k*_8_=210.0, *k*_9_=200.0, *k*_10_=1.0×10^−6^, *k*_11_=5.0×10^3^, *k*_12_=0.1. Initial concentrations (for all calculations): *A*_0_=2.0, *E*_1,0_=2.1×10^−5^, *E*_5,0_=0.1.

It was found that the system with four activation constants (Eqs 30-32) behaves analogous to the system with only two activation constants (Eqs 13-15). In order words, when 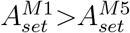 (row a, Fig 15) we have ‘delegated control’ with one controller’s compensatory flux fully contributing, while the other controller adjusts its compensatory flux in order to keep *A* at its setpoint.

When 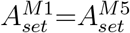 (row b, Fig 15) both controllers cooperate in their actions, i.e. while one controller (here M1) increases its compensatory flux *j*_3_ when moving along the blue line in panel b1, the other controller (M5) decreases its compensatory flux *j*_11_. When finally *j*_3_ reaches its maximum (panel b3) M5 controls *A* by further decreasing its compensatory flux (panel b4): the system shows ‘delegated control’.

When 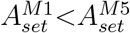 (row c, Fig 15) the system shows ‘isolated control’. Here, during control of *A* by M5 (between points 1 and 2), the compensatory flux of M1 (*j*_3_) is low and constant (panel c3) and does not significantly contribute to the control of *A*. The situation is reversed when *A* is controlled by M1: now M5’s compensatory flux (*j*_11_) is constant and negligible (panel c4).

In summary, although the introduction of the activation constants *k*_13_ and *k*_14_ affect the values of the M1 and M5 setpoints, the borderlines for delegated and isolated control are still described by respectively Eqs 19, 20, 22, and 23 and Eqs 25, 26, 27, and 28. In fact, the two sets of borderlines for delegated and isolated control remain the same also for the other controller combinations (see `Sup1.pdf`). The videos `Sup4.mp4`-`Sup7.mp4` illustrate how the control regions of the inflow and outflow controllers are affected during delegated and isolated control when rate constants *k*_3_ or *k*_11_ of the compensatory fluxes change.

#### 3.2.5. Compensatory first-order kinetics with respect to E_1_ and E_5_ leads to extended control regions

It is further found that the reaction order with respect to the controller variables affect the regions of control significantly. In the following example the compensatory fluxes of M1 and M5 are first-order with respect to *E*_1_ and *E*_5_ changing *A*’s rate equation to

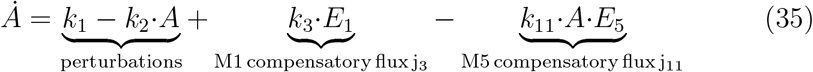

while the rate equations of *E*_1_ and *E*_5_ remain the same (Eqs 14 and 15). Enzyme-catalyzed reactions can show first-order (or close to first-order) kinetics during low-affinity binding between enzyme and substrate, i.e. when the *K*_*M*_ becomes much larger than an enzyme’s substrate concentration, i.e. when *v*=*V*_*max*_ · *S/*(*K*_*M*_ +*S*) ≈ (*V*_*max*_*/K*_*M*_) · *S*.

Fig 16 shows a calculation with the same parameters as in Fig 12 except that the rate equation of *A*, Eq 13, has been replaced by Eq 35. As the saturations in *j*_3_ and *j*_11_ are now missing the M1 and M5 control of *A* can extend along the *k*_1_ and *k*_2_ axes as long as no limits on the growth of *E*_1_ and *E*_5_ are encountered. Clearly, this is a highly idealized situation, but could occur in a biological system by different *E*_1_ and *E*_5_ species which are successively expressed when there is a need for an increased compensatory flux with lower affinity regulation. An analogous regulation is well-known to occur for low- or high affinity transport processes (Huang et al., 2011; Bosdriesz et al., 2018).

**Figure 16.**
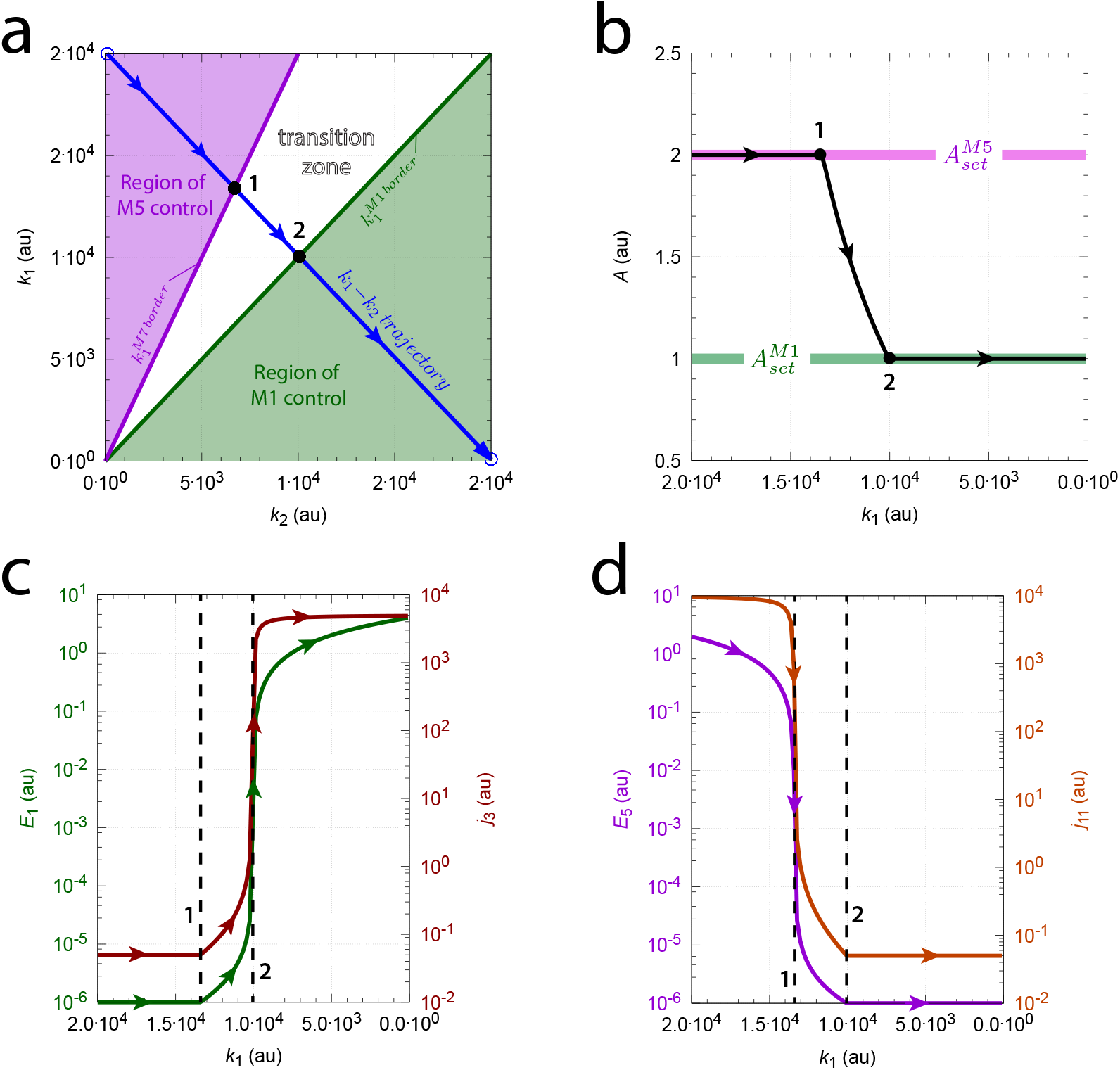
Extended regions of M1 and M5 control by first-order compensatory fluxes with respect to M1 and M4 (Eq 35). Rate parameters and initial concentrations as in Fig 12.

### 3.3. The combined controllers’ windup behaviors

In Fig 9 (and Figs S9 and S14, `Sup2.pdf`) it is shown that positive windup leads to very long settling times. On the other hand, when windups are negative settling times are short (Fig 11 and Figs S11, S16 in `Sup2.pdf`). Table 1 gives an overview of the positive and negative windup behaviors when controller combinations show delegated or isolated control. Controller combinations showing negative windup (both *Ė*_*inf low*_ and *Ė*_*outf low*_ are negative) are outlined in blue, while controller combinations with positive windup (both *Ė*_*inf low*_ and *Ė*_*outf low*_ are positive) are outlined in red. Controller combinations with mixed positive and negatived windups are indicated in yellow.

**Table 1:**
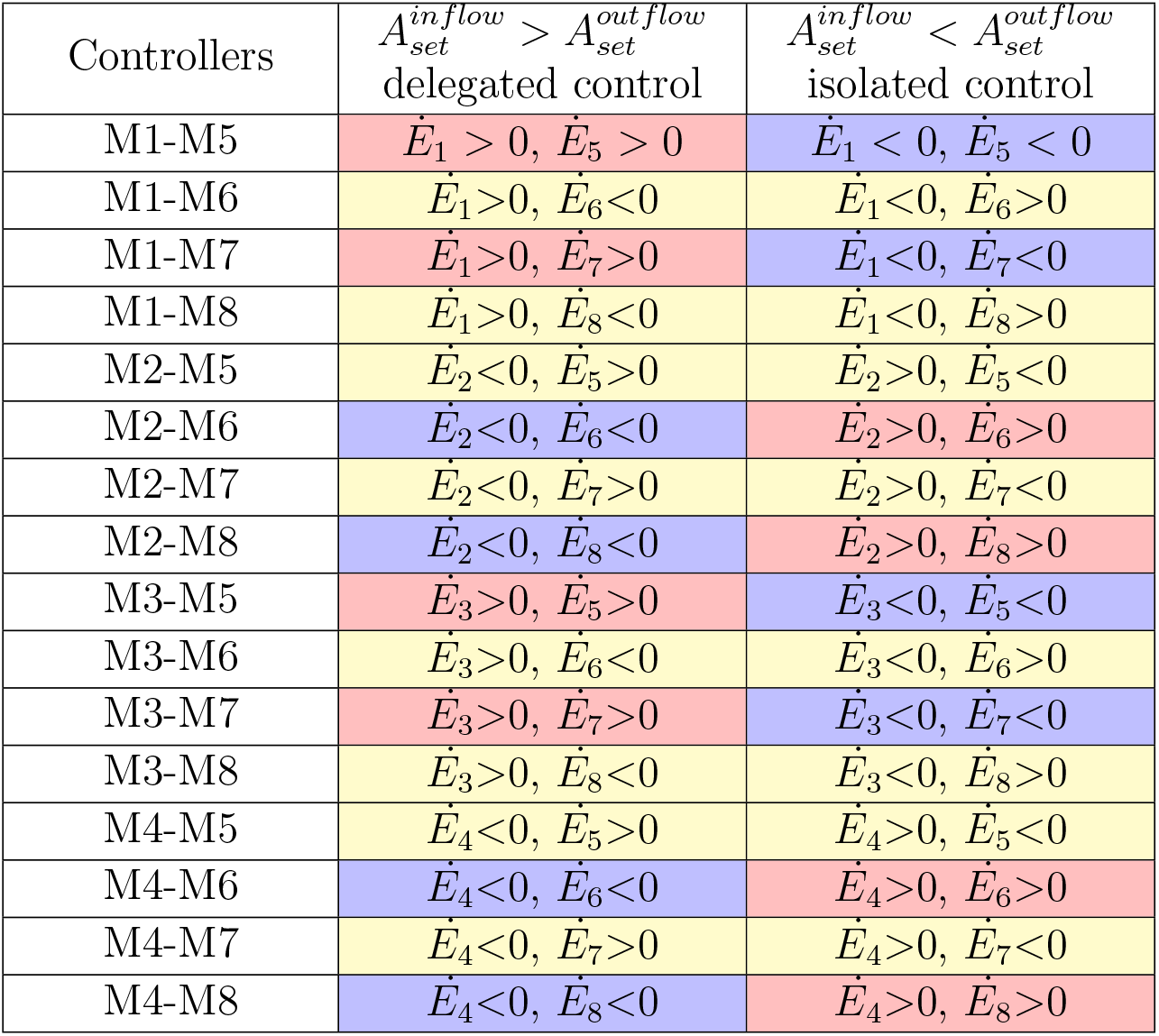
Overview of combined controllers’ windup behaviors

| Controllers | $A_{set}^{inflow} > A_{set}^{outflow}$<br>delegated control | $A_{set}^{inflow} < A_{set}^{outflow}$<br>isolated control |
| --- | --- | --- |
| M1-M5 | $\dot{E}_1 > 0, \dot{E}_5 > 0$ | $\dot{E}_1 < 0, \dot{E}_5 < 0$ |
| M1-M6 | $\dot{E}_1 > 0, \dot{E}_6 < 0$ | $\dot{E}_1 < 0, \dot{E}_6 > 0$ |
| M1-M7 | $\dot{E}_1 > 0, \dot{E}_7 > 0$ | $\dot{E}_1 < 0, \dot{E}_7 < 0$ |
| M1-M8 | $\dot{E}_1 > 0, \dot{E}_8 < 0$ | $\dot{E}_1 < 0, \dot{E}_8 > 0$ |
| M2-M5 | $\dot{E}_2 < 0, \dot{E}_5 > 0$ | $\dot{E}_2 > 0, \dot{E}_5 < 0$ |
| M2-M6 | $\dot{E}_2 < 0, \dot{E}_6 < 0$ | $\dot{E}_2 > 0, \dot{E}_6 > 0$ |
| M2-M7 | $\dot{E}_2 < 0, \dot{E}_7 > 0$ | $\dot{E}_2 > 0, \dot{E}_7 < 0$ |
| M2-M8 | $\dot{E}_2 < 0, \dot{E}_8 < 0$ | $\dot{E}_2 > 0, \dot{E}_8 > 0$ |
| M3-M5 | $\dot{E}_3 > 0, \dot{E}_5 > 0$ | $\dot{E}_3 < 0, \dot{E}_5 < 0$ |
| M3-M6 | $\dot{E}_3 > 0, \dot{E}_6 < 0$ | $\dot{E}_3 < 0, \dot{E}_6 > 0$ |
| M3-M7 | $\dot{E}_3 > 0, \dot{E}_7 > 0$ | $\dot{E}_3 < 0, \dot{E}_7 < 0$ |
| M3-M8 | $\dot{E}_3 > 0, \dot{E}_8 < 0$ | $\dot{E}_3 < 0, \dot{E}_8 > 0$ |
| M4-M5 | $\dot{E}_4 < 0, \dot{E}_5 > 0$ | $\dot{E}_4 > 0, \dot{E}_5 < 0$ |
| M4-M6 | $\dot{E}_4 < 0, \dot{E}_6 < 0$ | $\dot{E}_4 > 0, \dot{E}_6 > 0$ |
| M4-M7 | $\dot{E}_4 < 0, \dot{E}_7 > 0$ | $\dot{E}_4 > 0, \dot{E}_7 < 0$ |
| M4-M8 | $\dot{E}_4 < 0, \dot{E}_8 < 0$ | $\dot{E}_4 > 0, \dot{E}_8 > 0$ |

Since the results above indicate that the eight delegated and isolated control modes outlined in blue are expected to be more rapid in their switching/settling between and within control regions, and also more economic by avoiding time- and energy requiring windups, one might speculate whether these controller combinations may be more frequently represented in nature; required that control is mediated between flatly combined inflow and out-flow controllers as considered here. The isolated control mode of the M3-M5 controller will be used below to describe blood glucose homeostasis.

### 3.4. Metastability

The term ‘metastability’ or ‘metastable control’ refers to the observation that in certain cases one of the controllers can temporarily gain control over *A* when one of the controller variables (the *E*_*i*_’s) are suddenly added or removed. Fig 17 shows an example with the combined M1-M5 controller (Fig 8): By adding 1000 units of *E*_1_ to the system M5’s isolated control of *A* at 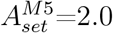 is turned into a temporary control by M1 at 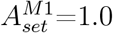. Panel a shows the system’s perturbation phase diagram with the *k*_1_ and *k*_2_ parameters indicated as a dot. Panel b shows the *A* behavior: Right after the *E*_1_ addition *A* goes into a steady state (induction period) as *j*_3_ is driven to its maximum *k*_3_, while *E*_1_ decreases (panel c). While *j*_3_ is at its maximum *E*_5_ increases, but its compensatory flux *j*_11_ is -like *j*_3_-fully saturated (panel d), i.e. for both M5 and M1 we have *E*_5_*/*(*k*_12_+*E*_5_) ≈ 1 and *E*_1_*/*(*k*_7_+*E*_1_) ≈ 1. The *A*-steady state can be determined form the rate equation of *A*:

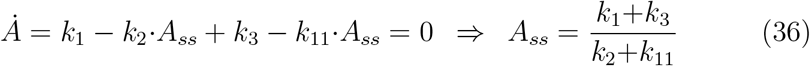

**Figure 17.**
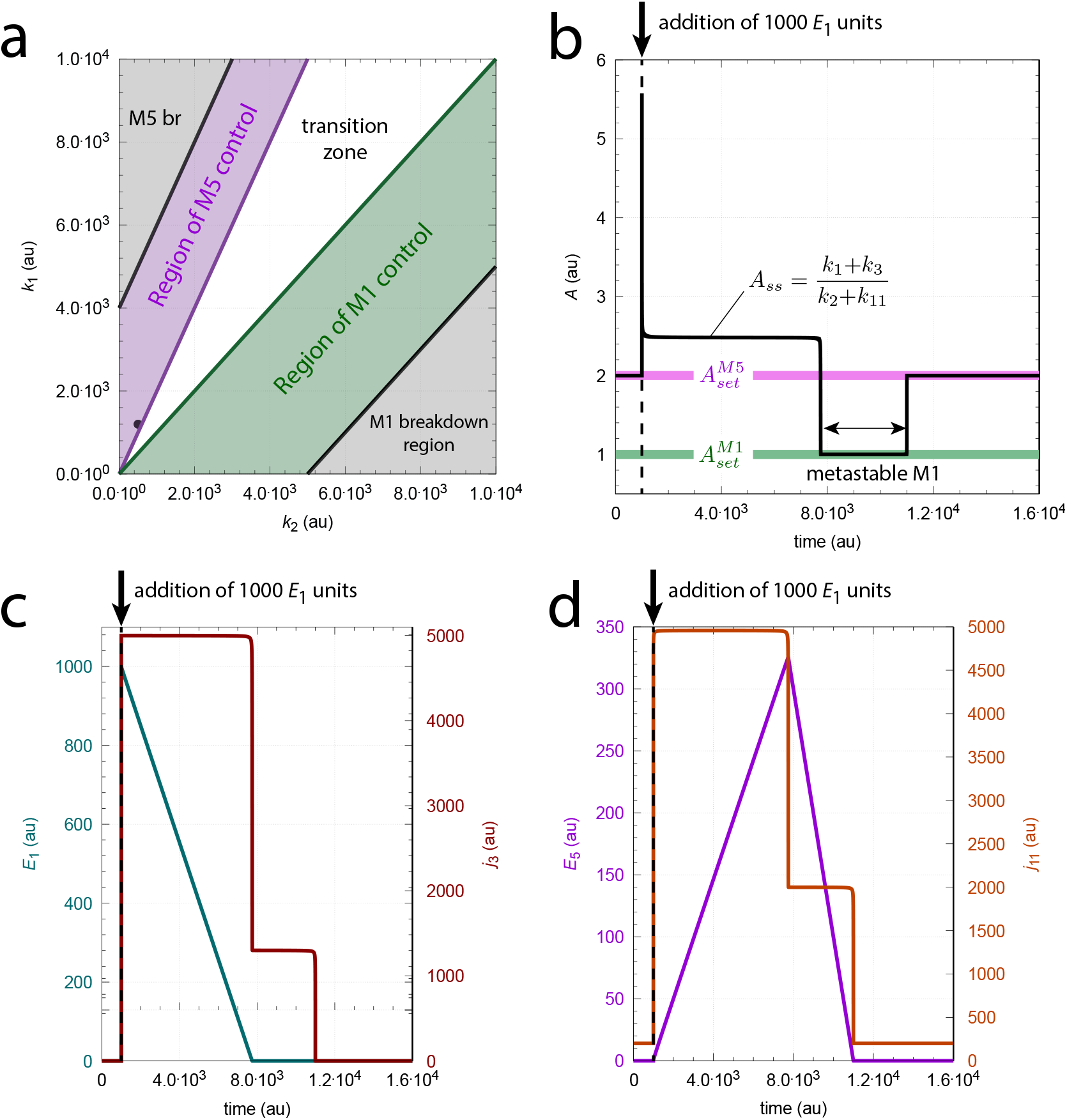
Metastable M1 control by adding *E*_1_ to isolated M5 control. Panel a: Perturbation phase diagram. The dot indicates the values *k*_1_=1.2 × 10^3^ and *k*_2_=5.0 × 10^2^. Panel b: *A* as a function of time. The metastable M1 region is indicated by the doublearrow. Panel c: *E*_1_ (left ordinate) and M2’s compensatory flux *j*_3_ (right ordinate) as a function of time. Panel d: *E*_5_ (left ordinate) and M5’s compensatory flux *j*_11_ (right ordinate) as a function of time. Vertical arrows indicate the addition of 1000 *E*_1_ units at t=1000.0. Other rate constants: *k*_3_=5.0×10^3^, *k*_4_=0.1, *k*_5_=0.1, *k*_6_=1.0×10^−6^, *k*_7_=0.1, *k*_8_=0.1, *k*_9_=0.2, *k*_10_=1.0×10^−6^, *k*_11_=2.0×10^3^, *k*_12_=0.1. Initial concentrations: *A*_0_=2.0, *E*_1,0_=1.0×10^−6^, and *E*_5,0_=5.27×10^−3^.

When *E*_1_ has reached its minimum again the growth in *E*_5_ stops and *E*_5_ starts to decrease. It is during the *E*_5_ decrease that the metastable M1 state appears (panel b). When *E*_5_ has reached its minimum *A*_*ss*_ returns to 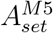.

Fig 18 shows the induction of the metastable M1 state by adding 1000 *E*_5_ units instead of adding *E*_1_, otherwise the system is the same as in Fig 17 (panel a). Here, the induction period characterized by *A*_*ss*_=(*k*_1_+*k*_3_)/(*k*_2_+*k*_11_) is missing prior to the appearing metastable M1 state (panel b). Instead, *E*_1_ and *j*_3_ go rapidly into steady states (panel c), while *E*_5_ is immediately driven to a maximum level and starts to decrease (panel d). As in Fig 17 it is during the decrease of *E*_5_ M1 metastability is observed (panel b).

**Figure 18.**
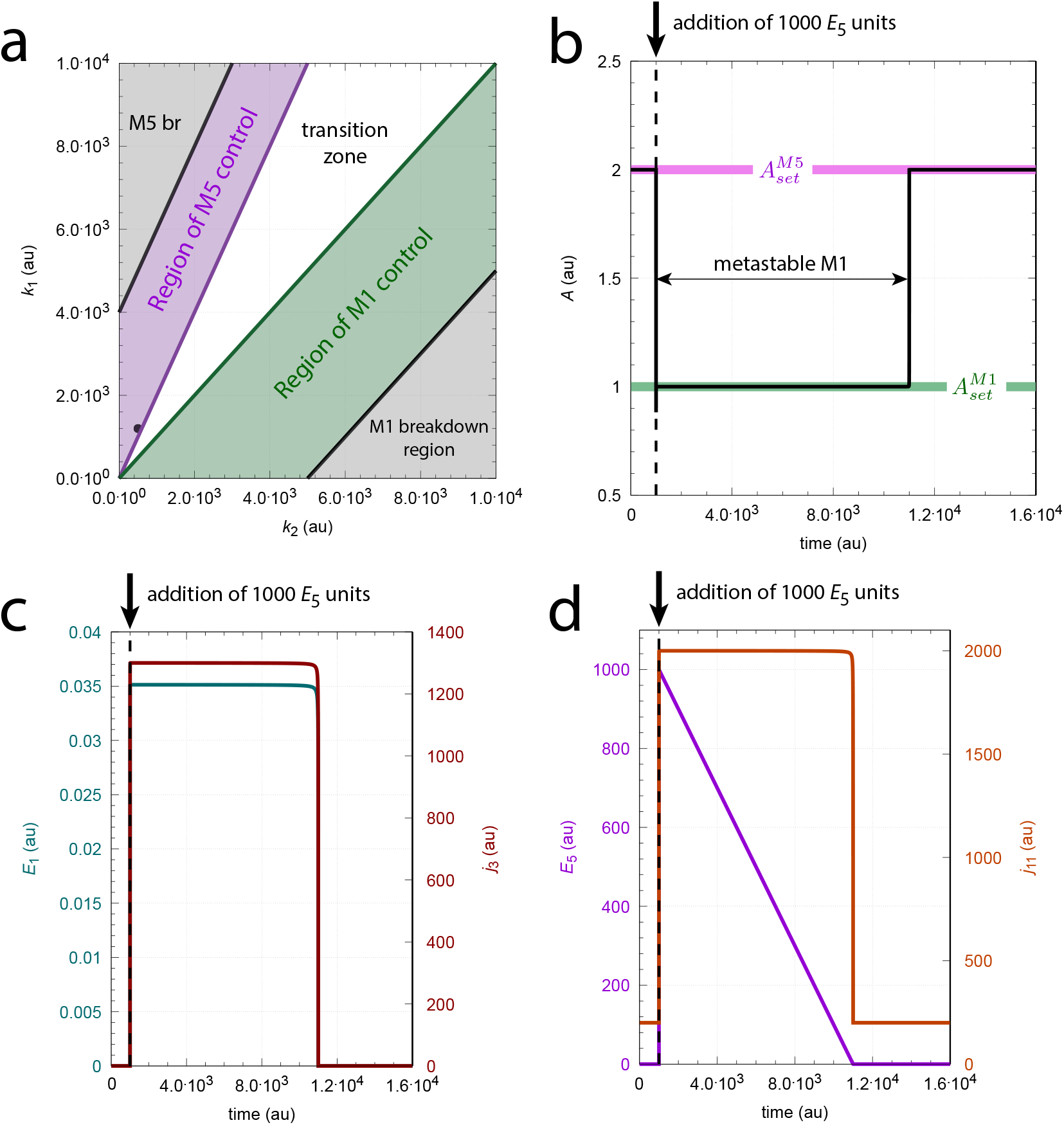
M1 metastability induced by adding *E*_5_ at isolated M5 control. Panel a: Perturbation phase diagram. The dot indicates the values *k*_1_=1.2 × 10^3^ and *k*_2_=5.0 × 10^2^. Panel b: *A* as a function of time. The metastable M1 region is indicated by the doublearrow. Panel c: *E*_1_ (left ordinate) and M2’s compensatory flux *j*_3_ (right ordinate) as a function of time. Panel d: *E*_5_ (left ordinate) and M5’s compensatory flux *j*_11_ (right ordinate) as a function of time. Vertical arrows indicate the addition of 1000 *E*_5_ units at t=1000.0. Other rate constants as in Fig 17.

There is also the peculiar case when the metastable state has the same setpoint as the original stable one. Figs 19 and 20 show two examples. In Figs 19 we start out with a stable M1 state indicated by the dot in panel a. When 1000 *E*_1_ units are added the same sequence of events occur as in Fig 16: directly after the *E*_1_ addition *A* goes into a steady state characterized by *A*_*ss*_=(*k*_1_+*k*_3_)*/*(*k*_2_+*k*_11_) (panel b) while *E*_1_ is decreasing and M2’s compensatory flux is at its maximum *j*_3_=*k*_3_ (panel c). During this period *E*_5_ increases (panel d). Once *E*_1_ has reached a minimum value *j*_3_ and *j*_11_ go into a new steady state and *E*_5_ starts to decrease. During the *E*_5_ decrease M1 is metastable (panels b and d). When *E*_5_ has reached its minimum value the end of the metastable state is indicated by a spike of *A* and the system returns to its original state.

**Figure 19.**
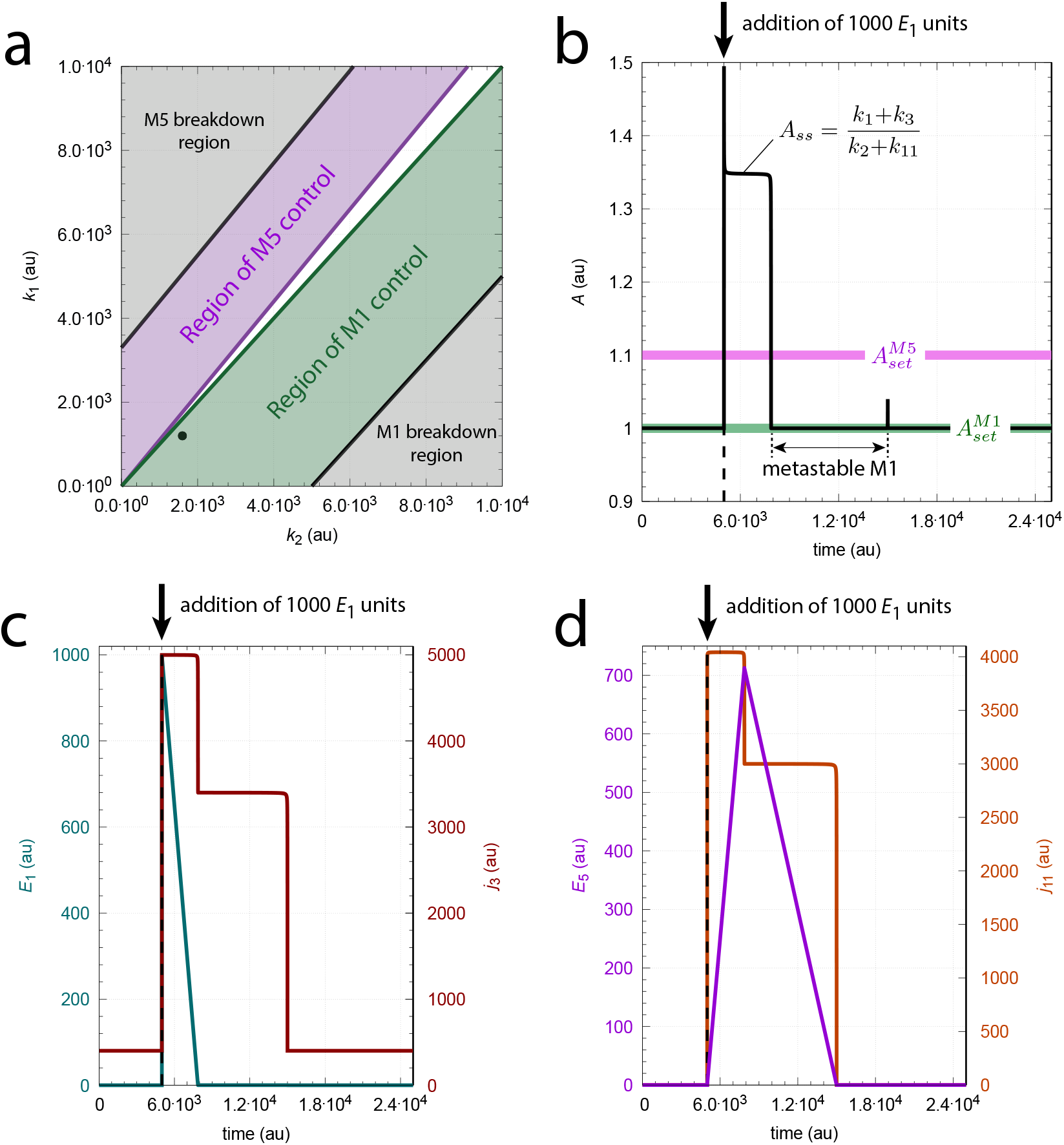
Transition between a M1 stable and a M1 metastable state by adding *E*_1_. Panel a: Perturbation phase diagram. The dot indicates the perturbations *k*_1_=1200 and *k*_2_=1600. Panel b: *A* levels as a function of time. Panel c: *E*_1_ (left ordinate) and M2’s compensatory flux *j*_3_ (right ordinate) as a function of time. Panel d: *E*_5_ (left ordinate) and M5’s compensatory flux *j*_11_ (right ordinate) as a function of time. Vertical arrows indicate the addition of 1000 *E*_1_ units at t=5000.0. Other rate constants: *k*_3_=5.0×10^3^, *k*_4_=1.0, *k*_5_=1.0, *k*_6_=1.0×10^−6^, *k*_7_=0.1, *k*_8_=1.0, *k*_9_=1.1, *k*_10_=1.0×10^−6^, *k*_11_=3.0×10^3^, *k*_12_=0.1. Initial concentrations: *A*_0_=1.0, *E*_1,0_=8.71×10^−3^, and *E*_5,0_=1.0×10^−5^.

**Figure 20.**
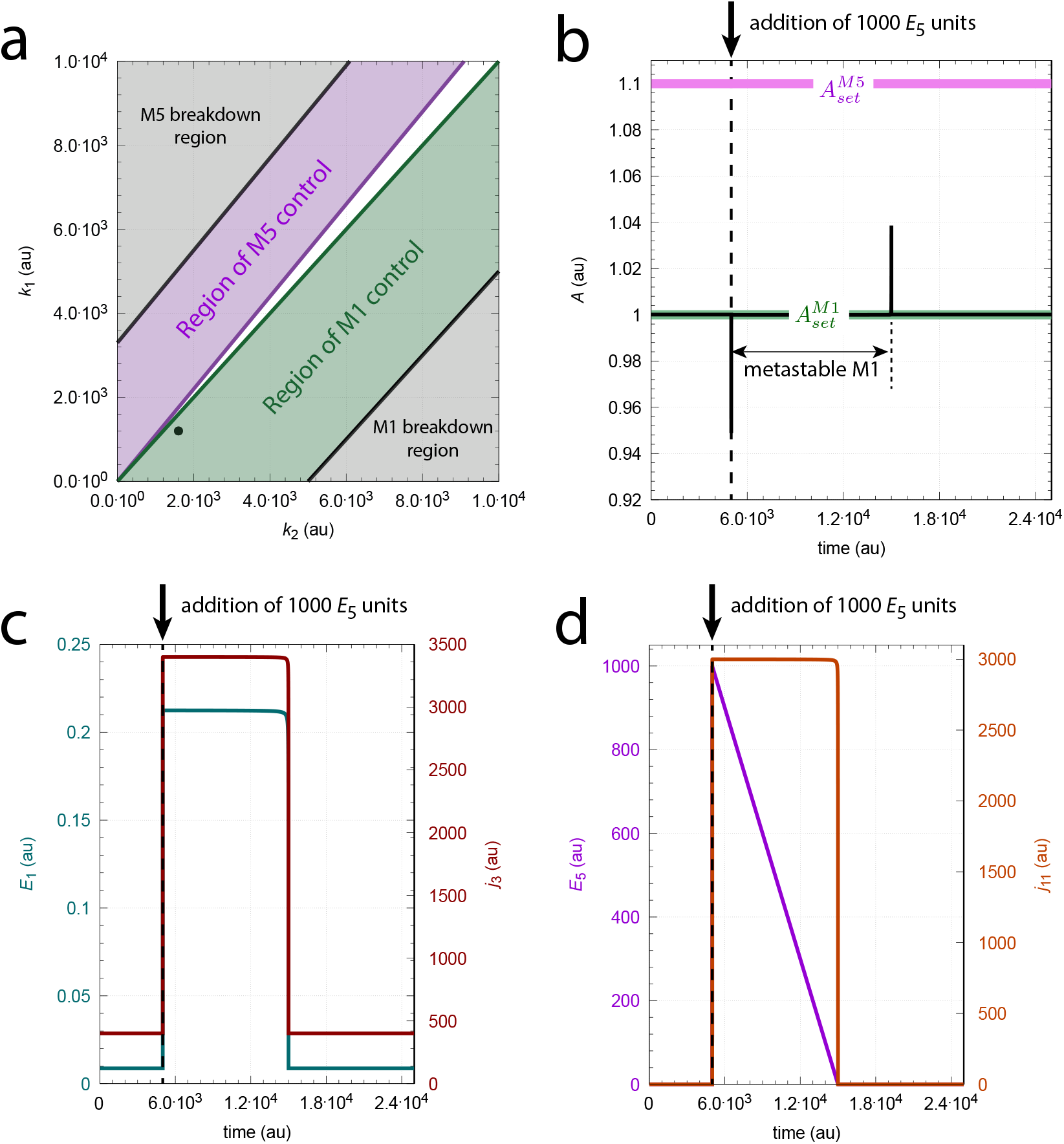
Transition between a M1 stable and a M1 metastable state by addition of *E*_5_. Panel a: Perturbation phase diagram. The dot indicates the perturbations *k*_1_=1200 and *k*_2_=1600. Panel b: *A* levels as a function of time. Panel c: *E*_1_ (left ordinate) and M2’s compensatory flux *j*_3_ (right ordinate) as a function of time. Panel d: *E*_5_ (left ordinate) and M5’s compensatory flux *j*_11_ (right ordinate) as a function of time. Vertical arrows indicate the addition of 1000 *E*_1_ units at t=5000.0. Rate constants and initial concentrations as in Fig 19.

Fig 20 starts out with the same M1 state as in Fig 19, but 1000 *E*_5_ units are added instead. The sequence of events is the same as in Fig 18: directly after the *E*_5_ addition *A* goes into the M1 metastable state (panel b) while *E*_1_ and *j*_3_ are in steady states (panel c). During the metastable M1 state *E*_5_ decreases until it reaches its minimum value and the end of the metastable state (panel d). The beginning and the end of the M1 metastable state are characterized by the negative and positive *A*-spikes (panel b).

In supporting material Sup3.pdf additional examples of metastability are given due to windup or by adding/removing a controller variable.

Concerning the question when metastability can be expected to occur or not, reference is made to Table 1. Loosely speaking, for all controller variables with positive windup no metastability is expected to occur by adding the windup species, since the addition of a controller variable does in this case not change the corresponding compensatory flux, which may already be saturated. Likewise for negative windups and removals, as removal of a controller variable already low is expected not to affect the steady state of *A*. Only during negative windup additions may lead to metastability as the four examples above for the isolated M1-M5 controller indicate. There, where the controller variable undergoes positive windup removal of this variable may temporarily drive *A* into the control domain of the other (antagonistic) controller. See for example Fig S3, panels a3 and b3 in Sup3.pdf. However, to predict the occurrence of a metastable state appears difficult since metastability is dependent on rate parameters and the relative positions of setpoints as shown in Sup3.pdf.

#### 3.4.1. Perfect adaptation without integral feedback

Robust perfect adaptation is closely associated with integral control. However, there are examples where perfect or near-perfect adaptation can occur in an open loop setting or by high and constant controller variables. I will use the combined M1-M5 controller as an illustration.

When, under delegated control, the compensatory fluxes *j*_3_ and *j*_11_ (Eq 13) are replaced by first-order expressions with respect to *E*_1_ and *E*_5_ (i.e. having low affinity compensatory fluxes), such that

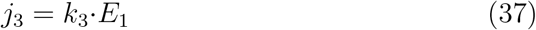

and

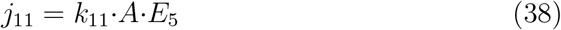

M1 and M5 work against each other and both *E*_1_ and *E*_5_ show linear integral windup as discussed in the supporting material by Drengstig et al. (2012a) (Fig S15, page 22ff). However, what was not realized at that time is that *A* in fact shows robust perfect adaptation. The setpoint

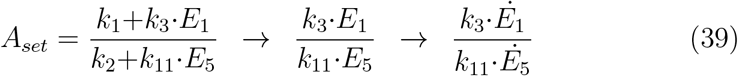

can be obtained by substituting Eqs 37 and 38 into Eq 13 and solving for *A*. The ratio after the first arrow in Eq 39 is a result of the linear increases of *E*_1_ and *E*_5_ such that the *k*_1_ and *k*_2_ terms eventually can be neglected. The ratio after the second arrow is due to the fact that *E*_1_ and *E*_5_ increase linearly with time. As *Ė*_1_ and *Ė*_5_ also depend on *k*_4_, *k*_5_, *k*_8_, and *k*_9_ *A*_*set*_ will in addition be implicitly determined by these parameters. Thus, the steady state of *A* will eventually reach *A*_*set*_ defined by

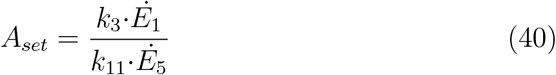

Fig 21 illustrates the behavior by using the *k*_1_/*k*_2_ values from Fig 9. The rate constants used give a *A*_*set*_ of precisely 2.0. A typical property of this type of robust control is that the *k*_1_/*k*_2_-induced *A* excursions diminish as *E*_1_ and *E*_2_ grow.

**Figure 21.**
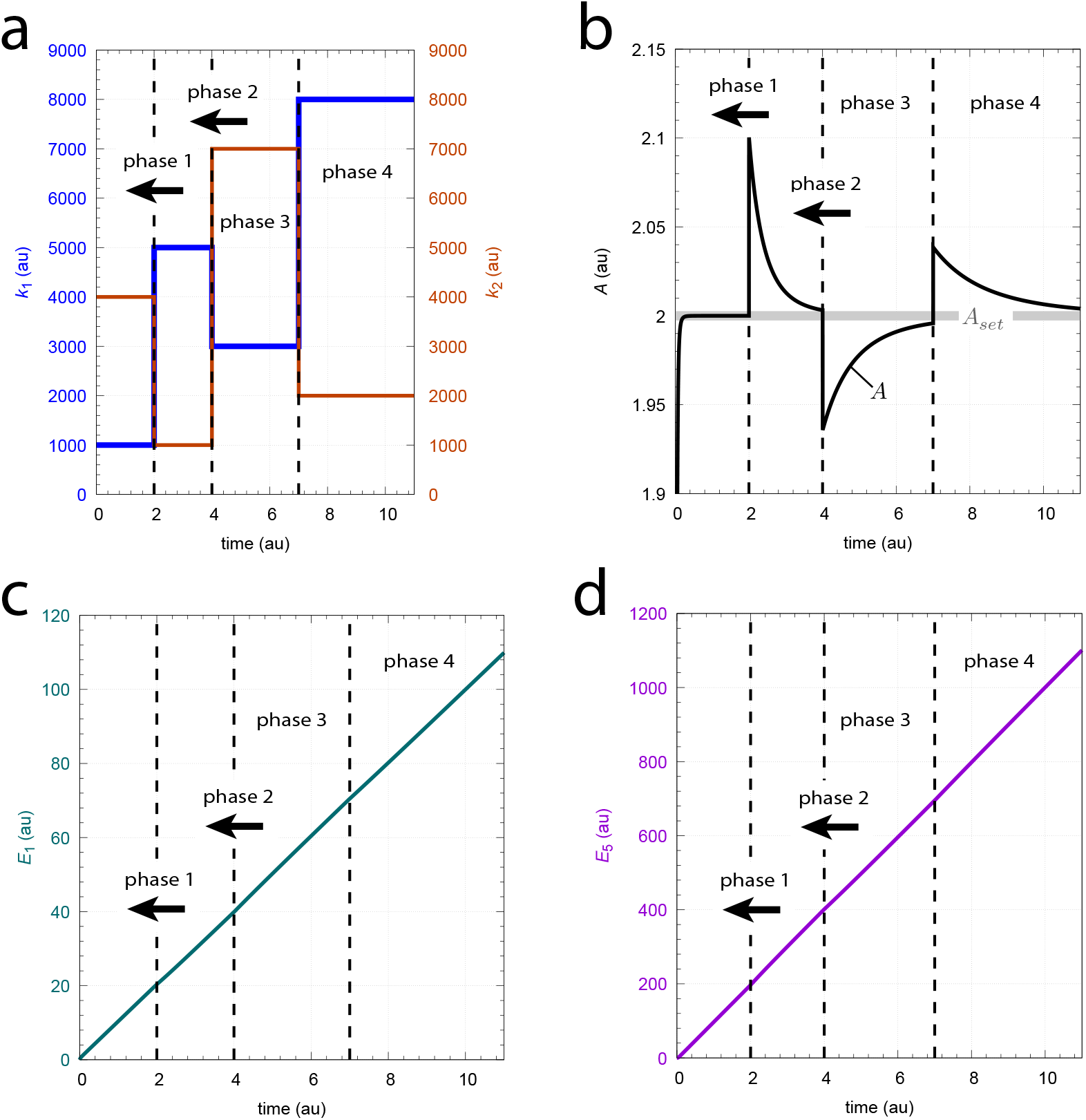
Robust perfect adaptation when 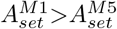 and compensatory fluxes follow first-order kinetics by Eqs 37 and 38. Panel a: *k*_1_ and *k*_2_ as a function of time. Phase 1: *k*_1_=1.0 × 10^3^, *k*_1_=4.0 × 10^3^; phase 2: *k*_1_=5.0 × 10^3^, *k*_1_=1.0 × 10^3^; phase 3: *k*_1_=3.0 × 10^3^, *k*_1_=7.0 × 10^3^; phase 4: *k*_1_=8.0 × 10^3^, *k*_1_=2.0 × 10^3^. Panel b: *A* as a function of time. Panels c and d show the linear increase of respectively *E*_1_ and *E*_5_. Rate constants: *k*_3_=1.0 × 10^4^, *k*_4_=30.0, *k*_5_=10.0, *k*_6_=1.0 × 10^−6^, *k*_7_ not used, *k*_8_=100.0, *k*_9_=100.0, *k*_10_=1.0 × 10^−6^, *k*_11_=50.0, *k*_12_ not used. Initial concentrations: *A*_0_=*E*_1,0_=*E*_5,0_=0.0. See Python code for details.

Since the M1 and M5 feedback structures in Fig 21 are broken due to the windup of *E*_1_ and *E*_5_ Fig 22 shows an analogous mechanism for robust perfect adaptation without being based on integral feedback. The rate equations are in this case:

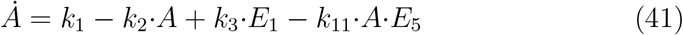

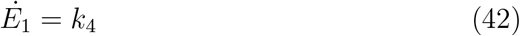

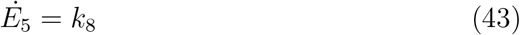

with the setpoint described by Eq 40. It may be mentioned that there occurs no ‘breakdown’ with respect to *k*_1_ and *k*_2_ perturbations as long as Eqs 41-43 are obeyed.

**Figure 22.**
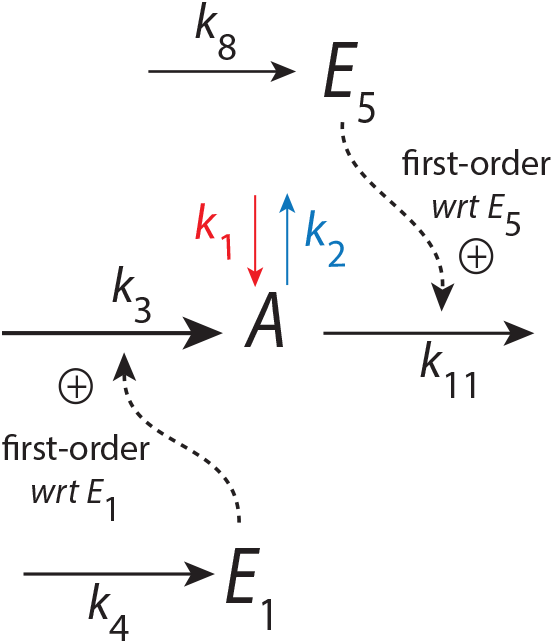
A mechanism for robust perfect adaptation without integral feedback. The mechanism is based on zero-order synthesis rates of *E*_1_ and *E*_5_ and first-order compensatory fluxes with respect to *E*_1_ and *E*_5_. For rate equations see Eqs 41-43.

#### 3.4.2. Compensatory first-order kinetics with large constant E_1_ and E_5_ leads to near perfect adaptation

The growth of the controller species (here *E*_1_ and *E*_5_) during windup cannot continue indefinitely in a physiological system. At some point resources to build *E*_1_ and *E*_5_ come to a limit. A possible scenario to maintain near perfect homeostasis (as this seem to occur in biological systems) is that *E*_1_ and *E*_5_ are kept at a constant steady-state, but that *E*_1_, *E*_5_ concentrations are such high that *A* can be kept near an unperturbed reference value (*A*_*ref*_) when *k*_1_=0.0 and *k*_2_=0.0. In other words, *A*_*ss*_ can be described during *k*_1_,*k*_2_ perturbations by Eq 39. Note that *A* does not approach a setpoint, but is kept close to *A*_*ref*_, i.e.

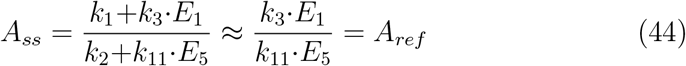

Fig 23 shows a calculation with the same perturbations and rate constants as in Fig 21, but with *E*_1_ and *E*_5_ kept constant at respectively 1 × 10^3^ and 1 × 10^5^ leading to *A*_*ref*_ =2.0. As this type of regulation can keep *A* within certain limits (of course, dependent on evolutionary anticipated *k*_1_ and *k*_2_ perturbation ranges) one may wonder to what extent such an approach may be used in physiology.

**Figure 23.**
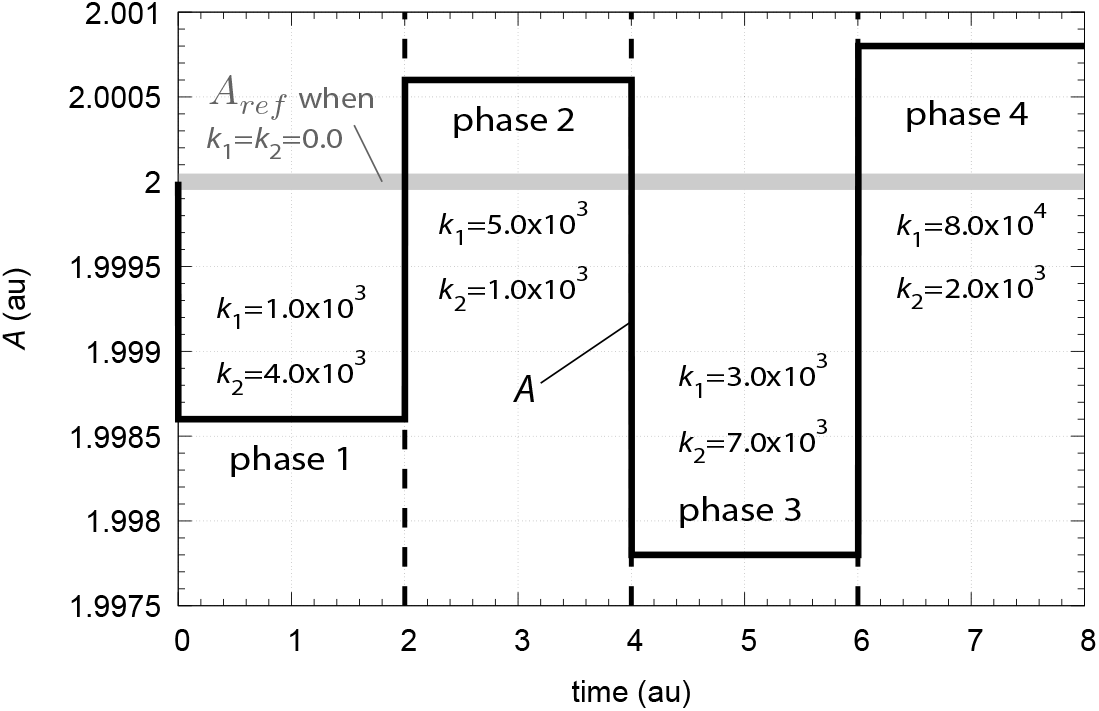
Constant *E*_1_ and *E*_5_ concentration can keep *A* levels relatively close to *A*_*ref*_ (Eq 44). Perturbations and rate constants *k*_3_ and *k*_11_ as in Fig 21. Constancy of *E*_1_ and *E*_11_ is achieved by setting in the calculations *Ė*_1_ and *Ė*_5_ to zero. Initial concentrations: *A*_0_=2.0, *E*_1,0_=1×10^3^, and *E*_5,0_=1×10^5^. See Python code for details.

## 4. Discussion: Two biological examples

Physiological regulations are extremely complex and the above described models will certainly lack many properties in an organism or cellular-wide regulatory setting. Still, two-setpoint models can provide essential regulatory behaviors as the two examples will demonstrate.

One of the distinctive properties of the above described controllers is the change of the controlled variable between the controllers’ two setpoints when environmental conditions cause a sweep across the perturbation phase diagram. A striking example of such behavior is the photoperiodic change of the Siberian hamster’s body weight.

The other example relates to blood glucose homeostasis. Although the major drivers of blood glucose regulation (i.e., insulin, glucagon, and somatostatin) are well established, there is still a debate about what determines the glycemic setpoint (Huang et al., 2024). Addressing this debate, a two-setpoint model is put forward to show how such a model in a straightforward way can describe several aspects of intact and diabetic blood glucose regulation.

### 4.1. Rheostasis by two-setpoint models: The Siberian hamster’s photoperiodic changes of body weight

An illustrative case of rheostasis, for example in comparison with Fig 16b, is the seasonal change of the Siberian hamster’s body weight. The white data points in Fig 24 show the average reference weight when hamsters were kept at summer day-night changes (16 hours light and 8 hours darkness). At week 0 hamsters were transferred to a winter light-dark regime (8 hours light and 16 hours darkness). Although food supply was not limited weights decreased and approached a new but lower level. Both the summer and the winter weights were found to be under homeostatic control. Although we do not know how *k*_1_ and *k*_2_ in Fig 16b relate to the hamster’s body weight it appears that weight changes during summer and winter periods are due to photoperiodic-induced regulations of the hamster’s food intake and consumption controlled via the hypothalamus (Bao et al., 2019).

**Figure 24.**
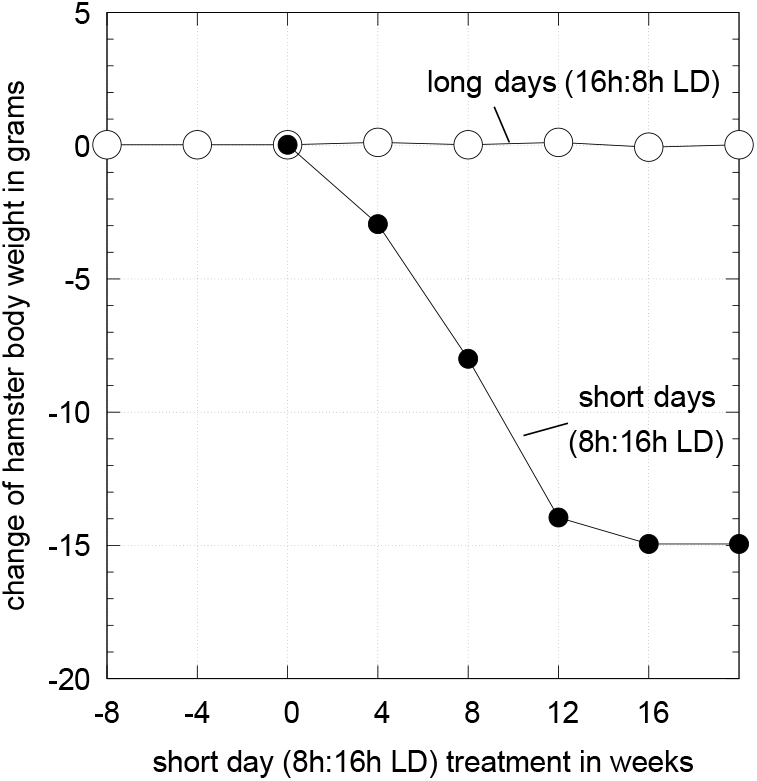
Photoperiod induced changes of the Siberian hamster’s body weight. White dots show the control when hamsters are kept at a summer 16h:8h light:dark (LD) regime. Black dots show the loss of body weight when hamsters were transferred to winter 8h:16h LD changes. Both summer and winter weights were found to be under homeostatic control. Redrawn from Stevenson (2024), Fig. 2.4.

The data in Fig 24 have been interpreted (Stevenson, 2024) to be due to rheostatic control, which includes an apparent change in setpoint. There is the claim (Mrosovsky, 1990; Stevenson, 2024) that homeostatic and rheostatic control are fundamentally different. Among the differences it has been postulated that rheostatic systems lack a defined setpoint. Although this may be possible, the combined M1-M5 controller of Fig 16b and the other examples above show that analogous behaviors to Fig 24 can indeed be described by homeostatic mechanisms with defined setpoints.

Concerning the physiological regulation of the Siberian hamster’s body weights, the two negative feedbacks which appear to be involved relate to two tissue-dependent circadian oscillators. They were identified in hamsters’ superchiasmatic nucleus (which is part of the hypothalamus) as morning (M) and evening (E) oscillators (Jagota et al., 2000), after postulating them by Pittendrigh and Daan (1976). Circadian oscillations are based on transcriptional-translational negative feedback loops (Dunlap, 1999), which have homeostatic properties (Saito et al., 2025) and show temperature compensation (Rensing et al., 1997). The circadian organization into two antagonistic negative feedbacks seems to be a general regulatory feature and has been found also in other organisms, ranging from protists (Morse et al., 1994) to fungi (De Paula et al., 2007), insects (Stoleru et al., 2007), and mammals (Helfrich-Förster, 2009). The body weight regulation in hamsters have been found to occur via melatonin, an important transducer of photoperiodic signals. Changes in hamsters body weight can in fact be mimicked by systemic administrations of melatonin (Bartness and Wade, 1985). By studying the enzyme arylalkylamine N-acetylytansferase (NAT), which takes part in the synthesis of melatonin, Illnerová (1991) concluded that there is strong support that NAT rhythmicity and melatonin production is due to the dual E-M negative feedback structure proposed by Pittendrigh and Daan. In this respect, *A* would refer to melatonin which determines hamster body weight while the perturbations would relate to the applied photoperiod. While our above two-feedback approach is clearly oversimplified with respect to hamster physiology, it would nevertheless be interesting to pursue this line of research under oscillatory circadian conditions.

Arguments brought forward against a homeostatic interpretation of rheostatic behaviors have implicitly assumed that homeostasis occurs only by single negative feedback loops (see for example chapter 1 in Mrosovsky (1990)). However, such an assumption has never been part of Cannon’s definition of homeostasis. In the definition chapter of his 1929 paper Cannon writes (page 251): *The coordinated physiological reactions which maintain most of the steady states in the body are so complex, and are so peculiar to the living organism, that it has been suggested Cannon (1926)* [see a facsimile of this citation in Langley (1973)] *that a specific destination for these states be employed–homeostasis*.

‘Rheostasis’ as a descriptive term to indicate apparent changes in a setpoint (or using the term *enantiostasis* (Mangum and Towle, 1977; Hochachka and Somero, 1984) to describe a preserved functional relationship with respect to environmental changes) is useful, but the declaration that the underlying control mechanisms of rheostasis are devoid of setpoints and of homeostatic mechanisms, (see description of Fig 2.5 in Stevenson (2024)), appears too dogmatic.

### 4.2. Blood sugar homeostasis: Predictions by a two-setpoint model

Insulin, glucagon and somatostatin are the major regulators of blood sugar homeostasis. They are produced by different cell types contained in the islets of Langerhans (Clancy and McVicar, 2002; Kumar et al., 2013). Beta cells produce insulin, which, among other roles, enables the entrance of glucose into cells to generate energy.

In this respect, insulin is part of an *outflow controller* which reduces the amount of blood sugar. Alpha cells produce glucagon, which have an glycogenolytic activity, i.e. glucagon increases blood glucose from stored glycogen in the liver. In this respect glucagon is part of an *inflow controller*. Delta cells produce somatostatin, which inhibits both insulin and glucagon release. The most important disease of blood sugar regulation is diabetes mellitus when insulin secretion or its action is defective causing an excess of glucose in the blood (hyperglycemia).

In healthy individuals blood glucose is robustly regulated between about 70 mg/dL (4 mM) and 130 mg/dL (10 mM), but allows for variations within these limits (Fig 25a). Glucose levels which stay above 180 mg/dL or below 40 mg/dL will cause serious health problems (Kumar et al., 2013).

**Figure 25.**
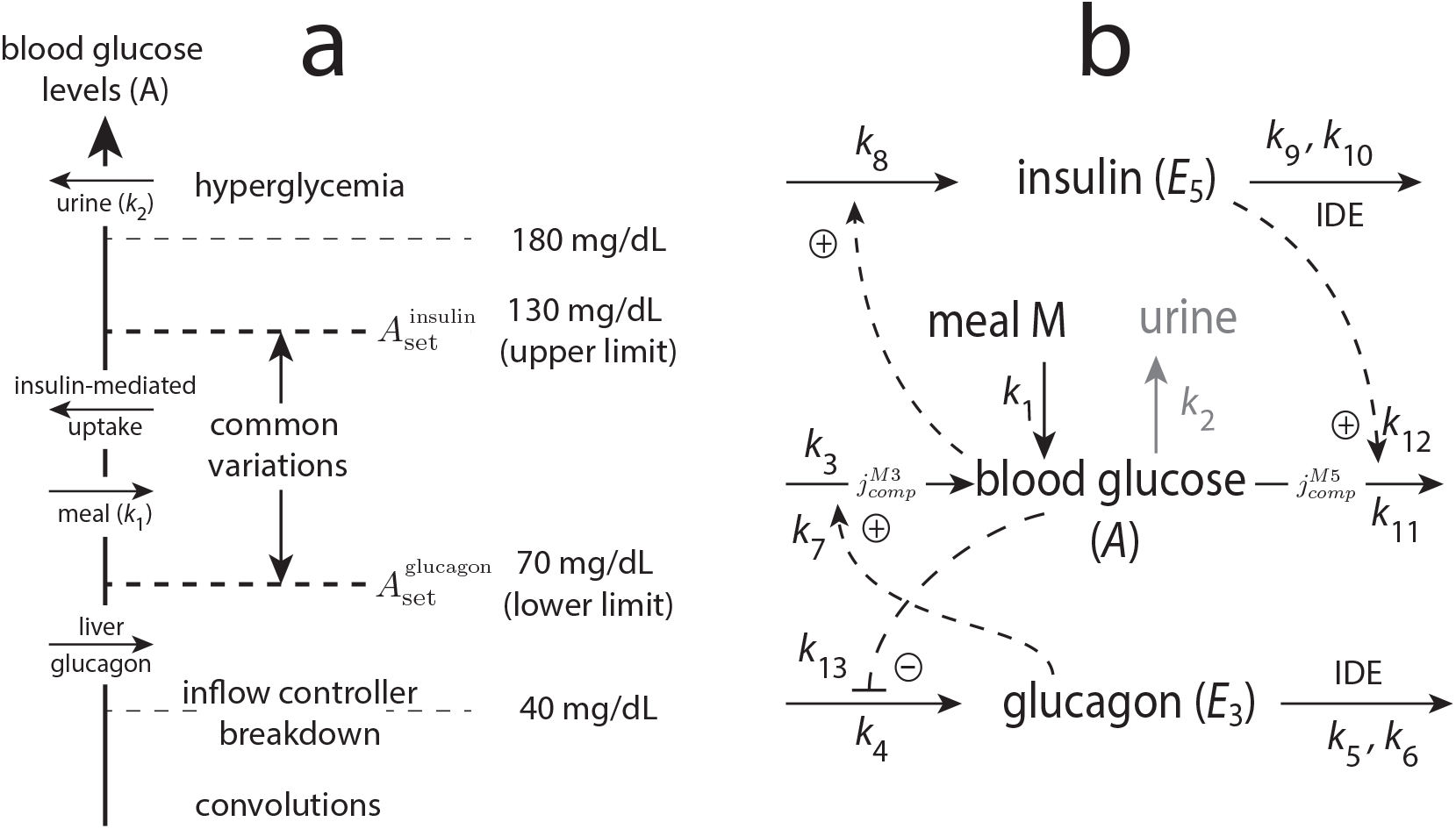
Blood glucose homeostasis described by a two-setpoint model. Panel a: Diagram of blood sugar levels based on Fig 1 in Cannon (1929). Common glucose variations, anticipated setpoints, glucose inflow/outflow fluxes, and areas of controller breakdown are indicated. 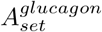 and 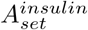 are the respective setpoints of inflow and outflow controllers in panel b. Perturbation *k*_1_ refers to the release of glucose by a meal. Since uptake of glucose into the organism’s various tissues is dependent on insulin, *k*_2_ is set to zero for healthy individuals. However, at high blood glucose levels glucose is excreted through the urine. The insulin-independent uptake of blood glucose by the kidneys and its recycling back into the blood stream is ignored. Glucagon activates the inflow of glucose into the blood by the liver when blood sugar levels are low. Panel b: Combined M3-M5 controller to describe blood glucose homeostasis. Degradations of insulin (*E*_5_) and glucagon (*E*_1_) are based on Michaelis-Menten kinetics by insulin degrading enzyme (IDE, insulysin). Compensatory flux 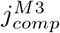 (see Table S25 in `Sup1.pdf`) refers to the glucagon-based inflow of glucose into the blood from the liver, while compensatory flux 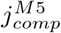 is related to the insulin-based incorporation of glucose into various tissues such as muscles, skin and liver. When blood glucose levels are high some of the glucose is excreted through the urine (outlined in gray).

To test a two-setpoint model the M3-M5 combined controller is used as a candidate since these feedback structures are close to that of insulin and glucagon. Rate equations are found in Table S25 (`Sup1.pdf`) and in the M3-M5 part of `Sup2.pdf`. To avoid unrealistically high settling times as in the M1-M5 pair (see Fig 9) the isolated control mode 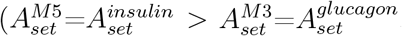, Table 1, M3-M5, blue entry) is used. *A* represents blood glucose concentration, *E*_3_ is glucagon, and *E*_5_ refers to insulin (Fig 25b). Setpoints 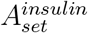 and 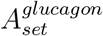

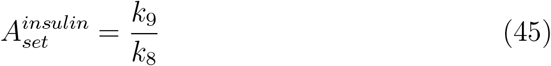

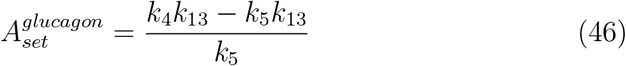

are put to respectively 10 mM (130 mg/dL) and 4 mM (70 mg/dL) blood glucose. Although calculations will refer to blood glucose level in the mM range other parts of the model, i.e. hormone concentrations and the time scale still need to be considered as qualitative properties with arbitrary units. The generation of glucose from a taken meal *M* is formulated as a simple first-order reaction with respect to *M* (Fig 25b), i.e. the conversion of *M* into glucose is written as:

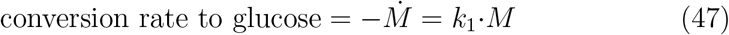

Fig 26 shows the influence of four parameters on blood glucose levels. Panel a indicates how different glucose release rates from a given meal M affect blood glucose profiles. The findings that a slower glucose release lowers the blood glucose amplitude after intake of the meal fits well with experimental results using continuous glucose profiles in nondiabetic individuals (see Fig 5 in Freckmann et al. (2007)).

**Figure 26.**
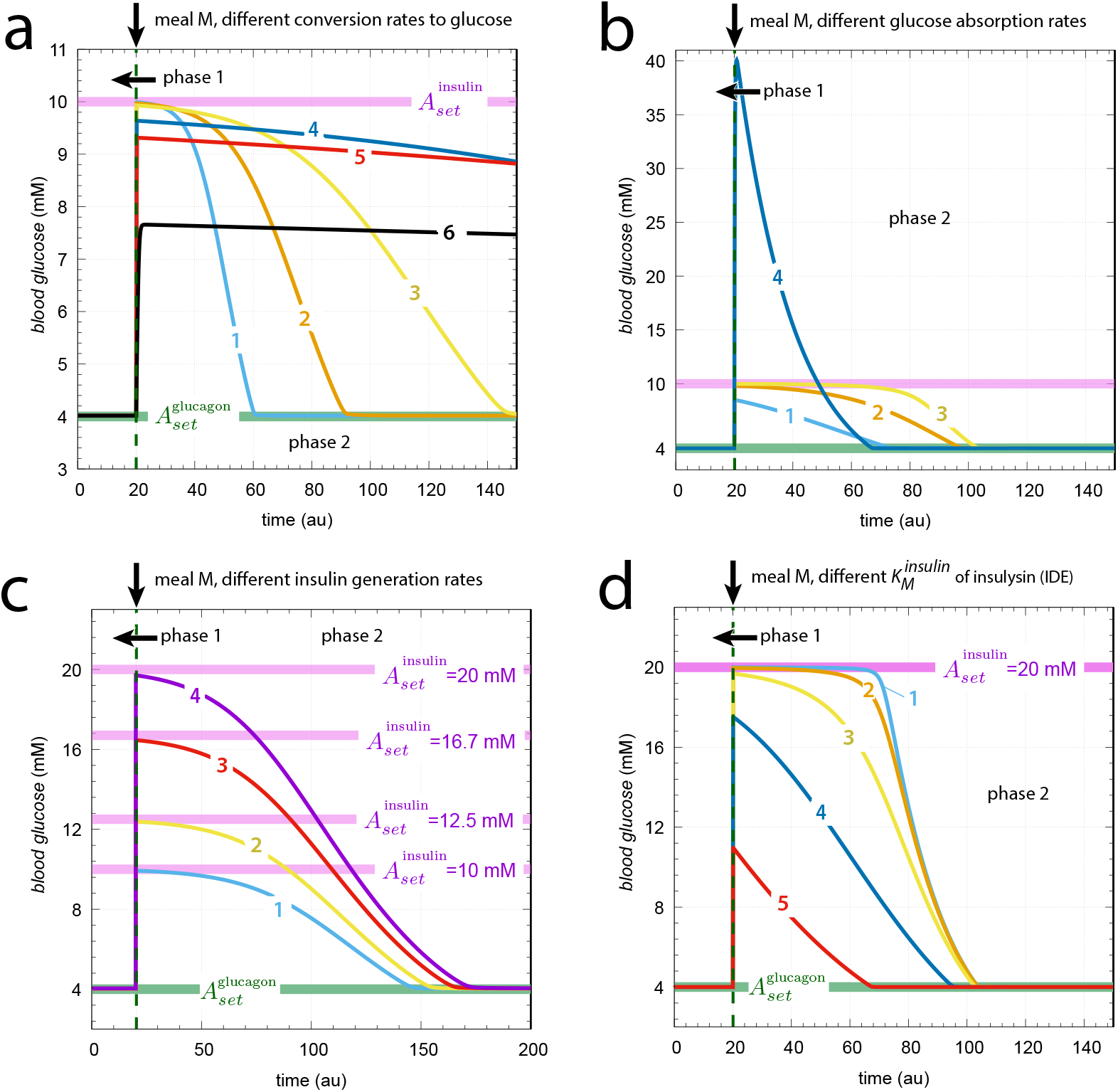
Parameters affecting blood glucose homeostasis in the M3-M5 model under isolated control. Panel a: Influence of a meal’s glucose release rate (Eq 47) on blood glucose levels. Phase 1: fasting conditions with *k*_1_=0.0. Starting phase 2 the meal is added at t=20 (vertical arrow) with an equivalent of 5M (5 × 10^3^ mM) glucose. Curves 1-6 relate to *k*_1_ values of respectively 0.2, 0.1, 0.05, 0.01, 0.005, and 0.001 inverse time units. Rate constants (phase 1 and phase 2): *k*_2_=0.0, *k*_3_=200.0, *k*_4_=400.0, *k*_5_=9.756098, *k*_6_=1.0 × 10^−6^, *k*_7_=0.1, *k*_8_=100.0, *k*_9_=1.0 × 10^3^, *k*_10_=1.0 × 10^−4^, *k*_11_=200.0, *k*_12_=*k*_13_=0.1. Initial concentrations: *A*_0_ (blood glucose)=4.0 mM, *E*_1,0_ (glucagon)=2.7 × 10^−4^, *E*_5,0_ (insulin)=6.7 × 10^−5^. Panel b: Influence of insulin-based glucose absorption rates *j*_11_ into the organism. Phase 1: fasting conditions with *k*_1_=0.0 and *k*_2_=1.0. Phase 2: *k*_1_=0.05 (added meal), *k*_2_=1.0. Curves 1-4 have *k*_11_ values of 5.0 × 10^3^, 5.0 × 10^2^, 50.0, and 5.0, respectively. Other rate constants and initial concentrations as in panel a. Panel c: Decrease of insulin generation rates increase insulin’s glucose setpoint. Phase 1 as in panel a. Phase 2: Rate constants and initial concentrations as in panel a, except *k*_1_=0.05. The *k*_8_ values of curves 1-4 are 100.0, 80.0, 60.0, and 50.0, respectively. Insulin-based setpoints are calculated by Eq 45. Panel d: Influence of 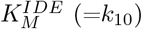 of insulin degrading enzyme (IDE, insulysin) on blood glucose levels. Phases 1 and 2 as in panel b, except varying values of *k*_10_ 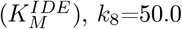, and *k*_11_=200.0. Curves 1-5 have *k*_10_ values of 1.0 × 10^−6^, 1.0 × 10^−5^, 1.0 × 10^−4^, 1.0 × 10^−3^, and 1.0 × 10^−2^, respectively. Other rate constants and initial concentrations as in panel b. See Python code for details.

Fig 26b shows the effect when insulin-based glucose incoorporation decreases (curves 1-4). The process involves an insulin-induced translocation of the glucose transporter GLUT4 Klip et al. (2019) to the cell surface such that glucose can enter the cell. Impairment of this process results in dysregulation of glucose homeostasis, insulin resistance (van Gerwen et al., 2023), and, as indicated in Fig 26b, to increased glucose levels.

An interesting prediction of the insulin (M5) part of the model is the increase of 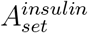 when production of insulin by decreased *k*_8_ values is reduced (Fig 26c). Although glucose levels barely reach 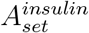 the insulin-related setpoint has a clear effect on the glucose amplitude after a meal.

Fig 26d shows the effect of different *K*_*M*_ values of the insulin degrading enzyme (IDE, insulysin). This *K*_*M*_ is of importance to define integral control and 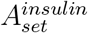 under ideal (low *K*_*M*_) conditions (curves 1 and 2, see also section 2.5). As *K*_*M*_ increases robust perfect adaptation will be lost and controller accuracy decreases (Drengstig et al., 2012a). The enzyme data base BRENDA (https://www.brenda-enzymes.org) has listed about 20 *K*_*M*_ entries for IDE with respect to insulin binding. For all organisms, including human IDE, the listed *K*_*M*_ values are relative low and lie in the range between 1 × 10^−5^ mM to 1 × 10^−4^ mM. A *K*_*M*_ of 1 × 10^−5^ mM is roughly 10 times higher than the average post meal insulin concentration (2h after the meal), which has been measured to about 152 ± 31.7 *µ*IU/mL ≈ 9×10^−7^ mM (Shulka et al., 2015). This indicates that the binding between insulin and IDE is relative tight, but IDE’s *K*_*M*_ is too high in comparison to insulin concentrations to obtain integral control/robust perfect adaptation as indicated by curves 1 or 2 in panel d. Curves 3-5 appear to be a more appropriate description for nondiabetic individuals.

In diabetics, ambulatory glucose profiles show occasional constant glucose levels (see Saturday 02 and 09 daily glucose profiles in Fig 1 of Johnson et al. (2019)), which may be interpreted that a setpoint has been reached when, due to medication, insulin levels have become tentatively higher than IDE’s *K*_*M*_. Since plateaus are also occasionally reached at lower glucose levels there appears to be a complex interplay between medication-induced insulin levels and related setpoint changes, and how fast glucose is generated from a meal.

The conclusion that is drawn from this relative simple model is: (i) There appears to be two setpoints, each relating to insulin and glucagon based feedback loops. In nondiabetic individuals it is 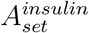 and 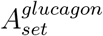 which are regulated. (ii) The adaptation to 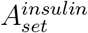 is not robustly perfect as glucose level are kept below 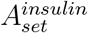. The reason for this behavior is that insulin levels are lower than IDE’s *K*_*M*_. This allows for the incorporation of glucose into the organism’s various tissues, but avoiding toxic blood glucose levels.

(iii) Diabetic individuals who have a lower insulin production rate will have a higher 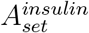 setpoint and blood glucose levels are therefore accordingly higher. However, an increased glucose setpoint may still be defended as ambulatory glucose profiles appear to indicate (Johnson et al., 2019). Medication, i.e. addition of insulin, will temporarily lower the setpoint.

#### 4.2.1. Somatostatin

While insulin and glucagon are considered to be the main drivers of blood glucose homeostasis somatostatin (SST), which is released by pancreatic delta cells and certain neurons in the brain, is recognized as an additional regulator of blood glucose levels. Somatostatin is known to inhibit both insulin and glucagon, as well as many other regulatory peptides (Bloom and Polak, 1987). Due to its inhibitory role on both insulin and glucagon somatostatin should have an effect on the two setpoints determined by the insulin and glucagon negative feedback loops.

The rate equation for insulin (*E*_5_) under steady state condition and SST inhibition can be written as

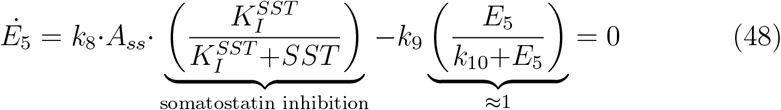

where 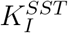 is somatostatin’s inhibition constant, *SST* denotes somatostatin concentration, and 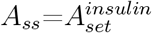 describes the glucose setpoint by the insulinbased M5 feedback. Solving for *A*_*ss*_ we get

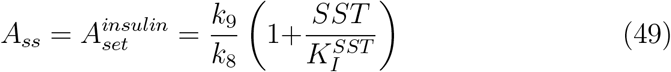

The influence of somatostatin inhibition on 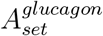 can be described similarly by using the rate equation for glucagon (*E*_3_), i.e.

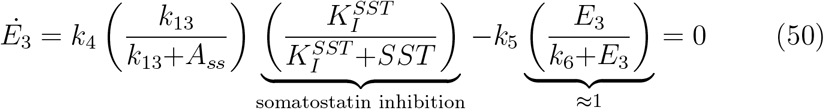

with 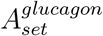 being

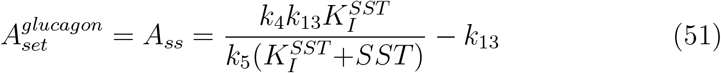

Considering Eqs 49 and 51 we see that increased somatostatin levels should increase 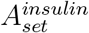, but decrease 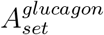. Fig 27 shows the effect of increasing somatostatin levels on 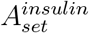 and 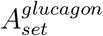.

**Figure 27.**
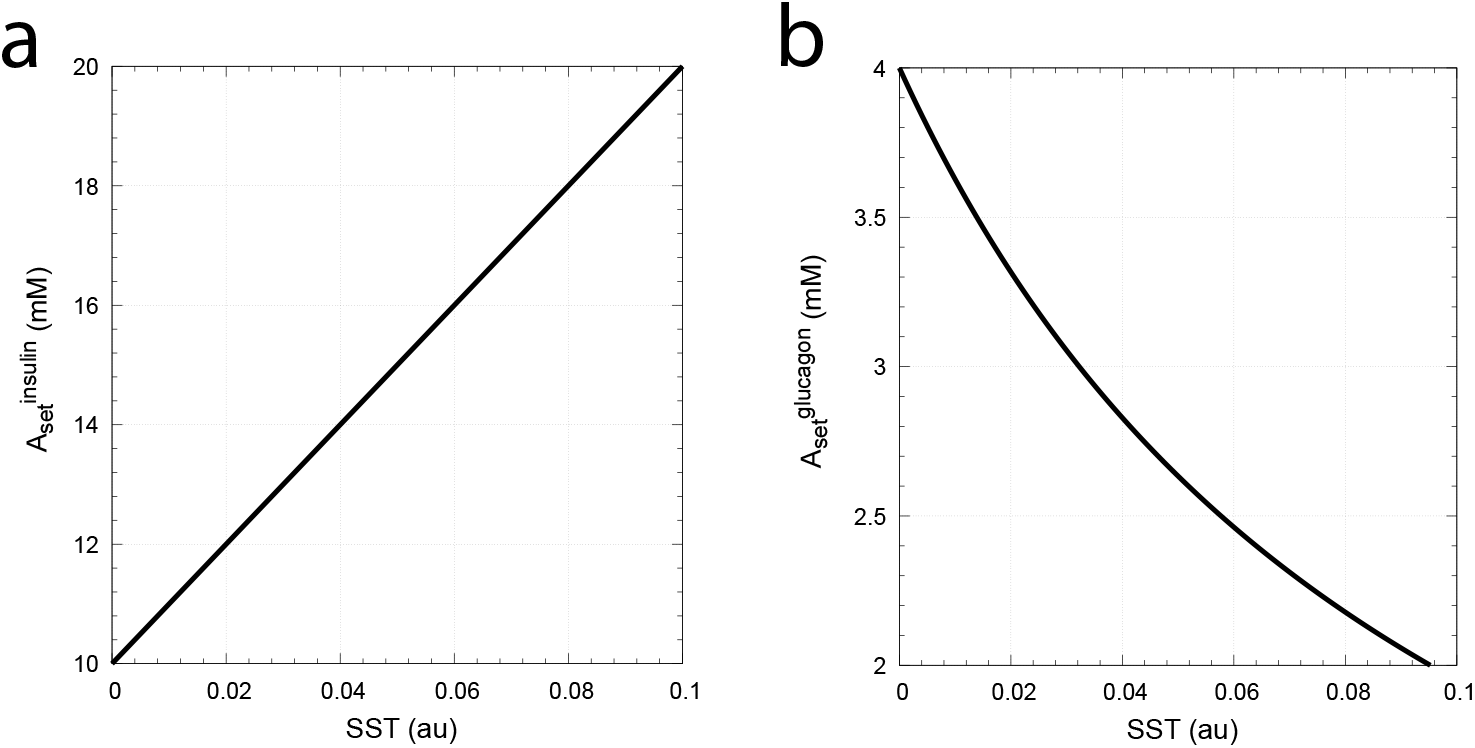
Increasing somatostatin levels (SST) lead to an increase and decrease of respectively 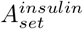 and 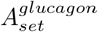. Panels a and b show respectively Eq 49 and Eq 51. Parameter values are the same as in Fig 26 with additional 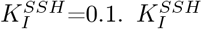 and SST levels are in arbitrary units.

The two-setpoint model thereby qualitatively predicts that under fasting conditions (blood sugar levels are determined by 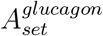) a knockout of SST should lead to an increase in glucose steady states, while during non-fasting conditions (blood sugar levels are determined by 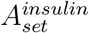) a SST knockout should decrease 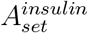 and glucose levels.

Huang et al. (2024) indeed found that in non-fasting *Sst* -Cre^TG*/*TG^ mice (homozygous transgenic mice which do not express somatostatin) a significant decrease in steady state glucose levels is observed. The authors concluded that the (single) glycemic setpoint in absence of somatostatin is decreased.

Concerning the effect of somatostatin on basal glucose levels 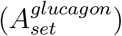 in fasting mice no direct results have been found in the literature. However, Xu et al. (2020) found that glucagon release from isolated perfused rat pancreas becomes high in presence of somatostatin blockers and that glucose lost its inhibitory effect on glucagon release in the presence of somatostatin blockers. Thus, somatostatin appears to be an essential part in the glucoseinduced inhibition mechanism of glucagon release. As a consequence, basal glucose levels 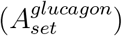 should increase when regulation by somatostatin is absent.

## 5. Summary and outlook

The subject of this work has been the study of two combined antagonistic (idealized) controllers with integral control which are (flatly) coupled through a common controlled variable *A*. The appearance of delegated and isolated control modes came to the author quite as a surprise. For each of the two control modes there are four instances with negative windup (blue entries in Table 1). As these combinations exhibit short settling times they appear to be good candidates to model antagonistic biochemical control systems. Whether metastable control or regulation by positive windup (yellow or red entries in Table 1 play a role in physiological regulation is not known.

Homeostatic systems are often considered to be related to single-loop negative feedbacks with only one setpoint. As shown by the above examples a possible way to understand rheostasis or antagonistic regulation like blood glucose is due to the interaction between two combined controllers. However, such an approach can be extended to include multiple feedbacks. For example, the use of three interlocked controller motifs (Ruoff and Nishiyama, 2020) can mimic the gradual change in p53 regulation for variable DNA-damaging stress. Although the p53 dynamics and the resulting setpoint and frequency changes are now quite complex, each of the three individual negative feedback controllers act as ‘building blocks’ in the overall homeostatic regulation of p53. Such an approach, i.e. the use of multiple combined controllers and how they interact appears to be an interesting avenue to understand homeostatic regulation in Cannon’s spirit.

## Glossary

A brief description of the concepts used in the paper.

### Controller or controller motif

A negative feedback arrangement.

### Controlled variable

The variable which is under homeostatic control with one or several setpoints.

### Controller variable or manipulated variable

The controller or manipulated variable generates a compensatory flux to oppose a perturbation applied on the controlled variable.

### High affinity kinetics

In high affinity kinetics a catalyzing enzyme binds tightly to its substrate with a low Michaelis (dissociation) constant *K*_*M*_.

### IDE

Insulin degrading enzyme. Also named insulysin.

### Integral control or integral feedback

In integral control the controller variable and its compensatory flux is related to the integrated error of the controller, i.e. the integrated difference between the actual value of the controlled variable (or a function of it) and the controller’s setpoint (or a reference). Integral feedback leads to perfect adaptation to a controller’s setpoint. Different kinetic mechanisms can result in integral control.

### Low affinity kinetics

In low affinity kinetics a catalyzing enzyme binds loosely to its substrate with a large Michaelis (dissociation) constant *K*_*M*_. Resembles often first-order or close to first-order kinetics.

### Relaxation or settling time

The time to reach a steady state after a perturbation has been applied.

### Reaction order

In a rate equation, the reaction order with respect to a chemical species is the exponent for that species.

### Steady state

The steady state of a compound *A* is reached when its concentration does not change, i.e. the time derivative 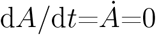.

### SST

Somatostatin.

### Windup

The continuous increase or decrease of a controller variable.

