## Supplementary material for "Flux organizations and control modes in antagonistically combined negative feedback loops": Python_ReadMe.txt

Running the Python scripts: 
=========================== 
The python codes were tested on a MacPro (Late 2013) with MacOS 12.7.6 using
python version 3.13.9.

Beside python, you need to have installed: numpy (https://numpy.org/), matplotlib (https://matplotlib.org/), and scipy (https://scipy.org/). 

To run the scripts point the terminal (or the CMD-prompt in Windows) to the appropriate folder 
which contains the scripts and execute one by writing:

python filename.py

where 'filename' is the name of the script you want to execute. This will create the graphical output. A plot in pdf format will be saved in the local 'plots' folders.


 

 
 
 
