## Supplementary material for "Flux organizations and control modes in antagonistically combined negative feedback loops": E5-time-plot_fig26d.py

```
import numpy as np
import matplotlib.pyplot as plt
from scipy.integrate import odeint
import csv
import os, sys

p1k1 = 0.0e0     #k1 during phase 1
p2k1 = 0.05      #k1 during phase 2, curve 1

p1k2 = 1.0   #k2 during phase 1
p2k2 = 1.0   #k2 during phase 2

##################### definition of model ######################
def model(x,t,vark1,vark2): #vark1, vark2: the four step perturbations in k1 and k2
                                  
    #parameters/constants:                                                                                            
    k1 = vark1 #step perturbations in k1
    k2 = vark2 #step perturbations in k2                             
    k3 = 2.0e2                                                
    k4 = 400.0                     
    k5 = 9.756098                                              
    k6 = 1.0e-6                                                 
    k7 = 0.1  
    k8 = 50.0
    k9 = 1000.0
    
    ##################### k10 values curves 1-5 ###############
    k10= 1.e-6 #curve 1
    # k10= 1.e-5 #curve 2
    # k10= 1.e-4 #curve 3
    # k10= 1.e-3 #curve 4
    # k10= 1.e-2 #curve 5
    ###########################################################   
   
    k11=2.e2   
    k12=0.1
    k13=0.1
    
    # assign each ODE to a vector:                                                         
    A = x[0]   # A (blood glucose)                                           
    E3 = x[1]  # E3 (glucagon)
    E5 = x[2]  # E5 (insulin)
    M = x[3]   # M  (meal)
                                      
    #define each ODE:                                                     
    dAdt=k1*M-k2*A+k3*E3/(k7+E3)-k11*A*E5/(k12+E5)    # rate equation for A  
    dE3dt=k4*k13/(k13+A)-k5*E3/(k6+E3)                # rate equation for E3
    dE5dt=k8*A-k9*E5/(k10+E5)                         #rate equation for E5
    dMdt=-k1*M                                        # rate equation for M
    return [dAdt, dE3dt, dE5dt, dMdt]                                   
###########################################################    
    
x0 = [4.01, 2.7e-4, 6.7e-5, 5.e3] # initial concentrations of A, E3, E5, and M

# declare a time vector (time window):
t1=np.linspace(0,20,2000)     #phase 1 time array
t2=np.linspace(20,180,18000)  #phase 2 time array
t = np.concatenate((t1,t2))   #total time array, phase 1 + phase 2

    
asetm5 = []
asetm3 = []
for i in range(0,20000):
    asetm5.append(20.0)
    asetm3.append(4.0)


#solve model for phases 1, 2, 3, and 4:
y = odeint(model, x0, t1, args=(p1k1,p1k2,)) #solve for phase 1
x1 = y[-1]       #get last element of array y; use as initial concentrtion for next phase calculation
# print (y[-1])  #print last element of array y 
y = np.concatenate((y, odeint(model, x1, t2, args=(p2k1,p2k2,)))) #solve for phase 2 and concatenate with phase 1  


A = y[:,0]   # A-array
E3 = y[:,1]  # E3-array
E5 = y[:,2]  # E5-array
M = y[:,3]   # M-array


#plot E5 (insulin) versus time:

plt.yscale("log")
plt.title("Fig 26d, $E_5$-trace; $A_{set}^{M5}$=$A_{set}^{insulin}$=20.0mM, $A_{set}^{M3}$=$A_{set}^{glucagon}$=4.0mM")
plt.plot(t,E5,label = 'E5', color='darkviolet', linewidth=3.0)
plt.tick_params(axis='y', colors='darkviolet', which='both')
plt.xlabel("time (au)")
plt.ylabel("$E_5$ (insulin) (au)", color='darkviolet')
plt.grid()
plt.xlim(0.0,150.0)
# plt.ylim(1.e-1,1.e8)
plt.axvline(x = 20, color = 'black', label = 'end phase 1', linestyle='dashed', linewidth=1.5)
plt.savefig("plots/E5-time-plot_fig26d.pdf") #saves pdf file of E5-time plot in local folder 'plots'.
plt.show()
```
