## Supplementary material for "Flux organizations and control modes in antagonistically combined negative feedback loops": E2-time-plot_figS6.py

```
import numpy as np
import matplotlib.pyplot as plt
from scipy.integrate import odeint
import csv
import os, sys

p1k1 = 3.7e3  #k1 during phase 1
p2k1 = 3.7e3  #k1 during phase 2
p3k1 = 5.0e3  #k1 during phase 3
p4k1 = 1.7e4  #k1 during phase 4

p1k2 = 4.2e3   #k2 during phase 1
p2k2 = 9.6e3  #k2 during phase 2
p3k2 = 2.e3  #k2 during phase 3
p4k2 = 2.e3  #k2 during phase 4

##################### definition of model ######################
def model(x,t,vark1,vark2): #vark1, vark2: the four step perturbations in k1 and k2
                                  
    #parameters/constants:                                                                                            
    k1 = vark1 #step perturbations in k1
    k2 = vark2 #step perturbations in k2                             
    k3 = 7.0e3                                                
    k4 = 10.0                     
    k5 = 10.0                                               
    k6 = 1.0e-6                                                 
    k7 = 0.1  
    k8 = 10.0
    k9 = 210.0
    k10= 1.e-6
    k11=7.e3
    k12=0.1
    k13=0.1
    
    # assign each ODE to a vector:
                                                          
    A = x[0]   # A                                            
    E2 = x[1]  # E2
    E7 = x[2]  # E7
                                      
    #define each ODE:                                                     
    dAdt=k1-k2*A+k3*k7/(k7+E2)-k11*A*E7/(k12+E7)    # rate equation for A  
    dE2dt=k4*A-k5*E2/(k6+E2)                        # rate equation for E2
    dE7dt=k8-k9*k13*E7/((k10+E7)*(k13+A))           # rate equation for E7
    return [dAdt, dE2dt, dE7dt]                                   
###########################################################    
    
x0 = [1.0, 1.3, 1.e-6] # initial concentrations of A, E2, and E7

# declare a time vector (time window):
t1=np.linspace(0,2000,200000)    #phase 1 time array
t2=np.linspace(2000,4000,200000)  #phase 2 time array
t3=np.linspace(4000,6000,200000)  #phase 3 time array
t4=np.linspace(6000,8000,200000)  #phase 4 time array
t = np.concatenate((t1,t2,t3,t4)) #total time array, phase 1 + phase 2 + phase 3 + phase 4

    
asetm7 = []
asetm2 = []
for i in range(0,800000):
    asetm7.append(2.0)
    asetm2.append(1.0)


#solve model for phases 1, 2, 3, and 4:
y = odeint(model, x0, t1, args=(p1k1,p1k2,)) #solve for phase 1
x1 = y[-1]       #get last element of array y; use as initial concentrtion for next phase calculation
# print (y[-1])  #print last element of array y 
y = np.concatenate((y, odeint(model, x1, t2, args=(p2k1,p2k2,)))) #solve for phase 2 and concatenate with phase 1  
x2 = y[-1] #find last concentrations; use as initial concs in the next phase calculation                    
y = np.concatenate((y, odeint(model, x2, t3, args=(p3k1,p3k2,)))) #solve for phase 3 and concatenate with phases 1 and 2  
x3 = y[-1] #find last concentrations; use as initial concs in the next phase calculation  

y = np.concatenate((y, odeint(model, x3, t4, args=(p4k1,p4k2,)))) #solve for phase 4 and concatenate with other phases                                                                                                  
A = y[:,0]   # A-array
E2 = y[:,1]  # E1-array
E7 = y[:,2]  # E5-array


    
#plot E2 versus time:
# plt.xscale("log")
plt.yscale("log")
plt.title("Fig S6, $E_2$-trace; $A_{set}^{M7}$=2.0, $A_{set}^{M2}$=1.0")
# plt.ticklabel_format(axis='y', style='sci',scilimits=(0,0))
plt.tick_params(axis='y', colors='teal', which='both')
plt.plot(t,E2,label = 'E2', color='teal', linewidth=3.0)
plt.xlabel("time (au)")
plt.ylabel("$E_2$ (au)", color='teal')
plt.grid()
plt.xlim(0.0,8000)
plt.ylim(1.e-2,1.e5)
plt.axvline(x = 2000, color = 'black', label = 'end phase 1', linestyle='dashed', linewidth=1.5)
plt.axvline(x = 4000, color = 'black', label = 'end phase 2', linestyle='dashed', linewidth=1.5)
plt.axvline(x = 6000, color = 'black', label = 'end phase 3', linestyle='dashed', linewidth=1.5)
plt.savefig("plots/E2-time-plot_figS6.pdf") #saves pdf file of E2-time plot in local folder 'plots'.
plt.show()
```
