## Supplementary material for "Flux organizations and control modes in antagonistically combined negative feedback loops": A-time-plot_figS14.py

```
import numpy as np
import matplotlib.pyplot as plt
from scipy.integrate import odeint
import csv
import os, sys

p1k1 = 5.0e3  #k1 during phase 1
p2k1 = 2.5e3  #k1 during phase 2
p3k1 = 1.25e4  #k1 during phase 3
p4k1 = 5.0e3  #k1 during phase 4

p1k2 = 1.0e3   #k2 during phase 1
p2k2 = 4.0e3  #k2 during phase 2
p3k2 = 4.e3  #k2 during phase 3
p4k2 = 8.e3  #k2 during phase 4

##################### definition of model ######################
def model(x,t,vark1,vark2): #vark1, vark2: the four step perturbations in k1 and k2
                                  
    #parameters/constants:                                                                                            
    k1 = vark1 #step perturbations in k1
    k2 = vark2 #step perturbations in k2                             
    k3 = 5.0e3                                                
    k4 = 100.0                     
    k5 = 1100.0                                               
    k6 = 1.0e-6                                                 
    k7 = 0.1  
    k8 = 2100.0
    k9 = 100.0
    k10= 1.e-6
    k11=5.e3
    k12=0.1
    k13=0.1
    k14=0.1
    
    # assign each ODE to a vector:
                                                          
    A = x[0]   # A                                            
    E4 = x[1]  # E4
    E8 = x[2]  # E8
                                      
    #define each ODE:                                                     
    dAdt=k1-k2*A+k3*k7/(k7+E4)-k11*A*k12/(k12+E8)    # rate equation for A  
    dE4dt=k4-k5*E4*k13/((k6+E4)*(k13+A))            # rate equation for E4
    dE8dt=k8*k14/(k14+A)-k9*E8/(k10+E8)             # rate equation for E8
    return [dAdt, dE4dt, dE8dt]                                   
###########################################################    
    
x0 = [2.0, 2.382E+04, 1.011] # initial concentrations of A, E4, and E8

# declare a time vector (time window):
t1=np.linspace(0,10,1000)    #phase 1 time array
t2=np.linspace(10,1000,99000)  #phase 2 time array
t3=np.linspace(1000,10000,900000)  #phase 3 time array
t4=np.linspace(10000,100000,9000000)  #phase 4 time array
t = np.concatenate((t1,t2,t3,t4)) #total time array, phase 1 + phase 2 + phase 3 + phase 4

    
asetm8 = []
asetm4 = []
for i in range(0,10000000):
    asetm8.append(2.0)
    asetm4.append(1.0)


#solve model for phases 1, 2, 3, and 4:
y = odeint(model, x0, t1, args=(p1k1,p1k2,)) #solve for phase 1
x1 = y[-1]       #get last element of array y; use as initial concentrtion for next phase calculation
# print (y[-1])  #print last element of array y 
y = np.concatenate((y, odeint(model, x1, t2, args=(p2k1,p2k2,)))) #solve for phase 2 and concatenate with phase 1  
x2 = y[-1] #find last concentrations; use as initial concs in the next phase calculation                    
y = np.concatenate((y, odeint(model, x2, t3, args=(p3k1,p3k2,)))) #solve for phase 3 and concatenate with phases 1 and 2  
x3 = y[-1] #find last concentrations; use as initial concs in the next phase calculation  

y = np.concatenate((y, odeint(model, x3, t4, args=(p4k1,p4k2,)))) #solve for phase 4 and concatenate with other phases                                                                                                  
A = y[:,0]   # A-array
E4 = y[:,1]  # E4-array
E8 = y[:,2]  # E8-array

#plot A versus time:
plt.xscale("log")
plt.plot(t, asetm8, color='orchid', linewidth=7.0)        # plotting AsetM8
plt.plot(t, asetm4, color='turquoise', linewidth=7.0)     # plotting AsetM4
plt.plot(t,A,label = 'A', color='black',linewidth=2.0)    # plotting A
plt.title("Fig S14, A-trace; $A_{set}^{M8}$=2.0, $A_{set}^{M4}$=1.0")
plt.xlabel("time (au)")
plt.ylabel("A (au)", color='black')
plt.grid()
plt.xlim(1.0,100000.0)
plt.ylim(0.5,3.5)
plt.axvline(x = 10, color = 'black', label = 'end phase 1', linestyle='dashed', linewidth=1.5)
plt.axvline(x = 1000, color = 'black', label = 'end phase 2', linestyle='dashed', linewidth=1.5)
plt.axvline(x = 10000, color = 'black', label = 'end phase 3', linestyle='dashed', linewidth=1.5)
plt.savefig("plots/A-time-plot_figS14.pdf") #saves pdf file of A-time plot in local folder 'plots'.
plt.show()
```
