## Supplementary material for "Flux organizations and control modes in antagonistically combined negative feedback loops": Sup1.pdf

Peter Ruoff

Department of Chemistry, Bioscience, and Environmental Engineering, University of Stavanger, Stavanger, Norway

### Supplementary material

#### Overview of all 16 combined controllers

The following 32 entries provide an overview of setpoints, windup properties, borderlines, and steady states of $A$ in the transition zones for both delegated and isolated control.

**Table S1.** Delegated control of combined M1 and M5 controllers ($A_{set}^{M1} > A_{set}^{M5}$)

| combined controllers | rate equations and compensatory fluxes | borderlines in phase diagram |
|---|---|---|
| 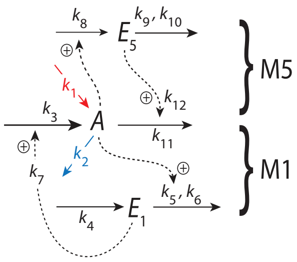 | Rate equations and set−points :<br>$\dot{A} = k_1 - k_2 A + \underbrace{k_3 \left(\frac{E_1}{k_7 + E_1}\right)}_{j_{comp}^{M1}} - \underbrace{\frac{k_{11} A \cdot E_5}{k_{12} + E_5}}_{j_{comp}^{M5}}$<br>$\dot{E_1} = k_4 - k_5 \underbrace{\left(\frac{E_1}{k_6 + E_1}\right)}_{\approx 1} A \Rightarrow A_{set}^{M1} = \frac{k_4}{k_5}$<br>$\dot{E_5} = k_8 A - k_9 \underbrace{\left(\frac{E_5}{k_{10} + E_5}\right)}_{\approx 1} \Rightarrow A_{set}^{M5} = \frac{k_9}{k_8}$<br><br>Windup conditions and generation<br>of max compensatory fluxes :<br>(i) M1 controls $A$ :<br>$A_{ss} = A_{set}^{M1} \Rightarrow \dot{E_5} > 0$<br>$\Rightarrow j_{comp,max}^{M5} = k_{11} A_{set}^{M1}$<br>− − − − − − −<br>(ii) M5 controls $A$ :<br>$A_{ss} = A_{set}^{M5} \Rightarrow \dot{E_1} > 0$<br>$\Rightarrow j_{comp,max}^{M1} = k_3$<br>− − − − − − −<br>(iii) $A_{ss}$ is in transition zone :<br>$A_{set}^{M1} > A_{ss} > A_{set}^{M5} \Rightarrow \dot{E_1} > 0 \;\&\; \dot{E_5} > 0$<br>$\Rightarrow j_{comp,max}^{M1} = k_3,$<br>$\& \; j_{comp,max}^{M5} = k_{11} A_{ss}.$<br>$A_{ss}$ is determined by $k_1$ and $k_2$, *i.e.*<br>$A_{ss} = \frac{k_1 + k_3}{k_2 + k_{11}}$ | schematic perturbation phase diagram<br>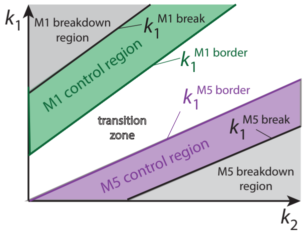<br>$k_1^{M1\,break}$ :<br>$j_{comp}^{M1} = 0 \;\&\; j_{comp}^{M5} = k_{11} \cdot A_{set}^{M1}$<br>$\Rightarrow k_1^{M1\,break} = (k_2 + k_{11}) A_{set}^{M1}$<br><br>$k_1^{M1\,border}$ :<br>$j_{comp}^{M1} = k_3 \;\&\; j_{comp}^{M5} = k_{11} \cdot A_{set}^{M1}$<br>$\Rightarrow k_1^{M1\,border} = (k_2 + k_{11}) A_{set}^{M1} - k_3$<br><br>$k_1^{M5\,border}$ :<br>$j_{comp}^{M1} = k_3 \;\&\; j_{comp}^{M5} = k_{11} \cdot A_{set}^{M5}$<br>$\Rightarrow k_1^{M5\,border} = (k_2 + k_{11}) A_{set}^{M5} - k_3$<br><br>$k_1^{M5\,break}$ :<br>$j_{comp}^{M1} = k_3 \;\&\; j_{comp}^{M5} = 0$<br>$\Rightarrow k_1^{M5\,break} = k_2 A_{set}^{M5} - k_3$ |

**Table S2.** Delegated control of combined M1 and M6 controllers ($A_{set}^{M1}>A_{set}^{M6}$)

| combined controllers | rate equations and compensatory fluxes | borderlines in phase diagram |
|---|---|---|
| 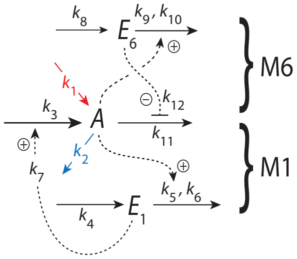<br> | Rate equations and set−points :<br>$\dot{A}=k_1-k_2A+k_3\underbrace{\left(\frac{E_1}{k_7+E_1}\right)}_{j_{comp}^{M1}}-\underbrace{\frac{k_{11}A\cdot k_{12}}{k_{12}+E_6}}_{j_{comp}^{M6}}$<br>$\dot{E}_1=k_4-k_5\underbrace{\left(\frac{E_1}{k_6+E_1}\right)}_{\approx 1}A \Rightarrow A_{set}^{M1}=\frac{k_4}{k_5}$<br>$\dot{E}_6=k_8-k_9A\underbrace{\left(\frac{E_6}{k_{10}+E_6}\right)}_{\approx 1} \Rightarrow A_{set}^{M6}=\frac{k_8}{k_9}$<br><br>Windup conditions and generation of max compensatory fluxes :<br>(i) M1 controls $A$ :<br>$A_{ss}=A_{set}^{M1} \Rightarrow \dot{E}_6<0$<br>$\Rightarrow j_{comp,max}^{M6}=k_{11}A_{set}^{M1}$<br>− − − − − − −<br>(ii) M6 controls $A$ :<br>$A_{ss}=A_{set}^{M6} \Rightarrow \dot{E}_1>0$<br>$\Rightarrow j_{comp,max}^{M1}=k_3$<br>− − − − − − −<br>(iii) $A_{ss}$ is in transition zone :<br>$A_{set}^{M1}>A_{ss}>A_{set}^{M6} \Rightarrow \dot{E}_1>0 \;\&\; \dot{E}_6<0$<br>$\Rightarrow j_{comp,max}^{M1}=k_3,$<br>$\& \; j_{comp,max}^{M6}=k_{11}A_{ss}.$<br>$A_{ss}$ is determined by $k_1$ and $k_2$, *i.e.*<br>$A_{ss}=\frac{k_1+k_3}{k_2+k_{11}}$ | schematic perturbation phase diagram<br>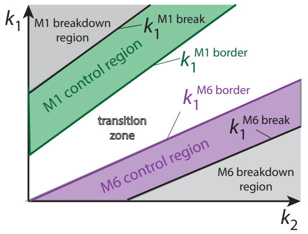<br><br><br>$k_1^{M1\,break}$ :<br>$j_{comp}^{M1}=0 \;\&\; j_{comp}^{M6}=k_{11}\cdot A_{set}^{M1}$<br>$\Rightarrow k_1^{M1\,break}=(k_2+k_{11})A_{set}^{M1}$<br><br>$k_1^{M1\,border}$ :<br>$j_{comp}^{M1}=k_3 \;\&\; j_{comp}^{M6}=k_{11}\cdot A_{set}^{M1}$<br>$\Rightarrow k_1^{M1\,border}=(k_2+k_{11})A_{set}^{M1}-k_3$<br><br>$k_1^{M6\,border}$ :<br>$j_{comp}^{M1}=k_3 \;\&\; j_{comp}^{M6}=k_{11}\cdot A_{set}^{M6}$<br>$\Rightarrow k_1^{M6\,border}=(k_2+k_{11})A_{set}^{M6}-k_3$<br><br>$k_1^{M6\,break}$ :<br>$j_{comp}^{M1}=k_3 \;\&\; j_{comp}^{M6}=0$<br>$\Rightarrow k_1^{M6\,break}=k_2A_{set}^{M6}-k_3$ |

**Table S3.** Delegated control of combined M1 and M7 controllers ($A^{M1}_{set}>A^{M7}_{set}$)

| combined controllers | rate equations and compensatory fluxes | borderlines in phase diagram |
|---|---|---|
| 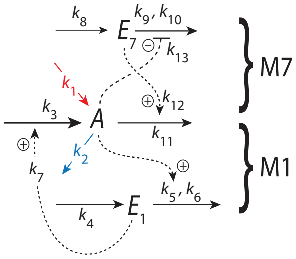<br> | Rate equations and set−points :<br>$\dot{A}=k_1-k_2A+\underbrace{k_3\left(\frac{E_1}{k_7+E_1}\right)}_{j^{M1}_{comp}}-\underbrace{\frac{k_{11}A\cdot E_7}{k_{12}+E_7}}_{j^{M7}_{comp}}$<br>$\dot{E}_1=k_4-k_5\underbrace{\left(\frac{E_1}{k_6+E_1}\right)}_{\approx 1}A \Rightarrow A^{M1}_{set}=\frac{k_4}{k_5}$<br>$\dot{E}_7=k_8-k_9\underbrace{\left(\frac{E_7}{k_{10}+E_7}\right)}_{\approx 1}\left(\frac{k_{13}}{k_{13}+A}\right)$<br>$\Rightarrow A^{M7}_{set}=\frac{k_9k_{13}-k_8k_{13}}{k_8}$<br><br>Windup conditions and generation of max compensatory fluxes :<br>(i) M1 controls $A$ :<br>$A_{ss}=A^{M1}_{set} \Rightarrow \dot{E}_7>0$<br>$\Rightarrow j^{M7}_{comp,max}=k_{11}A^{M1}_{set}$<br>− − − − − − −<br>(ii) M7 controls $A$ :<br>$A_{ss}=A^{M7}_{set} \Rightarrow \dot{E}_1>0$<br>$\Rightarrow j^{M1}_{comp,max}=k_3$<br>− − − − − − −<br>(iii) $A_{ss}$ is in transition zone :<br>$A^{M1}_{set}>A_{ss}>A^{M7}_{set} \Rightarrow \dot{E}_1>0 \,\&\, \dot{E}_7>0$<br>$\Rightarrow j^{M1}_{comp,max}=k_3,$<br>$\&\; j^{M7}_{comp,max}=k_{11}A_{ss}.$<br>$A_{ss}$ is determined by $k_1$ and $k_2$, *i.e.*<br>$A_{ss}=\frac{k_1+k_3}{k_2+k_{11}}$ | schematic perturbation phase diagram<br>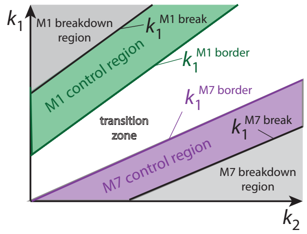<br><br>$k_1^{M1\,break}$ :<br>$j^{M1}_{comp}=0 \,\&\, j^{M7}_{comp}=k_{11}\cdot A^{M1}_{set}$<br>$\Rightarrow k_1^{M1\,break}=(k_2+k_{11})A^{M1}_{set}$<br><br>$k_1^{M1\,border}$ :<br>$j^{M1}_{comp}=k_3 \,\&\, j^{M7}_{comp}=k_{11}\cdot A^{M1}_{set}$<br>$\Rightarrow k_1^{M1\,border}=(k_2+k_{11})A^{M1}_{set}-k_3$<br><br>$k_1^{M7\,border}$ :<br>$j^{M1}_{comp}=k_3 \,\&\, j^{M7}_{comp}=k_{11}\cdot A^{M7}_{set}$<br>$\Rightarrow k_1^{M7\,border}=(k_2+k_{11})A^{M7}_{set}-k_3$<br><br>$k_1^{M7\,break}$ :<br>$j^{M1}_{comp}=k_3 \,\&\, j^{M7}_{comp}=0$<br>$\Rightarrow k_1^{M7\,break}=k_2A^{M7}_{set}-k_3$ |

**Table S4.** Delegated control of combined M1 and M8 controllers ($A^{M1}_{set}>A^{M8}_{set}$)

| combined controllers | rate equations and compensatory fluxes | borderlines in phase diagram |
|---|---|---|
| 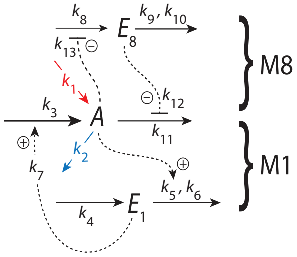<br> | Rate equations and set−points :<br>$\dot{A}=k_1-k_2A+\underbrace{k_3\left(\frac{E_1}{k_7+E_1}\right)}_{j^{M1}_{comp}}-\underbrace{\frac{k_{11}A\cdot k_{12}}{k_{12}+E_8}}_{j^{M8}_{comp}}$<br>$\dot{E}_1=k_4-k_5\underbrace{\left(\frac{E_1}{k_6+E_1}\right)}_{\approx 1}A \Rightarrow A^{M1}_{set}=\frac{k_4}{k_5}$<br>$\dot{E}_8=k_8\left(\frac{k_{13}}{k_{13}+A}\right)-k_9\underbrace{\left(\frac{E_8}{k_{10}+E_8}\right)}_{\approx 1}$<br>$\Rightarrow A^{M8}_{set}=\frac{k_8k_{13}-k_9k_{13}}{k_9}$<br><br>Windup conditions and generation<br>of max compensatory fluxes :<br>(i) M1 controls $A$ :<br>$A_{ss}=A^{M1}_{set} \Rightarrow \dot{E}_8<0$<br>$\Rightarrow j^{M8}_{comp,max}=k_{11}A^{M1}_{set}$<br>− − − − − − −<br>(ii) M8 controls $A$ :<br>$A_{ss}=A^{M8}_{set} \Rightarrow \dot{E}_1>0$<br>$\Rightarrow j^{M1}_{comp,max}=k_3$<br>− − − − − − −<br>(iii) $A_{ss}$ is in transition zone :<br>$A^{M1}_{set}>A_{ss}>A^{M8}_{set} \Rightarrow \dot{E}_1>0 \;\&\; \dot{E}_8<0$<br>$\Rightarrow j^{M1}_{comp,max}=k_3,$<br>$\&\; j^{M8}_{comp,max}=k_{11}A_{ss}.$<br>$A_{ss}$ is determined by $k_1$ and $k_2$, *i.e.*<br>$A_{ss}=\frac{k_1+k_3}{k_2+k_{11}}$ | schematic perturbation phase diagram<br>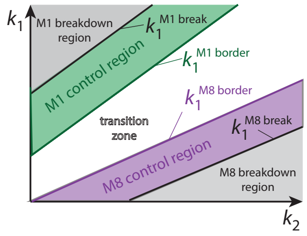<br><br><br>$k_1^{M1\,break}$ :<br>$j^{M1}_{comp}=0 \;\&\; j^{M8}_{comp}=k_{11}\cdot A^{M1}_{set}$<br>$\Rightarrow k_1^{M1\,break}=(k_2+k_{11})A^{M1}_{set}$<br><br>$k_1^{M1\,border}$ :<br>$j^{M1}_{comp}=k_3 \;\&\; j^{M8}_{comp}=k_{11}\cdot A^{M1}_{set}$<br>$\Rightarrow k_1^{M1\,border}=(k_2+k_{11})A^{M1}_{set}-k_3$<br><br>$k_1^{M8\,border}$ :<br>$j^{M1}_{comp}=k_3 \;\&\; j^{M8}_{comp}=k_{11}\cdot A^{M8}_{set}$<br>$\Rightarrow k_1^{M8\,border}=(k_2+k_{11})A^{M8}_{set}-k_3$<br><br>$k_1^{M8\,break}$ :<br>$j^{M1}_{comp}=k_3 \;\&\; j^{M8}_{comp}=0$<br>$\Rightarrow k_1^{M8\,break}=k_2A^{M8}_{set}-k_3$ |

**Table S5.** Delegated control of combined M2 and M5 controllers ($A_{set}^{M2}>A_{set}^{M5}$)

| combined controllers | rate equations and compensatory fluxes | borderlines in phase diagram |
|---|---|---|
| 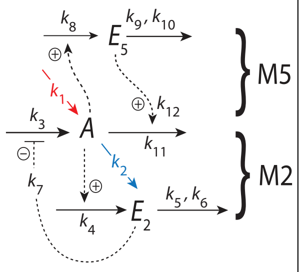<br> | Rate equations and set−points :<br>$\dot{A}=k_1-k_2A+\underbrace{k_3\left(\frac{k_7}{k_7+E_2}\right)}_{j_{comp}^{M2}}-\underbrace{\frac{k_{11}A\cdot E_5}{k_{12}+E_5}}_{j_{comp}^{M5}}$<br>$\dot{E_2}=k_4A-k_5\underbrace{\left(\frac{E_2}{k_6+E_2}\right)}_{\approx 1}\Rightarrow A_{set}^{M2}=\frac{k_5}{k_4}$<br>$\dot{E_5}=k_8A-k_9\underbrace{\left(\frac{E_5}{k_{10}+E_5}\right)}_{\approx 1}\Rightarrow A_{set}^{M5}=\frac{k_9}{k_8}$<br><br>Windup conditions and generation of max compensatory fluxes :<br>(i) M2 controls $A$ :<br>$A_{ss}=A_{set}^{M2}\Rightarrow \dot{E_5}>0$<br>$\Rightarrow j_{comp,max}^{M5}=k_{11}A_{set}^{M2}$<br>− − − − − − − −<br>(ii) M5 controls $A$ :<br>$A_{ss}=A_{set}^{M5}\Rightarrow \dot{E_2}<0$<br>$\Rightarrow j_{comp,max}^{M2}=k_3$<br>− − − − − − − −<br>(iii) $A_{ss}$ is in transition zone :<br>$A_{set}^{M2}>A_{ss}>A_{set}^{M5}\Rightarrow \dot{E_2}<0\ \&\ \dot{E_5}>0$<br>$\Rightarrow j_{comp,max}^{M2}=k_3,$<br>$\&\ j_{comp,max}^{M5}=k_{11}A_{ss}.$<br>$A_{ss}$ is determined by $k_1$ and $k_2$, *i.e.*<br>$A_{ss}=\frac{k_1+k_3}{k_2+k_{11}}$ | schematic perturbation phase diagram<br>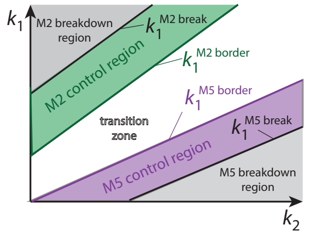<br><br>$k_1^{M2\,break}$ :<br>$j_{comp}^{M2}=0\ \&\ j_{comp}^{M5}=k_{11}\cdot A_{set}^{M2}$<br>$\Rightarrow k_1^{M2\,break}=(k_2+k_{11})A_{set}^{M2}$<br><br>$k_1^{M2\,border}$ :<br>$j_{comp}^{M2}=k_3\ \&\ j_{comp}^{M5}=k_{11}\cdot A_{set}^{M2}$<br>$\Rightarrow k_1^{M2\,border}=(k_2+k_{11})A_{set}^{M2}-k_3$<br><br>$k_1^{M5\,border}$ :<br>$j_{comp}^{M2}=k_3\ \&\ j_{comp}^{M5}=k_{11}\cdot A_{set}^{M5}$<br>$\Rightarrow k_1^{M5\,border}=(k_2+k_{11})A_{set}^{M5}-k_3$<br><br>$k_1^{M5\,break}$ :<br>$j_{comp}^{M2}=k_3\ \&\ j_{comp}^{M5}=0$<br>$\Rightarrow k_1^{M5\,break}=k_2A_{set}^{M5}-k_3$ |

**Table S6.** Delegated control of combined M2 and M6 controllers ($A^{M2}_{set}>A^{M6}_{set}$)

| combined controllers | rate equations and compensatory fluxes | borderlines in phase diagram |
|---|---|---|
| 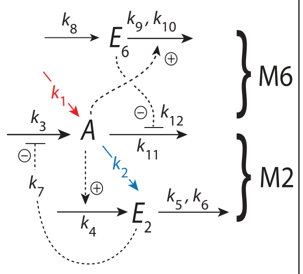 | Rate equations and set−points :<br>$\dot{A}=k_1-k_2A+\underbrace{k_3\left(\frac{k_7}{k_7+E_2}\right)}_{j^{M2}_{comp}}-\underbrace{\frac{k_{11}A\cdot k_{12}}{k_{12}+E_6}}_{j^{M6}_{comp}}$<br>$\dot{E}_2=k_4A-k_5\underbrace{\left(\frac{E_2}{k_6+E_2}\right)}_{\approx 1}\Rightarrow A^{M2}_{set}=\frac{k_5}{k_4}$<br>$\dot{E}_6=k_8-k_9A\underbrace{\left(\frac{E_6}{k_{10}+E_6}\right)}_{\approx 1}\Rightarrow A^{M6}_{set}=\frac{k_8}{k_9}$<br><br>Windup conditions and generation of max compensatory fluxes :<br>(i) M2 controls $A$ :<br>$A_{ss}=A^{M2}_{set}\Rightarrow \dot{E}_6<0$<br>$\Rightarrow j^{M6}_{comp,max}=k_{11}A^{M2}_{set}$<br>− − − − − − −<br>(ii) M6 controls $A$ :<br>$A_{ss}=A^{M6}_{set}\Rightarrow \dot{E}_2<0$<br>$\Rightarrow j^{M2}_{comp,max}=k_3$<br>− − − − − − −<br>(iii) $A_{ss}$ is in transition zone :<br>$A^{M2}_{set}>A_{ss}>A^{M6}_{set}\Rightarrow \dot{E}_2<0$ & $\dot{E}_6<0$<br>$\Rightarrow j^{M2}_{comp,max}=k_3$,<br>& $j^{M6}_{comp,max}=k_{11}A_{ss}$.<br>$A_{ss}$ is determined by $k_1$ and $k_2$, *i.e.*<br>$A_{ss}=\frac{k_1+k_3}{k_2+k_{11}}$ | schematic perturbation phase diagram<br>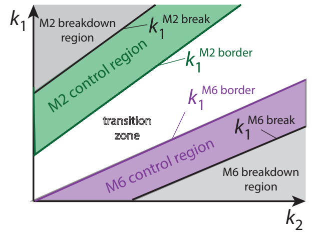<br>$k_1^{M2\,break}$ :<br>$j^{M2}_{comp}=0$ & $j^{M6}_{comp}=k_{11}\cdot A^{M2}_{set}$<br>$\Rightarrow k_1^{M2\,break}=(k_2+k_{11})A^{M2}_{set}$<br><br>$k_1^{M2\,border}$ :<br>$j^{M2}_{comp}=k_3$ & $j^{M6}_{comp}=k_{11}\cdot A^{M2}_{set}$<br>$\Rightarrow k_1^{M2\,border}=(k_2+k_{11})A^{M2}_{set}-k_3$<br><br>$k_1^{M6\,border}$ :<br>$j^{M2}_{comp}=k_3$ & $j^{M6}_{comp}=k_{11}\cdot A^{M6}_{set}$<br>$\Rightarrow k_1^{M6\,border}=(k_2+k_{11})A^{M6}_{set}-k_3$<br><br>$k_1^{M6\,break}$ :<br>$j^{M2}_{comp}=k_3$ & $j^{M6}_{comp}=0$<br>$\Rightarrow k_1^{M6\,break}=k_2A^{M6}_{set}-k_3$ |

**Table S7.** Delegated control of combined M2 and M7 controllers ($A_{set}^{M2} > A_{set}^{M7}$)

| combined controllers | rate equations and compensatory fluxes | borderlines in phase diagram |
|---|---|---|
| 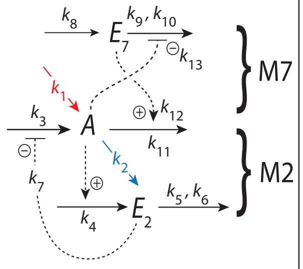<br> | Rate equations and set−points :<br>$\dot{A} = k_1 - k_2 A + \underbrace{k_3 \left(\frac{k_7}{k_7 + E_2}\right)}_{j_{comp}^{M2}} - \underbrace{\frac{k_{11} A \cdot E_7}{k_{12} + E_7}}_{j_{comp}^{M7}}$<br>$\dot{E_2} = k_4 A - k_5 \underbrace{\left(\frac{E_2}{k_6 + E_2}\right)}_{\approx 1} \Rightarrow A_{set}^{M2} = \frac{k_5}{k_4}$<br>$\dot{E_7} = k_8 - k_9 \underbrace{\left(\frac{E_7}{k_{10} + E_7}\right)}_{\approx 1} \left(\frac{k_{13}}{k_{13} + A}\right)$<br>$\Rightarrow A_{set}^{M7} = \frac{k_9 k_{13} - k_8 k_{13}}{k_8}$<br><br>Windup conditions and generation of max compensatory fluxes :<br>(i) M2 controls $A$ :<br>$A_{ss} = A_{set}^{M2} \Rightarrow \dot{E_7} > 0$<br>$\Rightarrow j_{comp,max}^{M7} = k_{11} A_{set}^{M2}$<br>− − − − − − −<br>(ii) M7 controls $A$ :<br>$A_{ss} = A_{set}^{M7} \Rightarrow \dot{E_2} < 0$<br>$\Rightarrow j_{comp,max}^{M2} = k_3$<br>− − − − − − −<br>(iii) $A_{ss}$ is in transition zone :<br>$A_{set}^{M2} > A_{ss} > A_{set}^{M7} \Rightarrow \dot{E_2} < 0$ & $\dot{E_7} > 0$<br>$\Rightarrow j_{comp,max}^{M2} = k_3$,<br>& $j_{comp,max}^{M7} = k_{11} A_{ss}$.<br>$A_{ss}$ is determined by $k_1$ and $k_2$, *i.e.*<br>$A_{ss} = \frac{k_1 + k_3}{k_2 + k_{11}}$ | schematic perturbation phase diagram<br>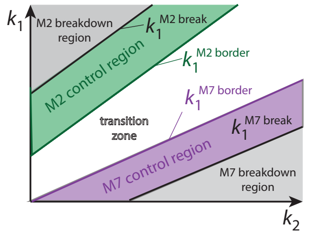<br><br>$k_1^{M2\,break}$ :<br>$j_{comp}^{M2} = 0$ & $j_{comp}^{M7} = k_{11} \cdot A_{set}^{M2}$<br>$\Rightarrow k_1^{M2\,break} = (k_2 + k_{11}) A_{set}^{M2}$<br><br>$k_1^{M2\,border}$ :<br>$j_{comp}^{M2} = k_3$ & $j_{comp}^{M7} = k_{11} \cdot A_{set}^{M2}$<br>$\Rightarrow k_1^{M2\,border} = (k_2 + k_{11}) A_{set}^{M2} - k_3$<br><br>$k_1^{M7\,border}$ :<br>$j_{comp}^{M2} = k_3$ & $j_{comp}^{M7} = k_{11} \cdot A_{set}^{M7}$<br>$\Rightarrow k_1^{M7\,border} = (k_2 + k_{11}) A_{set}^{M7} - k_3$<br><br>$k_1^{M7\,break}$ :<br>$j_{comp}^{M2} = k_3$ & $j_{comp}^{M7} = 0$<br>$\Rightarrow k_1^{M7\,break} = k_2 A_{set}^{M7} - k_3$ |

**Table S8.** Delegated control of combined M2 and M8 controllers ($A_{set}^{M2}>A_{set}^{M8}$)

| combined controllers | rate equations and compensatory fluxes | borderlines in phase diagram |
|---|---|---|
| 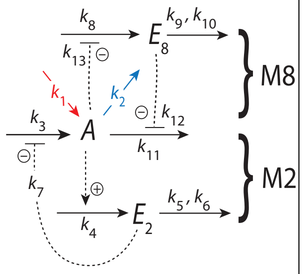 | Rate equations and set−points :<br>$\dot{A}=k_1-k_2A+\underbrace{k_3\left(\frac{k_7}{k_7+E_2}\right)}_{j_{comp}^{M2}}-\underbrace{\frac{k_{11}A\cdot k_{12}}{k_{12}+E_8}}_{j_{comp}^{M8}}$<br>$\dot{E_2}=k_4A-k_5\underbrace{\left(\frac{E_2}{k_6+E_2}\right)}_{\approx 1}\Rightarrow A_{set}^{M2}=\frac{k_5}{k_4}$<br>$\dot{E_8}=k_8\left(\frac{k_{13}}{k_{13}+A}\right)-k_9\underbrace{\left(\frac{E_8}{k_{10}+E_8}\right)}_{\approx 1}$<br>$\Rightarrow A_{set}^{M8}=\frac{k_8k_{13}-k_9k_{13}}{k_9}$<br><br>Windup conditions and generation of max compensatory fluxes :<br>(i) M2 controls $A$ :<br>$A_{ss}=A_{set}^{M2}\Rightarrow\dot{E_8}<0$<br>$\Rightarrow j_{comp,max}^{M8}=k_{11}A_{set}^{M2}$<br>− − − − − − −<br>(ii) M8 controls $A$ :<br>$A_{ss}=A_{set}^{M8}\Rightarrow\dot{E_2}<0$<br>$\Rightarrow j_{comp,max}^{M2}=k_3$<br>− − − − − − −<br>(iii) $A_{ss}$ is in transition zone :<br>$A_{set}^{M2}>A_{ss}>A_{set}^{M8}\Rightarrow\dot{E_2}<0\;\&\;\dot{E_8}<0$<br>$\Rightarrow j_{comp,max}^{M2}=k_3,$<br>$\&\; j_{comp,max}^{M8}=k_{11}A_{ss}.$<br>$A_{ss}$ is determined by $k_1$ and $k_2$, *i.e.*<br>$A_{ss}=\frac{k_1+k_3}{k_2+k_{11}}$ | schematic perturbation phase diagram<br>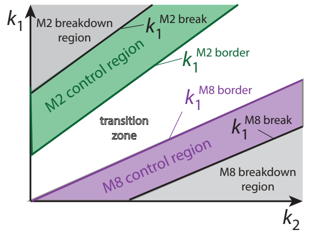<br>$k_1^{M2\,break}$ :<br>$j_{comp}^{M2}=0\;\&\;j_{comp}^{M8}=k_{11}\cdot A_{set}^{M2}$<br>$\Rightarrow k_1^{M2\,break}=(k_2+k_{11})A_{set}^{M2}$<br><br>$k_1^{M2\,border}$ :<br>$j_{comp}^{M2}=k_3\;\&\;j_{comp}^{M8}=k_{11}\cdot A_{set}^{M2}$<br>$\Rightarrow k_1^{M2\,border}=(k_2+k_{11})A_{set}^{M2}-k_3$<br><br>$k_1^{M8\,border}$ :<br>$j_{comp}^{M2}=k_3\;\&\;j_{comp}^{M8}=k_{11}\cdot A_{set}^{M8}$<br>$\Rightarrow k_1^{M8\,border}=(k_2+k_{11})A_{set}^{M8}-k_3$<br><br>$k_1^{M8\,break}$ :<br>$j_{comp}^{M2}=k_3\;\&\;j_{comp}^{M8}=0$<br>$\Rightarrow k_1^{M8\,break}=k_2A_{set}^{M8}-k_3$ |

**Table S9.** Delegated control of combined M3 and M5 controllers ($A^{M3}_{set}>A^{M5}_{set}$)

| combined controllers | rate equations and compensatory fluxes | borderlines in phase diagram |
|---|---|---|
| 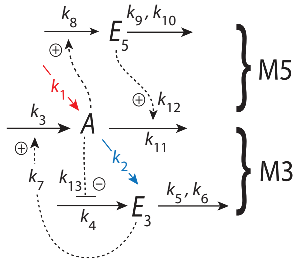  | Rate equations and set−points :<br>$\dot{A}=k_1-k_2A+\underbrace{k_3\left(\frac{E_3}{k_7+E_3}\right)}_{j^{M3}_{comp}}-\underbrace{\frac{k_{11}A\cdot E_5}{k_{12}+E_5}}_{j^{M5}_{comp}}$<br>$\dot{E}_3=k_4\left(\frac{k_{13}}{k_{13}+A}\right)-k_5\underbrace{\left(\frac{E_3}{k_6+E_3}\right)}_{\approx 1}$<br>$\Rightarrow A^{M3}_{set}=\frac{k_4k_{13}-k_5k_{13}}{k_5}$<br>$\dot{E}_5=k_8A-k_9\underbrace{\left(\frac{E_5}{k_{10}+E_5}\right)}_{\approx 1}\Rightarrow A^{M5}_{set}=\frac{k_9}{k_8}$<br>Windup conditions and generation of max compensatory fluxes :<br>(i) M3 controls $A$ :<br>$A_{ss}=A^{M3}_{set}\Rightarrow\dot{E}_5>0$<br>$\Rightarrow j^{M5}_{comp,max}=k_{11}A^{M3}_{set}$<br>− − − − − − −<br>(ii) M5 controls $A$ :<br>$A_{ss}=A^{M5}_{set}\Rightarrow\dot{E}_3>0$<br>$\Rightarrow j^{M3}_{comp,max}=k_3$<br>− − − − − − −<br>(iii) $A_{ss}$ is in transition zone :<br>$A^{M3}_{set}>A_{ss}>A^{M5}_{set}\Rightarrow\dot{E}_3>0\ \&\ \dot{E}_5>0$<br>$\Rightarrow j^{M3}_{comp,max}=k_3,$<br>$\&\ j^{M5}_{comp,max}=k_{11}A_{ss}.$<br>$A_{ss}$ is determined by $k_1$ and $k_2$, *i.e.*<br>$A_{ss}=\frac{k_1+k_3}{k_2+k_{11}}$ | schematic perturbation phase diagram<br>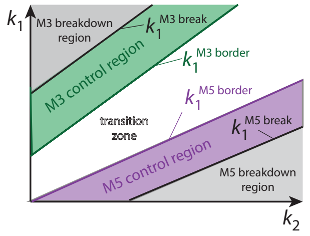 <br>$k_1^{M3\,break}$ :<br>$j^{M3}_{comp}=0\ \&\ j^{M5}_{comp}=k_{11}\cdot A^{M3}_{set}$<br>$\Rightarrow k_1^{M3\,break}=(k_2+k_{11})A^{M3}_{set}$<br>$k_1^{M3\,border}$ :<br>$j^{M3}_{comp}=k_3\ \&\ j^{M5}_{comp}=k_{11}\cdot A^{M3}_{set}$<br>$\Rightarrow k_1^{M3\,border}=(k_2+k_{11})A^{M3}_{set}-k_3$<br>$k_1^{M5\,border}$ :<br>$j^{M3}_{comp}=k_3\ \&\ j^{M5}_{comp}=k_{11}\cdot A^{M5}_{set}$<br>$\Rightarrow k_1^{M5\,border}=(k_2+k_{11})A^{M5}_{set}-k_3$<br>$k_1^{M5\,break}$ :<br>$j^{M3}_{comp}=k_3\ \&\ j^{M5}_{comp}=0$<br>$\Rightarrow k_1^{M5\,break}=k_2A^{M5}_{set}-k_3$ |

**Table S10.** Delegated control of combined M3 and M6 controllers ($A_{set}^{M3}>A_{set}^{M6}$)

| combined controllers | rate equations and compensatory fluxes | borderlines in phase diagram |
|---|---|---|
| 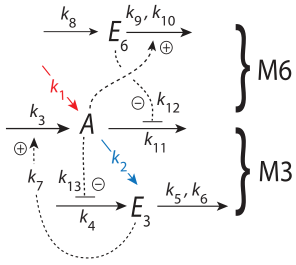<br> | Rate equations and set−points :<br>$\dot{A}=k_1-k_2A+\underbrace{k_3\left(\frac{E_3}{k_7+E_3}\right)}_{j_{comp}^{M3}}-\underbrace{\frac{k_{11}A\cdot k_{12}}{k_{12}+E_6}}_{j_{comp}^{M6}}$<br>$\dot{E}_3=k_4\left(\frac{k_{13}}{k_{13}+A}\right)-k_5\underbrace{\left(\frac{E_3}{k_6+E_3}\right)}_{\approx 1}$<br>$\Rightarrow A_{set}^{M3}=\frac{k_4k_{13}-k_5k_{13}}{k_5}$<br>$\dot{E}_6=k_8-k_9A\underbrace{\left(\frac{E_6}{k_{10}+E_6}\right)}_{\approx 1}\Rightarrow A_{set}^{M6}=\frac{k_8}{k_9}$<br><br>Windup conditions and generation of max compensatory fluxes :<br>(i) M3 controls $A$ :<br>$A_{ss}=A_{set}^{M3}\Rightarrow \dot{E}_6<0$<br>$\Rightarrow j_{comp,max}^{M6}=k_{11}A_{set}^{M3}$<br>– – – – – – –<br>(ii) M6 controls $A$ :<br>$A_{ss}=A_{set}^{M6}\Rightarrow \dot{E}_3>0$<br>$\Rightarrow j_{comp,max}^{M3}=k_3$<br>– – – – – – –<br>(iii) $A_{ss}$ is in transition zone :<br>$A_{set}^{M3}>A_{ss}>A_{set}^{M6}\Rightarrow \dot{E}_3>0\ \&\ \dot{E}_6<0$<br>$\Rightarrow j_{comp,max}^{M3}=k_3,$<br>$\&\ j_{comp,max}^{M6}=k_{11}A_{ss}.$<br>$A_{ss}$ is determined by $k_1$ and $k_2$, *i.e.*<br>$A_{ss}=\frac{k_1+k_3}{k_2+k_{11}}$ | schematic perturbation phase diagram<br>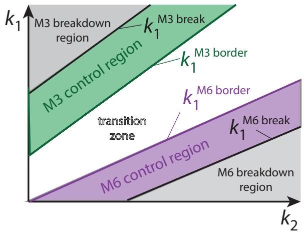<br><br><br>$k_1^{M3\,break}$ :<br>$j_{comp}^{M3}=0\ \&\ j_{comp}^{M6}=k_{11}\cdot A_{set}^{M3}$<br>$\Rightarrow k_1^{M3\,break}=(k_2+k_{11})A_{set}^{M3}$<br><br>$k_1^{M3\,border}$ :<br>$j_{comp}^{M3}=k_3\ \&\ j_{comp}^{M6}=k_{11}\cdot A_{set}^{M3}$<br>$\Rightarrow k_1^{M3\,border}=(k_2+k_{11})A_{set}^{M3}-k_3$<br><br>$k_1^{M6\,border}$ :<br>$j_{comp}^{M3}=k_3\ \&\ j_{comp}^{M6}=k_{11}\cdot A_{set}^{M6}$<br>$\Rightarrow k_1^{M6\,border}=(k_2+k_{11})A_{set}^{M6}-k_3$<br><br>$k_1^{M6\,break}$ :<br>$j_{comp}^{M3}=k_3\ \&\ j_{comp}^{M6}=0$<br>$\Rightarrow k_1^{M6\,break}=k_2A_{set}^{M6}-k_3$ |

**Table S11.** Delegated control of combined M3 and M7 controllers ($A_{set}^{M3}>A_{set}^{M7}$)

| combined controllers | rate equations and compensatory fluxes | borderlines in phase diagram |
|---|---|---|
| 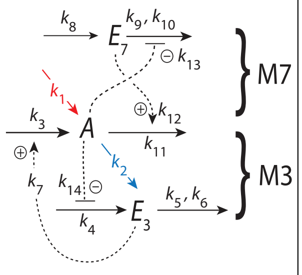<br> | Rate equations and set−points :<br>$\dot{A}=k_1-k_2A+\underbrace{k_3\left(\frac{E_3}{k_7+E_3}\right)}_{j_{comp}^{M3}}-\underbrace{\frac{k_{11}A\cdot E_7}{k_{12}+E_7}}_{j_{comp}^{M7}}$<br>$\dot{E_3}=k_4\left(\frac{k_{13}}{k_{13}+A}\right)-k_5\underbrace{\left(\frac{E_3}{k_6+E_3}\right)}_{\approx 1}$<br>$\Rightarrow A_{set}^{M3}=\frac{k_4k_{14}-k_5k_{14}}{k_5}$<br>$\dot{E_7}=k_8-k_9\left(\frac{E_7}{k_{10}+E_7}\right)\underbrace{\left(\frac{k_{13}}{k_{13}+A}\right)}_{\approx 1}$<br>$\Rightarrow A_{set}^{M7}=\frac{k_9k_{13}-k_8k_{13}}{k_8}$<br>Windup conditions and generation of max compensatory fluxes :<br>(i) M3 controls $A$ :<br>$A_{ss}=A_{set}^{M3}\Rightarrow \dot{E_7}>0$<br>$\Rightarrow j_{comp,max}^{M7}=k_{11}A_{set}^{M3}$<br>− − − − − − −<br>(ii) M7 controls $A$ :<br>$A_{ss}=A_{set}^{M7}\Rightarrow \dot{E_3}>0$<br>$\Rightarrow j_{comp,max}^{M3}=k_3$<br>− − − − − − −<br>(iii) $A_{ss}$ is in transition zone :<br>$A_{set}^{M3}>A_{ss}>A_{set}^{M7}\Rightarrow \dot{E_3}>0\ \&\ \dot{E_7}>0$<br>$\Rightarrow j_{comp,max}^{M3}=k_3,$<br>$\&\ j_{comp,max}^{M7}=k_{11}A_{ss}.$<br>$A_{ss}$ is determined by $k_1$ and $k_2$, *i.e.*<br>$A_{ss}=\frac{k_1+k_3}{k_2+k_{11}}$ | schematic perturbation phase diagram<br>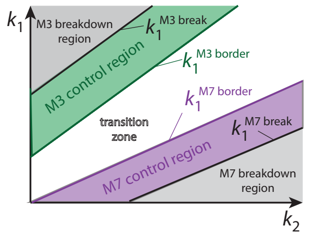<br><br>$k_1^{M3\,break}$ :<br>$j_{comp}^{M3}=0\ \&\ j_{comp}^{M7}=k_{11}\cdot A_{set}^{M3}$<br>$\Rightarrow k_1^{M3\,break}=(k_2+k_{11})A_{set}^{M3}$<br>$k_1^{M3\,border}$ :<br>$j_{comp}^{M3}=k_3\ \&\ j_{comp}^{M7}=k_{11}\cdot A_{set}^{M3}$<br>$\Rightarrow k_1^{M3\,border}=(k_2+k_{11})A_{set}^{M3}-k_3$<br>$k_1^{M7\,border}$ :<br>$j_{comp}^{M3}=k_3\ \&\ j_{comp}^{M7}=k_{11}\cdot A_{set}^{M7}$<br>$\Rightarrow k_1^{M7\,border}=(k_2+k_{11})A_{set}^{M7}-k_3$<br>$k_1^{M7\,break}$ :<br>$j_{comp}^{M3}=k_3\ \&\ j_{comp}^{M7}=0$<br>$\Rightarrow k_1^{M7\,break}=k_2A_{set}^{M7}-k_3$ |

**Table S12.** Delegated control of combined M3 and M8 controllers ($A^{M3}_{set}>A^{M8}_{set}$)

| combined controllers | rate equations and compensatory fluxes | borderlines in phase diagram |
|---|---|---|
| 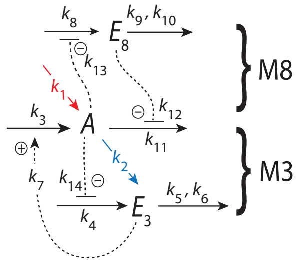<br> | Rate equations and set−points :<br>$\dot{A}=k_1-k_2A+k_3\underbrace{\left(\frac{E_3}{k_7+E_3}\right)}_{j^{M3}_{comp}}-\underbrace{\frac{k_{11}A\cdot k_{12}}{k_{12}+E_8}}_{j^{M8}_{comp}}$<br>$\dot{E}_3=k_4\left(\frac{k_{13}}{k_{13}+A}\right)-k_5\underbrace{\left(\frac{E_3}{k_6+E_3}\right)}_{\approx 1}$<br>$\Rightarrow A^{M3}_{set}=\frac{k_4k_{14}-k_5k_{14}}{k_5}$<br>$\dot{E}_8=k_8\left(\frac{k_{13}}{k_{13}+A}\right)-k_9\underbrace{\left(\frac{E_8}{k_{10}+E_8}\right)}_{\approx 1}$<br>$\Rightarrow A^{M8}_{set}=\frac{k_8k_{13}-k_9k_{13}}{k_9}$<br><br>Windup conditions and generation of max compensatory fluxes :<br>(i) M3 controls $A$ :<br>$A_{ss}=A^{M3}_{set}\Rightarrow \dot{E}_8<0$<br>$\Rightarrow j^{M8}_{comp,max}=k_{11}A^{M3}_{set}$<br>− − − − − − −<br>(ii) M8 controls $A$ :<br>$A_{ss}=A^{M8}_{set}\Rightarrow \dot{E}_3>0$<br>$\Rightarrow j^{M3}_{comp,max}=k_3$<br>− − − − − − −<br>(iii) $A_{ss}$ is in transition zone :<br>$A^{M3}_{set}>A_{ss}>A^{M8}_{set}\Rightarrow \dot{E}_3>0\ \&\ \dot{E}_8<0$<br>$\Rightarrow j^{M3}_{comp,max}=k_3,$<br>$\&\ j^{M8}_{comp,max}=k_{11}A_{ss}.$<br>$A_{ss}$ is determined by $k_1$ and $k_2$, *i.e.*<br>$A_{ss}=\frac{k_1+k_3}{k_2+k_{11}}$ | schematic perturbation phase diagram<br>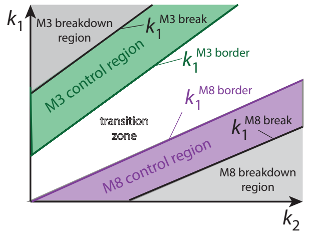<br><br><br>$k_1^{M3\,break}$ :<br>$j^{M3}_{comp}=0\ \&\ j^{M8}_{comp}=k_{11}\cdot A^{M3}_{set}$<br>$\Rightarrow k_1^{M3\,break}=(k_2+k_{11})A^{M3}_{set}$<br><br>$k_1^{M3\,border}$ :<br>$j^{M3}_{comp}=k_3\ \&\ j^{M8}_{comp}=k_{11}\cdot A^{M3}_{set}$<br>$\Rightarrow k_1^{M3\,border}=(k_2+k_{11})A^{M3}_{set}-k_3$<br><br>$k_1^{M8\,border}$ :<br>$j^{M3}_{comp}=k_3\ \&\ j^{M8}_{comp}=k_{11}\cdot A^{M8}_{set}$<br>$\Rightarrow k_1^{M8\,border}=(k_2+k_{11})A^{M8}_{set}-k_3$<br><br>$k_1^{M8\,break}$ :<br>$j^{M3}_{comp}=k_3\ \&\ j^{M8}_{comp}=0$<br>$\Rightarrow k_1^{M8\,break}=k_2A^{M8}_{set}-k_3$ |

**Table S13.** Delegated control of combined M4 and M5 controllers ($A_{set}^{M4}>A_{set}^{M5}$)

| combined controllers | rate equations and compensatory fluxes | borderlines in phase diagram |
|---|---|---|
| 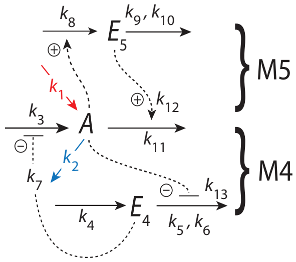<br> | Rate equations and set−points :<br>$\dot{A}=k_1-k_2A+k_3\underbrace{\left(\frac{k_7}{k_7+E_4}\right)}_{j_{comp}^{M4}}-\underbrace{\frac{k_{11}A\cdot E_5}{k_{12}+E_5}}_{j_{comp}^{M5}}$<br>$\dot{E}_4=k_4-k_5\underbrace{\left(\frac{E_4}{k_6+E_4}\right)}_{\approx 1}\cdot\left(\frac{k_{13}}{k_{13}+A}\right)$<br>$\Rightarrow A_{set}^{M4}=\frac{k_5k_{13}-k_4k_{13}}{k_4}$<br>$\dot{E}_5=k_8A-k_9\underbrace{\left(\frac{E_5}{k_{10}+E_5}\right)}_{\approx 1}\Rightarrow A_{set}^{M5}=\frac{k_9}{k_8}$<br><br>Windup conditions and generation of max compensatory fluxes :<br>(i) M4 controls $A$ :<br>$A_{ss}=A_{set}^{M4}\Rightarrow \dot{E}_5>0$<br>$\Rightarrow j_{comp}^{M5}=j_{comp,max}^{M5}=k_{11}A_{set}^{M4}$<br>− − − − − − −<br>(ii) M5 controls $A$ :<br>$A_{ss}=A_{set}^{M5}\Rightarrow \dot{E}_4<0$<br>$\Rightarrow j_{comp}^{M4}=j_{comp,max}^{M4}=k_3$<br>− − − − − − −<br>(iii) $A_{ss}$ is in transition zone :<br>$A_{set}^{M4}>A_{ss}>A_{set}^{M5}\Rightarrow \dot{E}_4<0\ \&\ \dot{E}_5>0$<br>$\Rightarrow j_{comp}^{M4}=k_3,$<br>$\&\ j_{comp}^{M5}=k_{11}A_{ss}.$<br>$A_{ss}$ is determined by $k_1$ and $k_2$, *i.e.*<br>$A_{ss}=\frac{k_1+k_3}{k_2+k_{11}}$ | schematic perturbation phase diagram<br>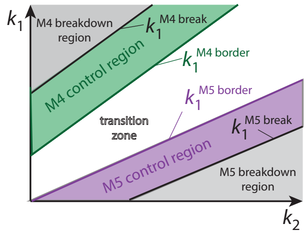<br><br>$k_1^{M4\,break}$ :<br>$j_{comp}^{M4}=0\ \&\ j_{comp}^{M5}=k_{11}\cdot A_{set}^{M4}$<br>$\Rightarrow k_1^{M4\,break}=(k_2+k_{11})A_{set}^{M4}$<br><br>$k_1^{M4\,border}$ :<br>$j_{comp}^{M4}=k_3\ \&\ j_{comp}^{M5}=k_{11}\cdot A_{set}^{M4}$<br>$\Rightarrow k_1^{M4\,border}=(k_2+k_{11})A_{set}^{M4}-k_3$<br><br>$k_1^{M5\,border}$ :<br>$j_{comp}^{M4}=k_3\ \&\ j_{comp}^{M5}=k_{11}\cdot A_{set}^{M5}$<br>$\Rightarrow k_1^{M5\,border}=(k_2+k_{11})A_{set}^{M5}-k_3$<br><br>$k_1^{M5\,break}$ :<br>$j_{comp}^{M4}=k_3\ \&\ j_{comp}^{M5}=0$<br>$\Rightarrow k_1^{M5\,break}=k_2A_{set}^{M5}-k_3$ |

**Table S14.** Delegated control of combined M4 and M6 controllers ($A^{M4}_{set}>A^{M6}_{set}$)

| combined controllers | rate equations and compensatory fluxes | borderlines in phase diagram |
|---|---|---|
| 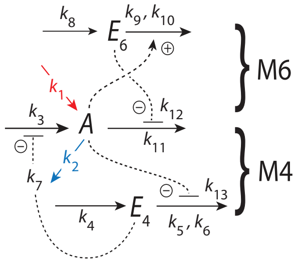 | Rate equations and set−points :<br>$\dot{A}=k_1-k_2A+\underbrace{k_3\left(\frac{k_7}{k_7+E_4}\right)}_{j^{M4}_{comp}}-\underbrace{\frac{k_{11}A\cdot k_{12}}{k_{12}+E_6}}_{j^{M6}_{comp}}$<br>$\dot{E}_4=k_4-k_5\underbrace{\left(\frac{E_4}{k_6+E_4}\right)}_{\approx 1}\cdot\left(\frac{k_{13}}{k_{13}+A}\right)$<br>$\Rightarrow A^{M4}_{set}=\frac{k_5k_{13}-k_4k_{13}}{k_4}$<br>$\dot{E}_6=k_8-k_9\underbrace{\left(\frac{E_6}{k_{10}+E_6}\right)}_{\approx 1}A\Rightarrow A^{M6}_{set}=\frac{k_8}{k_9}$<br><br>Windup conditions and generation of max compensatory fluxes :<br>(i) M4 controls $A$ :<br>$A_{ss}=A^{M4}_{set}\Rightarrow\dot{E}_6<0$<br>$\Rightarrow j^{M6}_{comp}=j^{M6}_{comp,max}=k_{11}A^{M4}_{set}$<br>− − − − − − −<br>(ii) M6 controls $A$ :<br>$A_{ss}=A^{M6}_{set}\Rightarrow\dot{E}_4<0$<br>$\Rightarrow j^{M4}_{comp}=j^{M4}_{comp,max}=k_3$<br>− − − − − − −<br>(iii) $A_{ss}$ is in transition zone :<br>$A^{M4}_{set}>A_{ss}>A^{M6}_{set}\Rightarrow\dot{E}_4<0\ \&\ \dot{E}_6<0$<br>$\Rightarrow j^{M4}_{comp}=k_3,$<br>$\&\ j^{M6}_{comp}=k_{11}A_{ss}.$<br>$A_{ss}$ is determined by $k_1$ and $k_2$, *i.e.*<br>$A_{ss}=\frac{k_1+k_3}{k_2+k_{11}}$ | schematic perturbation phase diagram<br>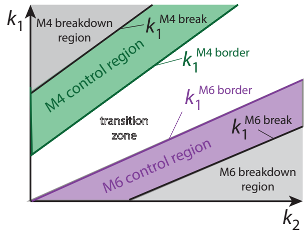<br>$k_1^{M4\,break}$ :<br>$j^{M4}_{comp}=0\ \&\ j^{M6}_{comp}=k_{11}\cdot A^{M4}_{set}$<br>$\Rightarrow k_1^{M4\,break}=(k_2+k_{11})A^{M4}_{set}$<br><br>$k_1^{M4\,border}$ :<br>$j^{M4}_{comp}=k_3\ \&\ j^{M6}_{comp}=k_{11}\cdot A^{M4}_{set}$<br>$\Rightarrow k_1^{M4\,border}=(k_2+k_{11})A^{M4}_{set}-k_3$<br><br>$k_1^{M6\,border}$ :<br>$j^{M4}_{comp}=k_3\ \&\ j^{M6}_{comp}=k_{11}\cdot A^{M6}_{set}$<br>$\Rightarrow k_1^{M6\,border}=(k_2+k_{11})A^{M6}_{set}-k_3$<br><br>$k_1^{M6\,break}$ :<br>$j^{M4}_{comp}=k_3\ \&\ j^{M6}_{comp}=0$<br>$\Rightarrow k_1^{M6\,break}=k_2A^{M6}_{set}-k_3$ |

**Table S15.** Delegated control of combined M4 and M7 controllers ($A_{set}^{M4}>A_{set}^{M7}$)

| combined controllers | rate equations and compensatory fluxes | borderlines in phase diagram |
|---|---|---|
| 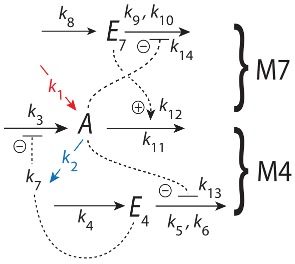<br> | Rate equations and set−points :<br>$\dot{A}=k_1-k_2A+k_3\underbrace{\left(\frac{k_7}{k_7+E_4}\right)}_{j_{comp}^{M4}}-\underbrace{\frac{k_{11}A\cdot E_7}{k_{12}+E_7}}_{j_{comp}^{M7}}$<br>$\dot{E}_4=k_4-k_5\underbrace{\left(\frac{E_4}{k_6+E_4}\right)}_{\approx 1}\cdot\left(\frac{k_{13}}{k_{13}+A}\right)$<br>$\Rightarrow A_{set}^{M4}=\frac{k_5k_{13}-k_4k_{13}}{k_4}$<br>$\dot{E}_7=k_8-k_9\underbrace{\left(\frac{E_7}{k_{10}+E_7}\right)}_{\approx 1}\left(\frac{k_{14}}{k_{14}+A}\right)$<br>$\Rightarrow A_{set}^{M7}=\frac{k_9k_{14}-k_8k_{14}}{k_8}$<br><br>Windup conditions and generation of max compensatory fluxes :<br>(i) M4 controls $A$ :<br>$A_{ss}=A_{set}^{M4}\Rightarrow \dot{E}_7>0$<br>$\Rightarrow j_{comp}^{M7}=j_{comp,max}^{M7}=k_{11}A_{set}^{M4}$<br>− − − − − − −<br>(ii) M7 controls $A$ :<br>$A_{ss}=A_{set}^{M7}\Rightarrow \dot{E}_4<0$<br>$\Rightarrow j_{comp}^{M4}=j_{comp,max}^{M4}=k_3$<br>− − − − − − −<br>(iii) $A_{ss}$ is in transition zone :<br>$A_{set}^{M4}>A_{ss}>A_{set}^{M7}\Rightarrow \dot{E}_4<0\ \&\ \dot{E}_7>0$<br>$\Rightarrow j_{comp}^{M4}=k_3,$<br>$\&\ j_{comp}^{M7}=k_{11}A_{ss}.$<br>$A_{ss}$ is determined by $k_1$ and $k_2$, *i.e.*<br>$A_{ss}=\frac{k_1+k_3}{k_2+k_{11}}$ | schematic perturbation phase diagram<br>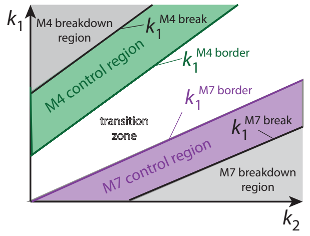<br><br>$k_1^{M4\,break}$ :<br>$j_{comp}^{M4}=0\ \&\ j_{comp}^{M7}=k_{11}\cdot A_{set}^{M4}$<br>$\Rightarrow k_1^{M4\,break}=(k_2+k_{11})A_{set}^{M4}$<br><br>$k_1^{M4\,border}$ :<br>$j_{comp}^{M4}=k_3\ \&\ j_{comp}^{M7}=k_{11}\cdot A_{set}^{M4}$<br>$\Rightarrow k_1^{M4\,border}=(k_2+k_{11})A_{set}^{M4}-k_3$<br><br>$k_1^{M7\,border}$ :<br>$j_{comp}^{M4}=k_3\ \&\ j_{comp}^{M7}=k_{11}\cdot A_{set}^{M7}$<br>$\Rightarrow k_1^{M7\,border}=(k_2+k_{11})A_{set}^{M7}-k_3$<br><br>$k_1^{M7\,break}$ :<br>$j_{comp}^{M4}=k_3\ \&\ j_{comp}^{M7}=0$<br>$\Rightarrow k_1^{M7\,break}=k_2A_{set}^{M7}-k_3$ |

**Table S16.** Delegated control of combined M4 and M8 controllers ($A^{M4}_{set}>A^{M8}_{set}$)

| combined controllers | rate equations and compensatory fluxes | borderlines in phase diagram |
|---|---|---|
| 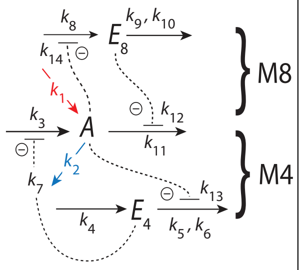<br> | Rate equations and set−points :<br>$\dot{A}=k_1-k_2A+k_3\underbrace{\left(\frac{k_7}{k_7+E_4}\right)}_{j^{M4}_{comp}}-\underbrace{\frac{k_{11}A\cdot k_{12}}{k_{12}+E_8}}_{j^{M8}_{comp}}$<br>$\dot{E}_4=k_4-k_5\underbrace{\left(\frac{E_4}{k_6+E_4}\right)}_{\approx 1}\cdot\left(\frac{k_{13}}{k_{13}+A}\right)$<br>$\Rightarrow A^{M4}_{set}=\frac{k_5k_{13}-k_4k_{13}}{k_4}$<br>$\dot{E}_8=k_8\left(\frac{k_{14}}{k_{14}+A}\right)-k_9\underbrace{\left(\frac{E_8}{k_{10}+E_8}\right)}_{\approx 1}$<br>$\Rightarrow A^{M8}_{set}=\frac{k_8k_{14}-k_9k_{14}}{k_9}$<br><br>Windup conditions and generation of max compensatory fluxes :<br>(i) M4 controls $A$ :<br>$A_{ss}=A^{M4}_{set} \Rightarrow \dot{E}_8<0$<br>$\Rightarrow j^{M8}_{comp}=j^{M8}_{comp,max}=k_{11}A^{M4}_{set}$<br>− − − − − − −<br>(ii) M8 controls $A$ :<br>$A_{ss}=A^{M8}_{set} \Rightarrow \dot{E}_4<0$<br>$\Rightarrow j^{M4}_{comp}=j^{M4}_{comp,max}=k_3$<br>− − − − − − −<br>(iii) $A_{ss}$ is in transition zone :<br>$A^{M4}_{set}>A_{ss}>A^{M8}_{set} \Rightarrow \dot{E}_4<0 \;\&\; \dot{E}_8<0$<br>$\Rightarrow j^{M4}_{comp}=k_3,$<br>$\&\; j^{M8}_{comp}=k_{11}A_{ss}.$<br>$A_{ss}$ is determined by $k_1$ and $k_2$, *i.e.*<br>$A_{ss}=\frac{k_1+k_3}{k_2+k_{11}}$ | schematic perturbation phase diagram<br>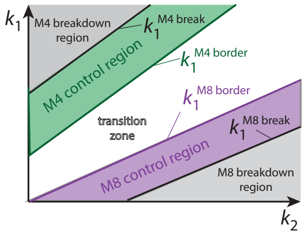<br><br><br>$k_1^{M4\,break}$ :<br>$j^{M4}_{comp}=0 \;\&\; j^{M8}_{comp}=k_{11}\cdot A^{M4}_{set}$<br>$\Rightarrow k_1^{M4\,break}=(k_2+k_{11})A^{M4}_{set}$<br><br>$k_1^{M4\,border}$ :<br>$j^{M4}_{comp}=k_3 \;\&\; j^{M8}_{comp}=k_{11}\cdot A^{M4}_{set}$<br>$\Rightarrow k_1^{M4\,border}=(k_2+k_{11})A^{M4}_{set}-k_3$<br><br>$k_1^{M8\,border}$ :<br>$j^{M4}_{comp}=k_3 \;\&\; j^{M8}_{comp}=k_{11}\cdot A^{M8}_{set}$<br>$\Rightarrow k_1^{M8\,border}=(k_2+k_{11})A^{M8}_{set}-k_3$<br><br>$k_1^{M8\,break}$ :<br>$j^{M4}_{comp}=k_3 \;\&\; j^{M8}_{comp}=0$<br>$\Rightarrow k_1^{M8\,break}=k_2A^{M8}_{set}-k_3$ |

**Table S17.** Isolated control of combined M1 and M5 controllers ($A_{set}^{M1} < A_{set}^{M5}$)

| combined controllers | rate equations and compensatory fluxes | borderlines in phase diagram |
|---|---|---|
| 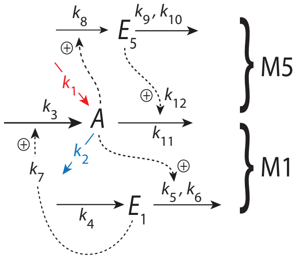 | Rate equations and set−points :<br>$\dot{A}=k_1-k_2A+\underbrace{k_3\left(\frac{E_1}{k_7+E_1}\right)}_{j_{comp}^{M1}}-\underbrace{\frac{k_{11}A\cdot E_5}{k_{12}+E_5}}_{j_{comp}^{M5}}$<br>$\dot{E_1}=k_4-k_5\underbrace{\left(\frac{E_1}{k_6+E_1}\right)}_{\approx 1}A \Rightarrow A_{set}^{M1}=\frac{k_4}{k_5}$<br>$\dot{E_5}=k_8A-k_9\underbrace{\left(\frac{E_5}{k_{10}+E_5}\right)}_{\approx 1} \Rightarrow A_{set}^{M5}=\frac{k_9}{k_8}$<br><br>Windup conditions and compensatory fluxes :<br>(i) M1 controls $A$ :<br>$A_{ss}=A_{set}^{M1} \Rightarrow \dot{E_5}<0$<br>$\Rightarrow j_{comp}^{M5} \to 0$<br>− − − − − − −<br>(ii) M5 controls $A$ :<br>$A_{ss}=A_{set}^{M5} \Rightarrow \dot{E_1}<0$<br>$\Rightarrow j_{comp}^{M1} \to 0$<br>− − − − − − −<br>(iii) $A_{ss}$ is in transition zone :<br>$A_{set}^{M1}<A_{ss}<A_{set}^{M5} \Rightarrow \dot{E_1}<0$ & $\dot{E_5}<0$<br>$\Rightarrow j_{comp}^{M1}=0,$<br>& $j_{comp}^{M5}=0.$<br>$A_{ss}$ is only determined by $k_1$ and $k_2$ :<br>$A_{ss}=\frac{k_1}{k_2}$ | schematic perturbation phase diagram<br>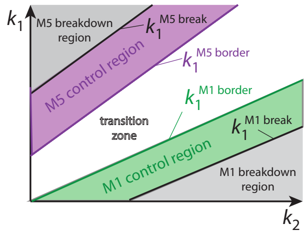<br>$k_1^{M1\,break}$ :<br>$j_{comp}^{M1}=j_{comp,\,max}^{M1}=k_3$ & $j_{comp}^{M5}=0$<br>$\Rightarrow k_1^{M1\,break}=k_2A_{set}^{M1}-k_3$<br><br>$k_1^{M1\,border}$ :<br>$j_{comp}^{M1}=0$ & $j_{comp}^{M5}=0$<br>$\Rightarrow k_1^{M1\,border}=k_2A_{set}^{M1}$<br><br>$k_1^{M5\,border}$ :<br>$j_{comp}^{M1}=0$ & $j_{comp}^{M5}=0$<br>$\Rightarrow k_1^{M5\,border}=k_2A_{set}^{M5}$<br><br>$k_1^{M5\,break}$ :<br>$j_{comp}^{M1}=0$ & $j_{comp}^{M5}=j_{comp\,max}^{M5}$<br>$=k_{11}A_{set}^{M5}$<br>$\Rightarrow k_1^{M5\,break}=(k_2+k_{11})A_{set}^{M5}$ |

**Table S18.** Isolated control of combined M1 and M6 controllers ($A^{M1}_{set}<A^{M6}_{set}$)

| combined controllers | rate equations and compensatory fluxes | borderlines in phase diagram |
|---|---|---|
| 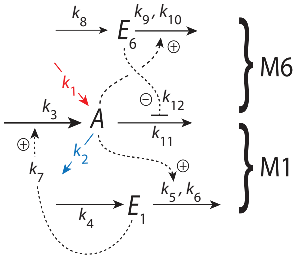<br> | Rate equations and set−points :<br>$\dot{A}=k_1-k_2A+\underbrace{k_3\left(\frac{E_1}{k_7+E_1}\right)}_{j^{M1}_{comp}}-\underbrace{\frac{k_{11}A\cdot k_{12}}{k_{12}+E_6}}_{j^{M6}_{comp}}$<br>$\dot{E}_1=k_4-k_5\underbrace{\left(\frac{E_1}{k_6+E_1}\right)}_{\approx 1}A \Rightarrow A^{M1}_{set}=\frac{k_4}{k_5}$<br>$\dot{E}_6=k_8-k_9A\underbrace{\left(\frac{E_6}{k_{10}+E_6}\right)}_{\approx 1} \Rightarrow A^{M6}_{set}=\frac{k_8}{k_9}$<br>Windup conditions and compensatory fluxes :<br>(i) M1 controls $A$ :<br>$A_{ss}=A^{M1}_{set} \Rightarrow \dot{E}_6>0$<br>$\Rightarrow j^{M6}_{comp} \to 0$<br>− − − − − − −<br>(ii) M6 controls $A$ :<br>$A_{ss}=A^{M6}_{set} \Rightarrow \dot{E}_1<0$<br>$\Rightarrow j^{M1}_{comp} \to 0$<br>− − − − − − −<br>(iii) $A_{ss}$ is in transition zone :<br>$A^{M1}_{set}<A_{ss}<A^{M6}_{set} \Rightarrow \dot{E}_1<0\ \&\ \dot{E}_6>0$<br>$\Rightarrow j^{M1}_{comp,max}=0,$<br>$\&\ j^{M6}_{comp,max}=0.$<br>$A_{ss}$ is determined by only $k_1$ and $k_2$ :<br>$A_{ss}=\frac{k_1}{k_2}$ | schematic perturbation phase diagram<br>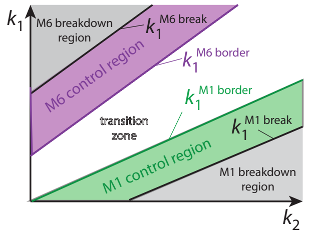<br><br>$k_1^{M1\,break}$ :<br>$j^{M1}_{comp}=j^{M1}_{comp\,max}=k_3\ \&\ j^{M6}_{comp}=0$<br>$\Rightarrow k_1^{M1\,break}=k_2A^{M1}_{set}-k_3$<br>$k_1^{M1\,border}$ :<br>$j^{M1}_{comp}=0\ \&\ j^{M6}_{comp}=0$<br>$\Rightarrow k_1^{M1\,border}=k_2A^{M1}_{set}$<br>$k_1^{M6\,border}$ :<br>$j^{M1}_{comp}=0\ \&\ j^{M6}_{comp}=0$<br>$\Rightarrow k_1^{M6\,border}=k_2A^{M6}_{set}$<br>$k_1^{M6\,break}$ :<br>$j^{M1}_{comp}=0\ \&\ j^{M6}_{comp}=j^{M6}_{comp\,max}$<br>$=k_{11}A^{M6}_{set}$<br>$\Rightarrow k_1^{M6\,break}=(k_2+k_{11})A^{M6}_{set}$ |

**Table S19.** Isolated control of combined M1 and M7 controllers ($A_{set}^{M1}<A_{set}^{M7}$)

| combined controllers | rate equations and compensatory fluxes | borderlines in phase diagram |
| --- | --- | --- |
| 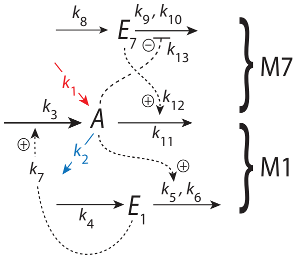<br> | Rate equations and set−points :<br>$\dot{A}=k_1-k_2A+\underbrace{k_3\left(\frac{E_1}{k_7+E_1}\right)}_{j_{comp}^{M1}}-\underbrace{\frac{k_{11}A\cdot E_7}{k_{12}+E_7}}_{j_{comp}^{M7}}$<br>$\dot{E}_1=k_4-k_5\underbrace{\left(\frac{E_1}{k_6+E_1}\right)}_{\approx 1}A\Rightarrow A_{set}^{M1}=\frac{k_4}{k_5}$<br>$\dot{E}_7=k_8-k_9\underbrace{\left(\frac{E_7}{k_{10}+E_7}\right)}_{\approx 1}\left(\frac{k_{13}}{k_{13}+A}\right)$<br>$\Rightarrow A_{set}^{M7}=\frac{k_9k_{13}-k_8k_{13}}{k_8}$<br>Windup conditions and compensatory fluxes :<br>(i) M1 controls $A$ :<br>$A_{ss}=A_{set}^{M1}\Rightarrow\dot{E}_7<0$<br>$\Rightarrow j_{comp}^{M7}\to 0$<br>− − − − − − −<br>(ii) M7 controls $A$ :<br>$A_{ss}=A_{set}^{M7}\Rightarrow\dot{E}_1<0$<br>$\Rightarrow j_{comp}^{M1}\to 0$<br>− − − − − − −<br>(iii) $A_{ss}$ is in transition zone :<br>$A_{set}^{M1}<A_{ss}<A_{set}^{M7}\Rightarrow\dot{E}_1<0\ \&\ \dot{E}_7<0$<br>$\Rightarrow j_{comp}^{M1}=0,$<br>$\&\ j_{comp}^{M7}=0.$<br>$A_{ss}$ is determined by only $k_1$ and $k_2$ :<br>$A_{ss}=\frac{k_1}{k_2}$ | schematic perturbation phase diagram<br>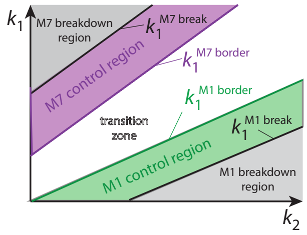<br><br>$k_1^{M1\,break}$ :<br>$j_{comp}^{M1}=j_{comp\,max}^{M1}=k_3\ \&\ j_{comp}^{M7}=0$<br>$\Rightarrow k_1^{M1\,break}=k_2A_{set}^{M1}-k_3$<br>$k_1^{M1\,border}$ :<br>$j_{comp}^{M1}=0\ \&\ j_{comp}^{M7}=0$<br>$\Rightarrow k_1^{M1\,border}=k_2A_{set}^{M1}$<br>$k_1^{M7\,border}$ :<br>$j_{comp}^{M1}=0\ \&\ j_{comp}^{M7}=0$<br>$\Rightarrow k_1^{M7\,border}=k_2A_{set}^{M7}$<br>$k_1^{M7\,break}$ :<br>$j_{comp}^{M1}=0\ \&\ j_{comp}^{M7}=j_{comp\,max}^{M7}$<br>$=k_{11}A_{set}^{M7}$<br>$\Rightarrow k_1^{M7\,break}=(k_2+k_{11})A_{set}^{M7}$ |

**Table S20.** Isolated control of combined M1 and M8 controllers ($A_{set}^{M1}<A_{set}^{M8}$)

| combined controllers | rate equations and compensatory fluxes | borderlines in phase diagram |
|---|---|---|
| 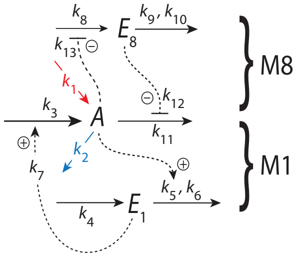 | Rate equations and set−points :<br>$\dot{A}=k_1-k_2A+\underbrace{k_3\left(\frac{E_1}{k_7+E_1}\right)}_{j_{comp}^{M1}}-\underbrace{\frac{k_{11}A\cdot k_{12}}{k_{12}+E_8}}_{j_{comp}^{M8}}$<br>$\dot{E}_1=k_4-k_5\underbrace{\left(\frac{E_1}{k_6+E_1}\right)}_{\approx 1}A \Rightarrow A_{set}^{M1}=\frac{k_4}{k_5}$<br>$\dot{E}_8=k_8\left(\frac{k_{13}}{k_{13}+A}\right)-k_9\underbrace{\left(\frac{E_8}{k_{10}+E_8}\right)}_{\approx 1}$<br>$\Rightarrow A_{set}^{M8}=\frac{k_8k_{13}-k_9k_{13}}{k_9}$<br>Windup conditions and compensatory fluxes :<br>(i) M1 controls $A$ :<br>$A_{ss}=A_{set}^{M1} \Rightarrow \dot{E}_8>0$<br>$\Rightarrow j_{comp}^{M8}\to 0$<br>– – – – – – –<br>(ii) M8 controls $A$ :<br>$A_{ss}=A_{set}^{M8}\Rightarrow \dot{E}_1<0$<br>$\Rightarrow j_{comp}^{M1}\to 0$<br>– – – – – – –<br>(iii) $A_{ss}$ is in transition zone :<br>$A_{set}^{M1}<A_{ss}<A_{set}^{M8}\Rightarrow \dot{E}_1<0\ \&\ \dot{E}_8>0$<br>$\Rightarrow j_{comp}^{M1}=0,$<br>$\&\ j_{comp}^{M8}=0.$<br>$A_{ss}$ is determined by only $k_1$ and $k_2$ :<br>$A_{ss}=\frac{k_1}{k_2}$ | schematic perturbation phase diagram<br>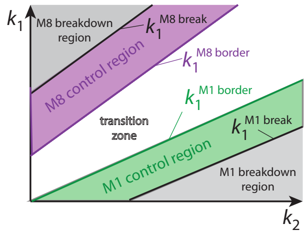<br>$k_1^{M1\,break}$ :<br>$j_{comp}^{M1}=j_{comp\,max}^{M1}=k_3\ \&\ j_{comp}^{M8}=0$<br>$\Rightarrow k_1^{M1\,break}=k_2A_{set}^{M1}-k_3$<br>$k_1^{M1\,border}$ :<br>$j_{comp}^{M1}=0\ \&\ j_{comp}^{M8}=0$<br>$\Rightarrow k_1^{M1\,border}=k_2A_{set}^{M1}$<br>$k_1^{M8\,border}$ :<br>$j_{comp}^{M1}=0\ \&\ j_{comp}^{M8}=0$<br>$\Rightarrow k_1^{M8\,border}=k_2A_{set}^{M8}$<br>$k_1^{M8\,break}$ :<br>$j_{comp}^{M1}=0\ \&\ j_{comp}^{M8}=j_{comp\,max}^{M8}$<br>$=k_{11}A_{set}^{M8}$<br>$\Rightarrow k_1^{M8\,break}=(k_2+k_{11})A_{set}^{M8}$ |

**Table S21.** Isolated control of combined M2 and M5 controllers ($A_{set}^{M2}<A_{set}^{M5}$)

| combined controllers | rate equations and compensatory fluxes | borderlines in phase diagram |
|---|---|---|
| 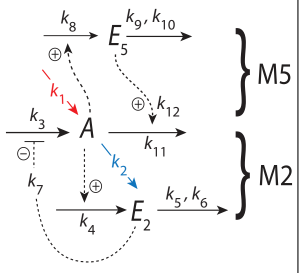<br> | Rate equations and set−points :<br>$\dot{A}=k_1-k_2A+k_3\underbrace{\left(\frac{k_7}{k_7+E_2}\right)}_{j_{comp}^{M2}}-\underbrace{\frac{k_{11}A\cdot E_5}{k_{12}+E_5}}_{j_{comp}^{M5}}$<br>$\dot{E_2}=k_4A-k_5\underbrace{\left(\frac{E_2}{k_6+E_2}\right)}_{\approx1}\Rightarrow A_{set}^{M2}=\frac{k_5}{k_4}$<br>$\dot{E_5}=k_8A-k_9\underbrace{\left(\frac{E_5}{k_{10}+E_5}\right)}_{\approx1}\Rightarrow A_{set}^{M5}=\frac{k_9}{k_8}$<br><br>Windup conditions and<br>compensatory fluxes :<br>(i) M2 controls $A$ :<br>$A_{ss}=A_{set}^{M2}\Rightarrow \dot{E_5}<0$<br>$\Rightarrow j_{comp}^{M5}\rightarrow 0$<br>− − − − − − −<br>(ii) M5 controls $A$ :<br>$A_{ss}=A_{set}^{M5}\Rightarrow \dot{E_2}>0$<br>$\Rightarrow j_{comp}^{M2}\rightarrow 0$<br>− − − − − − −<br>(iii) $A_{ss}$ is in transition zone :<br>$A_{set}^{M2}<A_{ss}<A_{set}^{M5}\Rightarrow \dot{E_2}>0\ \&\ \dot{E_5}<0$<br>$\Rightarrow j_{comp}^{M2}=0,$<br>$\&\ j_{comp}^{M5}=0.$<br>$A_{ss}$ is determined by only $k_1$ and $k_2$ :<br>$A_{ss}=\frac{k_1}{k_2}$ | schematic perturbation phase diagram<br>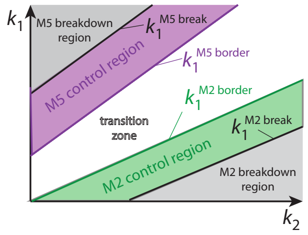<br><br>$k_1^{M2\,break}$ :<br>$j_{comp}^{M2}=j_{comp\,max}^{M2}=k_3\ \&\ j_{comp}^{M5}=0$<br>$\Rightarrow k_1^{M2\,break}=k_2A_{set}^{M2}-k_3$<br><br>$k_1^{M2\,border}$ :<br>$j_{comp}^{M2}=0\ \&\ j_{comp}^{M5}=0$<br>$\Rightarrow k_1^{M2\,border}=k_2A_{set}^{M2}$<br><br>$k_1^{M5\,border}$ :<br>$j_{comp}^{M2}=0\ \&\ j_{comp}^{M5}=0$<br>$\Rightarrow k_1^{M5\,border}=k_2A_{set}^{M5}$<br><br>$k_1^{M5\,break}$ :<br>$j_{comp}^{M2}=0\ \&\ j_{comp}^{M5}=j_{comp\,max}^{M5}$<br>$=k_{11}A_{set}^{M5}$<br>$\Rightarrow k_1^{M5\,break}=(k_2+k_{11})A_{set}^{M5}$ |

**Table S22.** Isolated control of combined M2 and M6 controllers ($A^{M2}_{set}<A^{M6}_{set}$)

| combined controllers | rate equations and compensatory fluxes | borderlines in phase diagram |
|---|---|---|
| 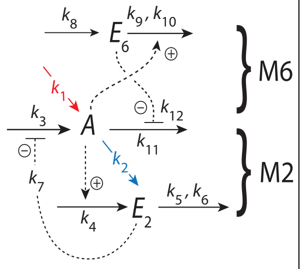<br> | Rate equations and set−points :<br>$\dot{A}=k_1-k_2A+k_3\underbrace{\left(\frac{k_7}{k_7+E_2}\right)}_{j^{M2}_{comp}}-\underbrace{\frac{k_{11}A\cdot k_{12}}{k_{12}+E_6}}_{j^{M6}_{comp}}$<br>$\dot{E}_2=k_4A-k_5\underbrace{\left(\frac{E_2}{k_6+E_2}\right)}_{\approx 1}\Rightarrow A^{M2}_{set}=\frac{k_5}{k_4}$<br>$\dot{E}_6=k_8-k_9A\underbrace{\left(\frac{E_6}{k_{10}+E_6}\right)}_{\approx 1}\Rightarrow A^{M6}_{set}=\frac{k_8}{k_9}$<br><br>Windup conditions and compensatory fluxes :<br>(i) M2 controls $A$ :<br>$A_{ss}=A^{M2}_{set}\Rightarrow\dot{E}_6>0$<br>$\Rightarrow j^{M6}_{comp}\rightarrow 0$<br>− − − − − − −<br>(ii) M6 controls $A$ :<br>$A_{ss}=A^{M6}_{set}\Rightarrow\dot{E}_2>0$<br>$\Rightarrow j^{M2}_{comp}\rightarrow 0$<br>− − − − − − −<br>(iii) $A_{ss}$ is in transition zone :<br>$A^{M2}_{set}<A_{ss}<A^{M6}_{set}\Rightarrow\dot{E}_2>0\ \&\ \dot{E}_6>0$<br>$\Rightarrow j^{M2}_{comp}=0,$<br>$\&\ j^{M6}_{comp}=0.$<br>$A_{ss}$ is determined by only $k_1$ and $k_2$ :<br>$A_{ss}=\frac{k_1}{k_2}$ | schematic perturbation phase diagram<br>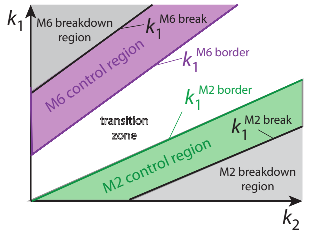<br><br>$k_1^{M2\,break}$ :<br>$j^{M2}_{comp}=j^{M2}_{comp\,max}=k_3\ \&\ j^{M6}_{comp}=0$<br>$\Rightarrow k_1^{M2\,break}=k_2A^{M2}_{set}-k_3$<br><br>$k_1^{M2\,border}$ :<br>$j^{M2}_{comp}=0\ \&\ j^{M6}_{comp}=0$<br>$\Rightarrow k_1^{M2\,border}=k_2A^{M2}_{set}$<br><br>$k_1^{M6\,border}$ :<br>$j^{M2}_{comp}=0\ \&\ j^{M6}_{comp}=0$<br>$\Rightarrow k_1^{M6\,border}=k_2A^{M6}_{set}$<br><br>$k_1^{M6\,break}$ :<br>$j^{M2}_{comp}=0\ \&\ j^{M6}_{comp}=j^{M6}_{comp\,max}$<br>$=k_{11}A^{M6}_{set}$<br>$\Rightarrow k_1^{M6\,break}=(k_2+k_{11})A^{M6}_{set}$ |

**Table S23.** Isolated control of combined M2 and M7 controllers ($A^{M2}_{set}<A^{M7}_{set}$)

| combined controllers | rate equations and compensatory fluxes | borderlines in phase diagram |
|---|---|---|
| 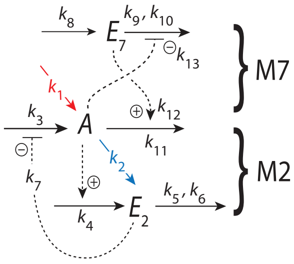<br> | Rate equations and set−points :<br>$\dot{A}=k_1-k_2A+\underbrace{k_3\left(\frac{k_7}{k_7+E_2}\right)}_{j^{M2}_{comp}}-\underbrace{\frac{k_{11}A\cdot E_7}{k_{12}+E_7}}_{j^{M7}_{comp}}$<br>$\dot{E}_2=k_4A-k_5\underbrace{\left(\frac{E_2}{k_6+E_2}\right)}_{\approx 1}\Rightarrow A^{M2}_{set}=\frac{k_5}{k_4}$<br>$\dot{E}_7=k_8-k_9\underbrace{\left(\frac{E_7}{k_{10}+E_7}\right)}_{\approx 1}\left(\frac{k_{13}}{k_{13}+A}\right)$<br>$\Rightarrow A^{M7}_{set}=\frac{k_9k_{13}-k_8k_{13}}{k_8}$<br><br>Windup conditions and<br>compensatory fluxes :<br>(i) M2 controls $A$ :<br>$A_{ss}=A^{M2}_{set}\Rightarrow\dot{E}_7<0$<br>$\Rightarrow j^{M7}_{comp,max}\to 0$<br>− − − − − − −<br>(ii) M7 controls $A$ :<br>$A_{ss}=A^{M7}_{set}\Rightarrow\dot{E}_2>0$<br>$\Rightarrow j^{M2}_{comp,max}\to 0$<br>− − − − − − −<br>(iii) $A_{ss}$ is in transition zone :<br>$A^{M2}_{set}<A_{ss}<A^{M7}_{set}\Rightarrow\dot{E}_2>0\,\&\,\dot{E}_7<0$<br>$\Rightarrow j^{M2}_{comp}=0,$<br>$\&\ j^{M7}_{comp}=0.$<br>$A_{ss}$ is only determined by $k_1$ and $k_2$ :<br>$A_{ss}=\frac{k_1}{k_2}$ | schematic perturbation phase diagram<br>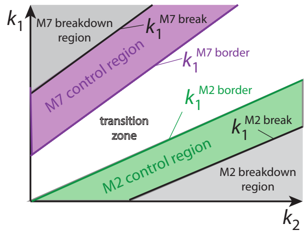<br><br>$k_1^{M2\,break}$ :<br>$j^{M2}_{comp}=j^{M2}_{comp\;max}=k_3\,\&\,j^{M7}_{comp}=0$<br>$\Rightarrow k_1^{M2\,break}=k_2A^{M2}_{set}-k_3$<br><br>$k_1^{M2\,border}$ :<br>$j^{M2}_{comp}=0\,\&\,j^{M7}_{comp}=0$<br>$\Rightarrow k_1^{M2\,border}=k_2A^{M2}_{set}$<br><br>$k_1^{M7\,border}$ :<br>$j^{M2}_{comp}=0\,\&\,j^{M7}_{comp}=0$<br>$\Rightarrow k_1^{M7\,border}=k_2A^{M7}_{set}$<br><br>$k_1^{M7\,break}$ :<br>$j^{M2}_{comp}=0\,\&\,j^{M7}_{comp}=j^{M7}_{comp\,max}$<br>$=k_{11}A^{M7}_{set}$<br>$\Rightarrow k_1^{M7\,break}=(k_2+k_{11})A^{M7}_{set}$ |

### Table S24. Isolated control of combined M2 and M8 controllers ($A^{M2}_{set}<A^{M8}_{set}$)

| combined controllers | rate equations and compensatory fluxes | borderlines in phase diagram |
|---|---|---|
| 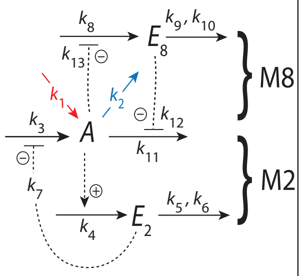  | Rate equations and set−points :<br>$\dot{A}=k_1-k_2A+\underbrace{k_3\left(\frac{k_7}{k_7+E_2}\right)}_{j^{M2}_{comp}}-\underbrace{\frac{k_{11}A\cdot k_{12}}{k_{12}+E_8}}_{j^{M8}_{comp}}$<br>$\dot{E}_2=k_4A-k_5\underbrace{\left(\frac{E_2}{k_6+E_2}\right)}_{\approx 1}\Rightarrow A^{M2}_{set}=\frac{k_5}{k_4}$<br>$\dot{E}_8=k_8\left(\frac{k_{13}}{k_{13}+A}\right)-k_9\underbrace{\left(\frac{E_8}{k_{10}+E_8}\right)}_{\approx 1}$<br>$\Rightarrow A^{M8}_{set}=\frac{k_8k_{13}-k_9k_{13}}{k_9}$<br>Windup conditions and compensatory fluxes :<br>(i) M2 controls $A$ :<br>$A_{ss}=A^{M2}_{set}\Rightarrow \dot{E}_8>0$<br>$\Rightarrow j^{M8}_{comp}\to 0$<br>− − − − − − −<br>(ii) M8 controls $A$ :<br>$A_{ss}=A^{M8}_{set}\Rightarrow \dot{E}_2>0$<br>$\Rightarrow j^{M2}_{comp}\to 0$<br>− − − − − − −<br>(iii) $A_{ss}$ is in transition zone :<br>$A^{M2}_{set}<A_{ss}<A^{M8}_{set}\Rightarrow \dot{E}_2>0\ \&\ \dot{E}_8>0$<br>$\Rightarrow j^{M2}_{comp,max}=0,$<br>$\&\ j^{M8}_{comp,max}=0.$<br>$A_{ss}$ is only determined by $k_1$ and $k_2$ :<br>$A_{ss}=\frac{k_1}{k_2}$ | schematic perturbation phase diagram<br>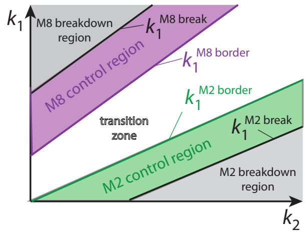 <br>$k_1^{M2\,break}$ :<br>$j^{M2}_{comp}=j^{M2}_{comp\ \max}=k_3\ \&\ j^{M8}_{comp}=0$<br>$\Rightarrow k_1^{M2\,break}=k_2A^{M2}_{set}-k_3$<br>$k_1^{M2\,border}$ :<br>$j^{M2}_{comp}=0\ \&\ j^{M8}_{comp}=0$<br>$\Rightarrow k_1^{M2\,border}=k_2A^{M2}_{set}$<br>$k_1^{M8\,border}$ :<br>$j^{M2}_{comp}=0\ \&\ j^{M8}_{comp}=0$<br>$\Rightarrow k_1^{M8\,border}=k_2A^{M8}_{set}$<br>$k_1^{M8\,break}$ :<br>$j^{M2}_{comp}=0\ \&\ j^{M8}_{comp}=j^{M8}_{comp\ max}$<br>$=k_{11}A^{M8}_{set}$<br>$\Rightarrow k_1^{M8\,break}=(k_2+k_{11})A^{M8}_{set}$ |

**Table S25.** Isolated control of combined M3 and M5 controllers ($A^{M3}_{set}<A^{M5}_{set}$)

| combined controllers | rate equations and compensatory fluxes | borderlines in phase diagram |
|---|---|---|
| 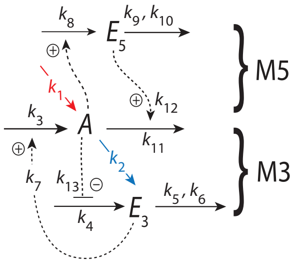 | Rate equations and set−points :<br>$\dot{A}=k_1-k_2A+\underbrace{k_3\left(\frac{E_3}{k_7+E_3}\right)}_{j^{M3}_{comp}}-\underbrace{\frac{k_{11}A\cdot E_5}{k_{12}+E_5}}_{j^{M5}_{comp}}$<br>$\dot{E}_3=k_4\left(\frac{k_{13}}{k_{13}+A}\right)-k_5\underbrace{\left(\frac{E_3}{k_6+E_3}\right)}_{\approx 1}$<br>$\Rightarrow A^{M3}_{set}=\frac{k_4k_{13}-k_5k_{13}}{k_5}$<br>$\dot{E}_5=k_8A-k_9\underbrace{\left(\frac{E_5}{k_{10}+E_5}\right)}_{\approx 1}\Rightarrow A^{M5}_{set}=\frac{k_9}{k_8}$<br><br>Windup conditions and<br>compensatory fluxes :<br>(i) M3 controls $A$ :<br>$A_{ss}=A^{M3}_{set}\Rightarrow \dot{E}_5<0$<br>$\Rightarrow j^{M5}_{comp}\rightarrow 0$<br>− − − − − − −<br>(ii) M5 controls $A$ :<br>$A_{ss}=A^{M5}_{set}\Rightarrow \dot{E}_3<0$<br>$\Rightarrow j^{M3}_{comp}\rightarrow 0$<br>− − − − − − −<br>(iii) $A_{ss}$ is in transition zone :<br>$A^{M3}_{set}<A_{ss}<A^{M5}_{set}\Rightarrow \dot{E}_3<0\ \&\ \dot{E}_5<0$<br>$\Rightarrow j^{M3}_{comp}=0,$<br>$\&\ j^{M5}_{comp}=0.$<br>$A_{ss}$ is only determined by $k_1$ and $k_2$ :<br>$A_{ss}=\frac{k_1}{k_2}$ | schematic perturbation phase diagram<br>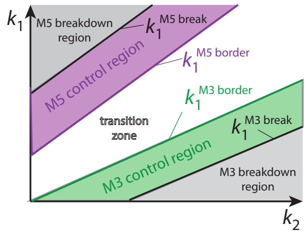<br>$k_1^{M3\,break}$ :<br>$j^{M3}_{comp}=j^{M3}_{comp}=k_3\ \&\ j^{M5}_{comp}=0$<br>$\Rightarrow k_1^{M3\,break}=k_2A^{M3}_{set}-k_3$<br><br>$k_1^{M3\,border}$ :<br>$j^{M3}_{comp}=0\ \&\ j^{M5}_{comp}=0$<br>$\Rightarrow k_1^{M3\,border}=k_2A^{M3}_{set}$<br><br>$k_1^{M5\,border}$ :<br>$j^{M3}_{comp}=0\ \&\ j^{M5}_{comp}=0$<br>$\Rightarrow k_1^{M5\,border}=k_2A^{M5}_{set}$<br><br>$k_1^{M5\,break}$ :<br>$j^{M3}_{comp}=0\ \&\ j^{M5}_{comp}=j^{M5}_{comp\ max}$<br>$=k_{11}A^{M5}_{set}$<br>$\Rightarrow k_1^{M5\,break}=(k_2+k_{11})A^{M5}_{set}$ |

**Table S26.** Isolated control of combined M3 and M6 controllers ($A^{M3}_{set}<A^{M6}_{set}$)

<table>
<tr><th>combined controllers</th><th>rate equations and compensatory fluxes</th><th>borderlines in phase diagram</th></tr>
<tr>
<td>
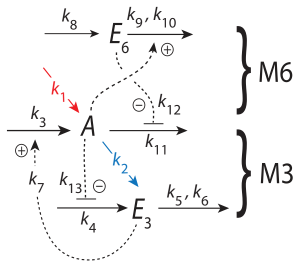

</td>
<td>
Rate equations and set−points :<br>
$\dot{A}=k_1-k_2A+\underbrace{k_3\left(\frac{E_3}{k_7+E_3}\right)}_{j^{M3}_{comp}}-\underbrace{\frac{k_{11}A\cdot k_{12}}{k_{12}+E_6}}_{j^{M6}_{comp}}$<br>
$\dot{E}_3=k_4\left(\frac{k_{13}}{k_{13}+A}\right)-k_5\underbrace{\left(\frac{E_3}{k_6+E_3}\right)}_{\approx 1}$<br>
$\Rightarrow A^{M3}_{set}=\frac{k_4k_{13}-k_5k_{13}}{k_5}$<br>
$\dot{E}_6=k_8-k_9A\underbrace{\left(\frac{E_6}{k_{10}+E_6}\right)}_{\approx 1}\Rightarrow A^{M6}_{set}=\frac{k_8}{k_9}$<br>
<br>
Windup conditions and<br>
compensatory fluxes :<br>
(i) M3 controls $A$ :<br>
$A_{ss}=A^{M3}_{set}\Rightarrow \dot{E}_6>0$<br>
$\Rightarrow j^{M6}_{comp}\to 0$<br>
– – – – – – –<br>
(ii) M6 controls $A$ :<br>
$A_{ss}=A^{M6}_{set}\Rightarrow \dot{E}_3<0$<br>
$\Rightarrow j^{M3}_{comp}\to 0$<br>
– – – – – – –<br>
(iii) $A_{ss}$ is in transition zone :<br>
$A^{M3}_{set}<A_{ss}<A^{M6}_{set}\Rightarrow \dot{E}_3<0\ \&\ \dot{E}_6>0$<br>
$\Rightarrow j^{M3}_{comp}=0,$<br>
$\&\ j^{M6}_{comp0}=0.$<br>
$A_{ss}$ is only determined by $k_1$ and $k_2$ :<br>
$A_{ss}=\frac{k_1}{k_2}$
</td>
<td>
schematic perturbation phase diagram<br>
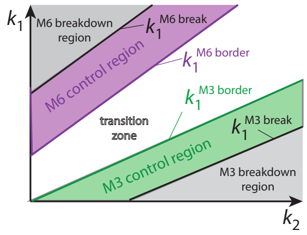

<br>
$k_1^{M3\,break}$ :<br>
$j^{M3}_{comp}=j^{M3}_{comp\,max}=k_3\ \&\ j^{M6}_{comp}=0$<br>
$\Rightarrow k_1^{M3\,break}=k_2A^{M3}_{set}-k_3$<br>
<br>
$k_1^{M3\,border}$ :<br>
$j^{M3}_{comp}=0\ \&\ j^{M6}_{comp}=0$<br>
$\Rightarrow k_1^{M3\,border}=k_2A^{M3}_{set}$<br>
<br>
$k_1^{M6\,border}$ :<br>
$j^{M3}_{comp}=0\ \&\ j^{M6}_{comp}=0$<br>
$\Rightarrow k_1^{M6\,border}=k_2A^{M6}_{set}$<br>
<br>
$k_1^{M6\,break}$ :<br>
$j^{M3}_{comp}=0\ \&\ j^{M6}_{comp}=j^{M6}_{comp\,max}$<br>
$=k_{11}A^{M6}_{set}$<br>
$\Rightarrow k_1^{M6\,break}=(k_2+k_{11})A^{M6}_{set}$
</td>
</tr>
</table>

**Table S27.** Isolated control of combined M3 and M7 controllers ($A_{set}^{M3}<A_{set}^{M7}$)

| combined controllers | rate equations and compensatory fluxes | borderlines in phase diagram |
|---|---|---|
| 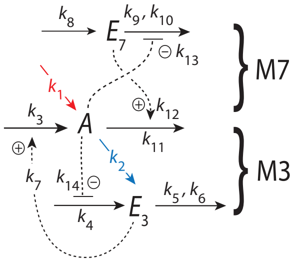<br> | Rate equations and set−points :<br>$\dot{A}=k_1-k_2A+\underbrace{k_3\left(\frac{E_3}{k_7+E_3}\right)}_{j_{comp}^{M3}}-\underbrace{\frac{k_{11}A\cdot E_7}{k_{12}+E_7}}_{j_{comp}^{M7}}$<br>$\dot{E}_3=k_4\left(\frac{k_{13}}{k_{13}+A}\right)-k_5\underbrace{\left(\frac{E_3}{k_6+E_3}\right)}_{\approx 1}$<br>$\Rightarrow A_{set}^{M3}=\frac{k_4k_{14}-k_5k_{14}}{k_5}$<br>$\dot{E}_7=k_8-k_9\left(\frac{E_7}{k_{10}+E_7}\right)\underbrace{\left(\frac{k_{13}}{k_{13}+A}\right)}_{\approx 1}$<br>$\Rightarrow A_{set}^{M7}=\frac{k_9k_{13}-k_8k_{13}}{k_8}$<br>Windup conditions and compensatory fluxes :<br>(i) M3 controls $A$ :<br>$A_{ss}=A_{set}^{M3}\Rightarrow \dot{E}_7<0$<br>$\Rightarrow j_{comp}^{M7}\to 0$<br>– – – – – – –<br>(ii) M7 controls $A$ :<br>$A_{ss}=A_{set}^{M7}\Rightarrow \dot{E}_3<0$<br>$\Rightarrow j_{comp}^{M3}\to 0$<br>– – – – – – –<br>(iii) $A_{ss}$ is in transition zone :<br>$A_{set}^{M3}<A_{ss}<A_{set}^{M7}\Rightarrow \dot{E}_3<0\ \&\ \dot{E}_7<0$<br>$\Rightarrow j_{comp,max}^{M3}=0,$<br>$\&\ j_{comp,max}^{M7}=0.$<br>$A_{ss}$ is only determined by $k_1$ and $k_2$ :<br>$A_{ss}=\frac{k_1}{k_2}$ | schematic perturbation phase diagram<br>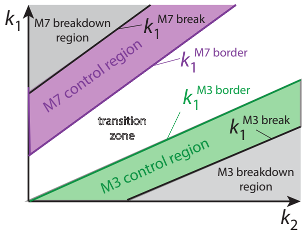<br><br>$k_1^{M3\,break}$ :<br>$j_{comp}^{M3}=j_{comp\,max}^{M3}=k_3\ \&\ j_{comp}^{M7}=0$<br>$\Rightarrow k_1^{M3\,break}=k_2A_{set}^{M3}-k_3$<br>$k_1^{M3\,border}$ :<br>$j_{comp}^{M3}=0\ \&\ j_{comp}^{M7}=0$<br>$\Rightarrow k_1^{M3\,border}=k_2A_{set}^{M3}$<br>$k_1^{M7\,border}$ :<br>$j_{comp}^{M3}=0\ \&\ j_{comp}^{M7}=0$<br>$\Rightarrow k_1^{M7\,border}=k_2A_{set}^{M7}$<br>$k_1^{M7\,break}$ :<br>$j_{comp}^{M3}=0\ \&\ j_{comp}^{M7}=j_{comp}^{M7}$<br>$=k_{11}A_{set}^{M7}$<br>$\Rightarrow k_1^{M7\,break}=(k_2+k_{11})A_{set}^{M7}$ |

**Table S28.** Isolated control of combined M3 and M8 controllers ($A_{set}^{M3}<A_{set}^{M8}$)

<table>
<tr><th>combined controllers</th><th>rate equations and compensatory fluxes</th><th>borderlines in phase diagram</th></tr>
<tr>
<td>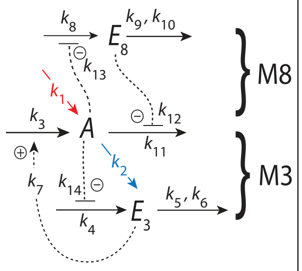
</td>
<td>Rate equations and set−points :<br>
$\dot{A}=k_1-k_2A+\underbrace{k_3\left(\frac{E_3}{k_7+E_3}\right)}_{j_{comp}^{M3}}-\underbrace{\frac{k_{11}A\cdot k_{12}}{k_{12}+E_8}}_{j_{comp}^{M8}}$<br>
$\dot{E}_3=k_4\left(\frac{k_{13}}{k_{13}+A}\right)-k_5\underbrace{\left(\frac{E_3}{k_6+E_3}\right)}_{\approx 1}$<br>
$\Rightarrow A_{set}^{M3}=\frac{k_4k_{14}-k_5k_{14}}{k_5}$<br>
$\dot{E}_8=k_8\left(\frac{k_{13}}{k_{13}+A}\right)-k_9\underbrace{\left(\frac{E_8}{k_{10}+E_8}\right)}_{\approx 1}$<br>
$\Rightarrow A_{set}^{M8}=\frac{k_8k_{13}-k_9k_{13}}{k_9}$<br>
Windup conditions and compensatory fluxes :<br>
(i) M3 controls $A$ :<br>
$A_{ss}=A_{set}^{M3}\Rightarrow \dot{E}_8>0$<br>
$\Rightarrow j_{comp}^{M8}\to 0$<br>
− − − − − − −<br>
(ii) M8 controls $A$ :<br>
$A_{ss}=A_{set}^{M8}\Rightarrow \dot{E}_3<0$<br>
$\Rightarrow j_{comp}^{M3}\to 0$<br>
− − − − − − −<br>
(iii) $A_{ss}$ is in transition zone :<br>
$A_{set}^{M3}<A_{ss}<A_{set}^{M8}\Rightarrow \dot{E}_3<0\ \&\ \dot{E}_8>0$<br>
$\Rightarrow j_{comp}^{M3}=0$,<br>
& $j_{comp}^{M8}=0$.<br>
$A_{ss}$ is only determined by $k_1$ and $k_2$ :<br>
$A_{ss}=\frac{k_1}{k_2}$</td>
<td>schematic perturbation phase diagram<br>
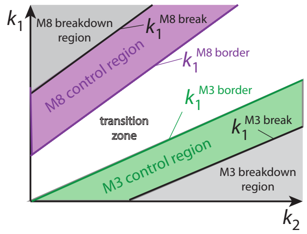

$k_1^{M3\,break}$ :<br>
$j_{comp}^{M3}=j_{comp\,max}^{M3}=k_3\ \&\ j_{comp}^{M8}=0$<br>
$\Rightarrow k_1^{M3\,break}=k_2A_{set}^{M3}-k_3$<br>
$k_1^{M3\,border}$ :<br>
$j_{comp}^{M3}=0\ \&\ j_{comp}^{M8}=0$<br>
$\Rightarrow k_1^{M3\,border}=k_2A_{set}^{M3}$<br>
$k_1^{M8\,border}$ :<br>
$j_{comp}^{M3}=0\ \&\ j_{comp}^{M8}=0$<br>
$\Rightarrow k_1^{M8\,border}=k_2A_{set}^{M8}$<br>
$k_1^{M8\,break}$ :<br>
$j_{comp}^{M3}=0\ \&\ j_{comp}^{M8}=j_{comp\,max}^{M8}$<br>
$=k_{11}A_{set}^{M8}$<br>
$\Rightarrow k_1^{M8\,break}=(k_2+k_{11})A_{set}^{M8}$</td>
</tr>
</table>

**Table S29.** Isolated control of combined M4 and M5 controllers ($A_{set}^{M4}<A_{set}^{M5}$)

| combined controllers | rate equations and compensatory fluxes | borderlines in phase diagram |
|---|---|---|
| 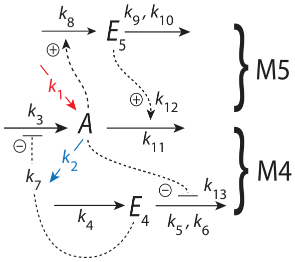<br> | Rate equations and set−points :<br>$\dot{A}=k_1-k_2A+\underbrace{k_3\left(\frac{k_7}{k_7+E_4}\right)}_{j_{comp}^{M4}}-\underbrace{\frac{k_{11}A\cdot E_5}{k_{12}+E_5}}_{j_{comp}^{M5}}$<br>$\dot{E_4}=k_4-\underbrace{k_5\left(\frac{E_4}{k_6+E_4}\right)}_{\approx 1}\cdot\left(\frac{k_{13}}{k_{13}+A}\right)$<br>$\Rightarrow A_{set}^{M4}=\frac{k_5k_{13}-k_4k_{13}}{k_4}$<br>$\dot{E_5}=k_8A-\underbrace{k_9\left(\frac{E_5}{k_{10}+E_5}\right)}_{\approx 1}\Rightarrow A_{set}^{M5}=\frac{k_9}{k_8}$<br><br>Windup conditions and compensatory fluxes :<br>(i) M4 controls $A$ :<br>$A_{ss}=A_{set}^{M4}\Rightarrow \dot{E_5}<0$<br>$\Rightarrow j_{comp}^{M5}\to 0$<br>− − − − − − −<br>(ii) M5 controls $A$ :<br>$A_{ss}=A_{set}^{M5}\Rightarrow \dot{E_4}>0$<br>$\Rightarrow j_{comp}^{M4}\to 0$<br>− − − − − − −<br>(iii) $A_{ss}$ is in transition zone :<br>$A_{set}^{M4}<A_{ss}<A_{set}^{M5}\Rightarrow \dot{E_4}>0\ \&\ \dot{E_5}<0$<br>$\Rightarrow j_{comp}^{M4}=0,$<br>$\&\ j_{comp}^{M5}=0.$<br>$A_{ss}$ is only determined by $k_1$ and $k_2$ :<br>$A_{ss}=\frac{k_1}{k_2}$ | schematic perturbation phase diagram<br>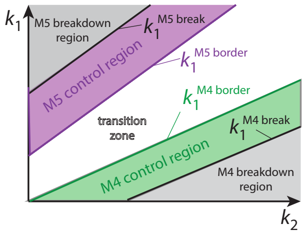<br><br><br>$k_1^{M4\,break}$ :<br>$j_{comp}^{M4}=j_{comp\ max}^{M4}=k_3\ \&\ j_{comp}^{M5}=0$<br>$\Rightarrow k_1^{M4\,break}=k_2A_{set}^{M4}-k_3$<br><br>$k_1^{M4\,border}$ :<br>$j_{comp}^{M4}=0\ \&\ j_{comp}^{M5}=0$<br>$\Rightarrow k_1^{M4\,border}=k_2A_{set}^{M4}$<br><br>$k_1^{M5\,border}$ :<br>$j_{comp}^{M4}=0\ \&\ j_{comp}^{M5}=0$<br>$\Rightarrow k_1^{M5\,border}=k_2A_{set}^{M5}$<br><br>$k_1^{M5\,break}$ :<br>$j_{comp}^{M4}=0\ \&\ j_{comp}^{M5}=j_{comp\ max}^{M5}$<br>$=k_{11}A_{set}^{M5}$<br>$\Rightarrow k_1^{M5\,break}=(k_2+k_{11})A_{set}^{M5}$ |

**Table S30.** Isolated control of combined M4 and M6 controllers ($A^{M4}_{set} < A^{M6}_{set}$)

<table>
<tr><th>combined controllers</th><th>rate equations and compensatory fluxes</th><th>borderlines in phase diagram</th></tr>
<tr>
<td>
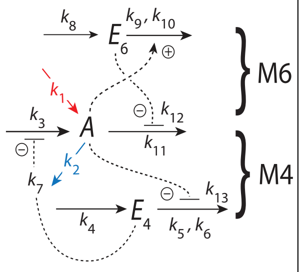

</td>
<td>
Rate equations and set−points :<br>
$\dot{A} = k_1 - k_2 A + \underbrace{k_3\left(\frac{k_7}{k_7+E_4}\right)}_{j^{M4}_{comp}} - \underbrace{\frac{k_{11} A \cdot k_{12}}{k_{12}+E_6}}_{j^{M6}_{comp}}$<br>
$\dot{E_4} = k_4 - k_5 \underbrace{\left(\frac{E_4}{k_6+E_4}\right)}_{\approx 1} \cdot \left(\frac{k_{13}}{k_{13}+A}\right)$<br>
$\Rightarrow A^{M4}_{set} = \frac{k_5 k_{13} - k_4 k_{13}}{k_4}$<br>
$\dot{E_6} = k_8 - k_9 \underbrace{\left(\frac{E_6}{k_{10}+E_6}\right)}_{\approx 1} A \Rightarrow A^{M6}_{set} = \frac{k_8}{k_9}$<br>
<br>
Windup conditions and<br>
compensatory fluxes :<br>
(i) M4 controls $A$ :<br>
$A_{ss} = A^{M4}_{set} \Rightarrow \dot{E_6} > 0$<br>
$\Rightarrow j^{M6}_{comp} \rightarrow 0$<br>
− − − − − − −<br>
(ii) M6 controls $A$ :<br>
$A_{ss} = A^{M6}_{set} \Rightarrow \dot{E_4} > 0$<br>
$\Rightarrow j^{M4}_{comp} \rightarrow 0$<br>
− − − − − − −<br>
(iii) $A_{ss}$ is in transition zone :<br>
$A^{M4}_{set} < A_{ss} < A^{M6}_{set} \Rightarrow \dot{E_4} > 0$ & $\dot{E_6} > 0$<br>
$\Rightarrow j^{M4}_{comp} = 0$,<br>
& $j^{M6}_{comp} = 0$.<br>
$A_{ss}$ is only determined by $k_1$ and $k_2$ :<br>
$A_{ss} = \frac{k_1}{k_2}$
</td>
<td>
schematic perturbation phase diagram<br>
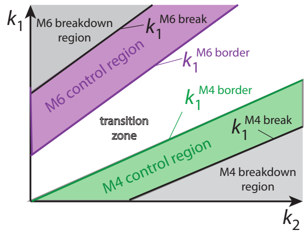

<br>
$k_1^{M4\,break}$ :<br>
$j^{M4}_{comp} = j^{M4}_{comp\,max} = k_3$ & $j^{M6}_{comp} = 0$<br>
$\Rightarrow k_1^{M4\,break} = k_2 A^{M4}_{set} - k_3$<br>
<br>
$k_1^{M4\,border}$ :<br>
$j^{M4}_{comp} = 0$ & $j^{M6}_{comp} = 0$<br>
$\Rightarrow k_1^{M4\,border} = k_2 A^{M4}_{set}$<br>
<br>
$k_1^{M6\,border}$ :<br>
$j^{M4}_{comp} = 0$ & $j^{M6}_{comp} = 0$<br>
$\Rightarrow k_1^{M6\,border} = k_2 A^{M6}_{set}$<br>
<br>
$k_1^{M6\,break}$ :<br>
$j^{M4}_{comp} = 0$ & $j^{M6}_{comp} = j^{M6}_{comp\,max}$<br>
$= k_{11} A^{M6}_{set}$<br>
$\Rightarrow k_1^{M6\,break} = (k_2 + k_{11}) A^{M6}_{set}$
</td>
</tr>
</table>

**Table S31.** Isolated control of combined M4 and M7 controllers ($A^{M4}_{set}<A^{M7}_{set}$)

| combined controllers | rate equations and compensatory fluxes | borderlines in phase diagram |
|---|---|---|
| 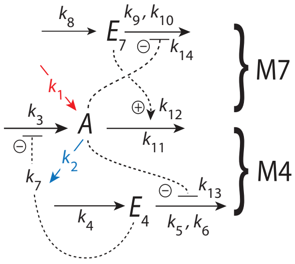<br> | Rate equations and set−points :<br>$\dot{A}=k_1-k_2A+\underbrace{k_3\left(\frac{k_7}{k_7+E_4}\right)}_{j^{M4}_{comp}}-\underbrace{\frac{k_{11}A\cdot E_7}{k_{12}+E_7}}_{j^{M7}_{comp}}$<br>$\dot{E}_4=k_4-k_5\underbrace{\left(\frac{E_4}{k_6+E_4}\right)}_{\approx 1}\cdot\left(\frac{k_{13}}{k_{13}+A}\right)$<br>$\Rightarrow A^{M4}_{set}=\frac{k_5k_{13}-k_4k_{13}}{k_4}$<br>$\dot{E}_7=k_8-k_9\underbrace{\left(\frac{E_7}{k_{10}+E_7}\right)}_{\approx 1}\left(\frac{k_{14}}{k_{14}+A}\right)$<br>$\Rightarrow A^{M7}_{set}=\frac{k_9k_{14}-k_8k_{14}}{k_8}$<br>Windup conditions and compensatory fluxes :<br>(i) M4 controls $A$ :<br>$A_{ss}=A^{M4}_{set}\Rightarrow\dot{E}_7<0$<br>$\Rightarrow j^{M7}_{comp}\to 0$<br>– – – – – – –<br>(ii) M7 controls $A$ :<br>$A_{ss}=A^{M7}_{set}\Rightarrow\dot{E}_4>0$<br>$\Rightarrow j^{M4}_{comp}\to 0$<br>– – – – – – –<br>(iii) $A_{ss}$ is in transition zone :<br>$A^{M4}_{set}<A_{ss}<A^{M7}_{set}\Rightarrow\dot{E}_4>0\ \&\ \dot{E}_7<0$<br>$\Rightarrow j^{M4}_{comp}=0,$<br>$\&\ j^{M7}_{comp}=0.$<br>$A_{ss}$ is only determined by $k_1$ and $k_2$ :<br>$A_{ss}=\frac{k_1}{k_2}$ | schematic perturbation phase diagram<br>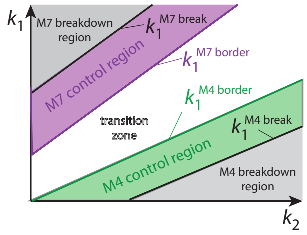<br><br>$k_1^{M4\,break}$ :<br>$j^{M4}_{comp}=j^{M4}_{comp\,max}=k_3\ \&\ j^{M7}_{comp}=0$<br>$\Rightarrow k_1^{M4\,break}=k_2A^{M4}_{set}-k_3$<br>$k_1^{M4\,border}$ :<br>$j^{M4}_{comp}=0\ \&\ j^{M7}_{comp}=0$<br>$\Rightarrow k_1^{M4\,border}=k_2A^{M4}_{set}$<br>$k_1^{M7\,border}$ :<br>$j^{M4}_{comp}=0\ \&\ j^{M7}_{comp}=0$<br>$\Rightarrow k_1^{M7\,border}=k_2A^{M7}_{set}$<br>$k_1^{M7\,break}$ :<br>$j^{M4}_{comp}=0\ \&\ j^{M7}_{comp}=j^{M7}_{comp\,max}$<br>$=k_{11}A^{M7}_{set}$<br>$\Rightarrow k_1^{M7\,break}=(k_2+k_{11})A^{M7}_{set}$ |

**Table S32.** Isolated control of combined M4 and M8 controllers ($A^{M4}_{set} < A^{M8}_{set}$)

| combined controllers | rate equations and compensatory fluxes | borderlines in phase diagram |
|---|---|---|
| 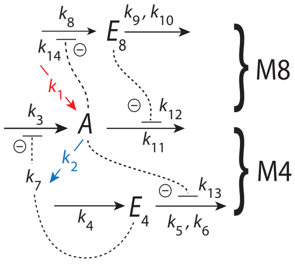<br> | Rate equations and set−points :<br>$\dot{A}=k_1-k_2A+\underbrace{k_3\left(\frac{k_7}{k_7+E_4}\right)}_{j^{M4}_{comp}}-\underbrace{\frac{k_{11}A\cdot k_{12}}{k_{12}+E_8}}_{j^{M8}_{comp}}$<br>$\dot{E_4}=k_4-k_5\underbrace{\left(\frac{E_4}{k_6+E_4}\right)}_{\approx 1}\cdot\left(\frac{k_{13}}{k_{13}+A}\right)$<br>$\Rightarrow A^{M4}_{set}=\frac{k_5k_{13}-k_4k_{13}}{k_4}$<br>$\dot{E_8}=k_8\left(\frac{k_{14}}{k_{14}+A}\right)-k_9\underbrace{\left(\frac{E_8}{k_{10}+E_8}\right)}_{\approx 1}$<br>$\Rightarrow A^{M8}_{set}=\frac{k_8k_{14}-k_9k_{14}}{k_9}$<br><br>Windup conditions and compensatory fluxes :<br>(i) M4 controls $A$ :<br>$A_{ss}=A^{M4}_{set} \Rightarrow \dot{E_8}>0$<br>$\Rightarrow j^{M8}_{comp} \rightarrow 0$<br>− − − − − − −<br>(ii) M8 controls $A$ :<br>$A_{ss}=A^{M8}_{set} \Rightarrow \dot{E_4}>0$<br>$\Rightarrow j^{M4}_{comp} \rightarrow 0$<br>− − − − − − −<br>(iii) $A_{ss}$ is in transition zone :<br>$A^{M4}_{set}<A_{ss}<A^{M8}_{set} \Rightarrow \dot{E_4}>0 \;\&\; \dot{E_8}>0$<br>$\Rightarrow j^{M4}_{comp}=0,$<br>$\&\; j^{M8}_{comp}=0.$<br>$A_{ss}$ is only determined by $k_1$ and $k_2$ :<br>$A_{ss}=\frac{k_1}{k_2}$ | schematic perturbation phase diagram<br>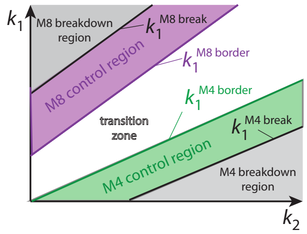<br><br><br>$k_1^{M4\,break}$ :<br>$j^{M4}_{comp}=j^{M4}_{comp\,max}=k_3 \;\&\; j^{M8}_{comp}=0$<br>$\Rightarrow k_1^{M4\,break}=k_2A^{M4}_{set}-k_3$<br><br>$k_1^{M4\,border}$ :<br>$j^{M4}_{comp}=0 \;\&\; j^{M8}_{comp}=0$<br>$\Rightarrow k_1^{M4\,border}=k_2A^{M4}_{set}$<br><br>$k_1^{M8\,border}$ :<br>$j^{M4}_{comp}=0 \;\&\; j^{M8}_{comp}=0$<br>$\Rightarrow k_1^{M8\,border}=k_2A^{M8}_{set}$<br><br>$k_1^{M8\,break}$ :<br>$j^{M4}_{comp}= \;\&\; j^{M8}_{comp}=j^{M8}_{comp\,max}$<br>$=k_{11}A^{M8}_{set}$<br>$\Rightarrow k_1^{M8\,break}=(k_2+k_{11})A^{M8}_{set}$ |
