## Supplementary material for "Flux organizations and control modes in antagonistically combined negative feedback loops": Sup2.pdf

Peter Ruoff

Department of Chemistry, Bioscience, and Environmental Engineering, University of Stavanger, Stavanger, Norway

### Supplementary material

#### Detailed description of combined controllers M2-M7, M3-M5, and M4-M8

#### Combined M2 and M7 controllers

Comparison of Fig 9 and Fig 11 shows that the M1-M5 combined controller varies dramatically in its settling times [1] dependent whether delegated control or isolated control led to positive or negative windups in both controller variables. The same behavior is also observed for the M4-M8 combined controller described below (compare there Fig S14 with Fig S16). Here the author investigates an example of 'mixed windup' for the M2-M7 combined controller, i.e. the situation when one of the $E_i$'s rate shows positive windup, while the other rate is negative (Table 1). The M2-M7 scheme is shown in Fig S1.

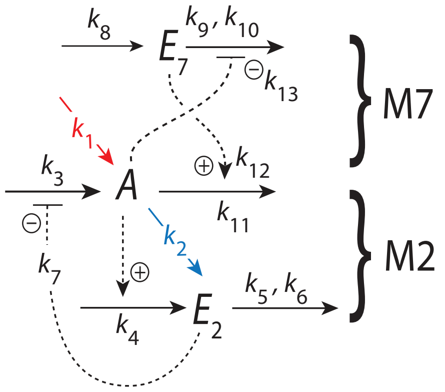


**Fig S1.** The combined M2-M7 controller. Outlined in red indicates the stepwise inflow ($k_1$) and outflow ($k_2$) perturbations applied to $A$. M7 is an outflow controller by direct activating its compensatory flux with activation constant $k_{12}$, while $A$ represses the removal of $E_7$ with inhibition constant $k_{13}$.

The rate equations are:

$$\dot{A} = \underbrace{k_1}_{\text{perturbation}} - \underbrace{k_2 \cdot A}_{\text{perturbation}} + \underbrace{k_3\left(\frac{k_7}{k_7+E_2}\right)}_{\text{M2 compensatory flux } j_3} - \underbrace{k_{11} \cdot A\left(\frac{E_7}{k_{12}+E_7}\right)}_{\text{M7 compensatory flux } j_{11}} \tag{S1}$$

[1]The settling (or response) time is defined as the time from start of a perturbation until the controlled variable $A$ has reached steady state.

$$\dot{E}_2 = k_4{\cdot}A - k_5{\cdot}\underbrace{\left(\frac{E_2}{k_6{+}E_2}\right)}_{\approx 1} \tag{S2}$$

$$\dot{E}_7 = k_8 - k_9{\cdot}\underbrace{\left(\frac{E_7}{k_{10}{+}E_7}\right)}_{\approx 1}{\cdot}\left(\frac{k_{13}}{k_{13}{+}A}\right) \tag{S3}$$

The activation of M7's compensatory flux $j_{11}$ is written as a mixed activation [1] with $k_{12}$ as the activation constant. Integral control of $A$ by M2 and M7 is incorporated by assuming zero-order kinetics with respect to $E_2$ (Eq S2) and $E_7$ (Eq S3) by setting $k_6$ and $k_{10}$ both to $1.0{\times}10^{-6}$. The setpoints of $A$ can then be calculated from the steady states of Eqs S2 and S3 by assuming that $E_2/(k_6{+}E_2)$ and $E_7/(k_{10}{+}E_7)$ are close to one, which leads to

$$A_{set}^{M2} = \frac{k_5}{k_4} \tag{S4}$$

$$A_{set}^{M7} = \frac{k_{13}}{k_8}\left(k_9 - k_8\right) \tag{S5}$$

In the following the combined M2-M7 controllers homeostatic behaviors are described as a function how $A_{set}^{M2}$ and $A_{set}^{M7}$ relate to each other.

### $A_{set}^{M2}{>}A_{set}^{M7}$: Delegated control with occasional $E_7$ windup

When $A_{set}^{M2}{>}A_{set}^{M7}$ we observe delegated control. In the example considered here $A_{set}^{M2}{=}2.0$ and $A_{set}^{M7}{=}1.0$.

Fig S2, rows a-d, shows the combined controllers' responses to different $k_1$, $k_2$ perturbations. Columns 1-4 show, respectively, the perturbation phase diagram, $A$, $E_2$ and $j_3$, and $E_7$ and $j_{11}$, the latter three all as a function of time. Row a shows four perturbations which occur *between* the regions of M2 and M7 control (plot a1). As expected (Table S7 in `Sup1.pdf`), $E_7$ shows windup when M2 controls $A$ in phases 2 and 4. When returning to the M7 controlled region from M2 control more time is needed to establish $A$ homeostasis at $A_{set}^{M7}$ (plot a2), because excess of $E_7$ needs to be removed (plot a4). In row b the perturbations occur *within* the M7 and M2 controlled regions (plot b1), which show rapid response times to achieve homeostasis at $A_{set}^{M7}$ or $A_{set}^{M2}$ (plot b2) even when $E_7$ undergoes windup in phases 3 and 4 (plot b4). In row c perturbations move the system between the M7 controlled region and the transition zone (plot c1). The response is rapid, but $E_7$ windup within the transition zone elongates the time to return back to M7 control (plots c2 and c4). Finally, row d shows that transitions between the M2 controlled region and the transition zone are rapid, despite the windup behavior of $E_7$ (plot d4).

Fig S3 shows the M2 and M7 behaviors during four phases.

In phases 1 and 2 M7 is in control of $A$ and since the steady state of $A$ is less than $A_{set}^{M2}$ (Fig S3b) $E_2$ is driven to a low value ($\dot{E}_2{<}0$, Eq S2) until $E_2$ is at a minimum (Fig S3c) and M2's compensatory flux is at its maximum $j_3{=}k_3$. The increase in $k_2$ when moving from phase 1 to phase 2 causes $E_7$ to decrease (Fig S3d) and to reduce M7's compensatory flux $j_{11}$ (Eq S1). When $k_2$ becomes too large M7 breaks down as $E_7$ and $j_{11}$ go to zero. In the phase diagram (Fig S3a) the borderline of M7's breakdown, $k_1^{M7\,break}$, is described by the relationship:

The border lines in the plots of column 1 separating regions of control, the transition zone, and regions of M2 and M7 breakdown are described by the following equations (see also Table S23 in `Sup1.pdf`).

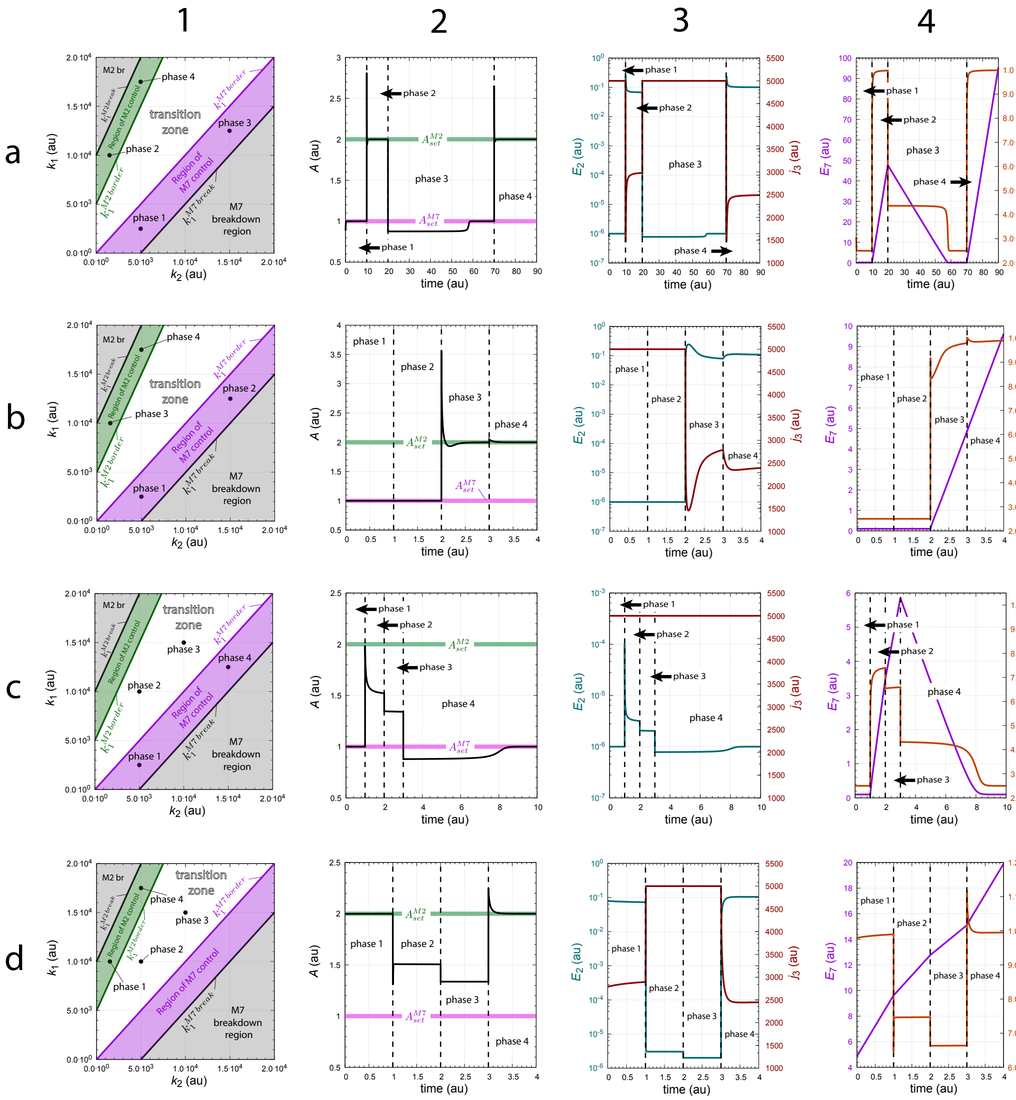


**Fig S2.** Response times in the combined M2M7 controller with occasional windup in $E_7$. Rows a-d show different perturbation sequences. Row a: phase 1, $k_1$=2.5×10$^3$, $k_2$=5.0×10$^3$; phase 2, $k_1$=1.0×10$^4$, $k_2$=1.5×10$^3$; phase 3, $k_1$=1.25×10$^4$, $k_2$=1.5×10$^4$; phase 4, $k_1$=1.75×10$^4$, $k_2$=5.0×10$^3$. Row b: phase 1, $k_1$=2.5×10$^3$, $k_2$=5.0×10$^3$; phase 2, $k_1$=1.25×10$^4$, $k_2$=1.5×10$^4$; phase 3, $k_1$=1.0×10$^4$, $k_2$=1.5×10$^3$; phase 4, $k_1$=1.75×10$^4$, $k_2$=5.0×10$^3$. Row c: phase 1, $k_1$=2.5×10$^3$, $k_2$=5.0×10$^3$; phase 2, $k_1$=1.0×10$^4$, $k_2$=5.0×10$^3$; phase 3, $k_1$=1.55×10$^4$, $k_2$=1.0×10$^4$; phase 4, $k_1$=1.25×10$^4$, $k_2$=1.5×10$^4$. Row d: phase 1, $k_1$=1.0×10$^4$, $k_2$=1.5×10$^3$; phase 2, $k_1$=1.0×10$^4$, $k_2$=5.05×10$^3$; phase 3, $k_1$=1.5×10$^4$, $k_2$=1.0×10$^4$; phase 4, $k_1$=1.75×10$^4$, $k_2$=5.0×10$^3$. Columns 1-4 show respectively perturbation phase diagrams, the concentration of $A$ as a unction of time, and $E_2$ (left ordinates), $j_3$ (right ordinates), as well as $E_7$ (left ordinates), $j_{11}$ (right ordinates) as a function of time. Rate constants: $k_3$=5.0×10$^3$, $k_4$=10.0, $k_5$=20.0, $k_6$=1.0×10$^{-6}$, $k_7$=0.1, $k_8$=10.0, $k_9$=110.0, $k_{10}$=1.0×10$^{-6}$, $k_{11}$=5.0×10$^3$, $k_{12}$=$k_{13}$=0.1. Initial concentrations, row a: $A_0$=1.0, $E_{2,0}$=1.0×10$^{-6}$, and $E_{7,0}$=0.23; initial concentrations, row b and c: $A_0$=1.0, $E_{2,0}$=1.0×10$^{-6}$, and $E_{7,0}$=0.1; Initial concentrations, row d: $A_0$=2.0, $E_{2,0}$=7.92×10$^{-2}$, and $E_{7,0}$=4.862.

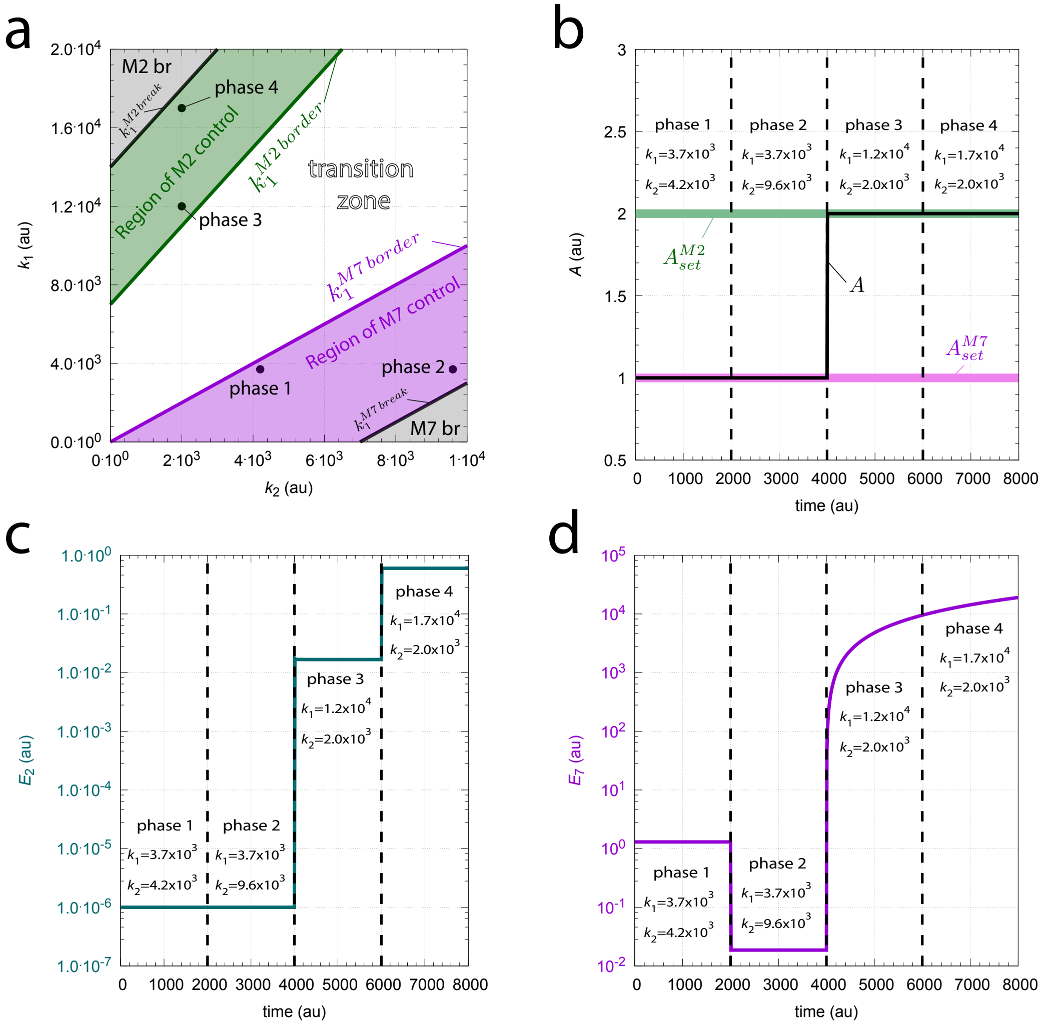


**Fig S3.** Delegated control between M2 and M7 when $A_{set}^{M2}>A_{set}^{M7}$. Panel a shows the $k_1$-$k_2$ phase diagram with regions of M2 and M7 control, the transition zone, the regions of M2 and M7 breakdown (respectively described as 'M2 br' and 'M7 br'), and the borderlines separating the different regions (see also main text). Panel b shows $A$ as a function of time during the four phases, while panels c and d show respectively $E_2$ and $E_7$ as a function of time. Values of $k_1$ and $k_2$ are indicated inside the figures. Other rate constants: $k_3$=7.0×10$^3$, $k_4$=10.0, $k_5$=20.0, $k_6$=1.0×10$^{-6}$, $k_7$=0.1, $k_8$=10, $k_9$=110.0, $k_{10}$=1.0×10$^{-6}$, $k_{11}$=7.0×10$^3$, $k_{12}$=$k_{13}$=0.1. Initial concentrations: $A_0$=1.0, $E_{2,0}$=1.0×10$^{-6}$, and $E_{7,0}$=1.30. See `Python code` for details.

When $k_2$ becomes too large M7 breaks down as $E_7$ and $j_{11}$ go to zero. In the phase diagrams the borderline of M7's breakdown, $k_1^{M7\,break}$, is described by the relationship:

$$k_1^{M7\,break} = k_2 \cdot A_{set}^{M7} - k_3 \tag{S6}$$

The border between the M7 control region and the transition zone, $k_1^{M7\,border}$, is characterized by $j_{11}$ reaching $j_{11}$=$k_{11} \cdot A_{set}^{M7}$, which defines this borderline to:

$$k_1^{M7\,border} = k_2 \cdot A_{set}^{M7} - k_3 + k_{11} \cdot A_{set}^{M7} \tag{S7}$$

When $k_1$, $k_2$ values move into the transition zone $E_7$ will undergo windup and keep M7's compensatory flux at a maximum while $A_{ss}$ values change according to the values of $k_1$ and $k_2$ to (check with $A$-plots c2 and d2, phases 2 and 3, in Fig S2):

$$A_{ss} = \frac{k_1+k_3}{k_2+k_{11}} \tag{S8}$$

At the border to M2's control region $j_3$ is still at its maximum, but $A_{ss}$ is now at $A_{set}^{M2}$. Accordingly, the borderline between the M2 control region and the transition zone, $k_1^{M2\,border}$ becomes:

$$k_1^{M2\,border} = k_2{\cdot}A_{set}^{M2} - k_3 + k_{11}{\cdot}A_{set}^{M2} \tag{S9}$$

Finally, when perturbative inflows to $A$ become too large M2's compensatory flux will go to zero and $E_2$ will show windup. The borderline of M2's breakdown, $k_1^{M2\,break}$, is described by

$$k_1^{M2\,break} = k_2{\cdot}A_{set}^{M2} + k_{11}{\cdot}A_{set}^{M2} \tag{S10}$$

In Fig S4 the behavior of the M2-M7 combined system is shown when the perturbation changes along a straight trajectory in the $k_1$-$k_2$ phase diagram. The system starts at $k_1^{start}$=1.0×10$^2$, $k_2^{start}$=1.0×10$^4$ and ends at $k_1^{end}$=1.0×10$^4$, $k_2^{end}$=1.0×10$^2$ (blue line in panel a). The black dots 1-4 indicate the (calculated) intersection points with the four borderlines Eqs S6-S10. The combined controller shows an apparent setpoint change (rheostasis [2,3]) as $k_1$ and $k_2$ change (Fig S4b). Figs S4c and d show the respective behaviors of $E_2$ and $E_7$ together with their compensatory fluxes $j_3$ and $j_{11}$. Note that during M7 control M2's compensatory flux is at its maximum, while during M2 control M7's compensatory flux is $j_{11}$=$k_{11}{\cdot}A_{set}^{M2}$.

#### $A_{set}^{M2}$<$A_{set}^{M7}$: Isolated control with occasional $E_2$ windup

When $A_{set}^{M2}$<$A_{set}^{M7}$ the controllers cooperate via isolated control, i.e. whenever M2 or M7 determines $A$'s setpoint the other controller will have a negligible compensatory flux. Thus, M2 or M7 control $A$ alone in isolation.

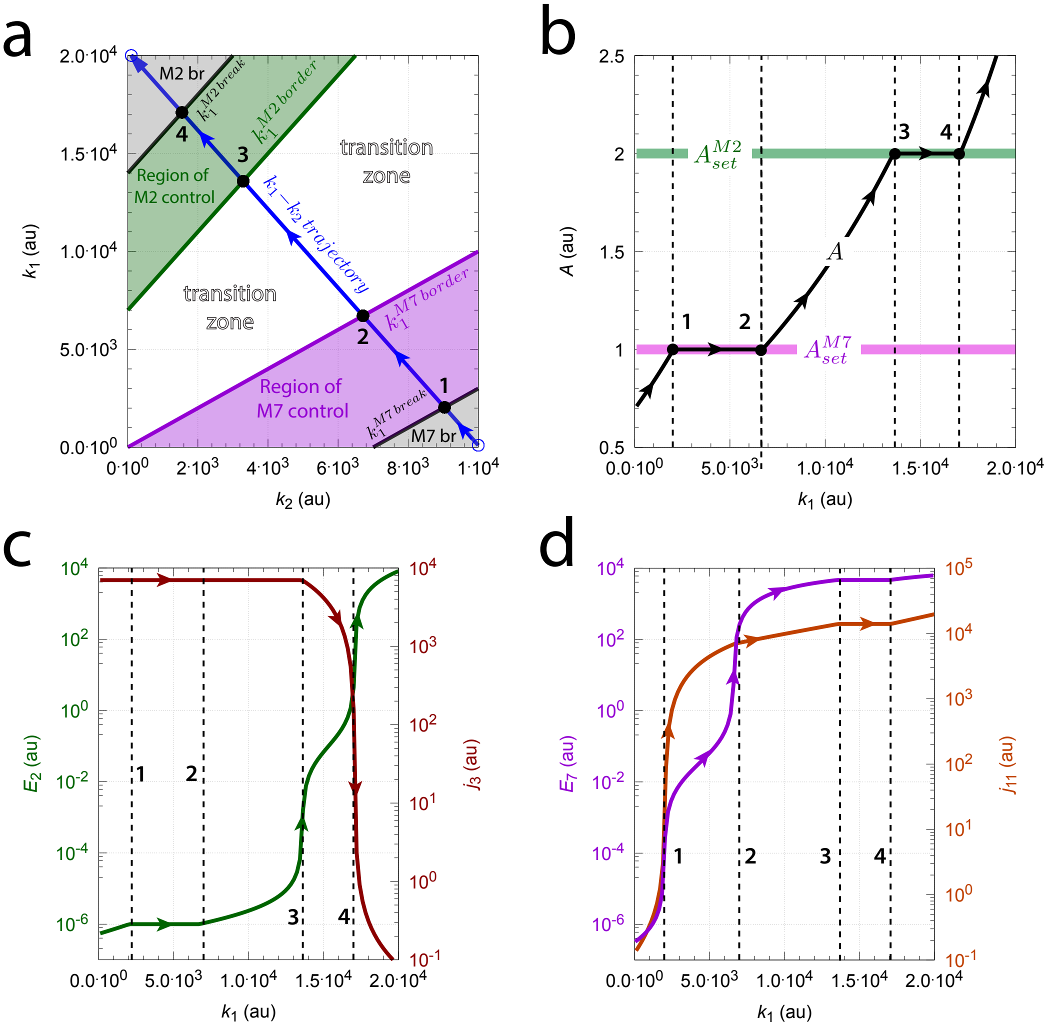


**Fig S4.** Apparent setpoint change (rheostasis) in $A$ when $k_1$ and $k_2$ move along a straight trajectory within the phase diagram. Panel a shows the trajectory in the phase diagram outlined in blue. Solid dots 1-4 indicate the intersections of the trajectory with the borderlines. Otherwise this phase diagram is identical with that in Fig S3a. Panel b shows the change of $A$ as a function of the trajectory's $k_1$ values. Panel c: $E_2$ (left ordinate) and M2's compensatory flux $j_3$ (right ordinate) as a function of $k_1$. Panel d: $E_7$ (left ordinate) and M7's compensatory flux $j_{11}$ as a function of $k_1$. Rate constants as in Fig S3. Initial concentrations: $A_0$=1.0, $E_{2,0}$=1.0×$10^{-6}$, and $E_{7,0}$=1.0. 'M2 br': M2 breakdown region; 'M7 br': M7 breakdown region.

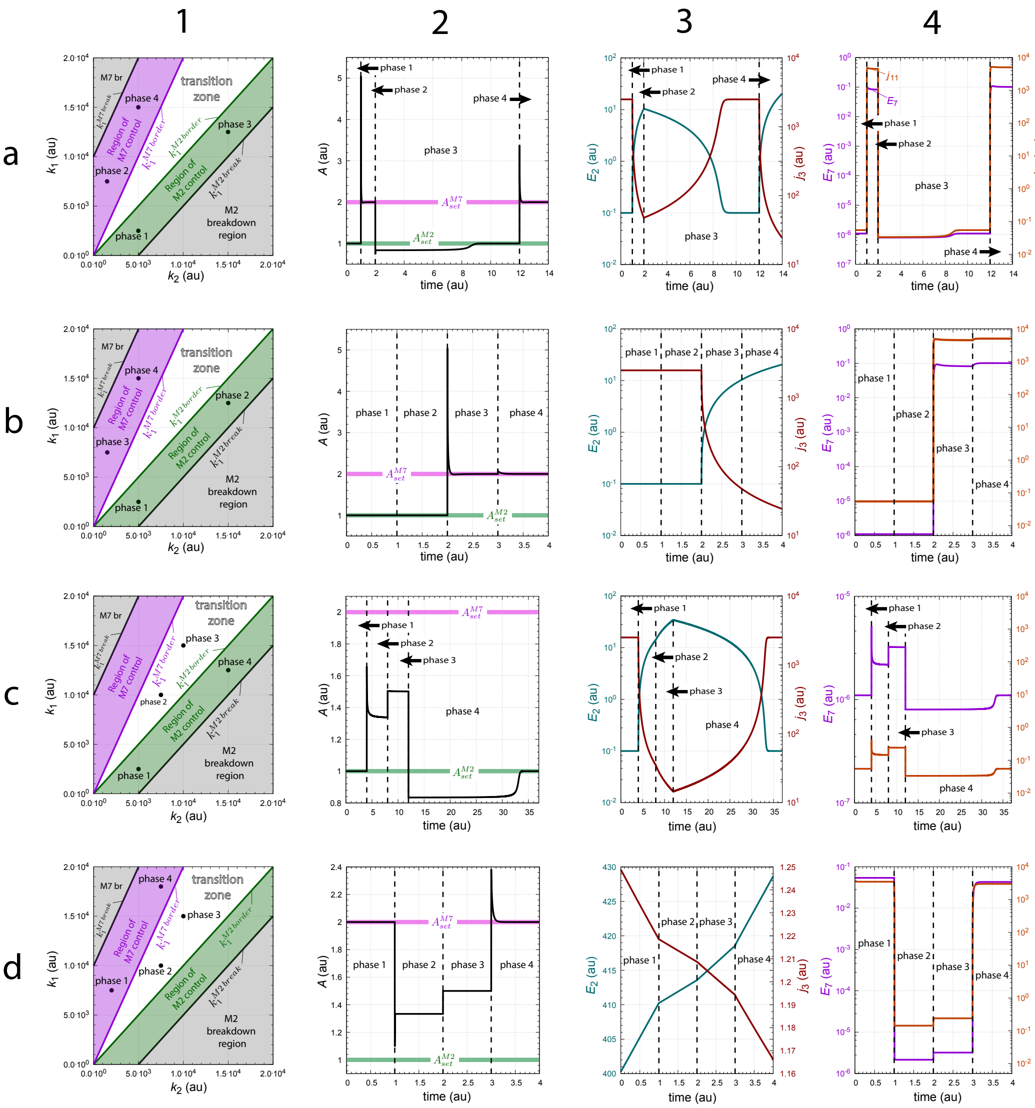


**Fig S5.** Response times in the combined M2M7 controller with occasional windup in $E_2$. Rows a-d show different perturbation sequences. Row a: phase 1, $k_1$=2.5×10$^3$, $k_2$=5.0×10$^3$; phase 2, $k_1$=7.5×10$^3$, $k_2$=1.52×10$^3$; phase 3, $k_1$=1.25×10$^4$, $k_2$=1.5×10$^4$; phase 4, $k_1$=1.5×10$^4$, $k_2$=5.0×10$^3$. Row b: phase 1, $k_1$=2.5×10$^3$, $k_2$=5.0×10$^3$; phase 2, $k_1$=1.25×10$^4$, $k_2$=1.5×10$^4$; phase 3, $k_1$=7.5×10$^3$, $k_2$=1.52×10$^3$; phase 4, $k_1$=1.5×10$^4$, $k_2$=5.0×10$^3$. Row c: phase 1, $k_1$=2.5×10$^3$, $k_2$=5.0×10$^3$; phase 2, $k_1$=1.0×10$^4$, $k_2$=7.5×10$^3$; phase 3, $k_1$=1.5×10$^4$, $k_2$=1.0×10$^4$; phase 4, $k_1$=1.25×10$^4$, $k_2$=1.5×10$^4$. Row d: phase 1, $k_1$=7.5×10$^3$, $k_2$=2.0×10$^3$; phase 2, $k_1$=1.0×10$^4$, $k_2$=7.5×10$^3$; phase 3, $k_1$=1.5×10$^4$, $k_2$=1.0×10$^4$; phase 4, $k_1$=1.8×10$^4$, $k_2$=7.5×10$^3$. Columns 1-4 show respectively perturbation phase diagrams, the concentration of $A$ as a function of time, and $E_2$ (left ordinates), $j_3$ (right ordinates), as well as $E_7$ (left ordinates), $j_{11}$ (right ordinates) as a function of time. Rate constants: $k_3$=5.0×10$^3$, $k_4$=10.0, $k_5$=10.0, $k_6$=1.0×10$^{-6}$, $k_7$=0.1, $k_8$=10.0, $k_9$=210.0, $k_{10}$=1.0×10$^{-6}$, $k_{11}$=5.0×10$^3$, $k_{12}$=$k_{13}$=0.1. Initial concentrations, rows a-c: $A_0$=1.0, $E_{2,0}$=1.0×10$^{-1}$, and $E_{7,0}$=1.1×10$^{-6}$; initial concentrations, row d: $A_0$=2.0, $E_{2,0}$=1.0×10$^{-6}$, and $E_{7,0}$=0.1; Initial concentrations, row d: $A_0$=2.0, $E_{2,0}$=4.0×10$^2$, and $E_{7,0}$=5.38×10$^{-2}$.

When $A_{set}^{M2}<A_{set}^{M7}$ the controllers cooperate via isolated control, i.e. whenever M2 or

M7 determines $A$'s setpoint the other controller will have a negligible compensatory flux. Thus, M2 or M7 control $A$ alone in isolation. Fig S6 shows the M2-M7 system's behavior when M2 or M7 controls $A$.

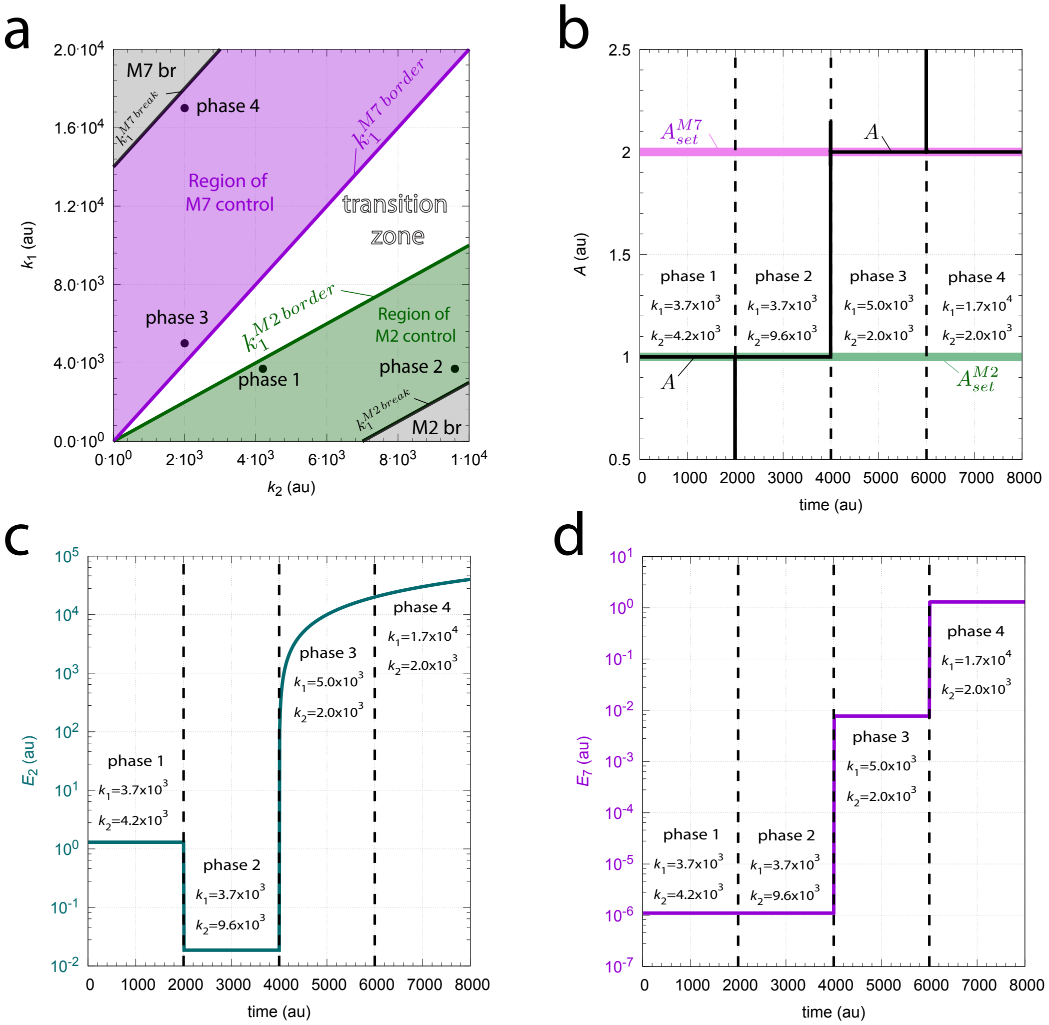


**Fig S6.** Isolated control in the combined M2-M7 controller. Panel a: $k_1$-$k_2$ phase diagram with regions of M2/M7 control, breakdown, and the transition zone. Panel b: During phases 1 and 2 $k_1$ and $k_2$ values are such that $A$ is controlled by M2, while during phases 3 and 4 M7 controls $A$. Panel c: Increased $k_2$ values are met by derepression of $E_2$ and thereby increasing M2's compensatory flux. During M7 control (phases 3 and 4) $E_2$ goes into windup leading to a negligible M2 compensatory flux. Panel d: Negligible $E_7$ is observed during control by M2 (phases 1 and 2). This leads to a low and negligible compensatory flux $j_{11}$. When M7 takes control of $A$ during phases 3 and 4 an increase of $E_7$ and M7's compensatory flux $j_{11}$ opposes the increased $k_1$ levels. Rate constants: $k_3$=7.0×10$^3$, $k_4$=10.0, $k_5$=10.0, $k_6$=1.0×10$^{-6}$, $k_7$=0.1, $k_8$=10.0, $k_9$=210.0, $k_{10}$=1.0×10$^{-6}$, $k_{11}$=7.0×10$^3$, $k_{12}$=$k_{13}$=0.1. Initial concentrations: $A_0$=1.0, $E_{2,0}$=1.3, and $E_{7,0}$=1.0×10$^{-6}$. See `Python code` for details.

Since M7's compensatory flux is negligible when M2 controls $A$ the breakdown border of M2 (upon increased $k_2$ values) is reached when $j_3$ is at its maximum compensatory flux $j_3$=$k_3$ and $A_{ss}$ is still at $A_{set}^{M2}$. This defines $k_1^{M2\,break}$ as:

$$k_1^{M2\,break} = k_2{\cdot}A_{set}^{M2} - k_3 \quad \text{(S11)}$$

The border between the M2 control region and the transition zone is characterized that $j_3$ is negligible which leads to

$$k_1^{M2\,border} = k_2{\cdot}A_{set}^{M2} \quad \text{(S12)}$$

The M7 controlled region is enclosed by two boundaries. The boundary adjacent to the transition zone, $k_1^{M2\,border}$, has a negligible compensatory flux $j_{11}$ such that

$$k_1^{M7\,border} = k_2{\cdot}A_{set}^{M7} \quad \text{(S13)}$$

Finally, when the inflow to $A$ becomes too large the M7 controller will reach the compensatory flux $j_{11}^{max}{=}k_{11}{\cdot}A_{set}^{M7}$ and results in the borderline $k_1^{M7\,break}$:

$$k_1^{M7\,break} = k_2{\cdot}A_{set}^{M7} + k_{11}A_{set}^{M7} = (k_2 + k_{11})A_{set}^{M7} \quad \text{(S14)}$$

Fig S7a shows a calculation where a $k_1$-$k_2$ trajectory covers all five regions in the phase diagram including the regions of breakdown 'M2 breakdown region' (abbreviated M2 br) and 'M7 breakdown region' (abbreviated M7 br). The solid dots 1-4 show the calculated intersection points with the four borderlines Eqs S11-S14.

Panel b of Fig S7 shows the change of $A$ as a function of $k_1$ showing the $k_1$ intervals of control by M2 and M7, the crossing of the transition zone and the controllers' breakdown in the regions M2 br and M7 br. Finally, panels c and d show that whenever M2 or M7 controls $A$, the compensatory flux of the antagonistic controller is low/negligible. In other words, panel c shows the low M2 compensatory flux $j_3$ between intersection points 3 and 4 while M7 has control of $A$. Panel d shows the low M7 compensatory flux between intersection points 1 and 2, when M2 has control of $A$.

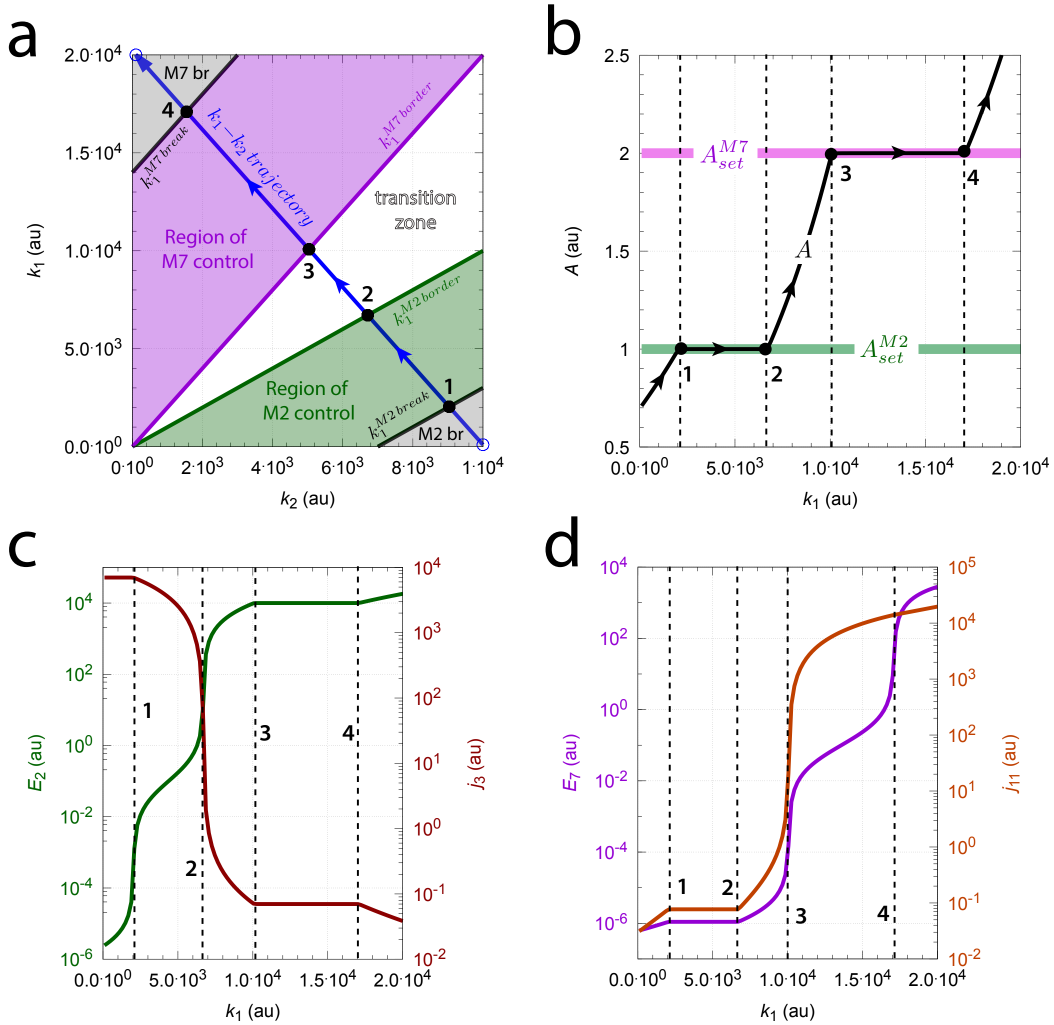


**Fig S7.** Apparent setpoint change and isolated control of $A$ along a $k_1$-$k_2$ trajectory in phase diagram. Start values: $k_1^{start}$=100.0, $k_2^{start}$=1×10$^4$; end values: $k_1^{end}$=2×10$^4$, $k_2^{end}$=100.0. Panel a: the $k_1$-$k_2$ phase diagram. The $k_1$-$k_2$ trajectory is outlined in blue. Solid dots 1-4 indicate the intersection points with the borderlines. Panel b: $A$ as a function of $k_1$. Panel c: $E_2$ as a function of $k_1$. Panel d: $E_7$ as a function of $k_1$. Other rate constants as in Fig S6. Initial concentrations: $A_0$=0.71, $E_{2,0}$=2.4×10$^{-6}$, and $E_{7,0}$=6.3×10$^{-7}$.

### Combined M3 and M5 controllers

Fig S8 shows the reaction scheme of the combined M3-M5 controller.

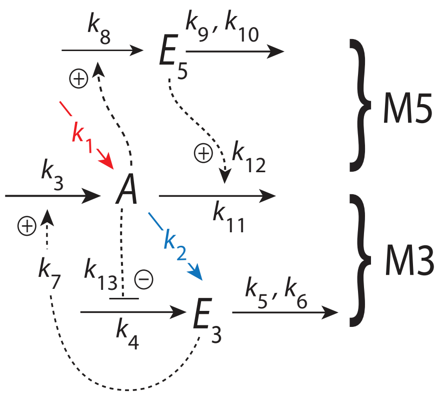


**Fig S8.** The combined M3-M5 controller. Outlined in red indicates the stepwise inflow ($k_1$) and outflow ($k_2$) perturbations applied to the system.

The rate equations are:

$$\dot{A} = \underbrace{k_1}_{\text{perturbation}} - \underbrace{k_2{\cdot}A}_{\text{perturbation}} + \underbrace{k_3\left(\frac{E_3}{k_7+E_3}\right)}_{\text{M3 compensatory flux } j_3} - \underbrace{k_{11}{\cdot}A\left(\frac{E_5}{k_{12}+E_5}\right)}_{\text{M5 compensatory flux } j_{11}} \tag{S15}$$

$$\dot{E}_3 = \left(\frac{k_{13}}{k_{13}+A}\right)k_4 - k_5\underbrace{\left(\frac{E_3}{k_6+E_3}\right)}_{\approx 1} \tag{S16}$$

$$\dot{E}_5 = k_8{\cdot}A - k_9\underbrace{\left(\frac{E_5}{k_{10}+E_5}\right)}_{\approx 1} \tag{S17}$$

Integral control and robust perfect adaptation is incorporated by using $k_6$=$1{\times}10^{-6}$ and $k_{10}$=$1{\times}10^{-6}$, which leads to $E_3/(k_6{+}E_3) \approx 1$ and $E_5/(k_{10}{+}E_5) \approx 1$. The setpoints of the two controllers are then found by setting Eqs S16 and S17 to zero and solving for the steady states values of $A$, which gives

$$A_{set}^{M3} = \frac{k_4 k_{13} - k_5 k_{13}}{k_5} \tag{S18}$$

and

$$A_{set}^{M5} = \frac{k_9}{k_8} \tag{S19}$$

**Delegated control ($A_{set}^{M3}$>$A_{set}^{M5}$)**

In the following calculations $A_{set}^{M3}$=2.0 and $A_{set}^{M5}$=1.0. When M3 controls $A$ $E_5$ shows (positive) windup, i.e. M5's compensatory flux is at its maximum with $j_{11}$=$k_{11}A_{set}^{M3}$ and $j_3$ is adjusted such that $A_{ss}$ is at $A_{set}^{M3}$ (Fig 6b). On the other hand, when M5 controls $A$ at $A_{set}^{M5}$ $E_3$ goes into windup $j_3$=$k_3$ is at its maximum and M3 adjusts its compensatory flux such that $A_{ss}$ is at $A_{set}^{M5}$ (Fig 6a). However, since both controller variables go into windup when M3 or M5 take control over $A$ (Table 1) the time required to settle between $A_{set}^{M3}$ and $A_{set}^{M5}$ increases with each transition.

The borderlines in Fig S9b can be derived as follows. When M3 is in control of $A$ $E_5$ shows windup which saturates $E_5/(k_{12}+E_5)$ such that compensatory flux becomes $j_{11}$=$k_{11}\cdot A_{set}^{M3}$ (Eq S15). With increasing inflows to $A$ by $k_1$ the adjusting compensatory flux $j_3$ by M3 (Eq S15) is diminished. M3 control breaks down when $j_3 \rightarrow 0$, which defines the borderline $k_1^{M3\,break}$. Finding $k_1$ as a function of $k_2$ from Eq S15 gives:

$$k_1^{M3\,break} = (k_2 + k_{11})A_{set}^{M3} \tag{S20}$$

Likewise, when M5 controls $A$ at $A_{set}^{M5}$ $E_3$ goes into windup such that M3's compensatory flux is at its maximum, i.e. $j_3$=$k_3$. As $k_2$ increases M5's compensatory flux $j_{11}$ decreases in order to keep $A$ at $A_{set}^{M5}$. Breakdown of M5 occurs when $j_{11} \rightarrow 0$. Solving for $k_1$ from Eq S15 when $j_{11}$ is formally zero leads to the borderline $k_1^{M5\,break}$:

$$k_1^{M5\,break} = k_2 A_{set}^{M3} - k_3 \tag{S21}$$

In the transition zone both $E_3$ and $E_5$ undergo windup, such that

$$k_1 - k_2 \cdot Ass + k_3 - k_{11}A_{ss} = 0 \;\;\Rightarrow\;\; A_{ss} = \frac{k_1 + k_3}{k_2 + k_{11}} \tag{S22}$$

Finally, the borderlines $k_1^{M3\,border}$ and $k_1^{M5\,border}$ adjacent to the transition zone are found by setting $A_{ss}$ in Eq S22 to either $A_{set}^{M3}$ or $A_{set}^{M5}$, which gives:

$$k_1^{M3\,border} = (k_2 + k_{11})A_{set}^{M3} - k_3 \tag{S23}$$

and

$$k_1^{M5\,border} = (k_2 + k_{11})A_{set}^{M5} - k_3 \tag{S24}$$

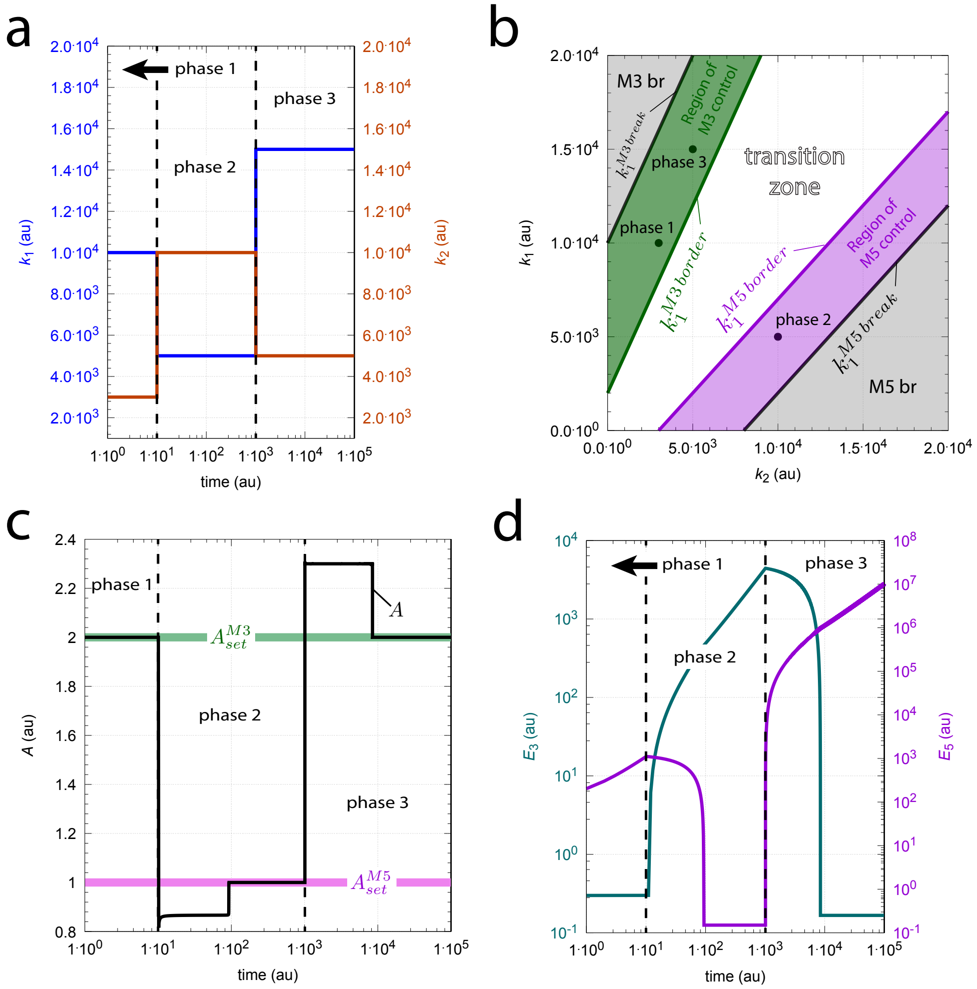


**Fig S9.** In the combined M3-M5 controller delegated control control leads to increased settling times when switching between M3 and M5 controlled states. Panel a: Steps of $k_1$ and $k_2$ as a function of time. Phase 1: $k_1$=1.0×10$^4$, $k_2$=3.0×10$^3$; phase 2: $k_1$=5.0×10$^3$, $k_2$=1.0×10$^4$; phase 3: $k_1$=1.5×10$^4$, $k_2$=5.0×10$^3$. Panel b: Perturbation phase diagram with indicated $k_1$ and $k_2$ values as solid dots. 'M3 br' and 'M5 br' indicated the breakdown regions of respectively M3 and M5 controllers. For borderline expressions see main text and Table S9. Panel c: $A$ as a function of time. Panel d: $E_3$ and $E_5$ as a function of time. Windup regions are indicated by the continuous increase of $E_3$ and $E_5$. Rate constants: $k_3$=8.0×10$^3$, $k_4$=100.0, $k_5$=4.7619, $k_6$=1.0×10$^{-6}$, $k_7$=0.1, $k_8$=$k_9$=100.0, $k_{10}$=1.0×10$^{-6}$, $k_{11}$=5.0×10$^3$, $k_{12}$=$k_{13}$=0.1. Initial concentrations: $A_0$=2.0, $E_{3,0}$=0.3, and $E_{5,0}$=1.0223×10$^2$. See `Python code` for details.

Fig S10 shows how $A$, the control variables $E_3$ and $E_5$, and the compensatory fluxes $j_3$ and $j_{11}$ change as $k_1$ and $k_2$ values move along the blue line across the perturbation phase diagram in panel a.

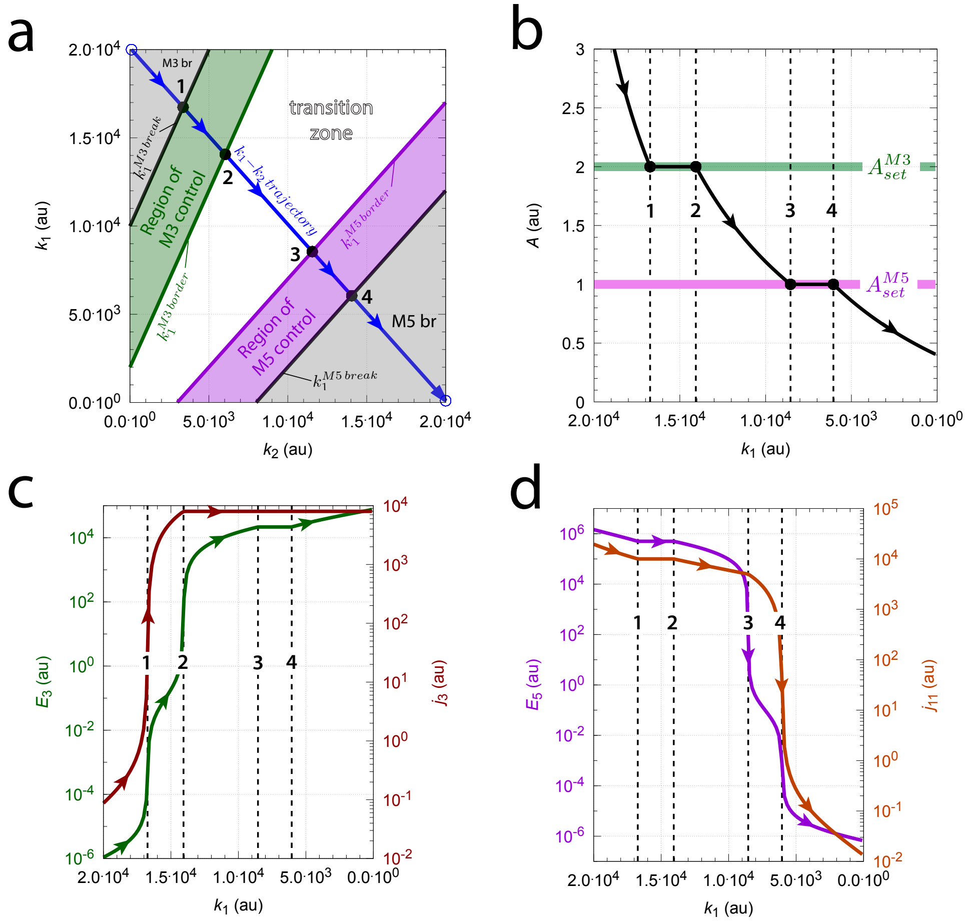


**Fig S10.** The compensatory fluxes and $A$ changes in a combined delegated M3-M5 controller when $k_1$ and $k_2$ move across the perturbation phase diagram. Panel a: Perturbation phase diagram. Blue trajectory starts at $k_{1,start}$=2.0×10$^4$ and $k_{2,start}$=100.0 and ends at $k_{1,final}$=100.0 and $k_{2,final}$=2.0×10$^4$. The line is divided into 100 data points and for each point $A$, $E_3$, $j_3$, $E_5$, and $j_{11}$ are successively calculated. Panel b: $A$ as a function of (decreasing) $k_1$. Vertical dashed lines 1-4 correspond to the $k_1$ values of the intersection points 1-4 in panel a. Panel c: $E_3$ (left ordinate) and $j_3$ (right ordinate) as a function of $k_1$. Panel d: $E_5$ (left ordinate) and $j_{11}$ (right ordinate) as a function of $k_1$. Rate constants and initial concentrations as in Fig S10.

Solid points 1-4 in Fig S10a are the calculated intersection points between the blue trajectory and the four border lines. Typically for delegated control: during M3 control $j_{11}$ is fully developed (Fig S10d) while $j_3$ increases with decreasing $k_1$ (Fig S10c). When M5 controls $A$ $j_3$ is fully developed while $j_{11}$ is adjusted to keep $A_{ss}$ at $A_{set}^{M5}$.

#### Isolated control ($A_{set}^{M3}$<$A_{set}^{M5}$)

In the following calculations setpoints are switched, i.e. $A_{set}^{M3}$=1.0 and $A_{set}^{M5}$=2.0. The system is now in isolated control mode and both controller show negative windup (Table 1, blue entry). Due to the negative windup behavior one of the controller stays silent when the other controller takes control of $A$. Fig S11 shows a computation when the same perturbations as in Fig S9 are applied. Settling times are remarkably shorter, because positive windup is absent.

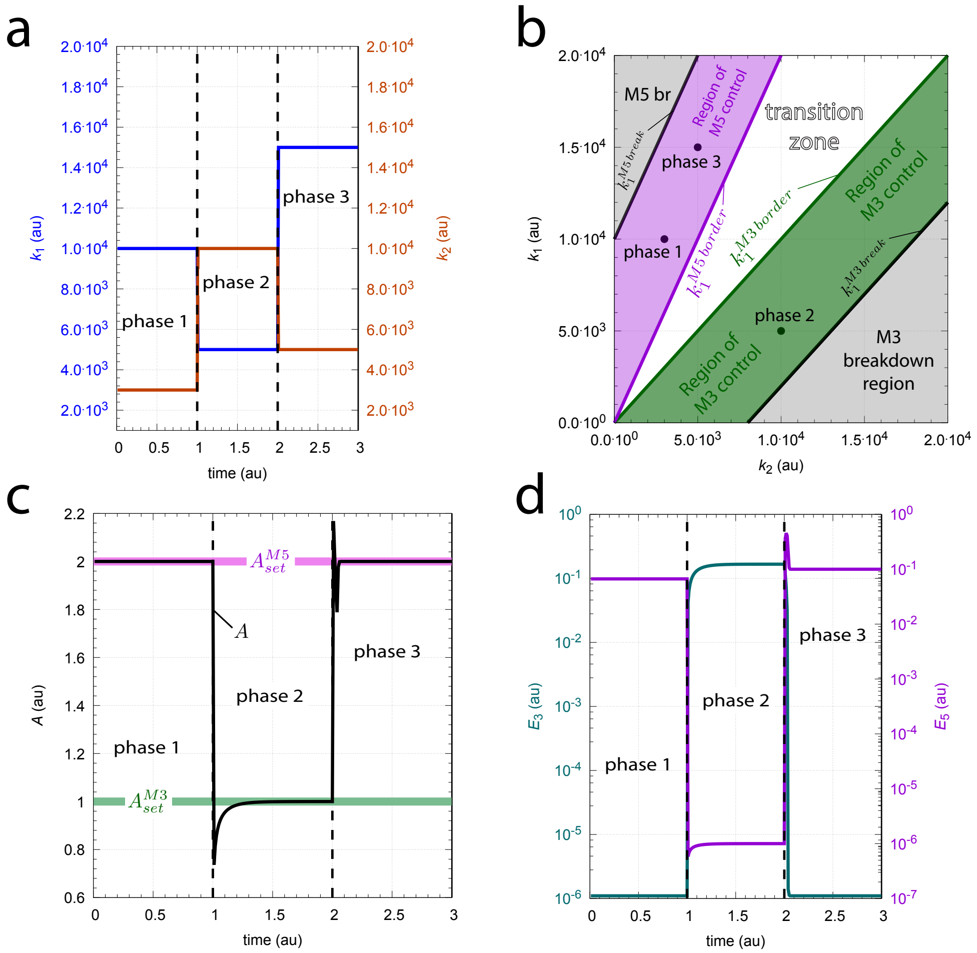


**Fig S11.** Isolated control mode in the M3-M5 controller has negative windups and shows rapid settling times. Panel a: Same $k_1$, $k_2$ steps as in Fig S9. Panel b: Perturbation phase diagram with borderlines. 'M5 br' denotes the M5 breakdown region. Panel c: $A$ as a function of time. Panel d: $E_3$ (left ordinate) and $E_5$ (right ordinate) as a function of time. Note the differences in settling times between panels c and d and Figs S9c, d. Rate constants: $k_3$=8.0×10$^3$, $k_4$=100.0, $k_5$=9.0909, $k_6$=1.0×10$^{-6}$, $k_7$=0.1, $k_8$=100.0, $k_9$=200.0, $k_{10}$=1.0×10$^{-6}$, $k_{11}$=5.0×10$^3$, $k_{12}$=$k_{13}$=0.1. Initial concentrations: $A_0$=2.0, $E_{3,0}$=1.1×10$^{-6}$, and $E_{5,0}$=6.667×10$^{-2}$. See `Python code` for details.

Characteristically, when $k_1$ and $k_2$ values stay within the transition zone both controllers are silent and the steady state of $A$ is determined by $A_{ss}$=$k_1/k_2$. From this relationship the borderlines of M3 and M5 control adjacent to the the transition zone can be determined by setting $A_{ss}$=$A_{set}^{M3}$ or $A_{ss}$=$A_{set}^{M5}$. This leads to:

$$k_1^{M3\,border} = k_2 \cdot A_{set}^{M3} \tag{S25}$$

and

$$k_1^{M5\,border} = k_2 \cdot A_{set}^{M5} \tag{S26}$$

The breakdown regions for M3 and M5 can be deduced as follows. When M3 controls $A$ compensatory flux $j_{comp}^{M5}$ (=$j_{11}$) is negligible. Thus, breakdown of M3 occurs when $j_{comp}^{M3}$ (=$j_3$=$k_3 \cdot E_3/(k_7+E_3)$) reaches its maximum value $k_3$ as $E_3/(k_7+E_3)$ becomes saturated ($\approx 1$). From the rate equation $\dot{A}$ we get:

$$\dot{A} = k_1 - k_2 \cdot A_{set}^{M3} + k_3 = 0 \;\Rightarrow\; k_1^{M3\,break} = k_2 \cdot A_{set}^{M3} - k_3 \tag{S27}$$

Likewise, when M5 controls $A$ M3's compensatory flux $j_3$ is negligible and M5 breakdown occurs when its compensatoty flux $j_{11}$ becomes saturated at $A_{ss}$=$A_{set}^{M5}$, i.e. when $E_5/(k_{12}+E_5) \approx 1$. From the rate equation of $A$ we can write

$$\dot{A} = k_1 - k_2 \cdot A_{set}^{M5} - k_{11} A_{set}^{M5} = 0 \;\Rightarrow\; k_1^{M5\,break} = (k_2 + k_{11}) \cdot A_{set}^{M3} \tag{S28}$$

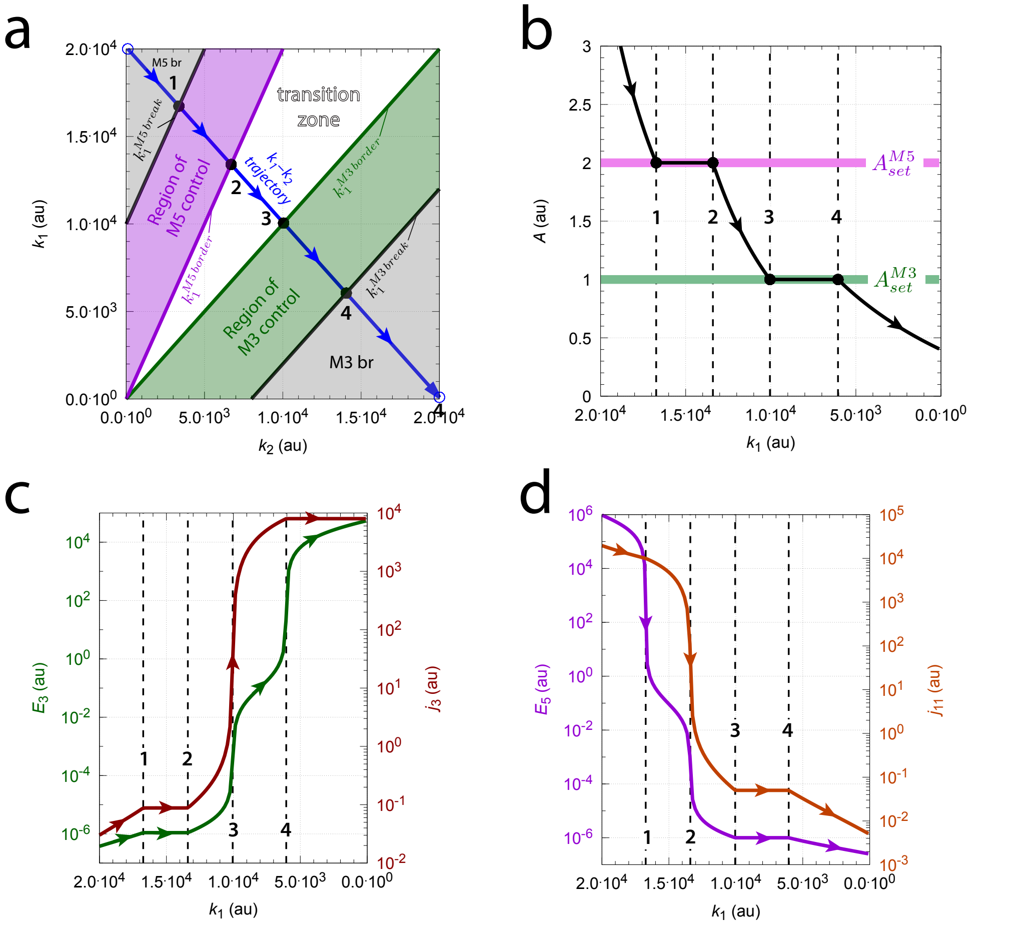


**Fig S12.** The compensatory fluxes and $A$ changes in a combined M3-M5 controller with isolated control mode when $k_1$ and $k_2$ move across the perturbation phase diagram. Panel a: Perturbation phase diagram. Blue trajectory starts at $k_{1,start}$=2.0×10$^4$ and $k_{2,start}$=100.0 and ends at $k_{1,final}$=100.0 and $k_{2,final}$=2.0×10$^4$. Same calculation procedure as described in Fig S10. Panel b: $A$ as a function of (decreasing) $k_1$. Vertical dashed lines 1-4 correspond to the $k_1$ values of the intersection points 1-4 in panel a. Panel c: $E_3$ (left ordinate) and $j_3$ (right ordinate) as a function of $k_1$. Panel d: $E_5$ (left ordinate) and $j_{11}$ (right ordinate) as a function of $k_1$. Rate constants and initial concentrations as in Fig S11.

Fig S12 shows $A$, $E_3$ and $E_5$, and compensatory flux behaviors when M3-M5 is in isolated control mode and $k_1$, $k_2$ sweep across the perturbation phase diagram along the indicated straight blue line. Typically for isolated control mode is that one of the compensatory fluxes is negligible when the other controller determines the steady state of $A$.

### Combined M4 and M8 controllers

Fig S13 shows the reaction scheme of the combined M2-M8 controller.

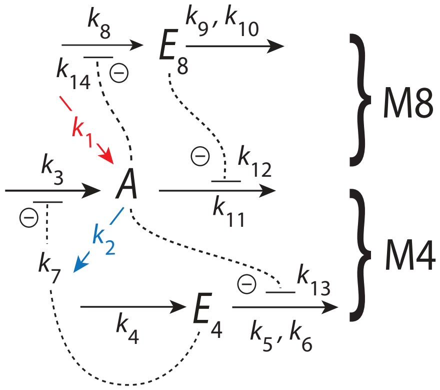


**Fig S13.** The combined M4-M8 controller. Outlined in red indicates the stepwise inflow ($k_1$) and outflow ($k_2$) perturbations applied to the system.

The rate equations are:

$$\dot{A} = \underbrace{k_1}_{\text{perturbation}} - \underbrace{k_2{\cdot}A}_{\text{perturbation}} + \underbrace{k_3\left(\frac{k_7}{k_7{+}E_4}\right)}_{\text{M4 compensatory flux } j_3} - \underbrace{k_{11}{\cdot}A\left(\frac{k_{12}}{k_{12}{+}E_8}\right)}_{\text{M8 compensatory flux } j_{11}} \tag{S29}$$

$$\dot{E}_4 = k_4 - k_5\underbrace{\left(\frac{E_4}{k_6{+}E_4}\right)}_{\approx 1}\left(\frac{k_{13}}{k_{13}{+}A}\right) \tag{S30}$$

$$\dot{E}_8 = k_8\left(\frac{k_{14}}{k_{14}{+}A}\right) - k_9\underbrace{\left(\frac{E_8}{k_{10}{+}E_8}\right)}_{\approx 1} \tag{S31}$$

The setpoints of the two controllers are calculated by setting Eqs S30 and S31 to zero and solving for $A$. This leads to:

$$A_{set}^{M4} = \frac{k_5 k_{13} - k_4 k_{13}}{k_4} \tag{S32}$$

and

$$A_{set}^{M8} = \frac{k_8 k_{14} - k_9 k_{14}}{k_9} \tag{S33}$$

#### $A_{set}^{M4} < A_{set}^{M8}$: Isolated control

When M4 controls $A$, i.e. when the steady state of $A$ is at $A_{set}^{M8}$ $E_8$ shows windup with $\dot{E}_8 > 0$ and M8's compensatory flux $j_{11}$ goes to zero. Similar when M8 controls $A$: in this case we have that $\dot{E}_4 > 0$ and M4's compensatory flux $j_3$ goes to zero. This situation of isolated control is illustrated in Fig S14 with four different $k_1$ and $k_2$ values at four phases.

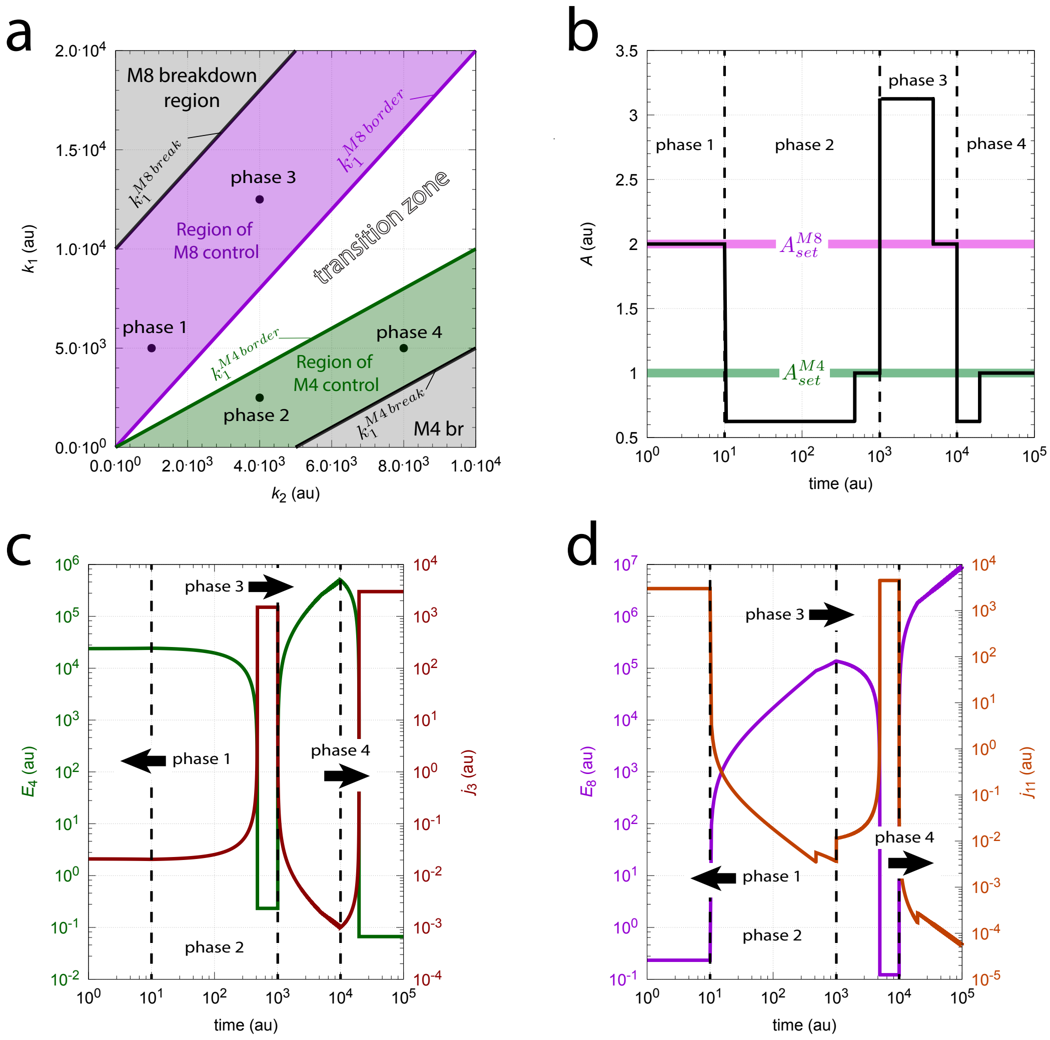


**Fig S14.** Isolated control of the combined M4-M8 controllers with $A_{set}^{M4}$=1.0 and $A_{set}^{M8}$=2.0. Perturbations phase 1: $k_1$=5.0×10$^3$, $k_2$=1.0×10$^3$; phase 2: $k_1$=2.5×10$^3$, $k_2$=4.0×10$^3$; phase 3: $k_1$=1.25×10$^4$, $k_2$=4.0×10$^3$; phase 4: $k_1$=5.0×10$^3$, $k_2$=8.0×10$^3$. Panel a: Perturbation phase diagram with regions of control, the transition zone, and region of M4 and M8 breakdown. The M4 breakdown region is indicated as 'M4 br'. Panel b: $A$ as a function of time during the four phases. Note the logarithmic time axis. Panel c: $E_4$ (left ordinate) and M4's compensatory flux $j_3$ (right ordinate) as a function of time. Panel d: $E_8$ (left ordinate) and M8's compensatory flux $j_{11}$ (right ordinate) as a function of time. Rate constants: $k_3$=5.0×10$^3$, $k_4$=100.0, $k_5$=1100.0, $k_6$=1.0×10$^{-6}$, $k_7$=0.1, $k_8$=2100.0, $k_9$=100.0, $k_{10}$=1.0×10$^{-6}$, $k_{11}$=5.0×10$^3$, $k_{12}$=0.1, $k_{13}$=0.1, and $k_{14}$=0.1. Initial concentrations: $A_0$=2.0, $E_{4,0}$=2.382×10$^4$, $E_{8,0}$=1.011. See `Python code` for details.

Panel a shows the perturbation phase diagram. The region of $A$ control by M8 and M4 are indicated in respectively violet and green, separated by the transition zone. The border of M8 breakdown is reached when M8's compensatory flux has reached its maximum while $A_{ss}$ is still at M8's setpoint, i.e. $j_{11}^{max}$=$k_{11}A_{set}^{M8}$ and M2's compensatory flux $j_3$ is negligible. Inserting these conditions into Eq S29 leads to the borderline between M8 control and its breakdown, i.e.

$$k_1^{M8\,break} = (k_2 + k_{11})A_{set}^{M8} \tag{S34}$$

Likewise, the breakdown of the M4 controller is characterized by having reached its maximum compensatory flux $j_3^{max}$=$k_3$ while M8's compensatory flux is negligible. This leads to the borderline between the M4 control region and the region of M4 breakdown ('M4 br'):

$$k_1^{M4\,break} = k_2 A_{set}^{M4} - k_3 \tag{S35}$$

The borderlines between M4, M8 control and the transition zone ($k_1^{M4\,border}$, $k_1^{M8\,border}$) are characterized by negligible M4, M8 compensatory fluxes, but $A$ having reached $A_{set}^{M4}$ or $A_{set}^{M8}$, which leads to:

$$k_1^{M4\,border} = k_2 A_{set}^{M4} \tag{S36}$$

and

$$k_1^{M8\,border} = k_2 A_{set}^{M8} \tag{S37}$$

Fig S14b shows $A$ as a function of time. When switching from M8 to M2 control the high $E_4$ value during phase 1 takes some time to reach a sufficiently low value (and a sufficiently high $j_3$) such that M2 can take control over $A$. During phase 2 $E_8$ goes into windup even if $A$ has reached $A_{set}^{M4}$. Switching back to M8 control during phase 3 needs even more time to reach low enough $E_8$ values such that M8 can exert control over $A$. Thus, for each switch to the other control regime it takes longer and longer times to get control over $A$. This is reflected by the logarithmic time scale.

Figs S14c and d show respectively $E_4$, $j_3$, and $E_8$, $j_{11}$ as a function of time indicating the mutual windup behaviors in $E_4/E_8$ when M8/M4 take control over $A$.

Fig S15 shows the combined M4-M8 controllers' behaviors when $k_1$ and $k_2$ move along a straight line across the perturbation phase diagram (panel a). At points 1, 2, 3, and 4 (panel a) the line crosses the respective borders $k_1^{M8\,break}$, $k_1^{M8\,border}$, $k_1^{M4\,border}$, and $k_1^{M4\,break}$. Panel b shows $A$ as a function of decreasing $k_1$. Vertical dashed lines correspond to the $k_1$ values of the points 1-4. Violet and green coloring indicate the regions when $A$ is controlled by respectively M8 and M4. Panels c and d show respectively $E_4$ and their compensatory fluxes as a function of $k_1$. The isolated control is clearly seen: When $A$ is controlled by M8 (violet shaded region) M4's compensatory flux $j_3$ is low (panel c), while $j_{11}$ is low when M4 is in control of $A$ (panel d). Note that these low compensatory fluxes are accomplished by windup in either $E_4$ or $E_8$.

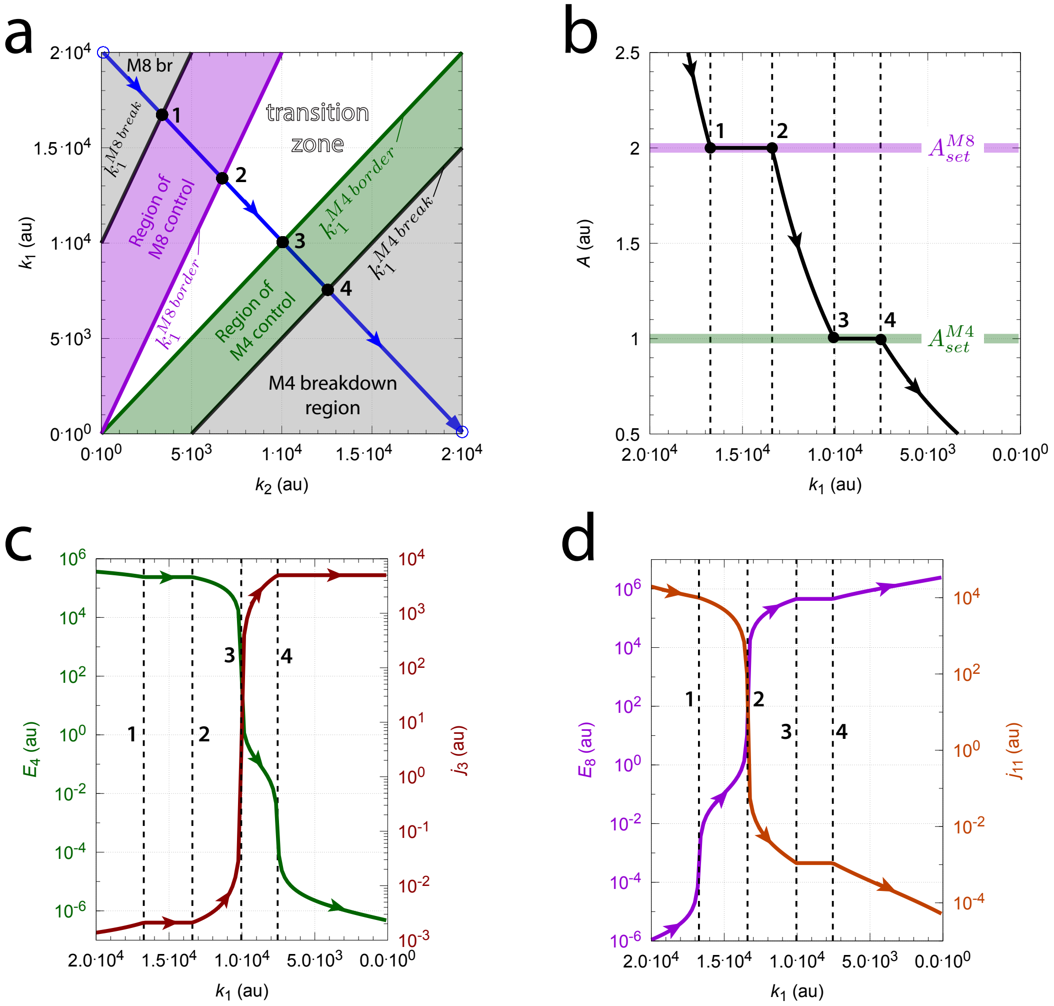


**Fig S15.** Moving across the perturbation phase diagram in the combined M4-M8 controllers when $A_{set}^{M4}$=1.0 and $A_{set}^{M8}$=2.0. Panel a shows the phase diagram. The blue straight line starts at $k_1$=1.0×10$^4$ and $k_2$=1.0×10$^2$ and ends at $k_1$=1.0×10$^2$ and $k_2$=1.0×10$^4$. The line is divided into 100 data points and the steady state of $A$ is calculated after 5000 time units. Panel b shows $A$ as a function of $k_1$. Panels c and d show respectively $E_4$ and $E_8$ (left ordinates) and the values of the compensatory fluxes $j_3$ and $j_{11}$ as a function of $k_1$. Rate constants as in Fig S14. Initial concentrations (all 100 data points): $A_0$=2.0, $E_{4,0}$=1.632×10$^{-6}$, $E_{8,0}$=4.901.

### $A_{set}^{M4}$>$A_{set}^{M8}$: Delegated control

Changing $k_5$ from 1.1×10$^3$ to 2.1×10$^3$ and $k_8$ from 2.1×10$^3$ to 1.1×10$^3$ switches setpoints to $A_{set}^{M4}$=2.0, and $A_{set}^{M8}$=1.0, and to delegated control. Fig S16 shows a calculation with 5 phases. Starting in phase 1 within the M4 breakdown region ('M4 br', panel a) $E_4$ shows windup with $A_{ss}$ being above $A_{set}^{M4}$ (panel b). As $E_4$ windup drives M4's compensatory flux $j_3$ to zero and $E_8$ is low with M8's compensatory flux being approximately $j_{11}$≈$k_{11}A_{ss}$ (as $k_{12}/(k_{12}$+$E_8)$≈1) $A$'s steady state inside the M4 breakdown region can be calculated from Eq S29 to be

$$A_{ss}^{M4\,br} \approx \frac{k_1}{k_2+k_{11}} \tag{S38}$$

which agrees with the numerically calculated value.

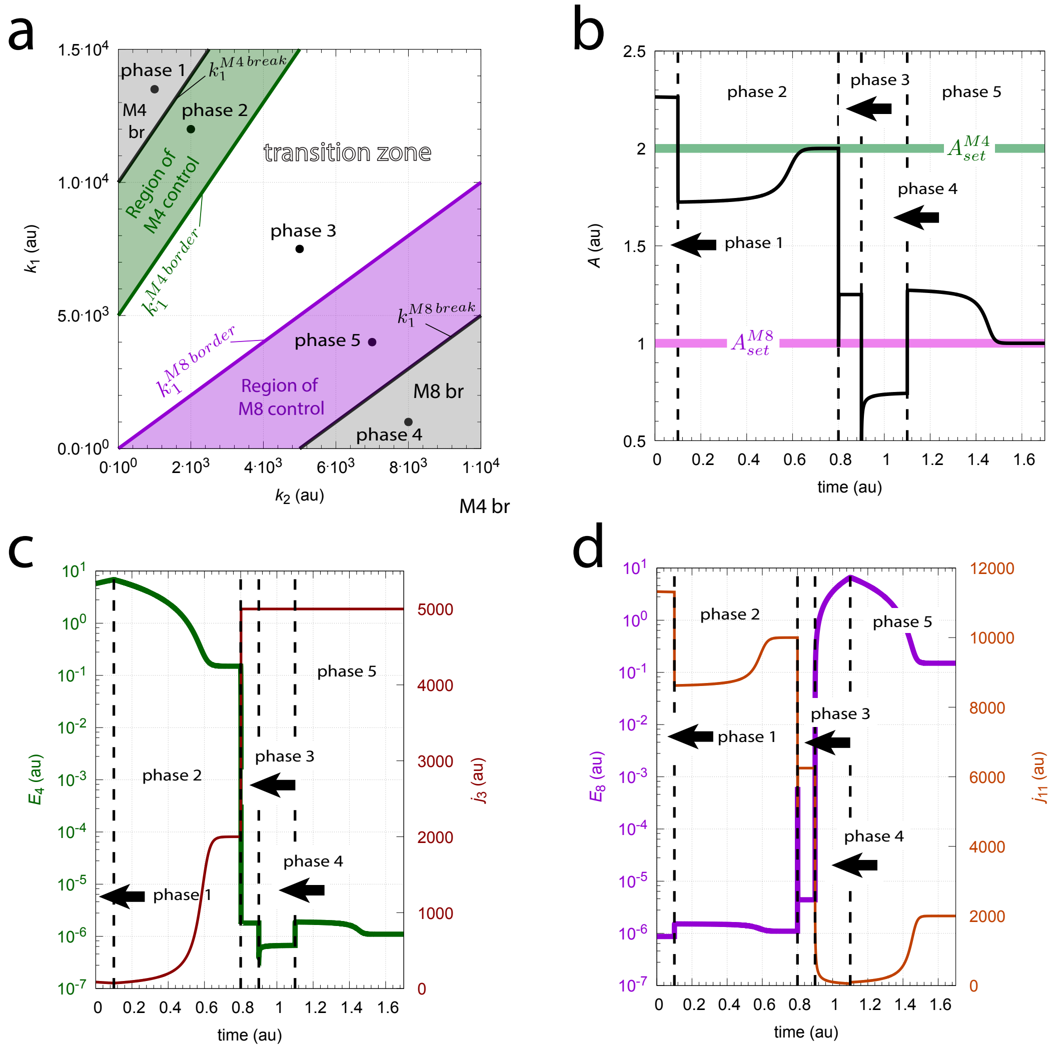


**Fig S16.** Time profiles for delegated control when $A_{set}^{M4}$=2.0, and $A_{set}^{M8}$=1.0. The calculation shows five different perturbations/phases for each region of control, the transition zone, and the two regions of M4 and M8 controller breakdown. Phase 1: $k_1$=1.35×10$^4$, $k_2$=1.0×10$^3$; phase 2: $k_1$=1.2×10$^4$, $k_2$=2.0×10$^3$; phase 3: $k_1$=7.5×10$^3$, $k_2$=5.0×10$^3$; phase 4: $k_1$=1.0×10$^3$, $k_2$=8.0×10$^3$; phase 5: $k_1$=4.0×10$^3$, $k_2$=7.0×10$^3$. Panel a: Perturbation phase diagram. Panel b: $A$ as a function of time. Panel c: $E_4$ (left ordinate) and $j_3$ (right ordinate) as a function of time. Panel d: $E_8$ (left ordinate) and $j_{11}$ (right ordinate) as a function of time. Rate constants: $k_3$=5.0×10$^3$, $k_4$=100.0, $k_5$=2100.0, $k_6$=1.0×10$^{-6}$, $k_7$=0.1, $k_8$=1100.0, $k_9$=100.0, $k_{10}$=1.0×10$^{-6}$, $k_{11}$=5.0×10$^3$, $k_{12}$=0.1, $k_{13}$=0.1, and $k_{14}$=0.1. Initial concentrations: $A_0$=2.26, $E_{4,0}$=5.729, $E_{8,0}$=8.7×10$^{-7}$. See `Python code` for details.

In phase 2 the system is in the M4 controlled region. Since $E_4$ underwent windup in phase 1 the M4 controller needs some time in phase 2 to remove the built-up $E_4$ and move $A$ to $A_{set}^{M4}$ (panels b and c). In phase 3 the system is in the transition zone with both compensatory fluxes fully developed, i.e. $j_3$=$k_3$ and $j_{11}$=$k_{11}A_{ss}$ (as $k_{12}/(k_{12}$+$E_8)\approx 1$). Using Eq S29 the steady state of $A$ inside the transition zone can be calculated to be

$$A_{ss}^{M4\,br} \approx \frac{k_1{+}k_3}{k_2{+}k_{11}} \tag{S39}$$

In phase 4 the system is in the M8 breakdown region (‘M8 br’, panel a). Now $E_8$ undergoes windup and M8’s compensatory flux $j_{11}$ goes to zero (panel d) while $j_3$ is at its maximum value $k_3$ (panel c). This allows to calculate the steady state of $A$ in the M8 breakdown region to be

$$A_{ss}^{M4\,br} \approx \frac{k_1{+}k_3}{k_2} \tag{S40}$$

Finally, in phase 5 the system is in the M8 controlled region. Also here it takes some time to move $E_8$ to low enough values such that $A$ reaches $A_{set}^{M8}$ while $j_3{=}k-3$ is at its maximum value (panels b and d).

The borderline between the regions of M4 breakdown and M4 control, $k_1^{M4\,break}$ (Fig S16a), is characterized that $A_{ss}{=}A_{set}^{M4}$, M8’s compensatory flux is fully developed, i.e. $j_{11}{=}k_{11}A_{set}^{M4}$, and M4’s compensatory flux $j_3$ has gone to zero and can’t any longer compensate the excess of $j_{11}$. Inserting these conditions into the steady state of Eq S29, and writing $k_1$ as a function of $k_2$, leads to the borderline equation:

$$k_1^{M4\,break} = (k_2 + k_{11})A_{set}^{M4} \tag{S41}$$

The border $k_1^{M4\,border}$ between the region of M4 control and the transition zone has fully developed compensatory fluxes of $j_3{=}k_3$ and $j_{11}{=}k_{11}A_{set}^{M4}$ at $A_{ss}{=}A_{set}^{M4}$. This gives the borderline equation:

$$k_1^{M4\,border} = (k_2 + k_{11})A_{set}^{M4} - k_3 \tag{S42}$$

Analogously to $k_1^{M4\,border}$, at the border $k_1^{M8\,border}$ the compensatory fluxes $j_3$ and $j_{11}$ are at their maxima, but $A_{ss}$ is now at $A_{set}^{M8}$. The leads to the $k_1^{M4\,border}$ borderline equation:

$$k_1^{M8\,border} = (k_2 + k_{11})A_{set}^{M8} - k_3 \tag{S43}$$

The border between the region of M8 control and M8 breakdown is characterized that $A_{ss}{=}A_{set}^{M8}$, $j_3{=}k_3$, and $E_8$ shows windup such that $j_{11} \to 0$. This leads to the borderline equation of $k_1^{M8\,break}$

$$k_1^{M8\,break} = k_2A_{set}^{M8} - k_3 \tag{S44}$$

Fig S17 shows a calculation when moving along a straight line inside the perturbation phase diagram (outlined in blue in panel a). The system starts in the M4 breakdown region (‘M4 br’) at $k_1{=}2.0{\times}10^4$ and $k_2{=}1.0{\times}10^2$, moves across the region of M4 control between points 1 and 2, passes the transition zone and moves across the region of M8 control between points 3 and 4. Finally, it moves into the M8 breakdown region and ends at $k_1{=}1.0{\times}10^2$ and $k_2{=}2.0{\times}10^4$. Points 1-4 in panel a are the calculated intersection points between the blue line and the border lines Eqs S41-S44. Fig S17b shows $A$ as a function of $k_1$. Dashed vertical lines represent the $k_1$ values for points 1-4. Green and purple colors indicate the $k_1$ intervals for respectively the region of M4 and M8 control. Panels c and d show $E_4$ and $E_8$, and their respective compensatory fluxes $j_3$ and $j_{11}$ as a function of $k_1$. The delegated control is clearly seen in panels c and d: As M4 control $A$ between points 1 and 2 M8’s compensatory flux $j_{11}$ is high and constant, while during M8 control between points 3 and 4 M4’s compensatory flux is at its maximum.

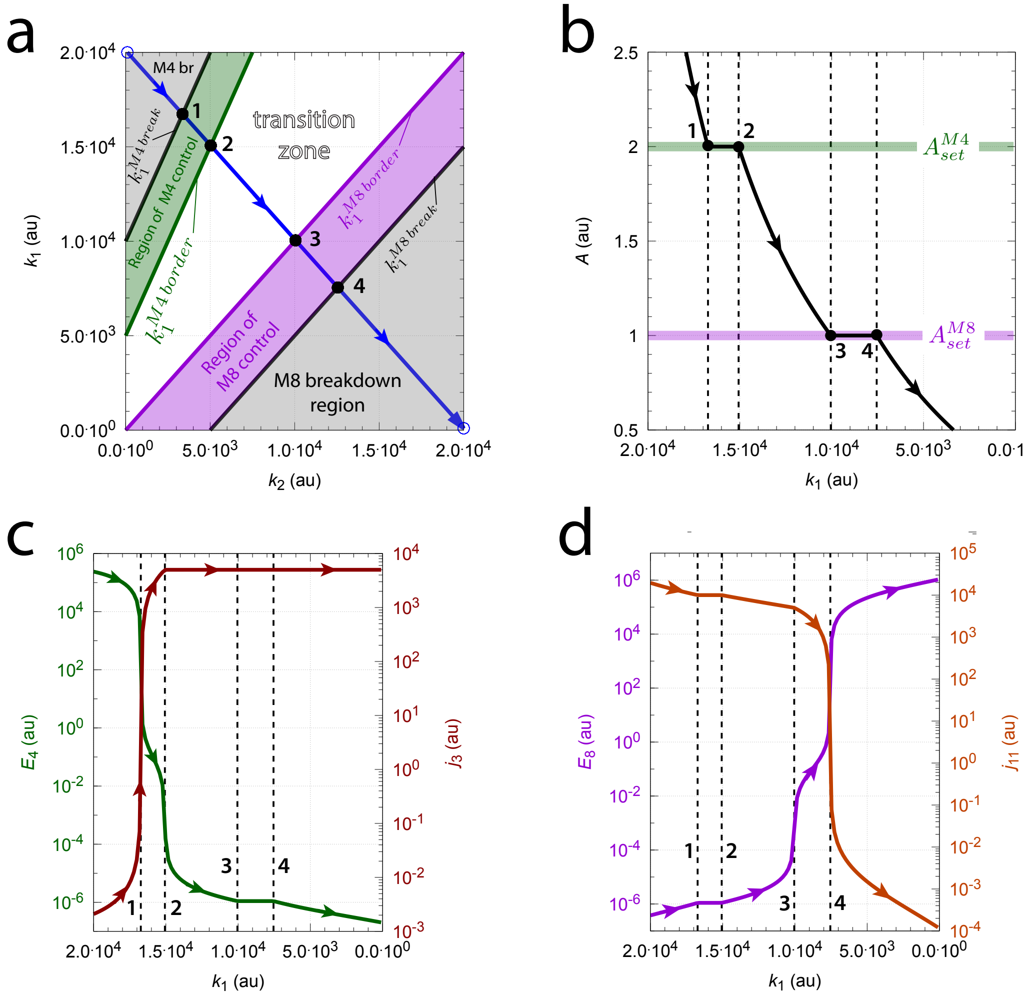


**Fig S17.** Moving across the perturbation phase diagram when $A_{set}^{M4}$=2.0 and $A_{set}^{M8}$=1.0. The blue straight line starts at $k_1$=2.0×10$^4$ and $k_2$=1.0×10$^2$ and ends at $k_1$=1.0×10$^2$ and $k_2$=2.0×10$^4$. The line is divided into 100 data points and the steady state of $A$, $E_4$, $E_8$ and the compensatory fluxes $j_3$ and $j_{11}$ (Eq S29) are determined for each line point after 5000 time units. Panel b shows $A$ as a function of $k_1$. Panels c and d show respectively $E_4$ and $E_8$ (left ordinates) and the values of the compensatory fluxes $j_3$ and $j_{11}$ as a function of $k_1$. The vertical dashed lines 1-4 refer to the $k_1$ values of points 1-4 in panel a. Rate constants and initial concentrations (all 100 calculations) as in Fig S16.
