## Supplementary material for "Flux organizations and control modes in antagonistically combined negative feedback loops": Sup3.pdf

Peter Ruoff

Department of Chemistry, Bioscience, and Environmental Engineering, University of Stavanger, Stavanger, Norway

### Supplementary material

#### Additional examples of metastable control

In the following additional examples of metastable control are given.

##### The combined M2-M6 controller

Fig S1 shows the reaction scheme of the combined M2-M6 controller.

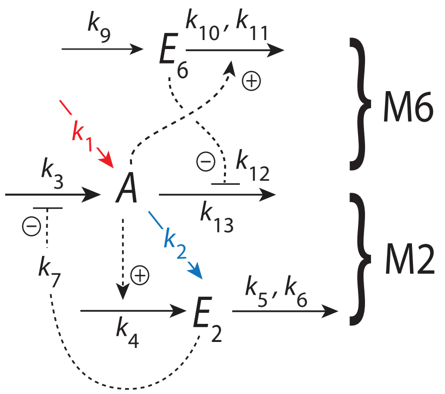


**Fig S1.** The combined M2-M6 controller. Outlined in red and blue indicate the respective stepwise inflow ($k_1$) and outflow ($k_2$) perturbations applied to $A$.

The rate equations are:

$$\dot{A} = \underbrace{k_1}_{\text{perturbation}} - \underbrace{k_2{\cdot}A}_{\text{perturbation}} + \underbrace{k_3\left(\frac{k_7}{k_7{+}E_2}\right)}_{\text{M2 compensatory flux } j_3} - \underbrace{k_{13}{\cdot}A\left(\frac{k_{12}}{k_{12}{+}E_6}\right)}_{\text{M6 compensatory flux } j_{13}} \tag{S1}$$

$$\dot{E}_2 = k_4{\cdot}A - k_5{\cdot}\left(\frac{E_2}{k_6{+}E_2}\right) \tag{S2}$$

$$\dot{E}_6 = k_9 - k_{10}{\cdot}A{\cdot}\left(\frac{E_6}{k_{11}{+}E_6}\right) \tag{S3}$$

Using zero-order kinetics with respect of $E_2$ and $E_6$ in the degradations of $E_2$ and $E_2$, i.e. setting $k_6{=}k_{11}{=}1\times10^{-6}$ such that $E_2/(k_6{+}E_2){\approx}1$ and $E_6/(k_{11}{+}E_6){\approx}1$, the set-points for $A$ of M2 and M6 are

$$A_{set}^{M2} = \frac{k_5}{k_4} \quad \text{(S4)}$$

$$A_{set}^{M6} = \frac{k_9}{k_{10}} \quad \text{(S5)}$$

For more information on the M2-M6 system see Tables S6 and S22.

**Metastability in the combined M2-M6 controllers**

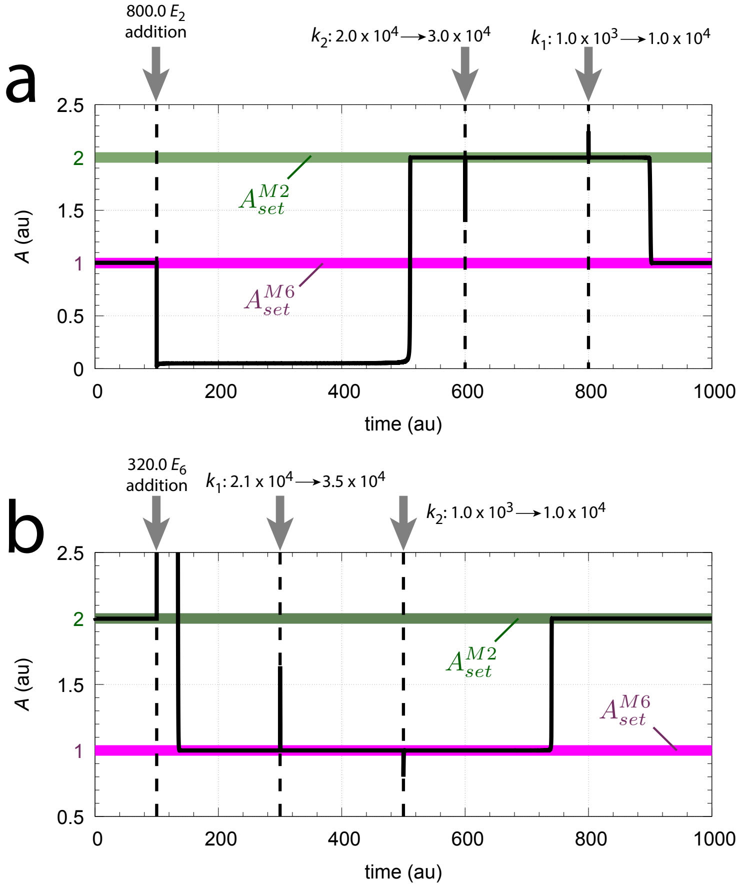


**Fig S2.** Metastability in the combined M2-M6 controller with $A_{set}^{M2}$=2.0 and $A_{set}^{M6}$=1.0. Panel a: Induced metastability of the M2 controller by adding 800.0 concentration units of $E_2$ to the system at t=100.0. The metastable M2 state appears at around 500 time units and is stable against $k_2$ and $k_1$ steps as indicated in the figure. Rate constants: $k_1$=1.0×10$^3$ (t=0), $k_2$=2.0×10$^4$ (t=0), $k_3$=6.0×10$^4$, $k_4$=1.0, $k_5$=2.0, $k_6$=1.0×10$^{-6}$, $k_7$=$k_8$=0.1, $k_9$=$k_{10}$=0.5, $k_{11}$=1.0×10$^{-6}$, $k_{12}$=0.1, and $k_{13}$=4.1×10$^4$. Initial concentrations: $A_0$=1.0, $E_{2,0}$=1.0×10$^{-6}$, $E_{6,0}$=8.183×10$^{-2}$. Panel b: Induced M6 metastability by the addition of 320.0 concentration units of $E_6$ at t=100.0. The M6 state is stable against applied $k_1$ and $k_2$ step perturbations. Rate constants: $k_1$=2.1×10$^4$ (t=0), $k_2$=1.0×10$^3$ (t=0), $k_{13}$=3.5×10$^4$. Other rate constants as in panel a. Initial concentrations: $A_0$=2.0, $E_{2,0}$=7.864×10$^{-3}$, $E_{6,0}$=1.0×10$^{-6}$.

Metastability of M2 or M6 can be observed by the addition of either $E_2$ or $E_6$ to the combined M2-M6 system. Fig S2a shows the effect when 800 $E_2$ concentration units are added to the system when M6 has originally control on $A$ under delegated control. Like for the metastable M2M7 controller (see Fig26, panels a1 and a2) the addition of $E_2$ causes both compensatory fluxes to disappear and $A$ is initially determined by the ratio $k_1/k_2$. When both compensatory fluxes $j_3$ and $j_{13}$ reappear after a relatively long transit ($E_2$ is slowly removed) M2 dominates and keeps $A$ temporarily at its set-point. Step perturbations in $k_2$ and $k_1$ show that M2 is (temporarily) stable. When $E_2$ has finally been assimilated and $j_3$ has reached its maximum level (=$k_3$) M6 regains delegated control over $A$.

Panel b shows the induced metastable $A$ control by M6 when 320 units of $E_6$ are added under M2 control. The addition of $E_6$ causes a rapid increase in $A$ and an initial block of the two compensatory fluxes. As $E_6$ slowly decreases the two compensatory fluxes $j_3$ and $j_{13}$ increase correspondingly, but the M6's compensatory flux $j_{13}$ is the dominating one such that M6 controls $A$. Small step-wise perturbations show that the M6 state is temporarily stable. When the excess of $E_6$ is depleted the system returns to the M2-controlled state at $A_{ss}$=$A_{set}^{M2}$=2.0 with $j_{13}$ at its (delegated) maximum, i.e. $j_{13}$=$k_{11}A_{set}^{M2}$.

### Metastability in the combined M2-M7 controller

Fig S3 illustrates induced metastability by using the combined M2-M7 controller when $k_1$=1.0 and $k_2$=2.0. Panel a1 shows the induced metastability of M2 at $A_{set}^{M2} = 1.1$ when 800 units of $E_2$ are added to the system controlled by M7 at $A_{set}^{M7} = 1.0$. The $E_2$ addition is indicated by arrow 1. The system responds by switching from M7 control to a temporary state where both M2 and M7 have negligible compensatory fluxes and $A$ is given by the ratio between $k_1$ and $k_2$, i.e. $A_{ss} = k_1/k_2$=0.5 (indicated by arrow 2). The cause of the lack of compensatory fluxes is due to the effective inhibition of M2's compensatory flux $j_3$, which leads to a low $A$ level and subsequently to low $E_7$ and a low compensatory flux $j_{11}$. When both compensatory fluxes $j_3$ and $j_{11}$ start to grow again as $E_2$ decreases (with the dominance of $j_3$), $A$ switches to temporary M2 control at $A_{set}^{M2} = 1.1$ and stays there until M2's compensatory flux $j_3$ has reached its maximum value ($k_3$) and delegated control by M7 is reestablished. Small step perturbations in $k_1$ (arrow 3) and $k_2$ (arrow 4) indicate that the metastable M2 state is locally stable. Note, however, that larger step perturbations, for example in $k_1$, will move the control of $A$ towards M2 control or into M2's breakdown zone.

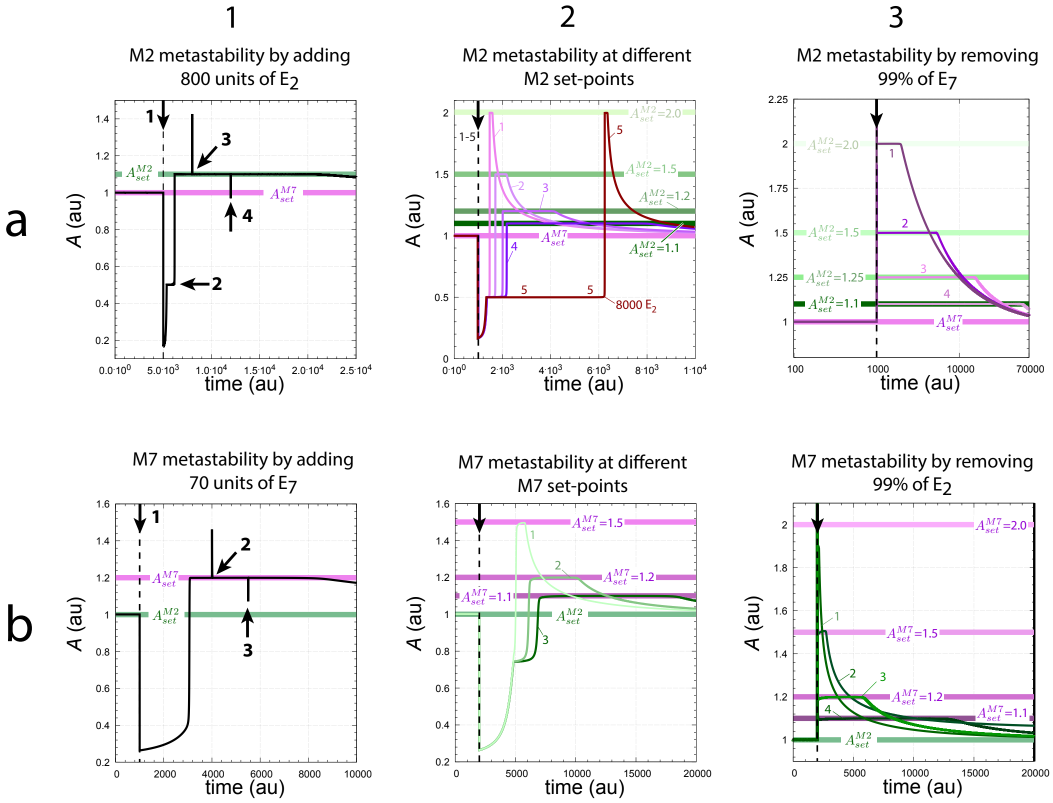


**Fig S3.** Induced M2 and M7 metastable states in the M2M7 controller by adding/removing $E_2$ or $E_7$. Panel a1: Induced M2 metastability by adding 800 $E_2$ units at t=5000 (arrow 1). Rate parameters: $k_1$=1000 (inflow perturbation), $k_2$=2000 (outflow perturbation), $k_3$=$k_{11}$=5.0×$10^3$, $k_4$=1.0, $k_5$=1.1, $k_6$=$k_{10}$=1.0×$10^{-6}$, $k_7$=$k_{12}$=$k_{13}$=0.1, $k_8$=4.5455×$10^{-4}$, and $k_9$=5.0×$10^{-3}$. Initial concentrations: $A_0$=1.0, $E_{2,0}$=1.0×$10^{-6}$, and $E_{7,0}$=0.4. At time t=8000 (arrow 3) $k_1$ is changed to 3000 while $k_2$ remains unchanged. At t=1.2×$10^4$ $k_2$ is changed to 3000. Around t=2×$10^4$ $A$ begins to decrease and moving towards $A_{set}^{M7}$. Panel a2: Same parameters and initial concentrations as in panel a1, except that $E_2$ addition occurs at t=1000 and the $A_{set}^{M2}$'s are changed via altered $k_5$ values. Curve 1, $k5$=2.0; curve 2, $k5$=1.5; curve 3, $k5$=1.2; curve 4, $k5$=1.1. Curve 5, as curve 1, but 8000 $E_2$ units are added. Panel a3: Initial conditions similar to panel a2, but instead of $E_2$ addition 99% of $E_7$ is removed. Changed rate constants: $k_4$=0.1, $k_8$=4.5455×$10^{-5}$, and $k_9$=5.0×$10^{-4}$. The $A_{set}^{M2}$ values are: Curve 1, $A_{set}^{M2}$=2.0 with $k5$=0.2; curve 2, $A_{set}^{M2}$=1.5 with $k5$=0.15; curve 3, $A_{set}^{M2}$=1.25 with $k5$=0.125, and curve 4 with $A_{set}^{M2}$=1.1 with $k5$=0.1. Note the logarithmic scale of the time axis. Panel b1: Induced M7 metastability by adding 70 $E_7$ units at t=1000 (arrow 1). Rate parameters: $k_1$=2000 (inflow perturbation), $k_2$=2500 (outflow perturbation), $k_3$=5.0×$10^3$, $k_4$=$k_5$=5.0×$10^{-4}$, $k_6$=$k_{10}$=1.0×$10^{-6}$, $k_7$=$k_{12}$=$k_{13}$=0.1, $k_8$=1.53846×$10^{-2}$, $k_9$=0.2, $k_{11}$=7.0×$10^3$. Initial concentrations: $A_0$=1.0, $E_{2,0}$=8.9343×$10^{-1}$, and $E_{7,0}$=1.11329×$10^{-5}$. At time t=4000 $k_1$ is changed to 3000 (arrow 2), while $k_2$ is changed to 3000 at t=5500. At about t=8000 $A$ starts to decrease again towards $A_{set}^{M2}$=1.0. Panel b2: Conditions as in panel b1, but $A_{set}^{M7}$ is changed via changed $k_8$ values. Curve 1, $A_{set}^{M7}$=1.5 by $k_8$=9.375×$10^{-3}$; curve 2, $A_{set}^{M7}$=1.2 by $k_8$=1.15385×$10^{-2}$; curve 3, $A_{set}^{M7}$=1.1 by $k_8$=1.25×$10^{-2}$. Panel b3: Parameters as panel b2, but M7 metastability is induced by removing 99% of $E_2$. Curve 1, $A_{set}^{M7}$=2.0 by $k_8$=7.14286×$10^{-3}$ ($A$ goes immediately back to $A_{set}^{M2}$=1.0); curve 2, $A_{set}^{M7}$=1.5 by $k_8$=9.375×$10^{-3}$; curve 3, $A_{set}^{M7}$=1.2 by $k_8$=1.15385×$10^{-2}$; curve 4, $A_{set}^{M7}$=1.1 by $k_8$=1.25×$10^{-2}$.

Panel a2 shows that the residence time in the M2 metastable state increases as M2's set-point decreases. Lines 1-4 show the same conditions as in panel a1, but the set-point of M2 is respectively 2.0, 1.5, 1.2, and 1.1. Line 5 shows the system's response when 8000 $E_2$ units are added instead of 800 and when $A_{set}^{M2}$=2.0. The system stays now longer at $A_{ss} = k_1/k_2$=0.5, but the residence time at the M2 metastable state remains unaltered, as for line 1.

Panel a3 shows that M2 metastability can be initiated by removing $E_7$. Removing most of $E_7$ (here 99%, indicated by the vertical arrow) leads to a negligible M7 compensatory flux $j_{11}$, which in turn leads to M2 as the only temporary controller of $A$. Lines 1-4 show the resulting $A$ profiles when $A_{set}^{M7}$=1.0 (horizontal violet line) and $A_{set}^{M2}$ is changed respectively to 2.0, 1.5, 1.25, and 1.1 (horizontal green lines). Also here the residence time at the metastable M2 state increases with decreasing $A_{set}^{M2}$.

Panels b1-b2 show M7 metastable states when 70 units of $E_7$ are added to a state with $A$ stable at $A_{set}^{M2}$. With decreasing $A_{set}^{M7}$ the residence time at metastable M7 increases. Panel b3 shows that M7 metastability can also be induced by removal of $E_2$. Curves 1-4 show that M7's metastable residence time of $A$ increases as $A_{set}^{M7}$ decreases. Note that $A$ in curve 1 does not reach $A_{set}^{M7}$=2.0.

### Windup-induced metastability in the combined M2-M8 controller

Fig S4 shows the reaction scheme of the combined M2-M8 controller.

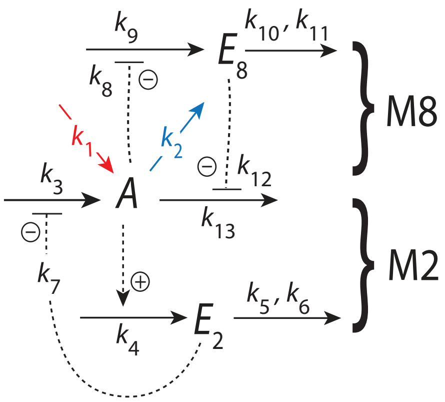


**Fig S4.** The combined M2-M8 controller. Red and blue outlines indicate the respective stepwise inflow ($k_1$) and outflow ($k_2$) perturbations applied on $A$.

The rate equations are:

$$\dot{A} = \underbrace{k_1}_{\text{perturbation}} - \underbrace{k_2 \cdot A}_{\text{perturbation}} + \underbrace{k_3 \left( \frac{k_7}{k_7 + E_2} \right)}_{\text{M2 compensatory flux } j_3} - \underbrace{k_{13} \cdot A \left( \frac{k_{12}}{k_{12} + E_8} \right)}_{\text{M8 compensatory flux } j_{13}} \tag{S6}$$

$$\dot{E}_2 = k_4 \cdot A - \frac{k_5 \cdot E_2}{k_6 + E_2} \tag{S7}$$

$$\dot{E}_8 = k_9 \left( \frac{k_8}{k_8 + A} \right) - \frac{k_{10} \cdot E_8}{k_{11} + E_8} \tag{S8}$$

Assuming zero-order conditions in the removal of $E_2$ and $E_8$, i.e. $E_2/(k_6{+}E_2){\cong}1$ and $E_8/(k_{11}{+}E_8){\cong}1$ (this can practically be achieved by setting $k_6$ and $k_{11}$ both to $1{\times}10^{-6}$), the set-points of the individual M2 and M8 controllers can be determined by the steady state conditions $\dot{E}_2{=}\dot{E}_8{=}0$ and solving for $A$ from Eqs S7 and S8. This leads to two set-points, one for M2 and another for M8:

$$A_{set}^{M2} = \frac{k_5}{k_4} \tag{S9}$$

$$A_{set}^{M8} = \frac{k_8 k_9 - k_8 k_{10}}{k_{10}} \tag{S10}$$

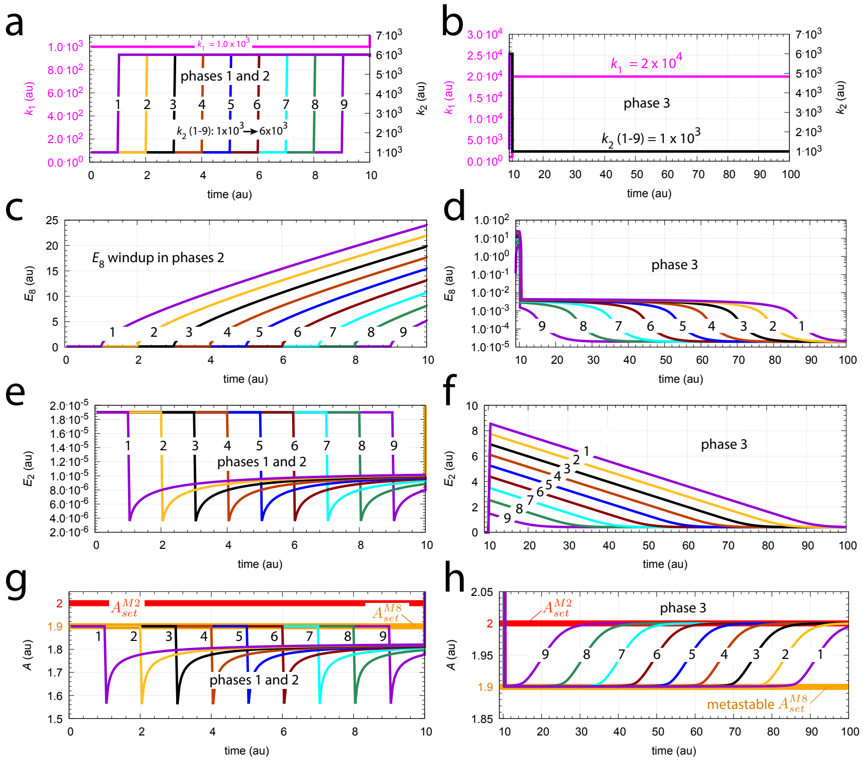

**Fig S5.** Metastable M8 set-point by increased $E_8$ concentrations through windup. Panel a: $k_1$ and $k_2$ values as a function of time. Numbers 1-9 indicate the stepwise changes of $k_2$ from $1\times10^3$ to $6\times10^3$ at times 1.0, 2.0,..., and 9.0, which define the length of phase 1. The lengths of phase 2 when $k_2$=$6\times10^3$ are 9.0, 8.0,..., and 1.0 such that the total lengths of phases 1 and 2 are 10.0. $k_1$ stays constant during phases 1 and 2 at $1\times10^3$. Panel b shows the change to constant $k_1$=$2\times10^4$ and to constant $k_2$=$1\times10^3$ in phase 3. Panel c: Windup of $E_8$ during phases 2. Panel d: $E_8$ concentrations for situations 1-9 as a function of time during phase 3. Note how the relaxation times back to M2 control depends upon the length of phase 1. Panel e: $E_2$ concentrations as a function of time during phases 1 and 2. Panel f: $E_2$ concentrations in phase 3 as a function of time. Panel g: Concentration of $A$ for phases 1 and 2. Due to windup in $E_8$ and breakdown of M8 the concentration of $A$ in phases 2 falls below $A_{set}^{M8}$. Panel h: Concentration of $A$ during phase 3. $A$ stays at the metastable M8's set-point $A_{set}^{M8}$ once windup has finished and $E_8$ decreases. When $E_8$ has reached its minimum value $A$ is controlled by M2. Rate constants: $k_3$=$1.0\times10^4$, $k_4$=1.0, $k_5$=2.0, $k_6$=$1.0\times10^{-6}$, $k_7$=$k_8$=0.1, $k_9$=$1.0\times10^3$, $k_{10}$=50.0, $k_{11}$=$1.0\times10^{-6}$, $k_{12}$=0.1, and $k_{13}$=$1.0\times10^4$. Initial concentrations: $A_0$=1.9, $E_{2,0}$=$1.9\times10^{-5}$, $E_{8,0}$=$1.09\times10^{-1}$.

Here an example of $E_8$ windup-induced metastable control of $A$ by M8 for delegated control ($A_{set}^{M2}$=2.0 and $A_{set}^{M8}$=1.9). The system starts out with stable control of $A$ by M8 when $k_1$=$k_2$=$1.0\times10^3$. This defines phase 1 of the system. Then $k_2$ is increased at different times to $k_2$=$6.0\times10^3$ (curves 1-9 in Fig S5a). This increase of $k_2$ pushes the system into the M8 breakdown region (Table S8) and causes windup in $E_8$ ($\dot{E}_8 > 0$). The system stays then up to 10 time units in the $E_8$ windup state, which defines phase 2 (panel c). After 10 time units the $E_8$ windup is stopped and the system is moved into the M2 controlled region by increasing $k_1$ to $2.0\times10^4$ and reducing $k_2$ back to $1.0\times10^3$.

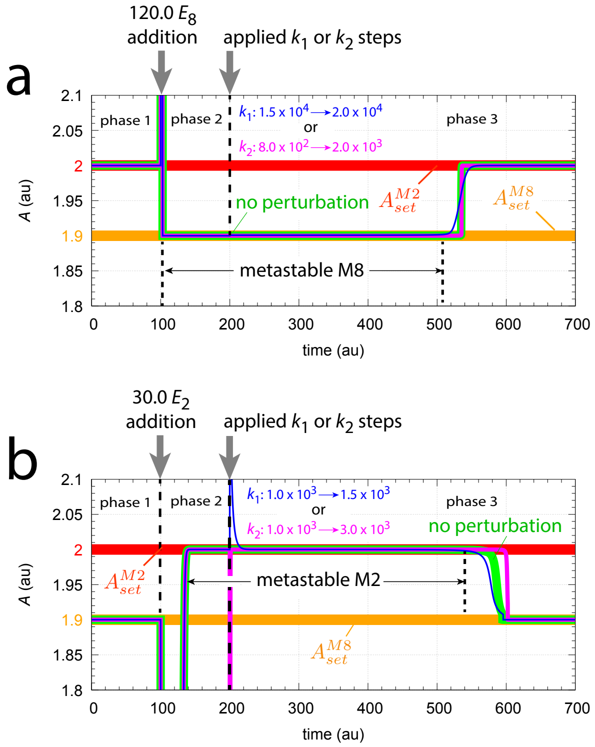


**Fig S6.** Metastable states defend their temporary set-points. Panel a: The control delegated to M2 with set-point $A_{set}^{M2}$ in phase 1 is temporarily changed into a metastable M8 controlled state with set-point $A_{set}^{M8}$ by the addition of 120 concentration units of $E_8$ at $t$=100.0 (indicated by the grey vertical arrow). Steps of $k_1$: $1.5{\times}10^4 \rightarrow 2.0{\times}10^4$ or $k_2$: $8.0{\times}10^2 \rightarrow 2.0{\times}10^3$ applied at $t$=200.0 (indicated by respectively blue and magenta curves) do not alter the regulated $A$ by the metastable M8 controller. At approximately 540 time units the metastable state disappears as $E_8$ returns to its minimum value of $2.0{\times}10^{-5}$ and control is delegated again to M2. Rate constants: $k_1$=$1.5{\times}10^4$ ($t$=0), $k_2$=$8.0{\times}10^2$ ($t$=0), $k_3$=$1.0{\times}10^4$, $k_4$=1.0, $k_5$=2.0, $k_6$=$1.0{\times}10^{-6}$, $k_7$=$k_8$=0.1, $k_9$=$1.0{\times}10^3$, $k_{10}$=50.0, $k_{11}$=$1.0{\times}10^{-6}$, $k_{12}$=0.1, and $k_{13}$=$1.0{\times}10^4$. Initial concentrations: $A_0$=2.0, $E_{2,0}$=$5.162{\times}10^{-2}$, $E_{8,0}$=$2.0{\times}10^{-5}$. Panel b: Phase 1 shows the control of $A$ by M8 at $A_{set}^{M8}$=1.9. At time $t$=100.0 $E_2$ is added (30 concentration units) and the system goes into a metastable M2 state. The system returns to the M8 controlled state once $E_2$ has returned to its minimum value of $1.9{\times}10^{-5}$. The green curve shows the system's behavior in absence of a perturbation, while blue and magenta curves show $A$ as a function of time when respectively $k_1$: $1.0{\times}10^3 \rightarrow 1.5{\times}10^3$ or $k_2$: $1.0{\times}10^3 \rightarrow 3.0{\times}10^3$ steps are applied at $t$=200.0. $k_1$=$k_2$=$1.0{\times}10^3$ ($t$=0); other rate constants as for panel a. Initial concentrations: $A_0$=1.9, $E_{2,0}$=$1.9{\times}10^{-5}$, $E_{8,0}$=$2.586{\times}10^{-1}$.

The system is then followed for 100 time units (which defines phase 3, panels b, d, f, and h). While this $k_1$, $k_2$ change should move $A$ to M2 control, $A$ stays temporarily at $A_{set}^{M8}$=1.9 as long as $E_2$ and $E_8$ stay high (panels d and f). When $E_8$ has sufficiently decreased $A$ is controlled by M2, as expected (panel h). As M2 is in control of $A$ and $A_{ss}$>$A_{set}^{M8}$ the rate of forming $E_8$ is less than its degradation (due to the stronger inhibition of the rate forming $E_8$) $E_8$ shows negative windup ($\dot{E}_8$<0) and moves to a minimum value of $E_8^{min}$=$2{\times}10^{-5}$ (panel d). $E_8^{min}$ can be calculated from Eq S8 by setting $A$=$A_{set}^{M2}$ and solving for $E_8$, which leads to

$$E_8^{min} = \frac{k_8 k_9 k_{11}}{k_8 k_{10} + k_{10} A_{set}^{M2} - k_8 k_9} \tag{S11}$$

Fig S6 shows that the metastable M2 and M8 controllers are able to defend their set-points. In phase 1 of Fig S6a delegated M2 control keeps $A$ at $A_{set}^{M2}$=2.0 ($A_{set}^{M2}$ is indicated by the thick red line). At time $t$=100.0 (beginning of phase 2) 120.0 concentration units of $E_8$ are added which moves $A$'s steady state temporarily to $A_{set}^{M8}$ (indicated by the thick orange line). This M8 state is metastable and $E_8$ will decrease similarly as shown in Fig S5d. The spontaneous return of $A$ to $A_{set}^{M2}$ with no applied perturbation is shown by the thick green line. On top of the green line two $A$ curves are added. In one curve, outlined in blue, $k_1$ undergoes a stepwise change from $1.5{\times}10^4$ to $2.0{\times}10^4$ at t=200.0 (which is the beginning of phase 3). The other curve, outlined in magenta, shows the effect of a stepwise $k_2$ change from $8.0{\times}10^2$ to $2.0{\times}10^3$ (also applied at t=200.0).

In Fig S6b $k_1$ and $k_2$ steps are applied when M2 is metastable. Here the system starts out with $A$ at $A_{set}^{M8}$. At t=100.0 30 concentration units of $E_2$ are added which moves the system into a metastable M2 state with $A$ at $A_{set}^{M2}$. As in panel a, the green thick line shows $A$ when the system returns spontaneously back to $A_{set}^{M8}$ without any applied perturbation. Stepwise perturbations are applied at t=200.0. The blue curve shows the response of a $k_1$ step from $1.0{\times}10^3$ to $1.5{\times}10^3$, while the magenta line shows the response of a $k_2$: $1.0{\times}10^3 \rightarrow 3.0{\times}10^3$ step.
