## Supplementary figures and images for "Flux organizations and control modes in antagonistically combined negative feedback loops"

### A-time-plot_fig9.pdf

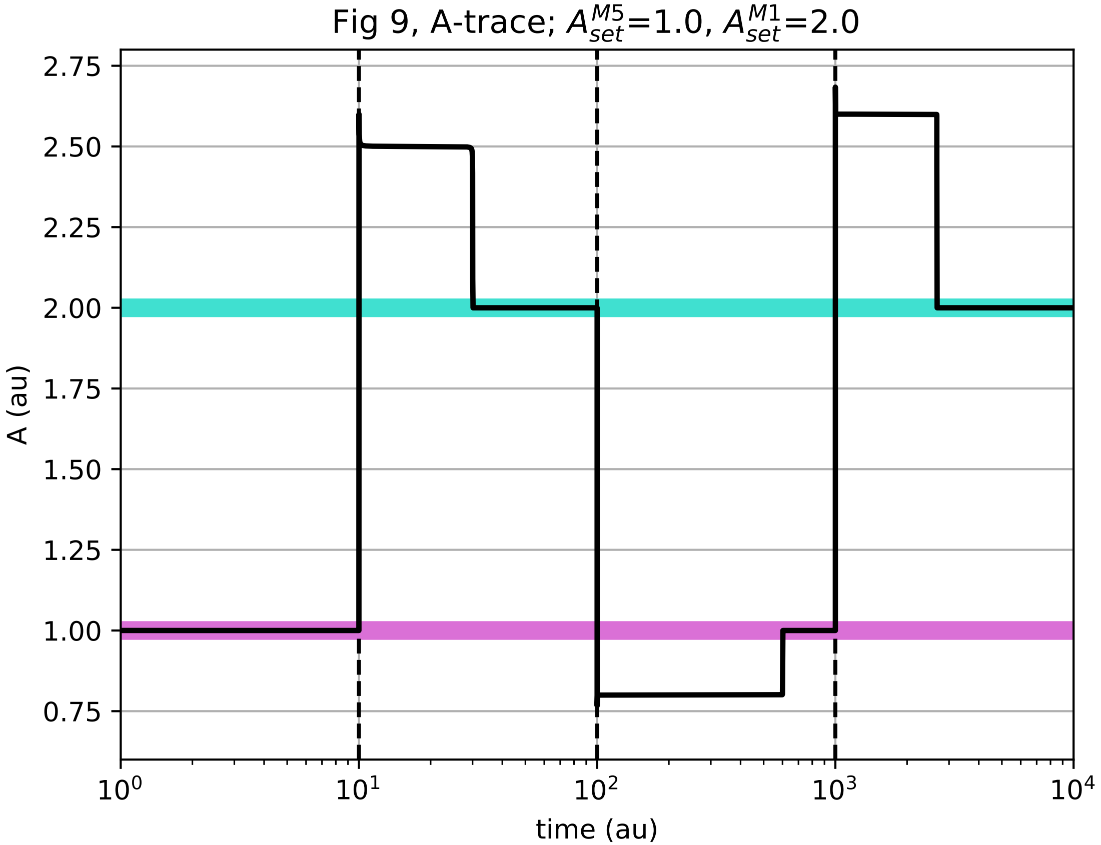

### A-time-plot_fig21.pdf

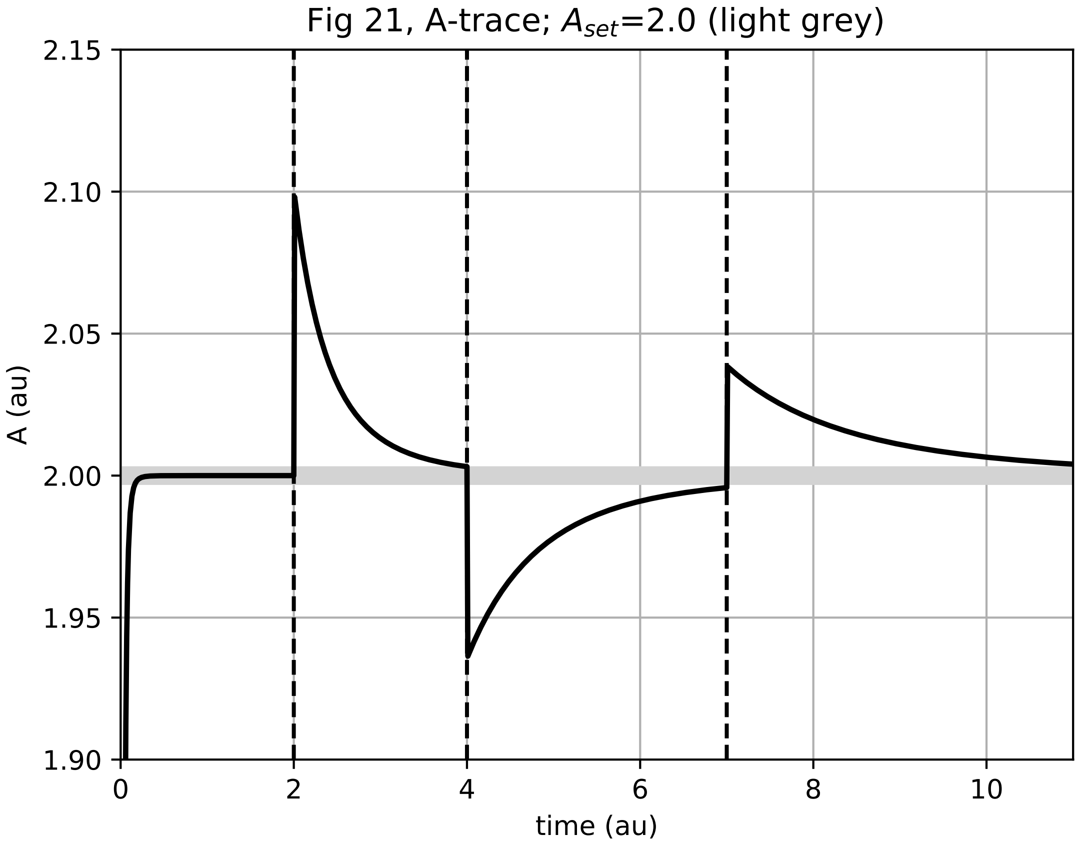

### A-time-plot_fig26a.pdf

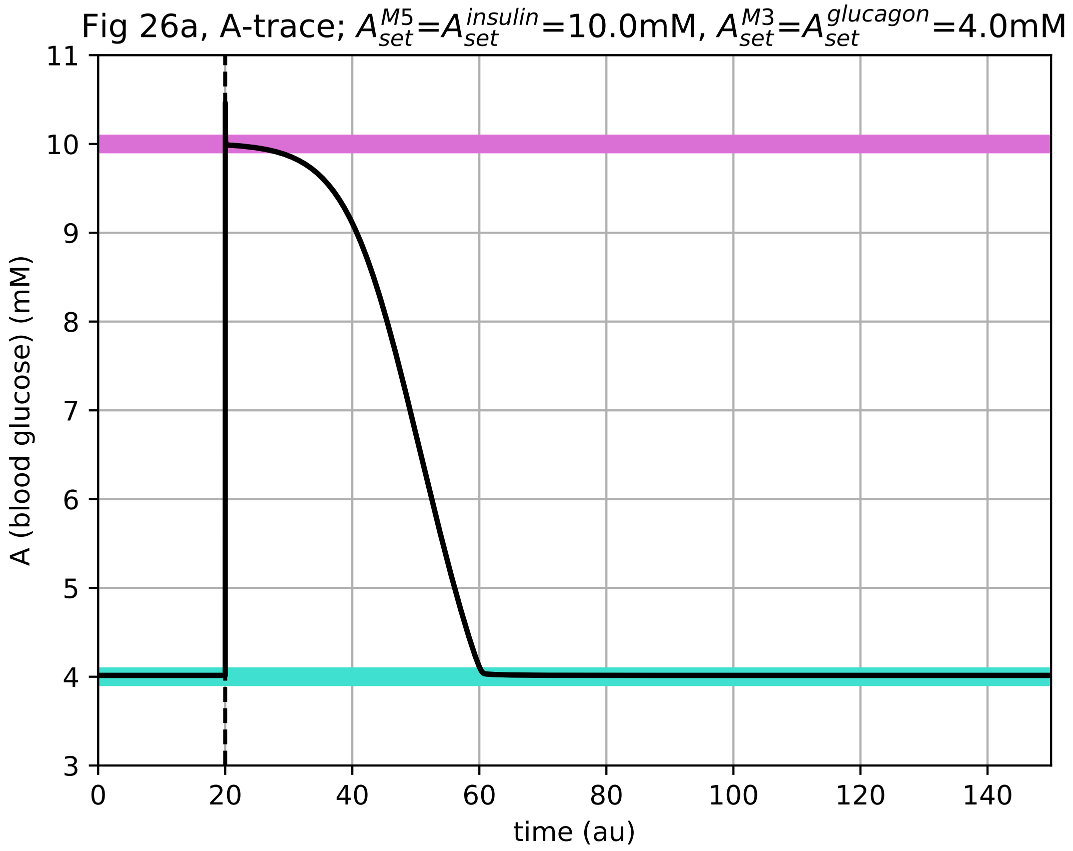

### A-time-plot_fig26b.pdf

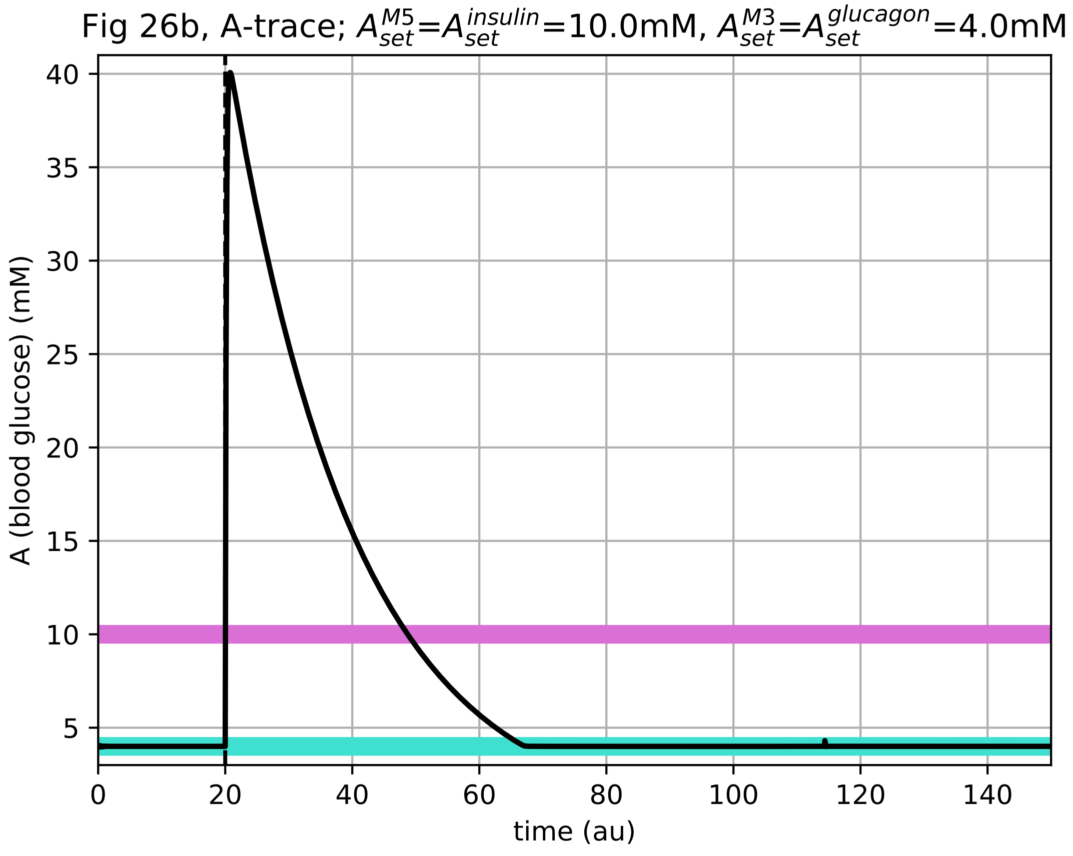

### A-time-plot_fig26c.pdf

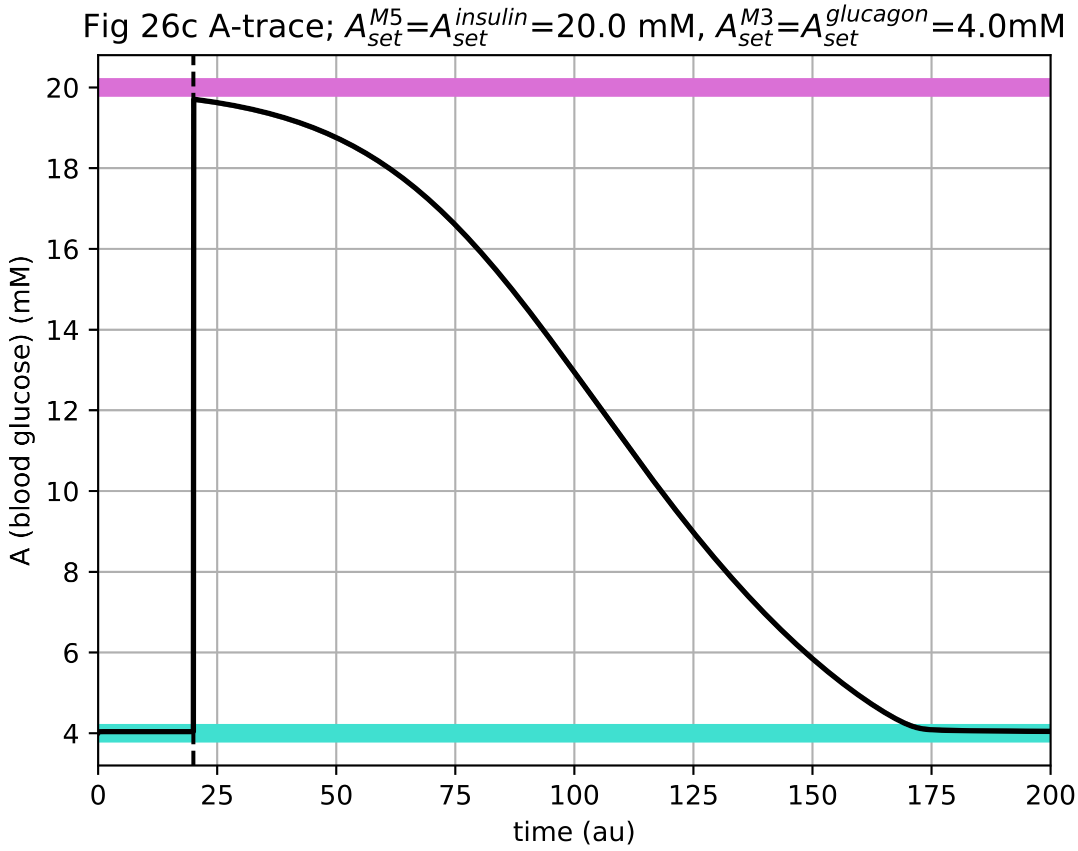

### A-time-plot_fig26d.pdf

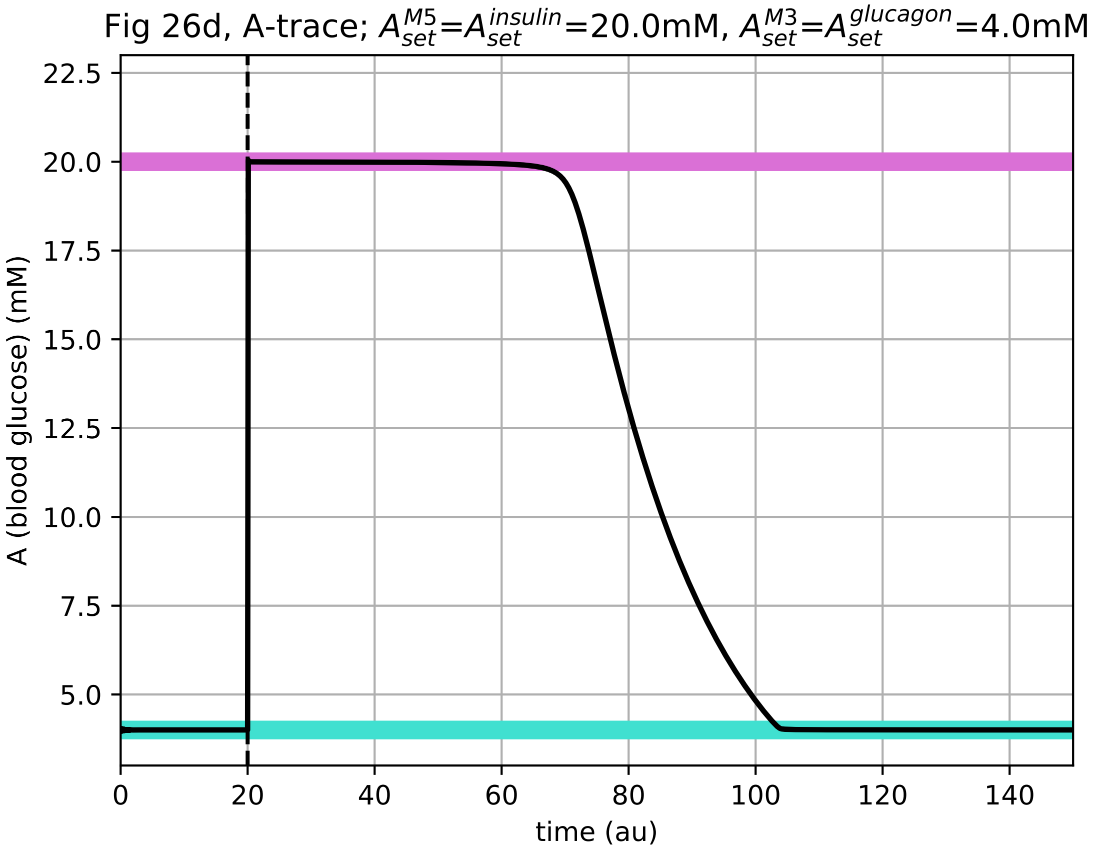

### A-time-plot_figS3.pdf

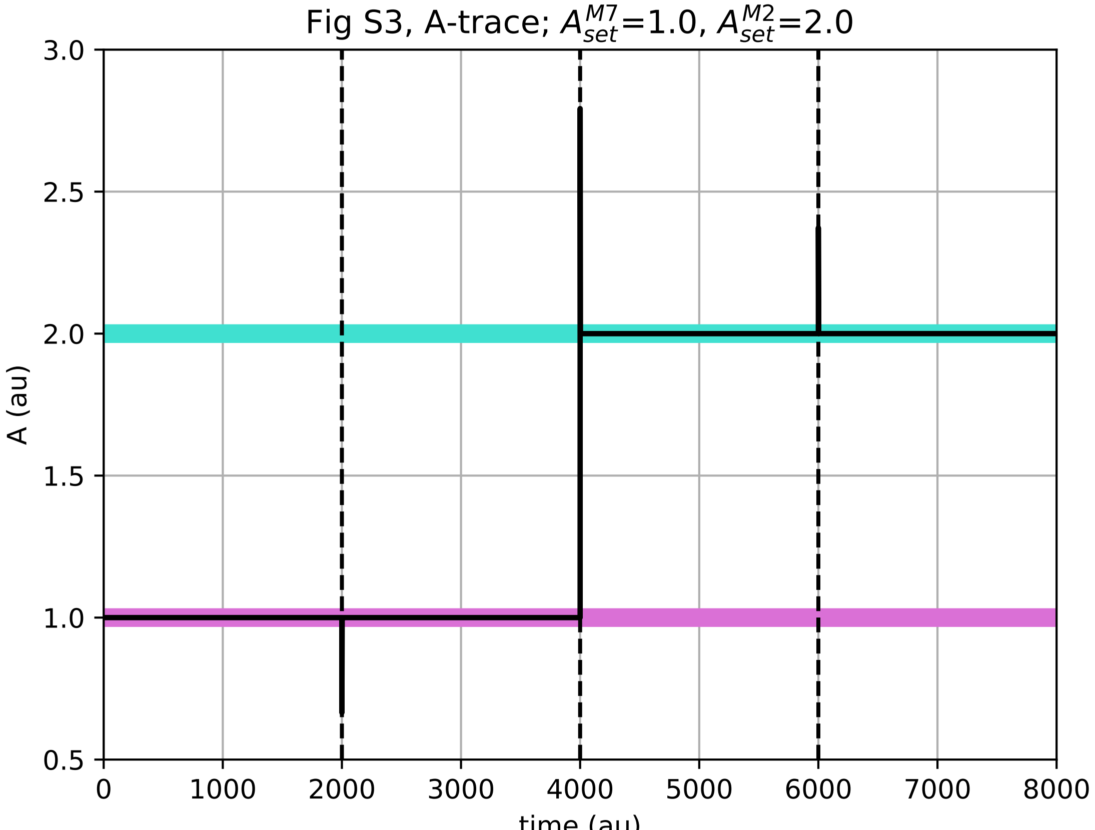

### A-time-plot_figS6.pdf

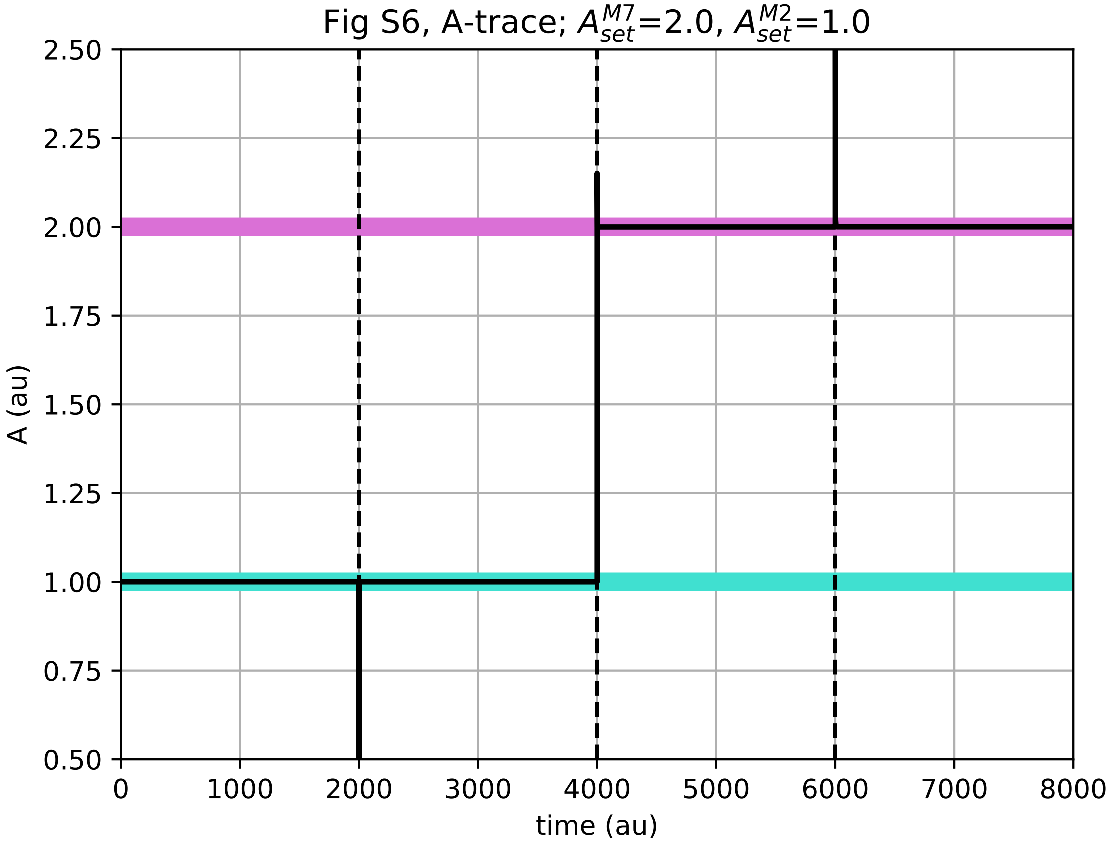

### A-time-plot_figS9.pdf

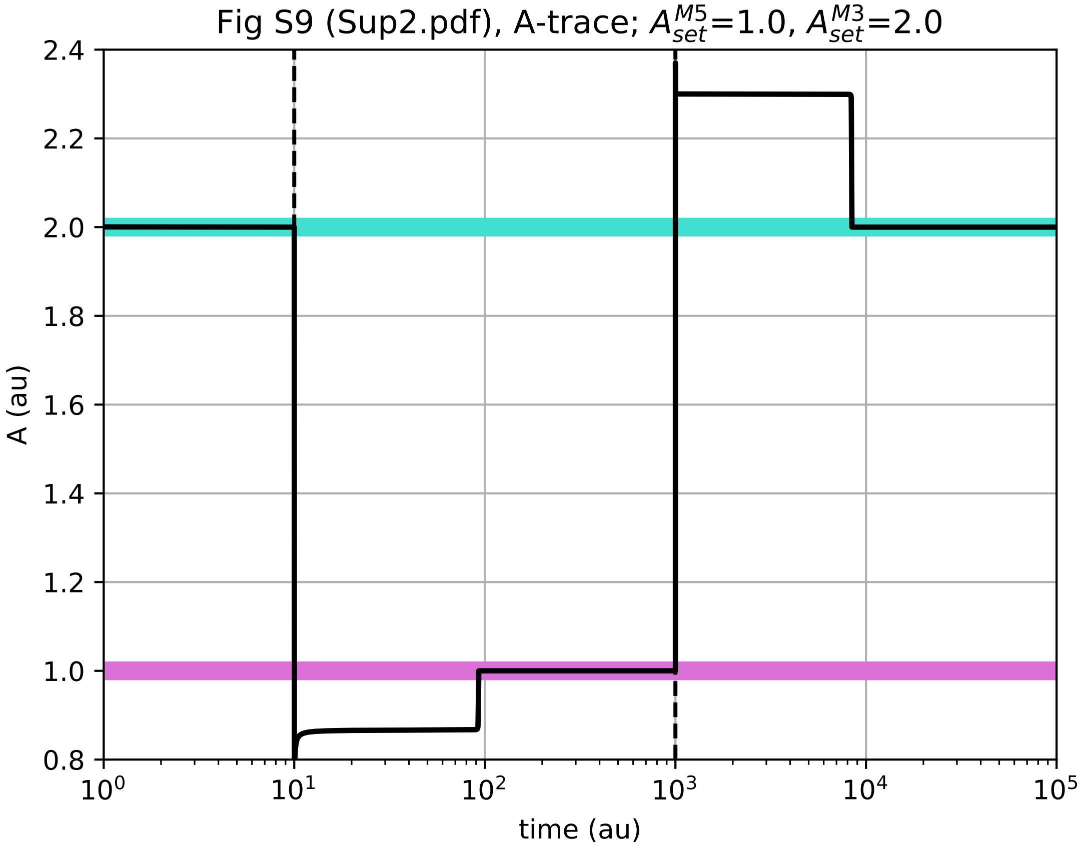

### A-time-plot_figS11.pdf

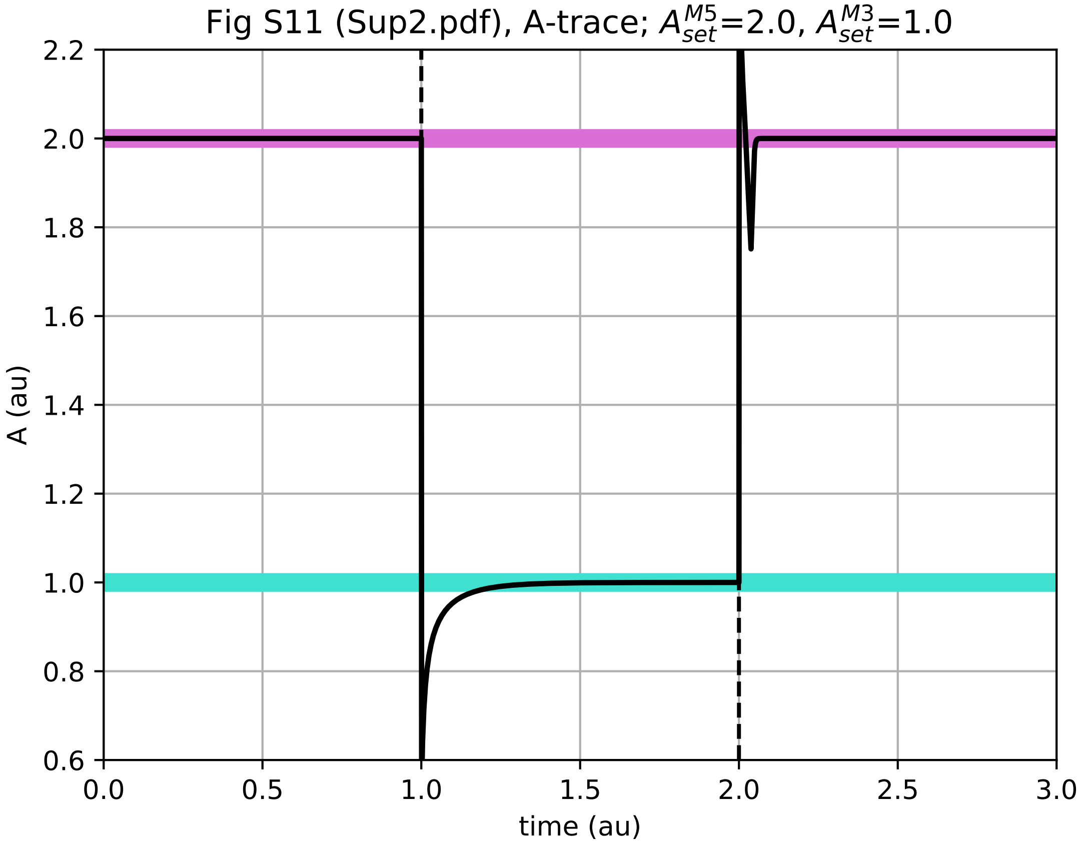

### A-time-plot_figS14.pdf

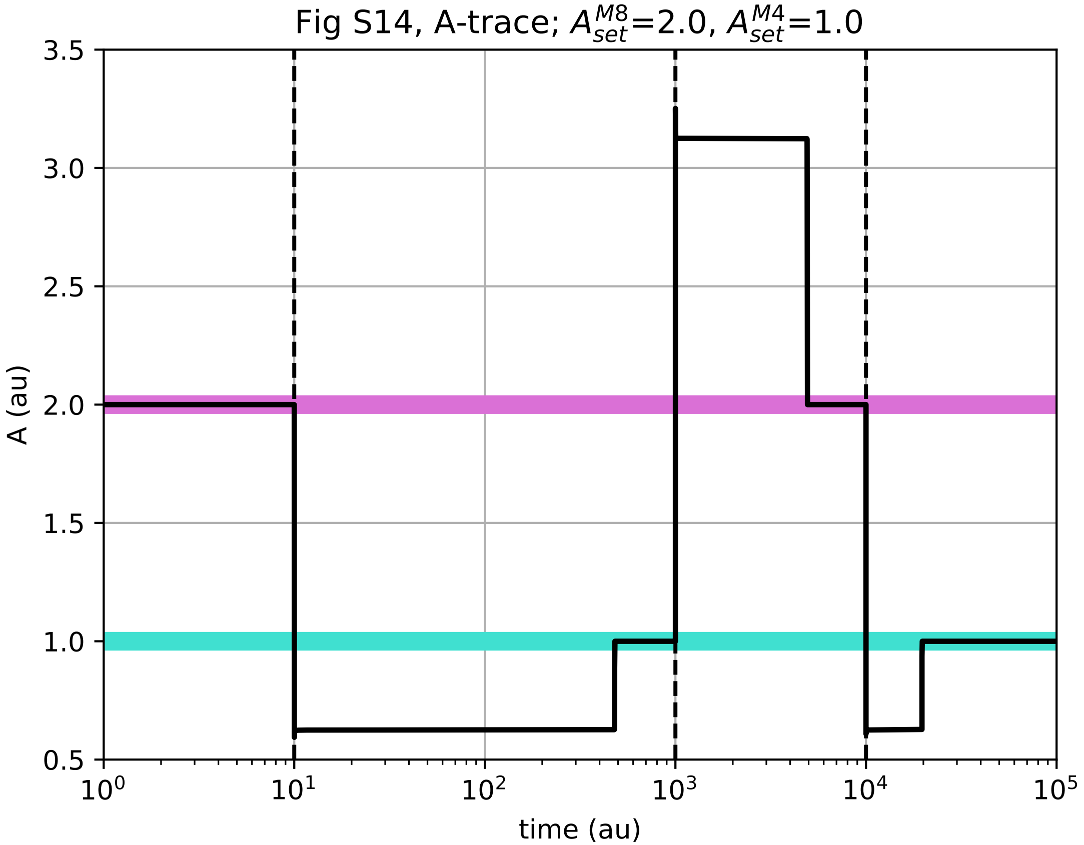

### A-time-plot_figS16.pdf

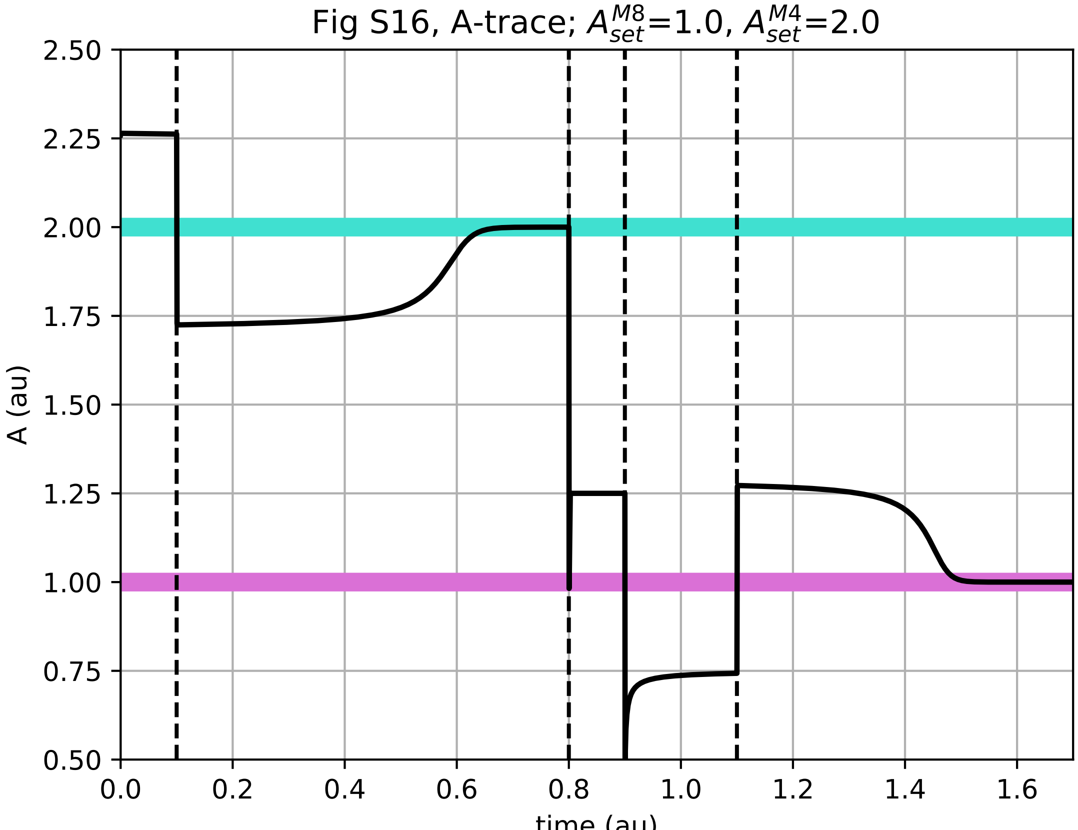

### E1-time-plot_fig9.pdf

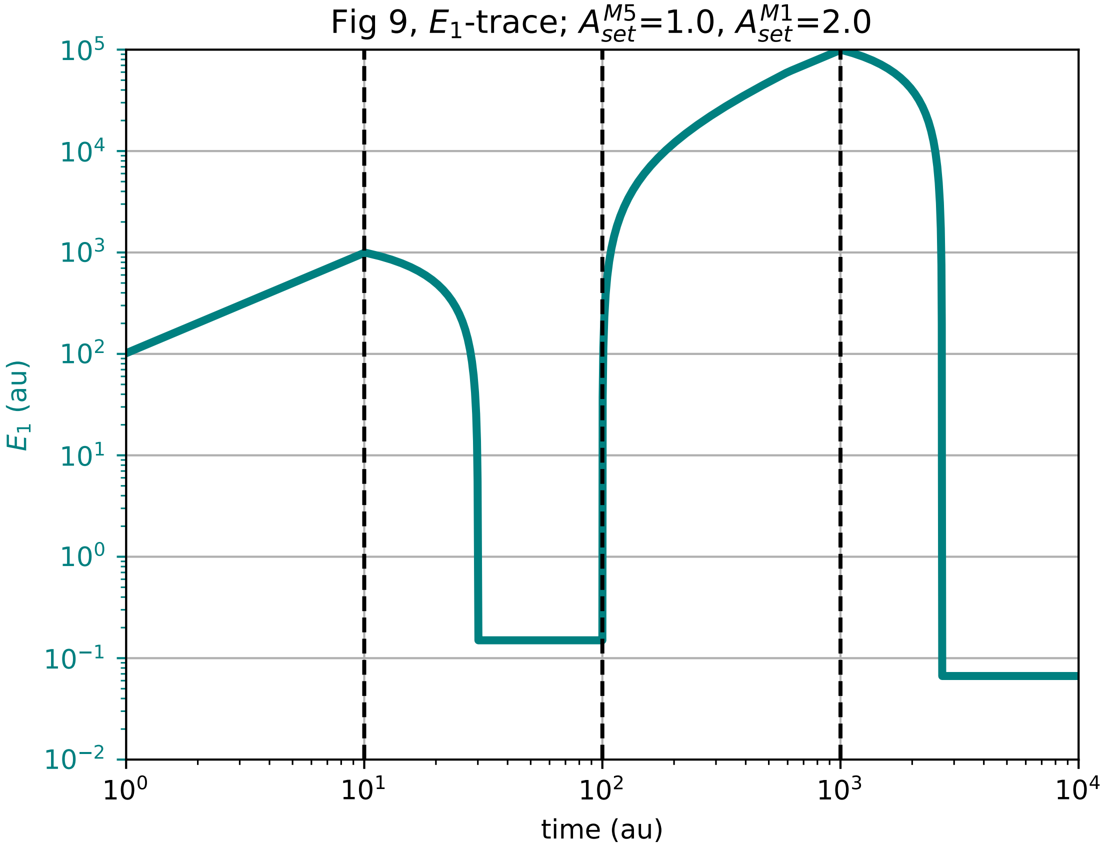

### E1-time-plot_fig21.pdf

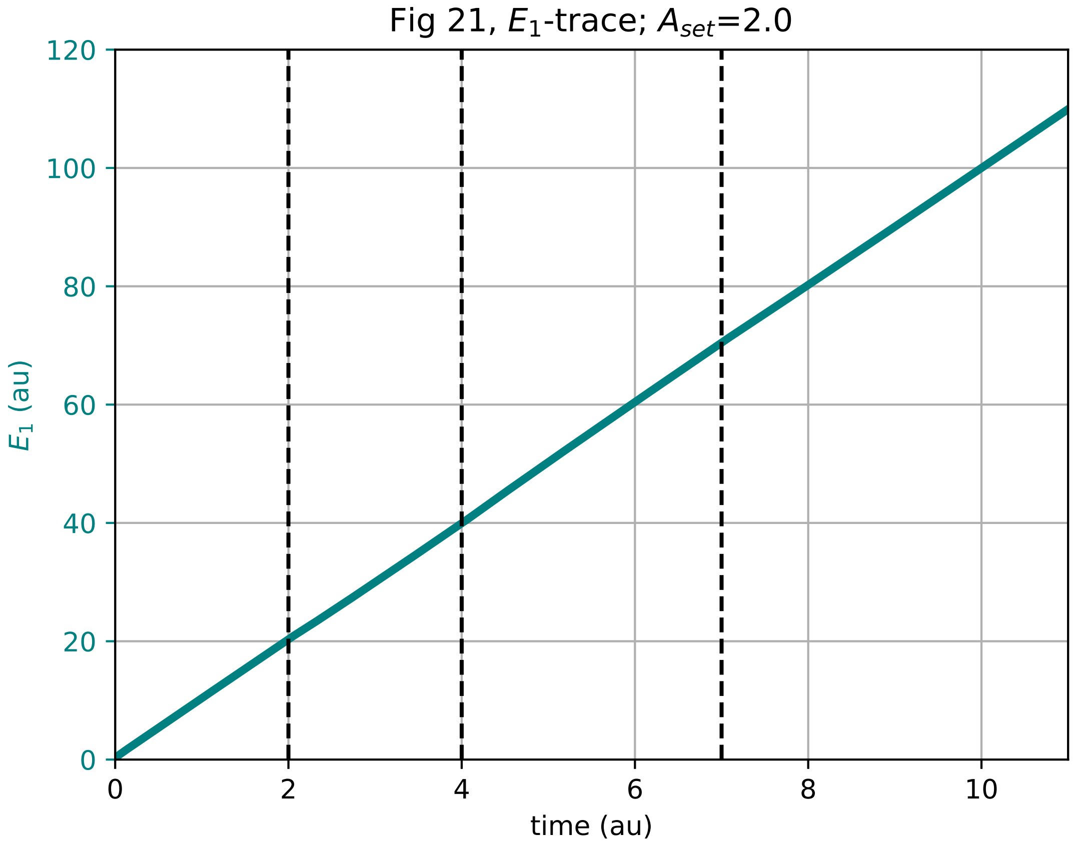

### E2-time-plot_figS3.pdf

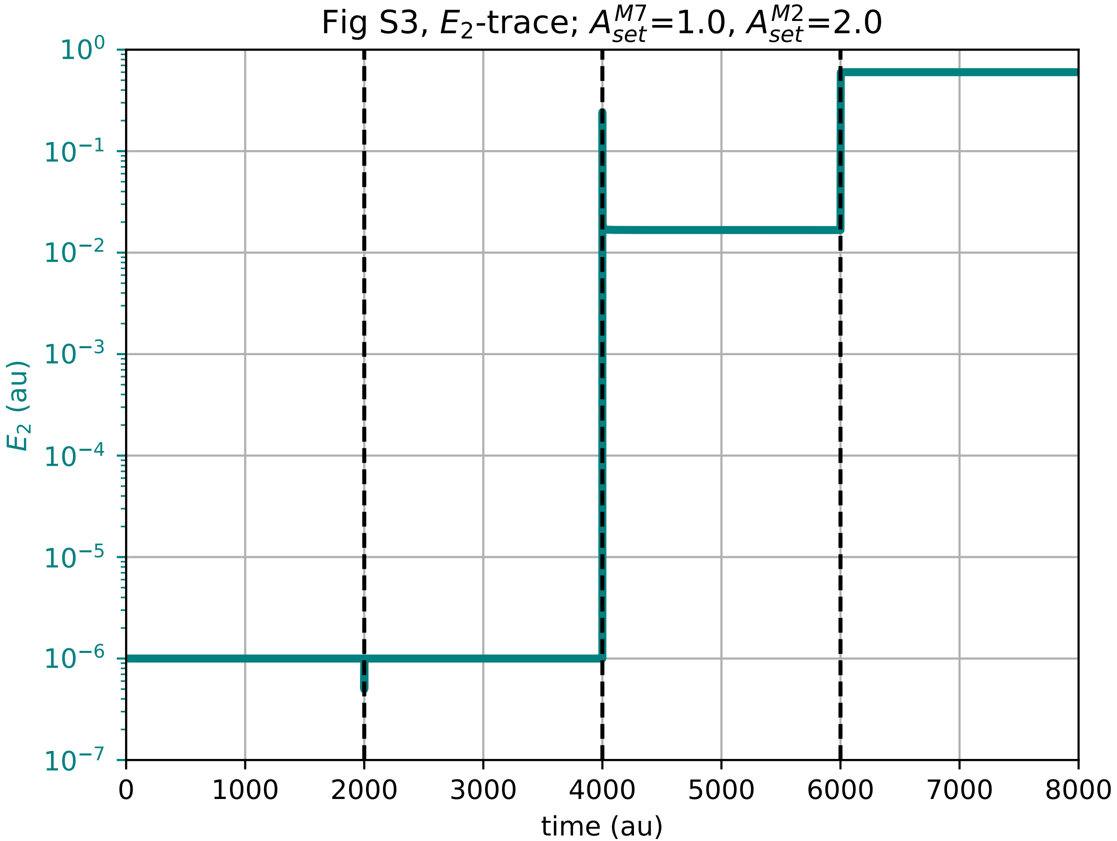

### E2-time-plot_figS6.pdf

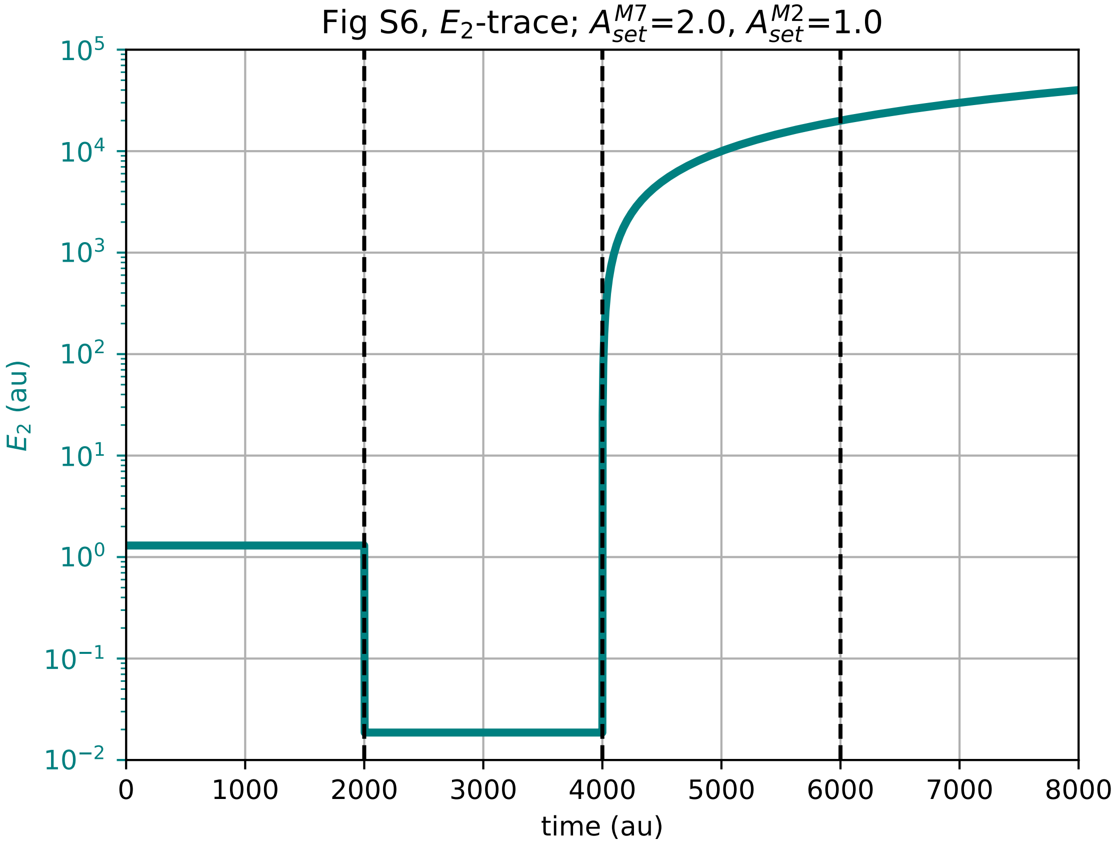

### E3-time-plot_fig26a.pdf

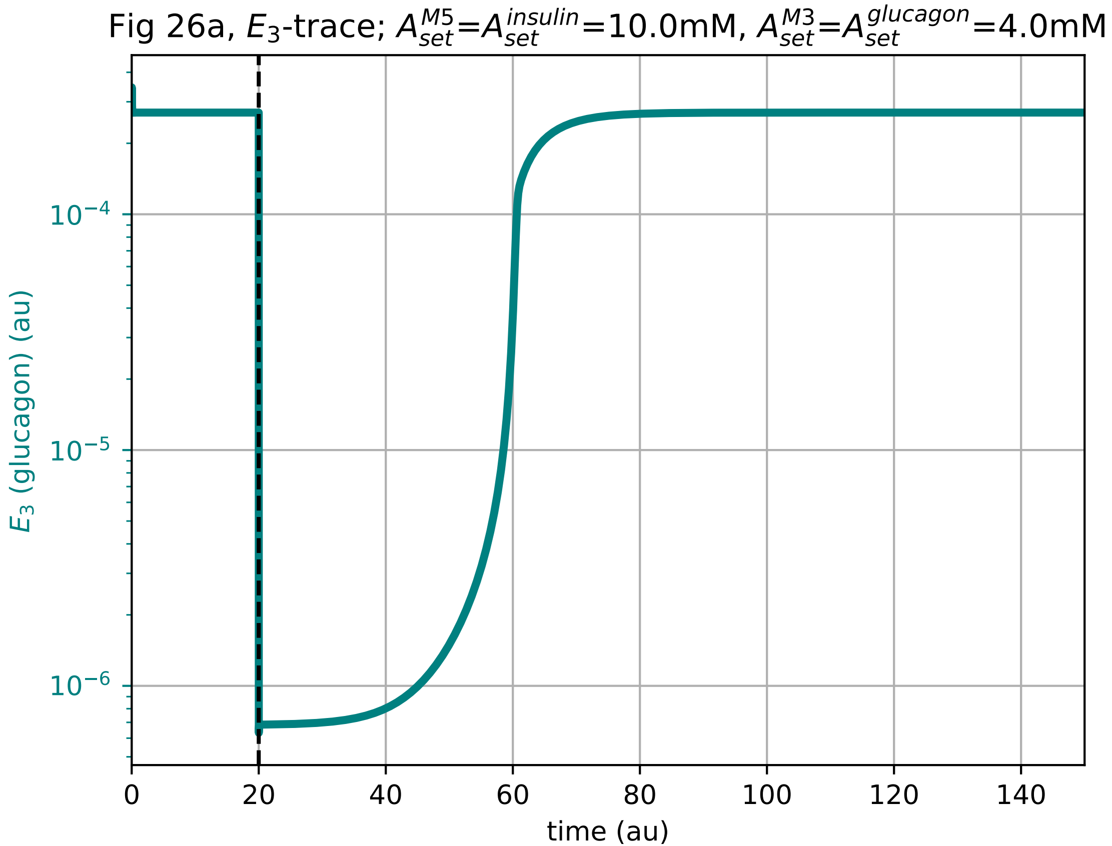

### E3-time-plot_fig26b.pdf

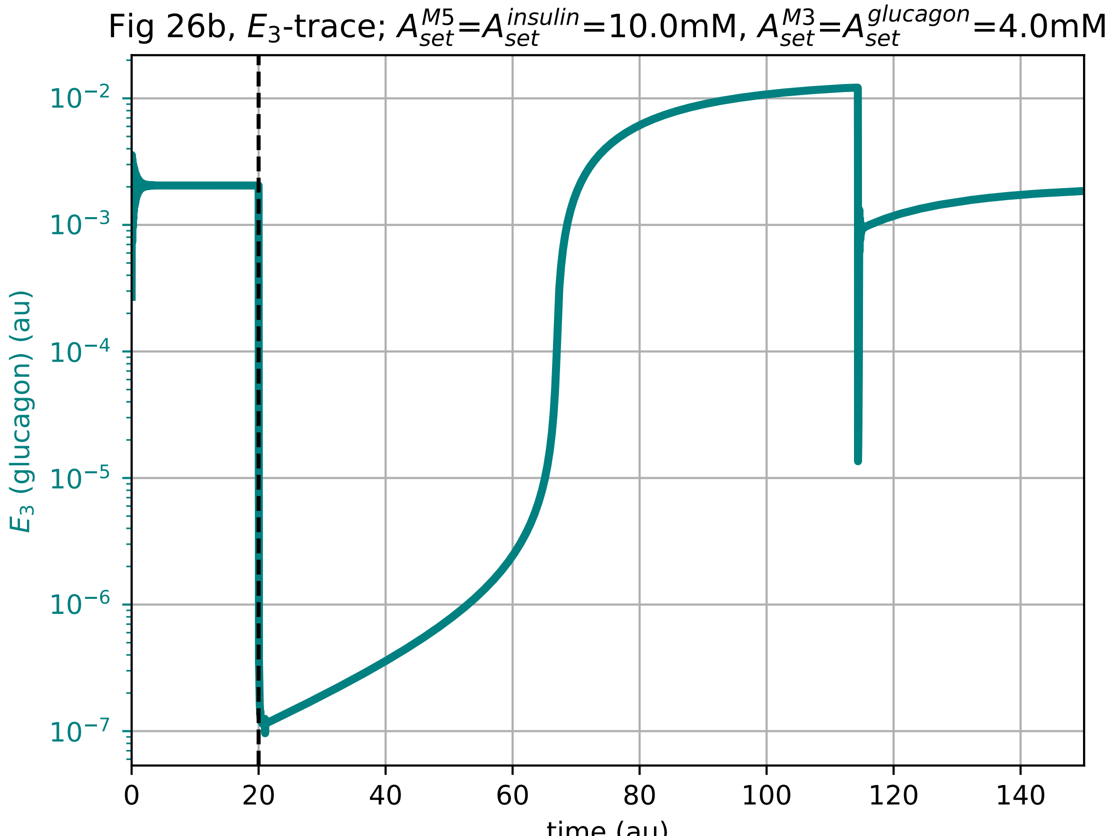

### E3-time-plot_fig26c.pdf

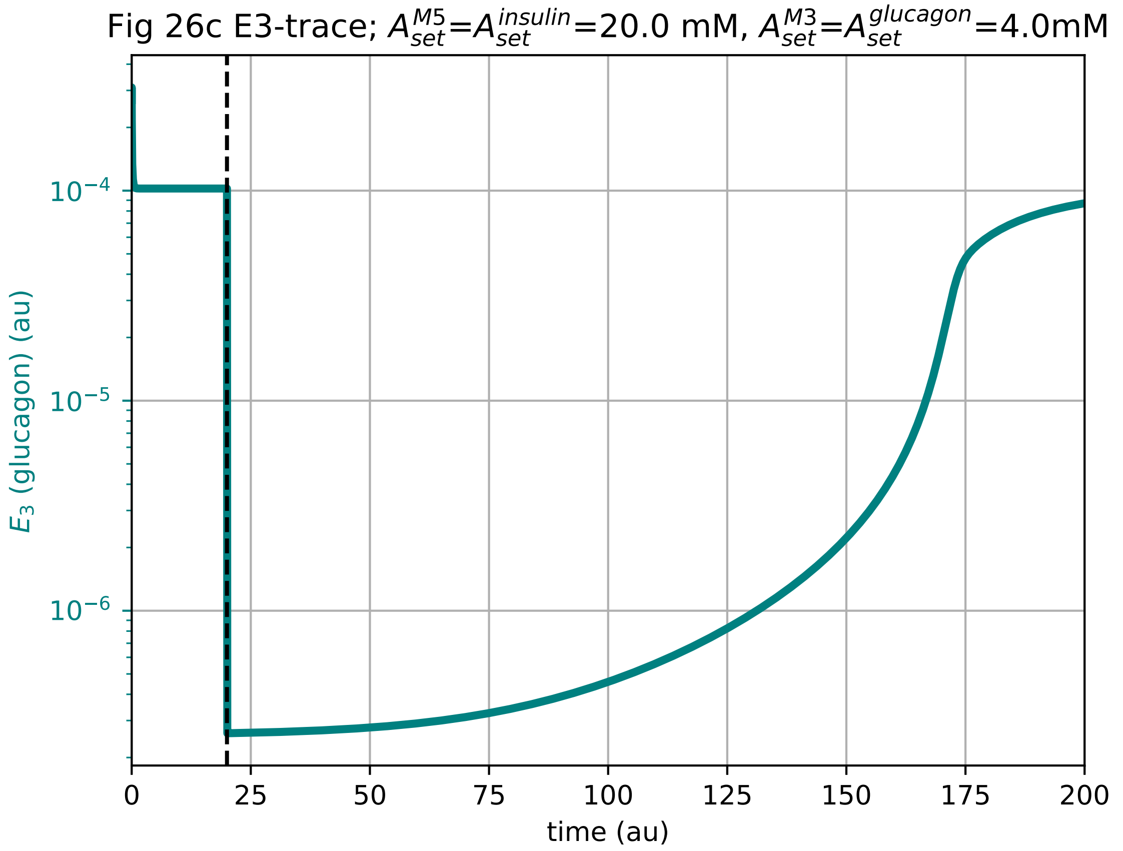

### E3-time-plot_fig26d.pdf

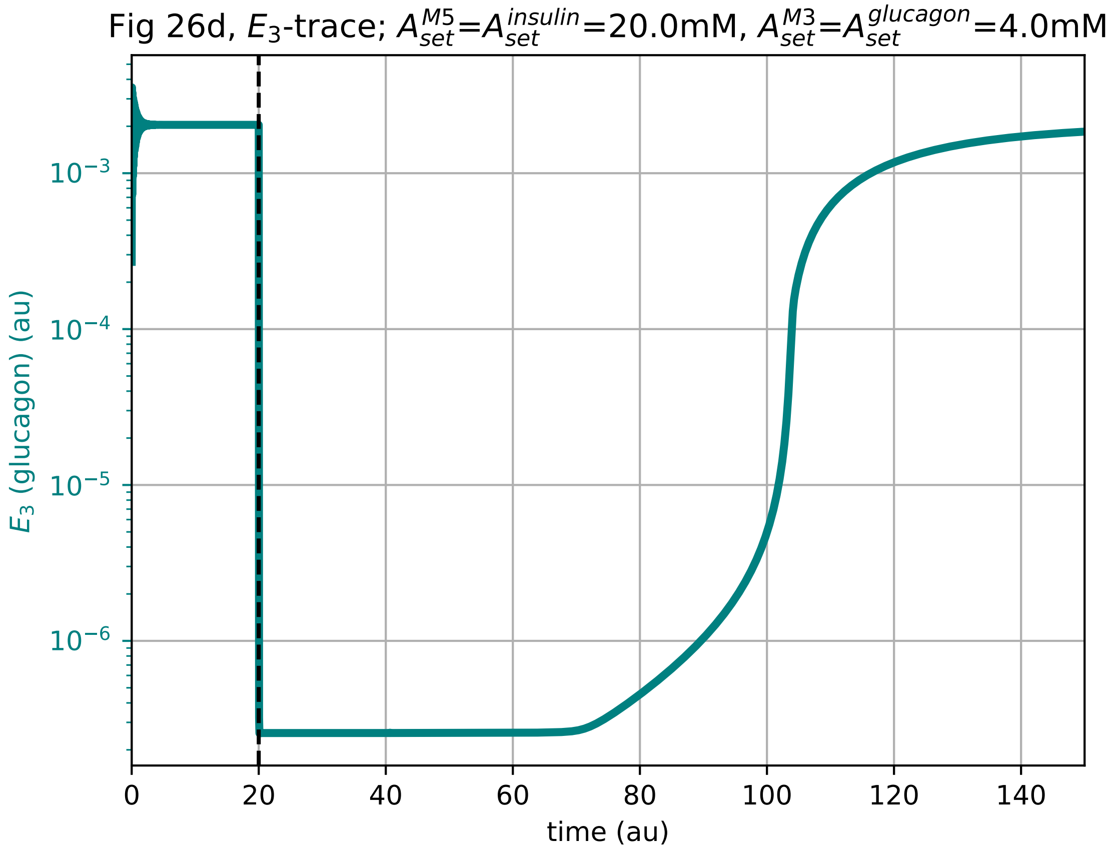

### E3-time-plot_figS9.pdf

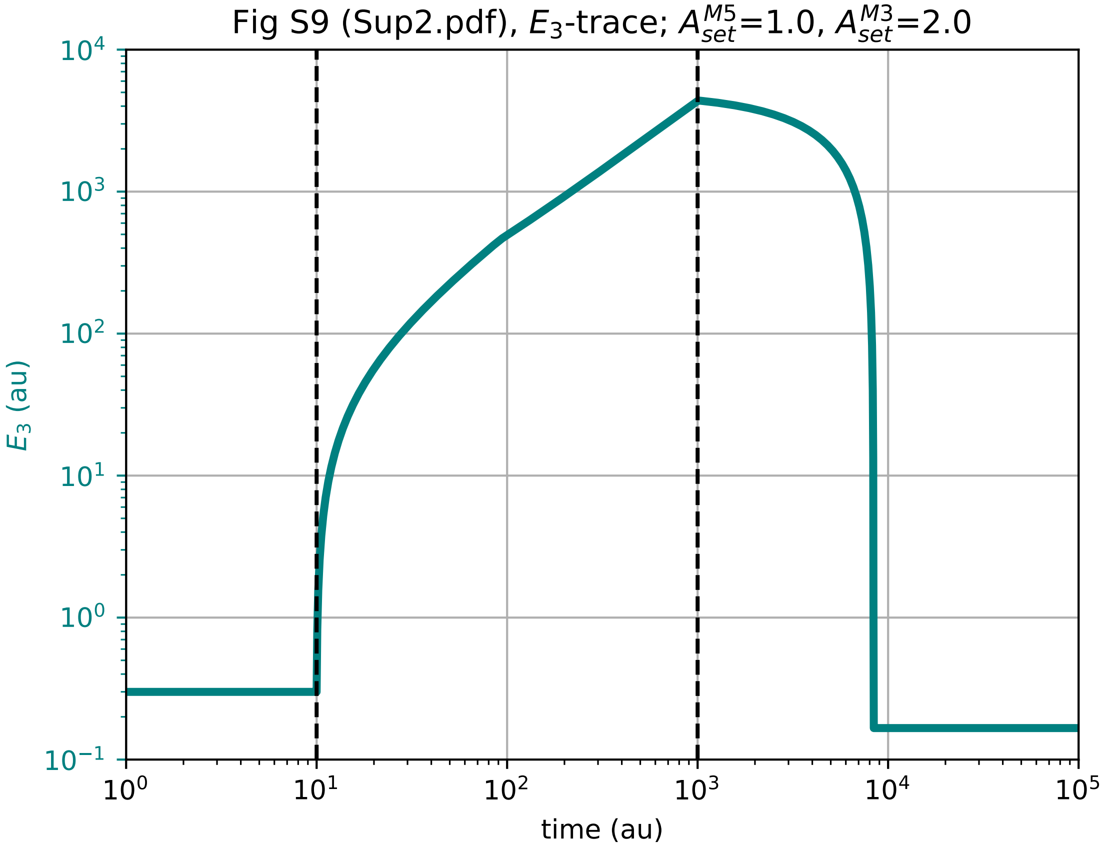

### E3-time-plot_figS11.pdf

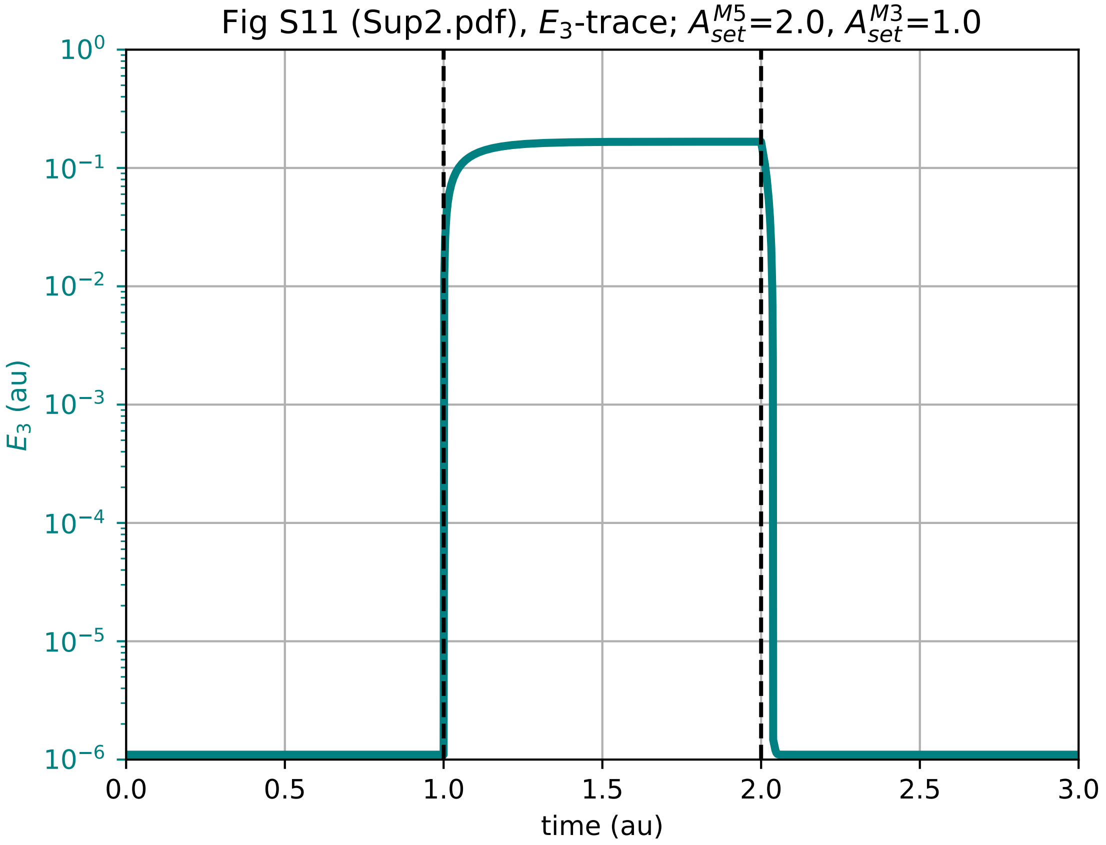

### E4-time-plot_figS14.pdf

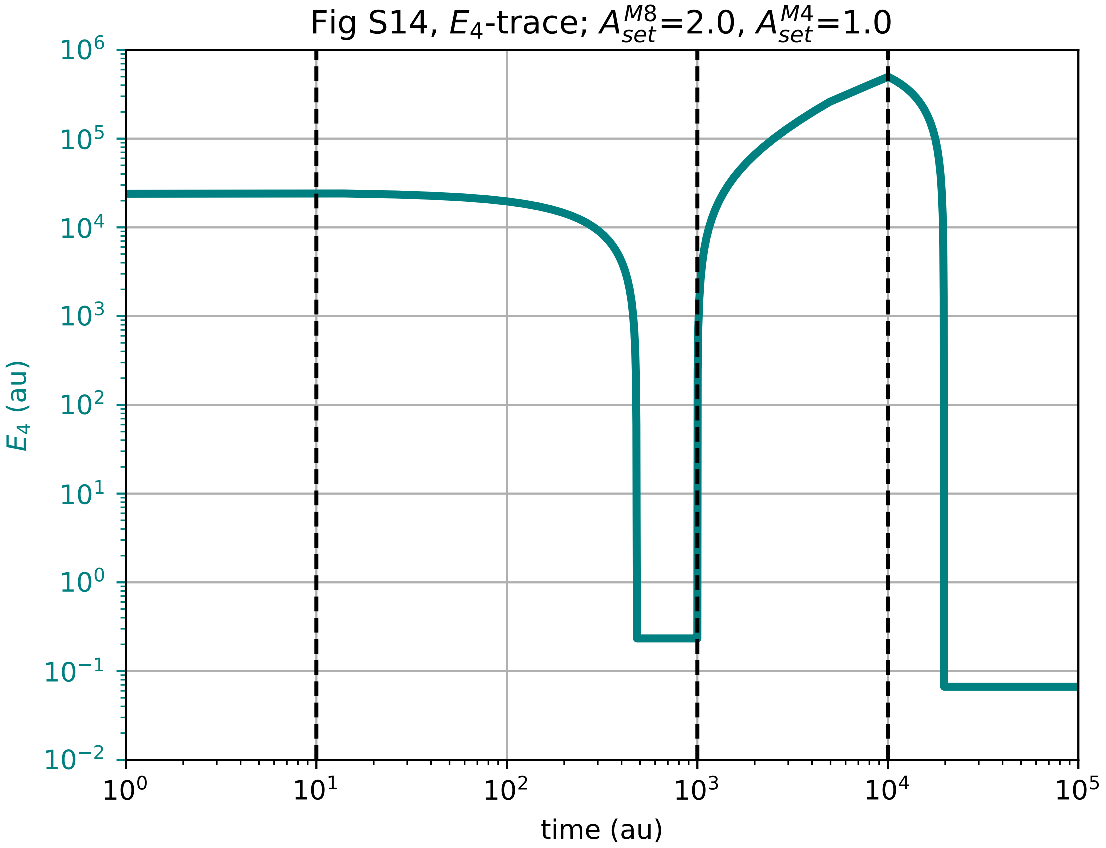

### E4-time-plot_figS16.pdf

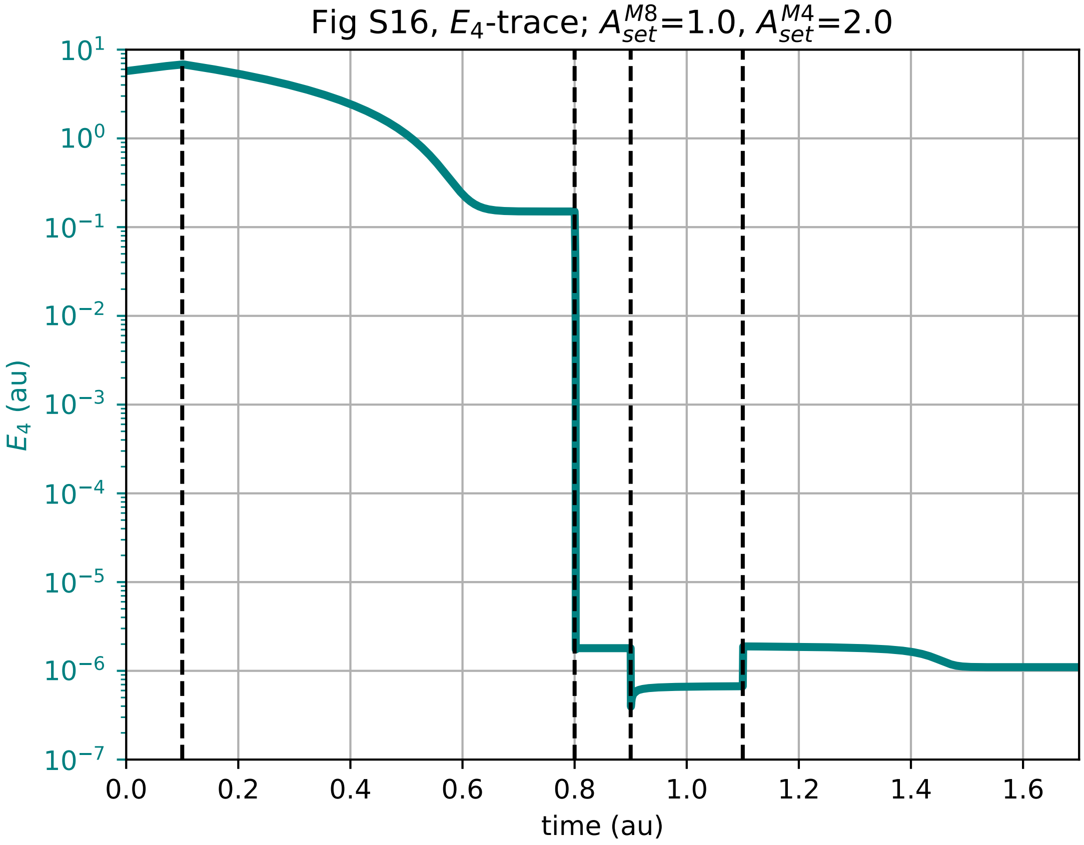

### E5-time-plot_fig9.pdf

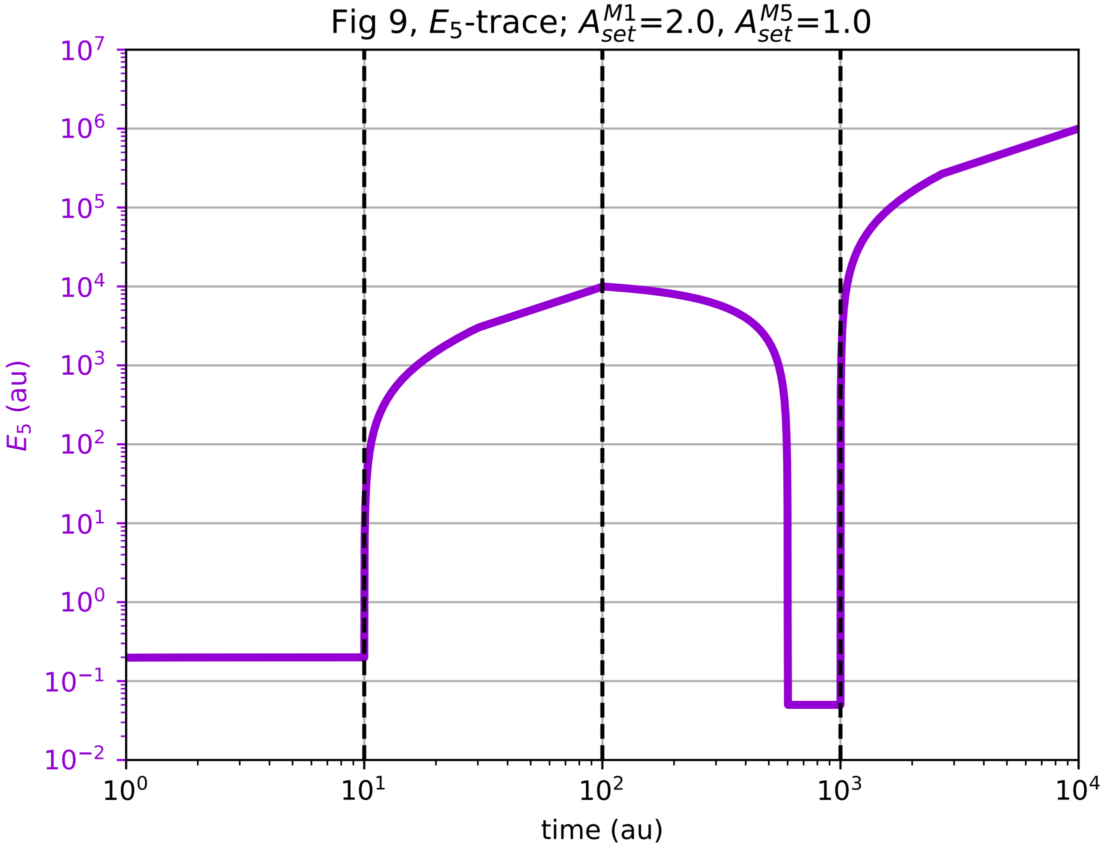

### E5-time-plot_fig21.pdf

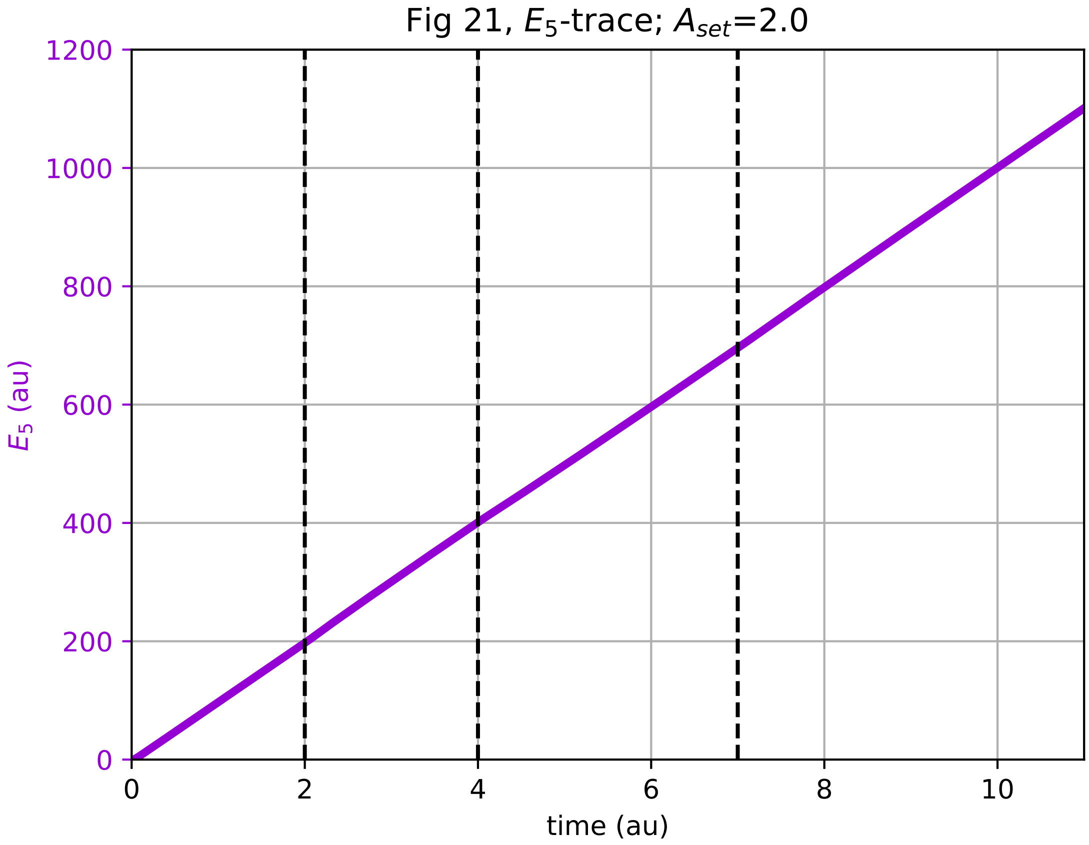

### E5-time-plot_fig26a.pdf

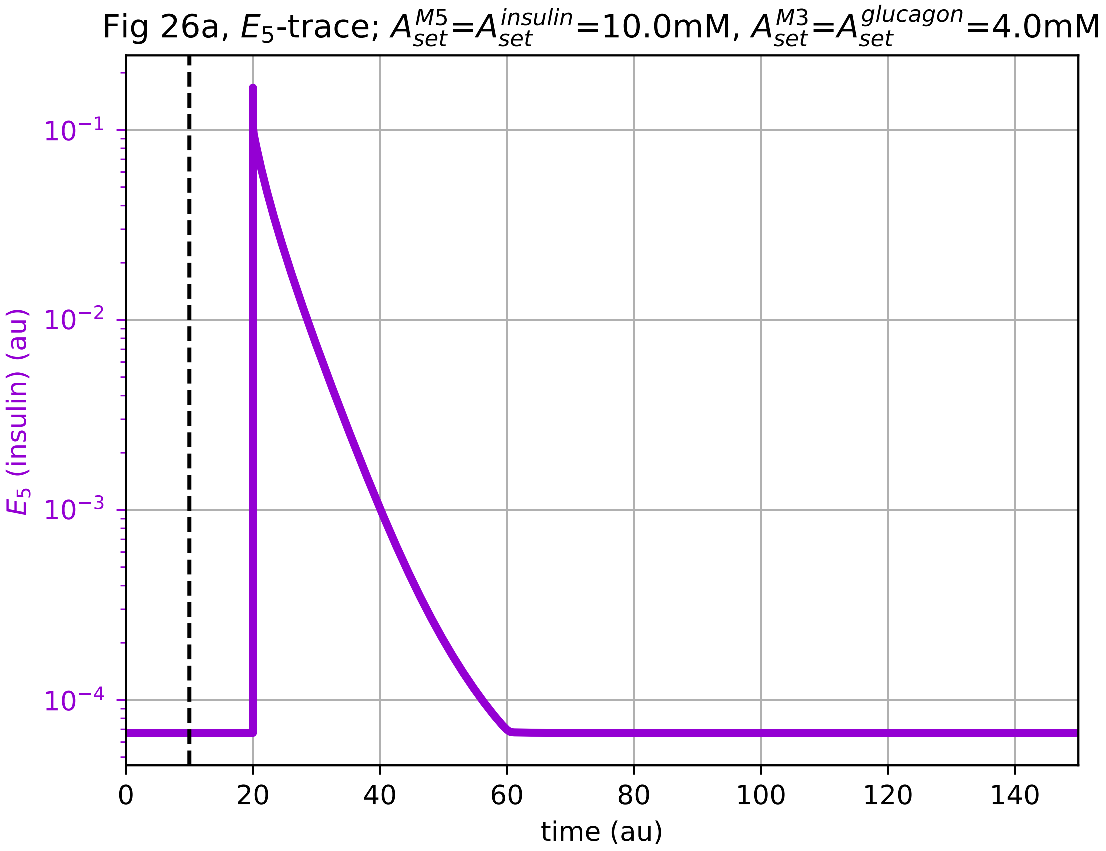

### E5-time-plot_fig26b.pdf

Fig 26b, $E_5$-trace; $A^{M5}_{set}$=$A^{insulin}_{set}$=10.0mM, $A^{M3}_{set}$=$A^{glucagon}_{set}$=4.0mM

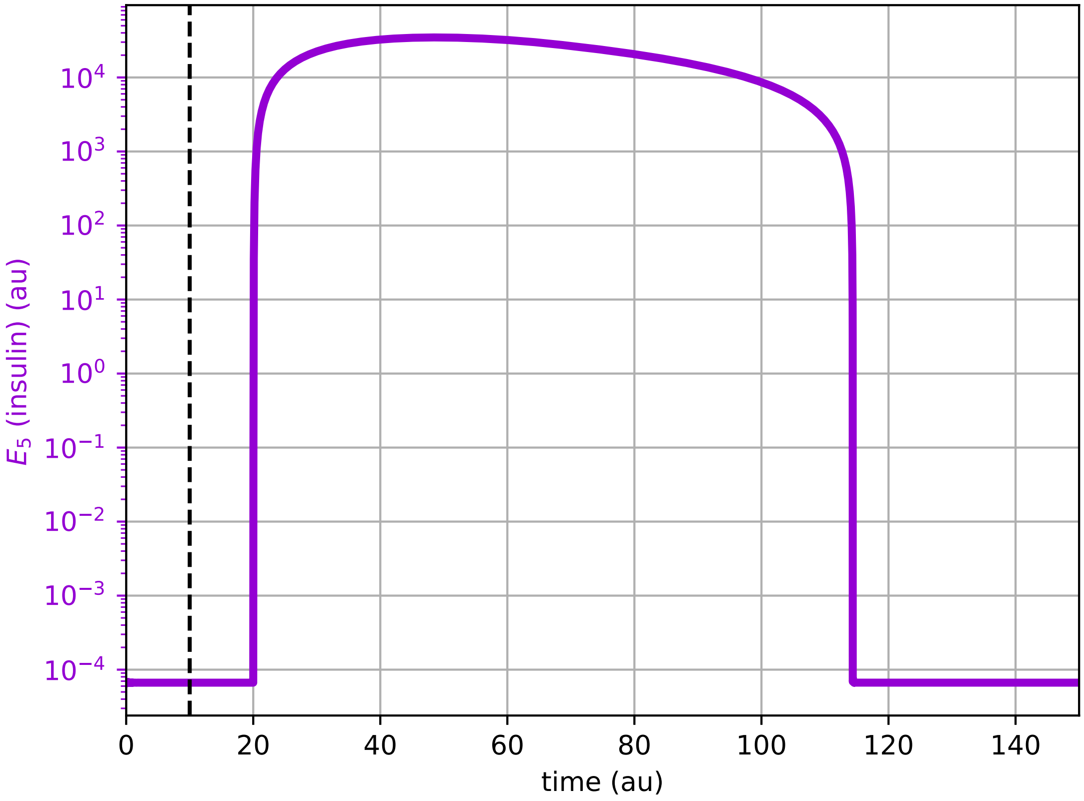

### E5-time-plot_fig26c.pdf

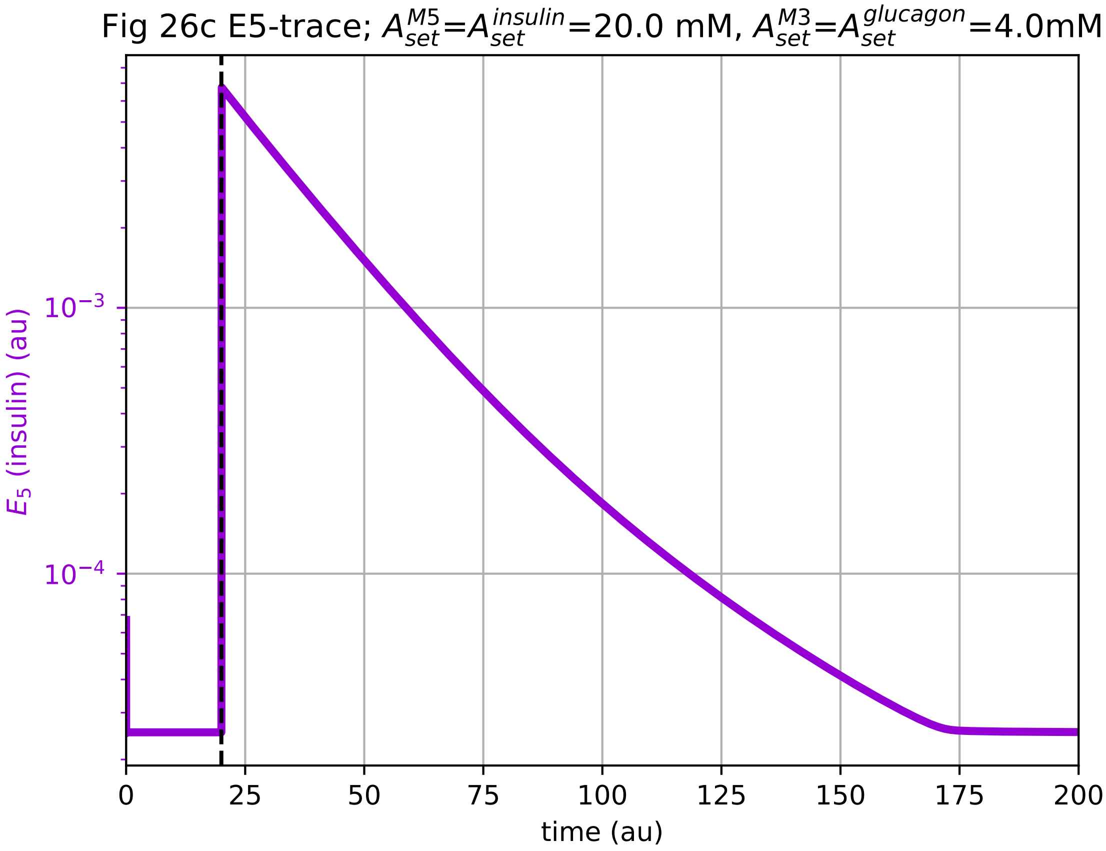

### E5-time-plot_fig26d.pdf

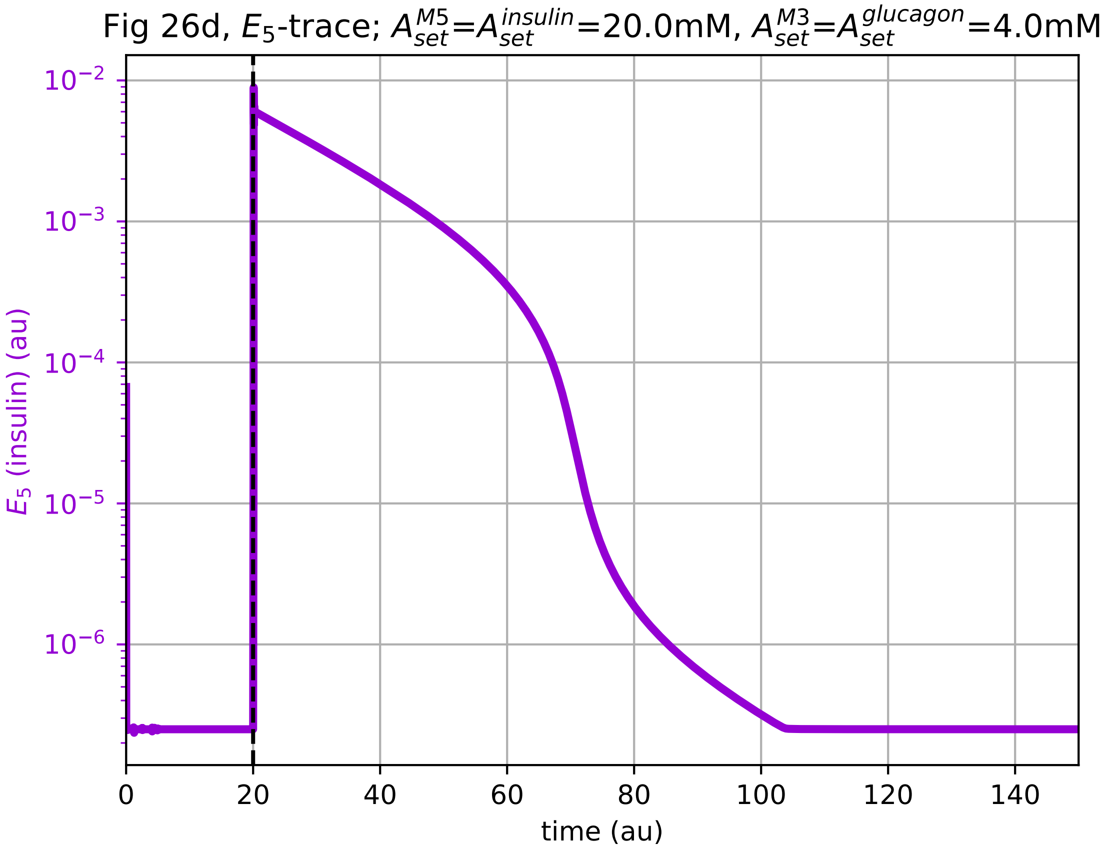

### E5-time-plot_figS9.pdf

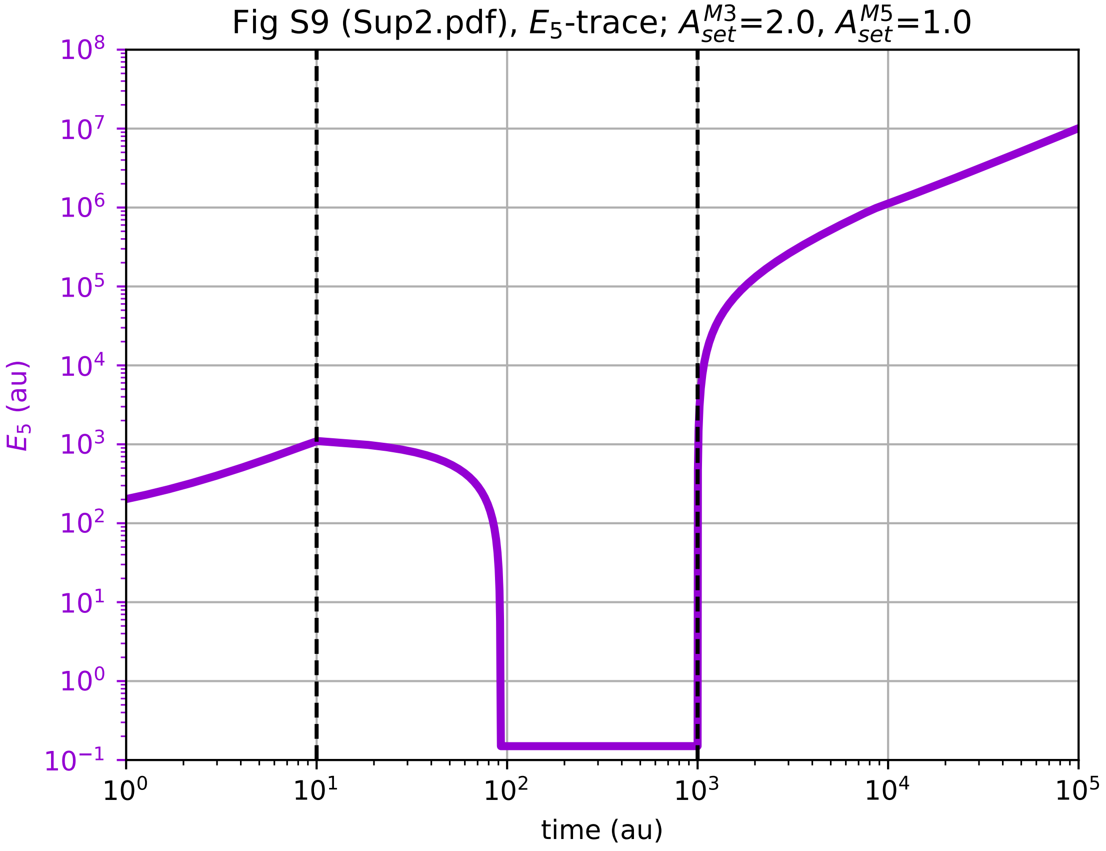

### E5-time-plot_figS11.pdf

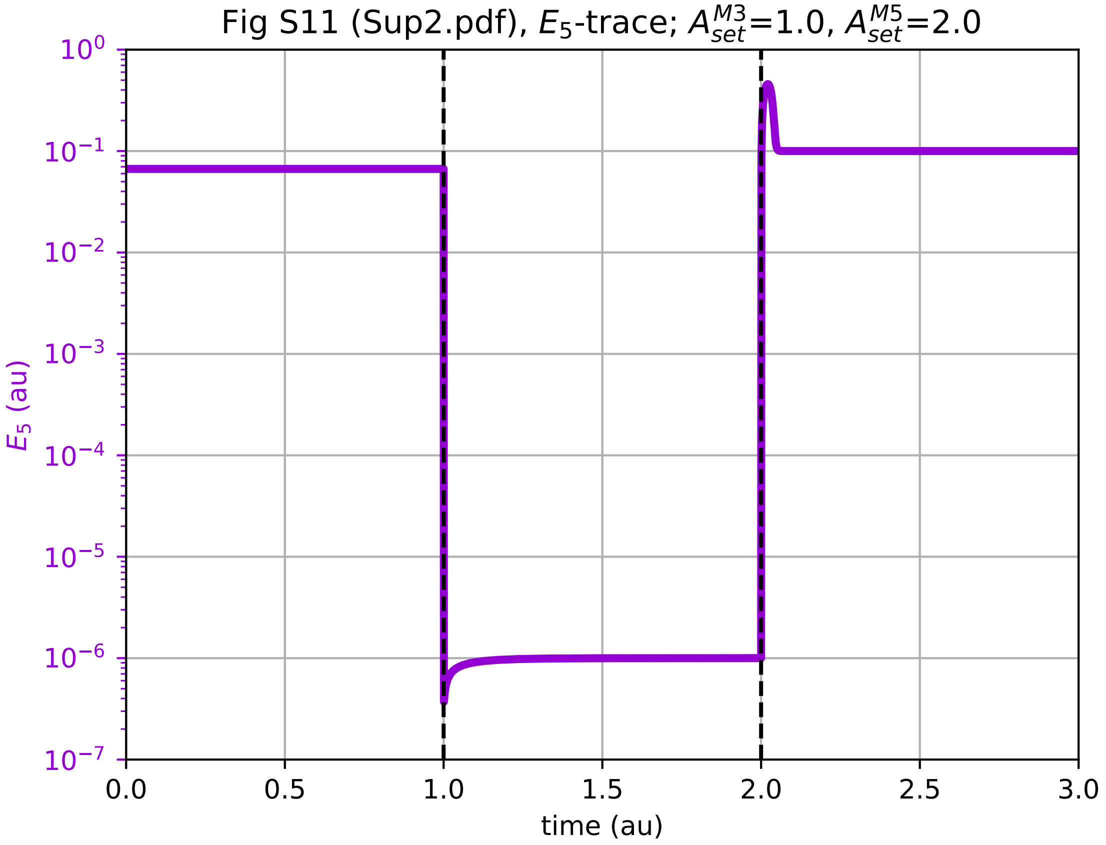

### E7-time-plot_figS3.pdf

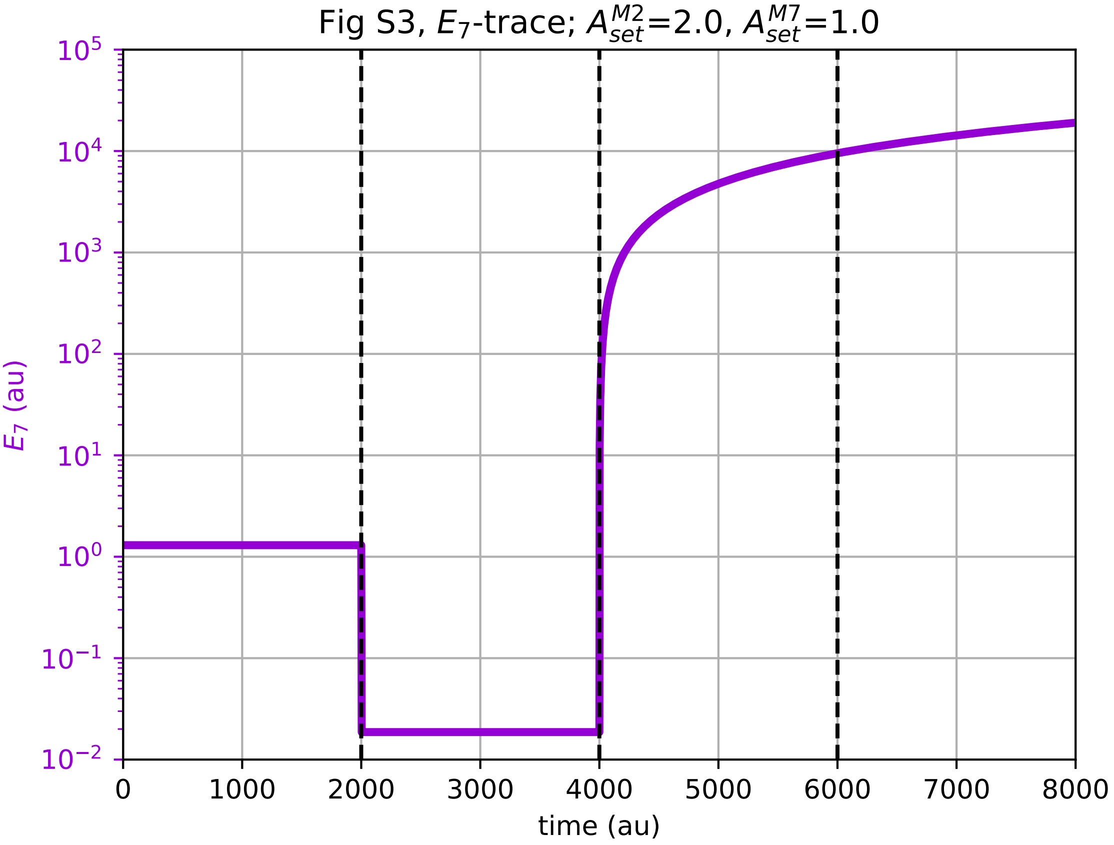

### E7-time-plot_figS6.pdf

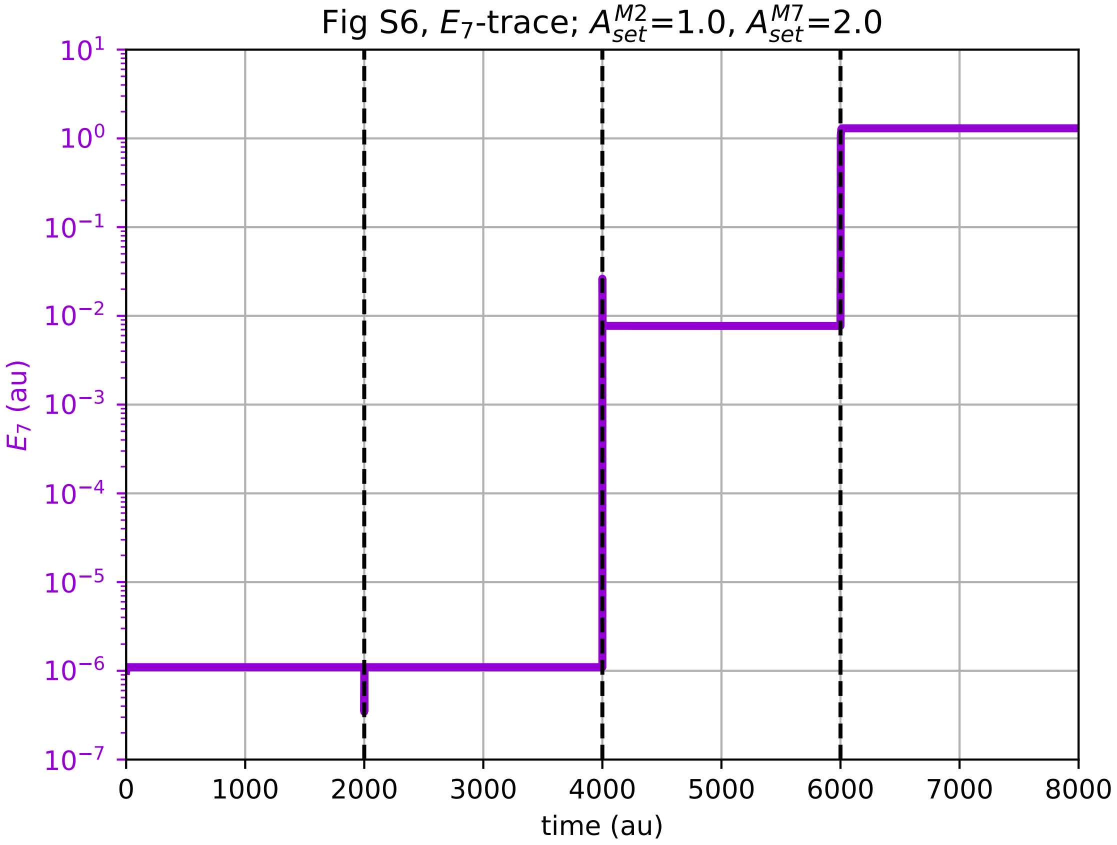

### E8-time-plot_figS14.pdf

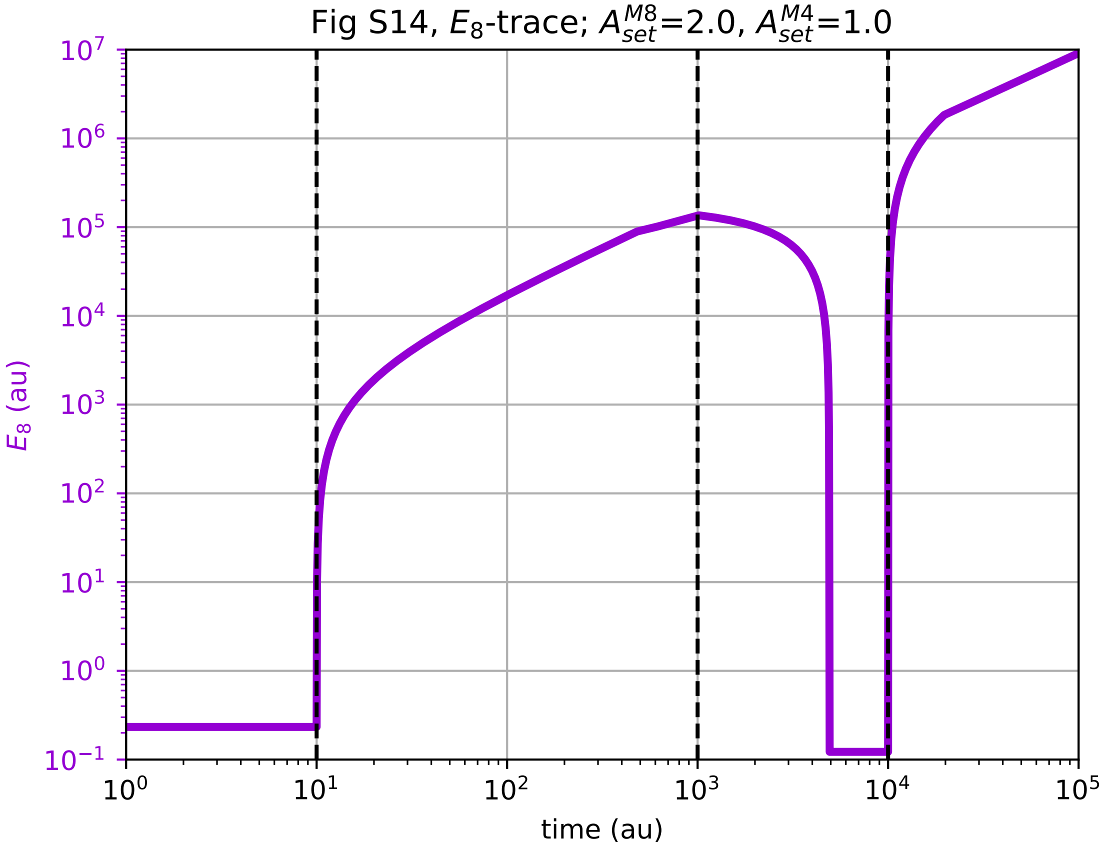

### E8-time-plot_figS16.pdf

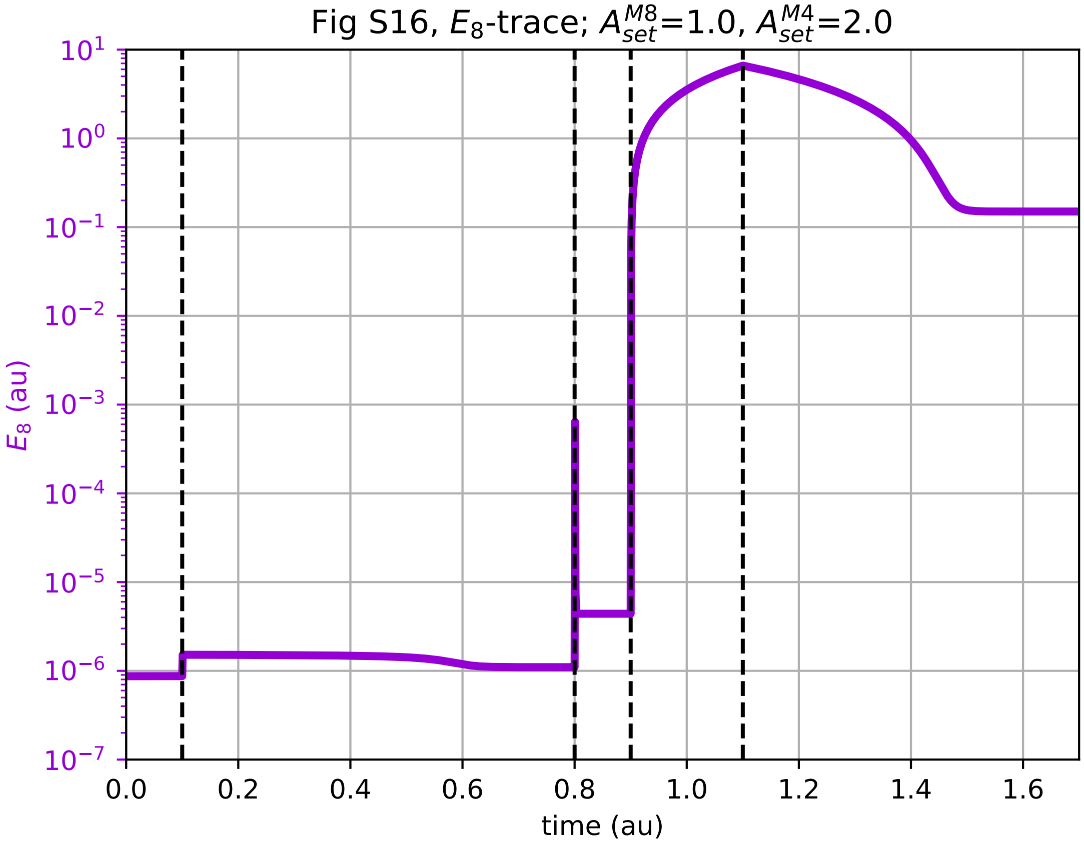

### Fig11-A-time-plot.pdf

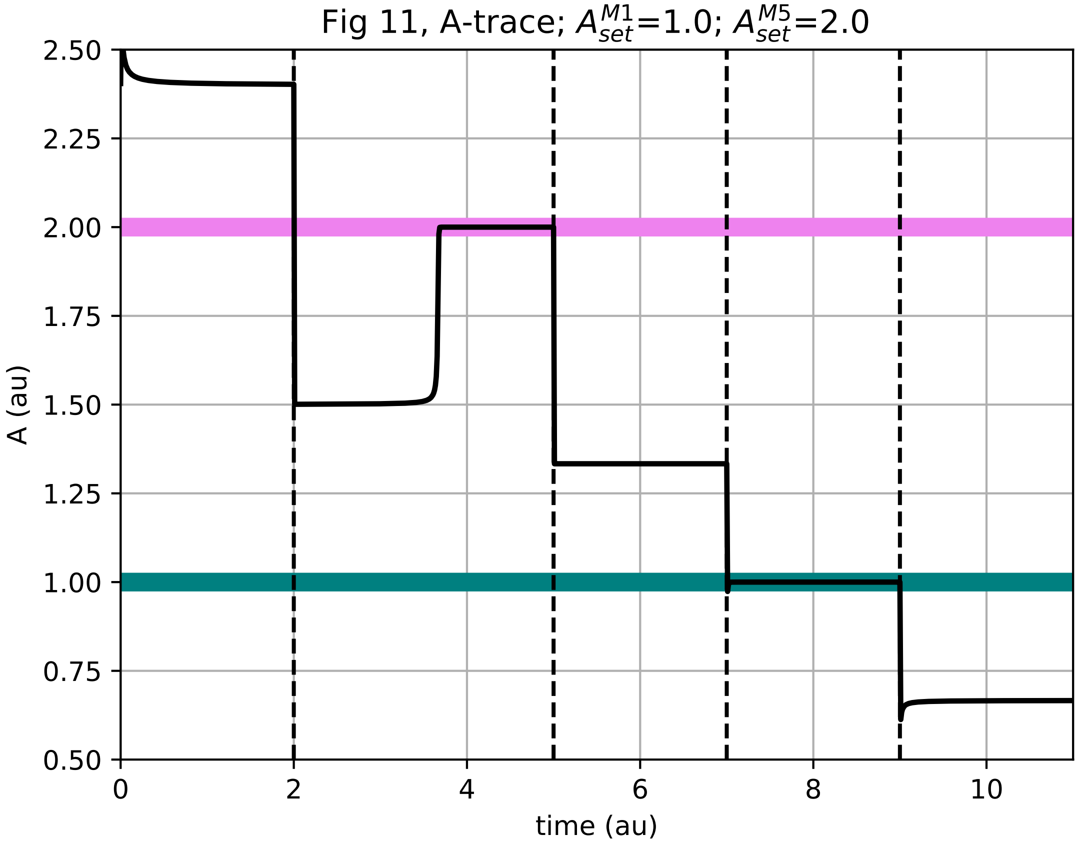

### Fig11-E1-time-plot.pdf

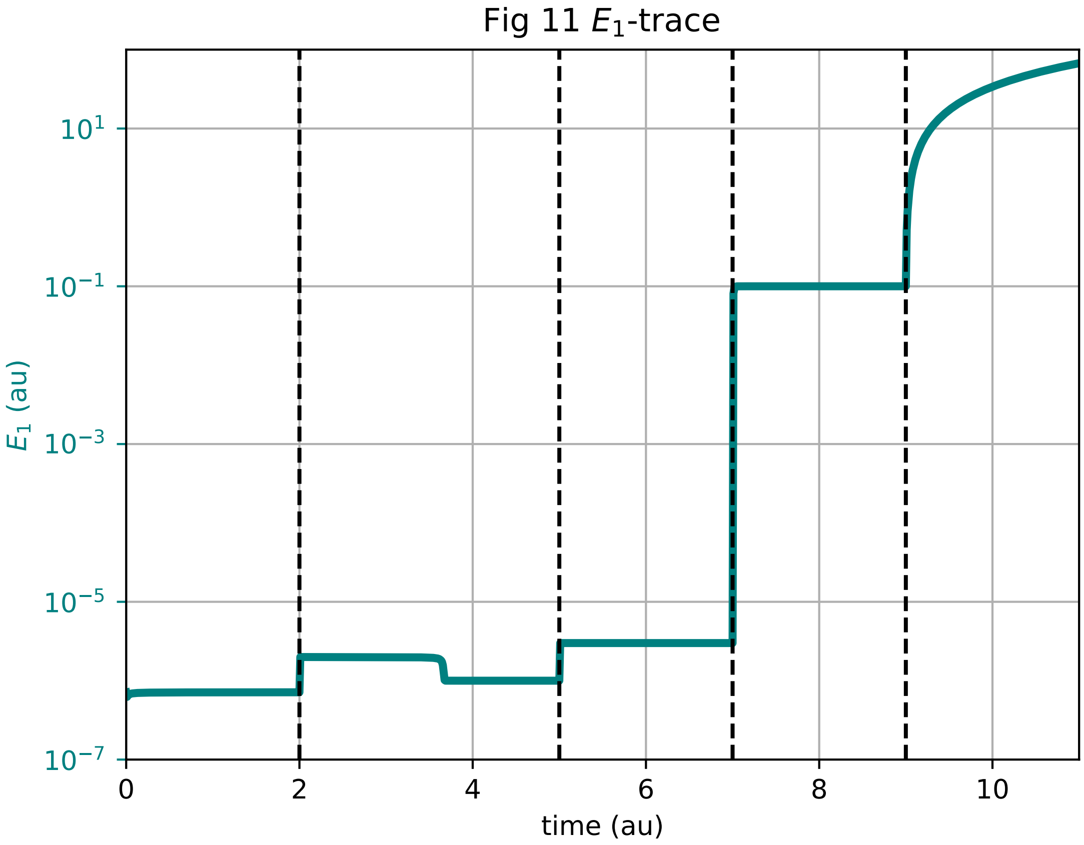

### Fig11-E5-time-plot.pdf

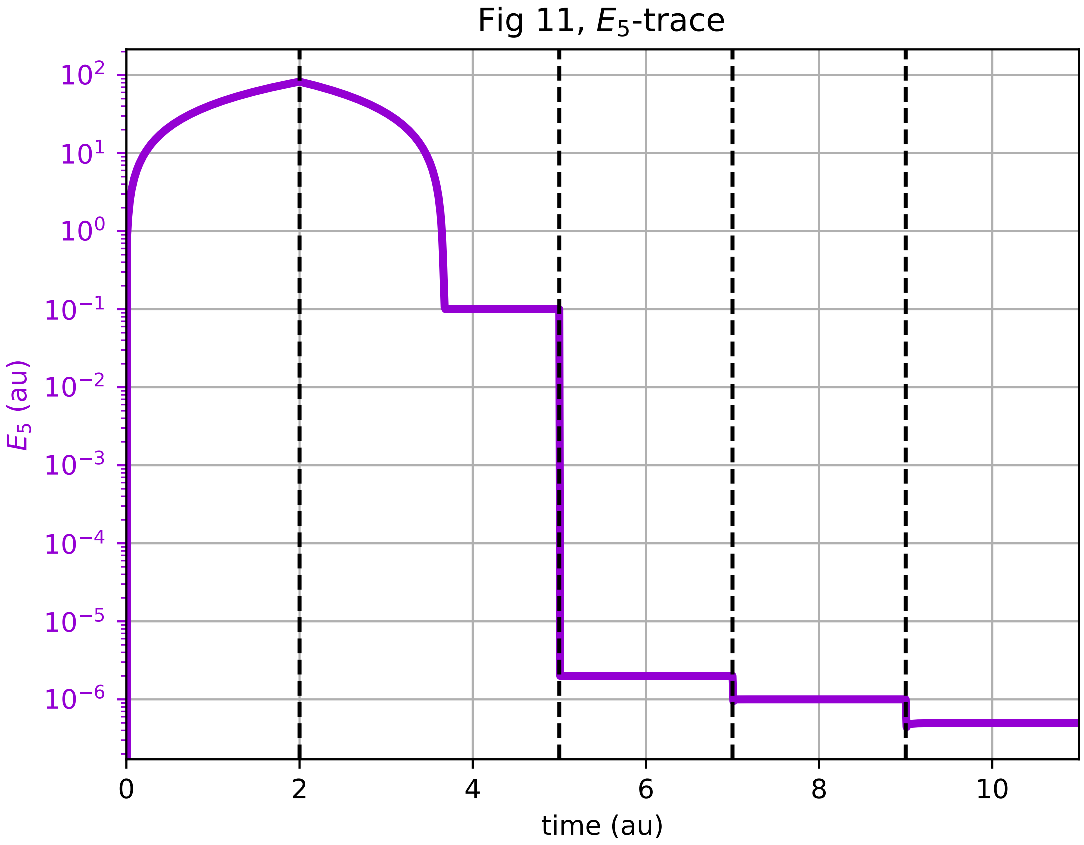

### Fig23-A-time-plot.pdf

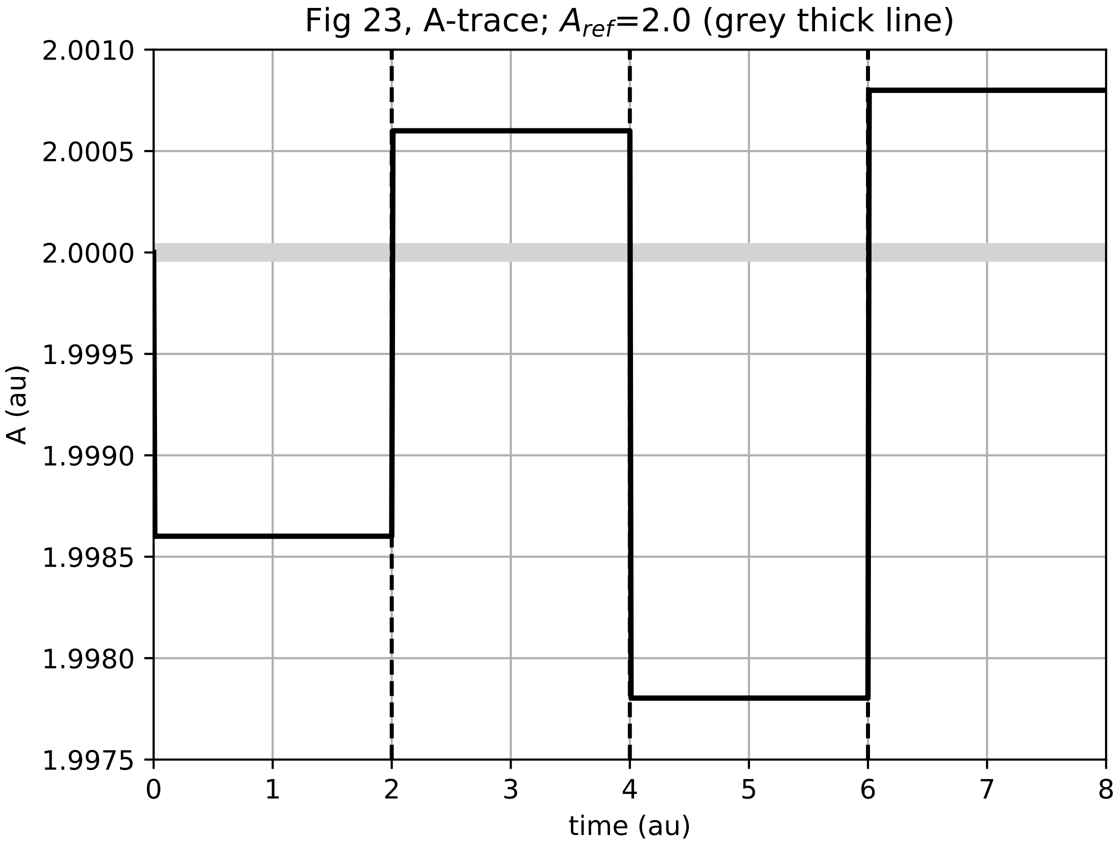
